# Adaptive structural and functional evolution of the placenta protects fetal growth in high elevation deer mice

**DOI:** 10.1101/2022.09.27.509814

**Authors:** Kathryn Wilsterman, Emily C. Moore, Rena M. Schweizer, Kirksey Cunningham, Jeffrey M. Good, Zachary A. Cheviron

## Abstract

Environmental hypoxia challenges female reproductive physiology in placental mammals, increasing rates of gestational complications. Adaptation to high elevation has limited many of these effects in humans and other mammals, offering potential insight into the developmental processes that lead to and protect against hypoxia-related gestational complications. However, our understanding of these adaptations has been hampered by a lack of experimental work linking the functional, regulatory, and genetic underpinnings of gestational development in locally-adapted populations. Here, we dissect high-elevation adaptation in the reproductive physiology of deer mice, (*Peromyscus maniculatus*), a rodent species with an exceptionally broad elevational distribution that has emerged as a model for hypoxia adaptation. Using experimental acclimations, we show that lowland mice experience pronounced fetal growth restriction when challenged with gestational hypoxia, while highland mice maintain normal growth by expanding the compartment of the placenta that facilitates nutrient and gas exchange between dam and fetus. We then use compartment-specific transcriptome analyses to show that adaptive structural remodeling of the placenta is coincident with widespread changes in gene expression within this same compartment. Genes associated with fetal growth in deer mice significantly overlap with genes involved in human placental development, pointing to conserved or convergent pathways underlying these processes. Finally, we overlay our results with genetic data from natural populations to identify candidate genes and genomic features that contribute to these placental adaptations. Collectively, these experiments advance our understanding of adaptation to hypoxic environments by revealing physiological and genetic mechanisms that shape fetal growth trajectories under maternal hypoxia.

**Significance Statement:** Residence at high elevations is associated with higher risk pregnancies and low birth weight, yet the causal mechanisms remain poorly understood. Using a high elevation-adapted rodent model, we investigated the physiological traits that explain fetal growth trajectories in low oxygen environments, and how evolutionary adaptation has modified these traits. We showed that high- and low-elevation populations of deer mice differ in their susceptibility to fetal growth restriction during gestational hypoxia and that these population-level differences are associated with structural and transcriptomic changes in the placenta. We further link placental gene expression to genomic features under selection at high elevation. Our findings identify adaptations that are likely relevant to offsetting the effects of hypoxia on fetal and placental development across mammals.

---

High elevation (> 2500 m) environments fundamentally challenge terrestrial life through an inescapable and pervasive reduction in oxygen availability. Low oxygen directly limits individual performance, and common physiological responses to low oxygen can further exacerbate these performance decrements (1–3). Low oxygen also compromises reproduction (4–6); in humans, residence at high elevations is associated with reduced birth weight and increased risks for birth complications, including intra-uterine growth restriction (4). Compromised reproductive function should have significant consequences for populations because reproductive outcomes are the ultimate arbiters of Darwinian fitness. Indeed, humans with altitude-adapted ancestry (e.g., Tibetan and Quechua peoples) experience reduced risk for these complications (7), presumably reflecting local adaptation in reproductive physiology.

Comparative analyses within and between species adapted to high elevations can provide clues about how the challenges of hypoxia are surmounted by adaptive evolution. Such analyses are common in comparative physiology, but these analyses have rarely considered prenatal reproductive physiology outside of humans (5, 6). In contrast to well-studied cardiopulmonary and metabolic adaptations to hypoxia, the reproductive physiology that influences fetal growth outcomes at high elevations remains poorly understood. To date, maternal traits, including ventilatory characteristics, uterine artery diameter, and microvascular structure in the placenta, have been hypothesized to contribute to fetal growth restriction (and protection thereof at elevation) (5, 7). However, mechanistic links between these traits and fetal growth remain limited.

The deer mouse, *Peromyscus maniculatus*, is a promising model system for investigating how adaptive evolution has mediated the fundamental challenges hypoxia poses to gestational physiology. Several aspects of deer mouse biology make them a useful comparative model for such questions. First, deer mice are a well-established model for studying adaptive evolution (8), including adaptation to high elevation environments (reviewed in (9–11)), though prenatal reproductive adaptations have not yet been explored in this system (6). Second, genetic differentiation and variation among wild deer mouse populations is well-characterized (12–14), and genetic signatures of local adaptation in highland deer mice persist in the face of high rates of gene flow (12, 15). The low genome-wide genetic differentiation among deer mouse populations allows for fine-scale resolution of genomic regions that have experienced a history of natural selection at high elevation, which can be informative for understanding the physiological basis of adaptation (e.g., (12, 15)). Third, high elevation deer mouse populations are derived from lowland populations (13), which mirrors the biogeographic history of many highland human populations (i.e., lowland individuals moving into and adapting to highland environments). Related, deer mice and humans share fundamental aspects of placental structure (16, 17), and many of the maternal traits that are adjusted to support pregnancy in humans are also remodeled in deer mice (16). Together, the similarities in biogeographic history and reproductive physiology provide an opportunity to identify conserved or convergent solutions to the challenges that high elevation places on reproductive physiology.

Here, we used hypoxia acclimation experiments to link population-specific reproductive outcomes to subordinate physiological traits. We further investigated transcriptomic variation in the placenta, linking expression patterns to fetal growth and population-specific hypoxia responses, and we asked whether these transcriptomic signatures were associated with genomic targets of local adaptation (12, 14, 15). Our analyses identify mechanisms by which placental physiology and maternal hypoxia interact to influence fetal growth, and they highlight both new and established genic targets that are closely tied to these developmental processes.

## Results and Discussion

### Fetal growth under maternal hypoxia in deer mice

If fetal growth restriction is a fundamental challenge that impacts survival and fitness of mammals at high elevation, we expected that (a) lowland-derived deer mice should give birth to smaller pups when gestating under hypobaric hypoxia (hereafter, hypoxia), and (b) highland-adapted deer mice should protect fetal growth under the same conditions. We compared birth weights of deer mouse pups born to dams gestating under normoxia to those born to dams held in hypoxia, using deer mice derived from Ann Arbor, Michigan (appx. 250 m ASL, lowlanders) and Mt. Evans, CO (4300 m ASL, highlanders; Table 1, Fig. 1A). As predicted, pups from lowland dams gestating under hypoxia were nearly 20% smaller (in mass) than their normoxia-gestated counterparts (Fig. 1B; Lowland N vs Lowland H: P<0.05, see fig. caption). Strikingly, highland dams gestating under hypoxia gave birth to pups that did not differ in mass from their normoxia-gestated counterparts (Fig. 1B; Highland N vs Highland H: P=0.99, see fig. caption), suggesting that local adaptation to high elevation environments has involved changes to reproductive physiology that prevent hypoxia-dependent fetal growth restriction.

**Table 1.**
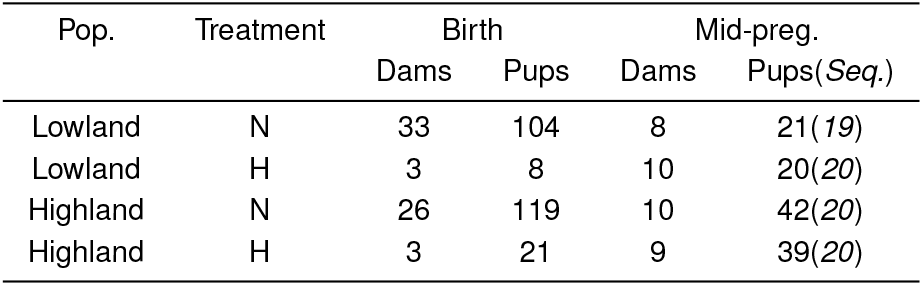
Sample sizes across populations and treatments. For mid-pregnancy, the subsets of total pups used for sequencing experiments are indicated in parentheses

| Pop. | Treatment | Birth |  | Mid-preg. |  |
| --- | --- | --- | --- | --- | --- |
|  |  | Dams | Pups | Dams | Pups(Seq.) |
| Lowland | N | 33 | 104 | 8 | 21(19) |
| Lowland | H | 3 | 8 | 10 | 20(20) |
| Highland | N | 26 | 119 | 10 | 42(20) |
| Highland | H | 3 | 21 | 9 | 39(20) |

**Fig. 1.**
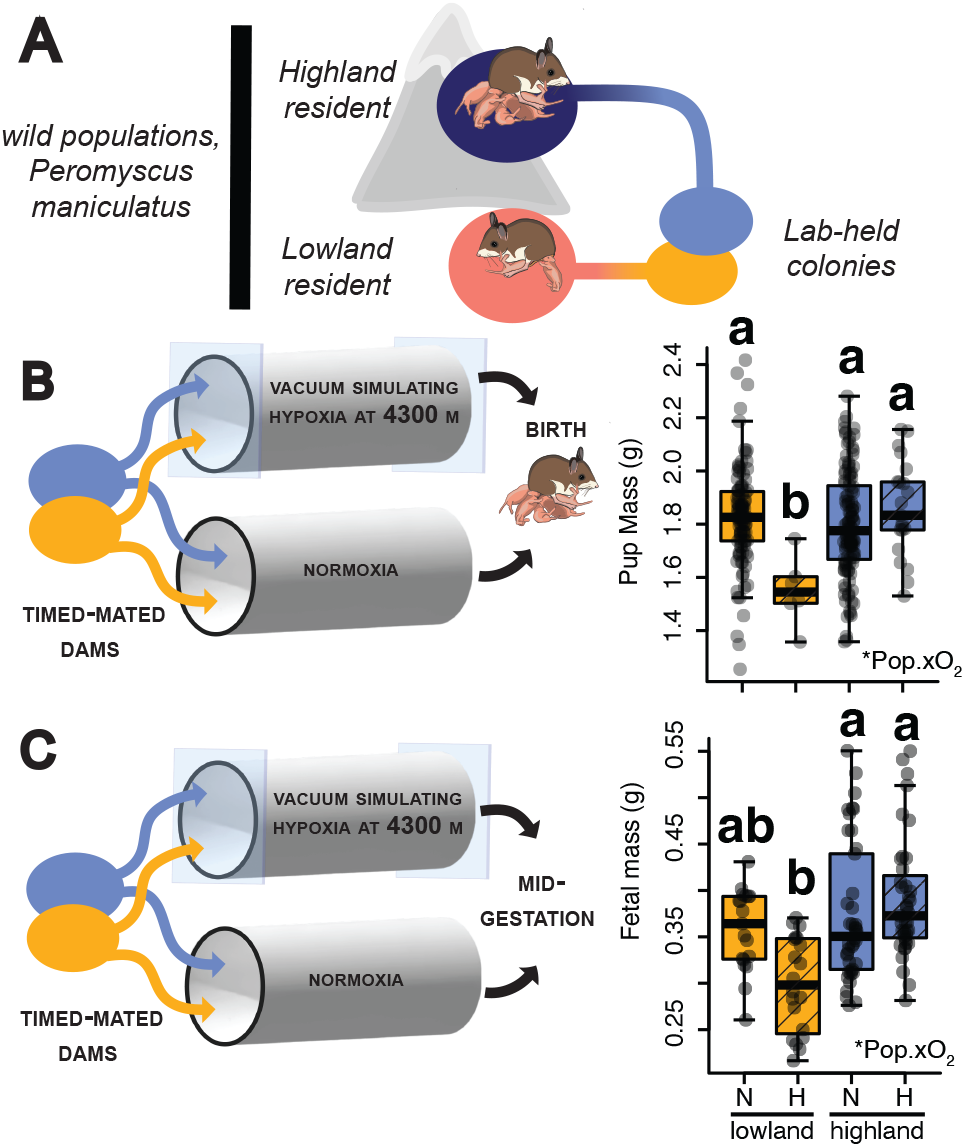
(A) Lowland and highland deer mice were derived from wild-caught populations from low (pink/orange) and high (blue) elevations that have been maintained in lab colonies at low elevation for at least two generations. (B) To test for population-specific effects of hypoxia on birth weight, time-mated dams from each population were held under hypoxia or normoxia until birth in ventilated chambers made from large PVC pipes. While lowland dams gave birth to smaller pups under hypoxia (right; linear mixed model [LMM] overall Pop.xO2 F_1,145.69_=7.13, P=0.008; Low. N vs. Low. H: t_129.4_=2.89, P=0.02), pups born to highland dams were not affected (High. N vs. High. H: t_169.7_=−0.2, P=0.99). Each point represents a single pup from a litter; LMM controls for litter size and includes dam ID as a random effect. (C) By mid-gestation, pups from lowland dams displayed fetal growth restriction (right; LMM, overall Pop.xO2 F_1,35.13_=4.94, P=0.03; Low. N vs. Low. H: t_35.9_=2.31, P=0.05), whereas pups from highland dams did not (High. N vs. High. H: t_32.1_=−0.77, P=0.54). Each point represents a single pup from a litter; LMM includes dam ID as a random effect. Litter size did not affect pup weight at mid-gestation (Table S1). Significant interaction terms are indicated in the bottom right (*P <0.05). Different letters in (B) and (C) indicate significant (P <0.05) differences between group means in post-hoc tests. See Table 1 for sample sizes.

### Maternal physiology and fetal growth under hypoxia

Next, we asked whether maternal physiology could explain population differences in fetal growth outcomes at mid-gestation (day 18.5/19.5 of a 23-24 day gestation; Theiler stage 23/24; N = 8-10 dams per group, Table 1), at which point the fetal growth phenotype is already apparent (Fig. 1C; LMM, Pop.xO2: P=0.03). One way in which maternal physiology may mediate fetal growth is via nutrient acquisition. Most rodents restrict food intake when held under hypoxia, which can limit fetal growth (22). However, food consumption was not suppressed by hypoxia in lowland or highland deer mice (Fig. S1; Table S2). Similarly, gestational hypoxia did not reduce mass gained in lowland or highland dams across pregnancy (Fig. S1, Table S2) nor did maternal body condition explain fetal growth outcomes (Table S3).

Alternatively, maternal cardiopulmonary function could shape fetal growth trajectories under hypoxia (5, 7). We thus tested associations between fetal mass and a range of traits tied to cardiopulmonary function, including hematocrit, lung tissue mass, and heart mass, using linear models (LMs) accounting for maternal body size and *in utero* litter size (See SI Methods). Although we found that several of these traits were indeed affected by hypoxia (Fig. S1, Table S2), these traits failed to explain population differences in fetal growth (Table S3).

We next reasoned that small or interactive contributions of multiple maternal cardiopulmonary traits might influence fetal growth. To test this, we asked whether a combination of maternal physiological parameters explained fetal growth using a principal components analysis (PCA), the output of which is a set of principal components that represent multi-variate metrics (18, 19). However, these multi-variate metrics still failed to explain significant variation in fetal growth among populations and treatments (Table S3).

Our findings thus suggest that maternal physiology does not explain either hypoxia-dependent fetal growth restriction or protection thereof in adapted populations. This result contrasts with findings in humans, where maternal traits like hematocrit or pulmonary function do provide predictive value with regards to fetal growth (4, 5). Because maternal physiology interacts broadly with fetal growth and development (20), it seems unlikely that maternal physiology is entirely inconsequential for fetal growth outcomes in deer mice. Other traits that we did not measure here (e.g., blood pressure or vascular tone in the uterine artery, (4, 7)) may still directly contribute to fetal growth outcomes. Alternatively, maternal traits may work in concert with placental development and function to mediate fetal growth.

### Structure of the placenta and fetal growth

The placenta performs several functions that are essential for successful pregnancies, including gas and nutrient exchange between the maternal and fetal circulatory systems. Altered placental development has been repeatedly linked to fetal growth restriction associated with chronic gestational hypoxia in humans and rodents (5, 21), so we next asked whether placental structure or function explained fetal growth in deer mice. Indeed, we found that placenta mass was positively correlated with fetal mass at mid-gestation in highlanders (linear mixed model [LMM], P=0.03; Fig. 2A,B, Table S4), suggesting that placenta growth or structure contributes to protecting fetal growth under maternal hypoxia.

**Fig. 2.**
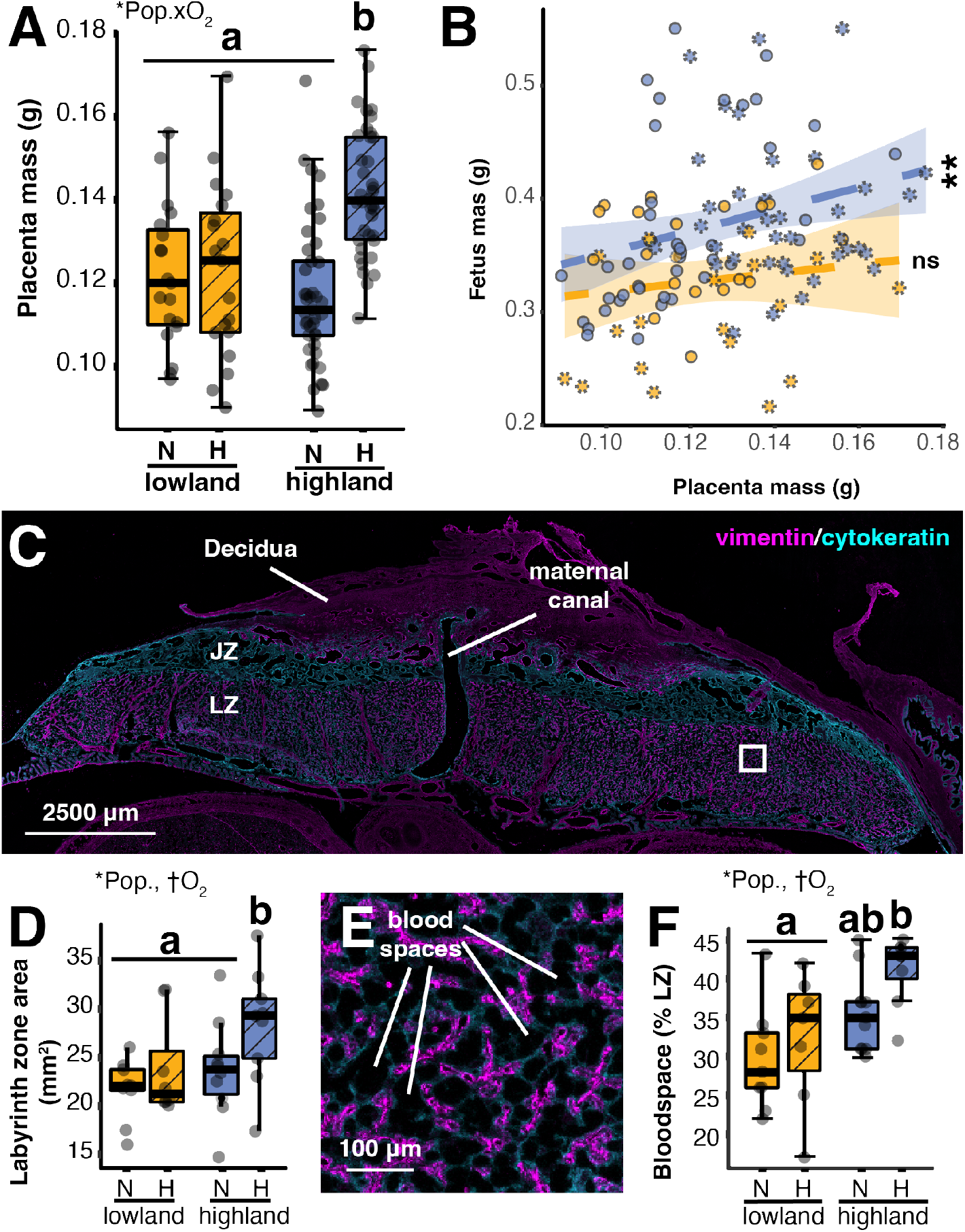
Structural variation in the placenta explains fetal growth protection in highland deer mice. (A) Placenta mass at mid-gestation is larger specifically in highlanders gestating under hypoxia (LMM: Overall Pop.xO_2_, F_1,33.52_=7.65, P<0.01) (B) Placental mass positively correlates with fetal mass, but only in highland deer mice (LMM, P=0.03, Table S4). (C) Compartment sizes and vascular structure within the placenta were quantified from mid-line cryosections that were fluorescently labeled with vimentin and cytokeratin. Maternal canal, decidua, labyrinth zone [LZ], and junctional zone [JZ] are indicated in the figure. White box outlines area expanded in panel E. (D) The labyrinth zone was relatively larger in highlanders under hypoxia than lowlander placentas (LM: Pop., F_1,30_=6.2, P=0.018; O_2_, F_1,30_=3.53, P=0.07; Pop.xO_2_, F_1,30_=3.01, P=0.09). (E) Expanded box from panel C showing blood space in labyrinth zone in detail. (F) Hypoxia increases blood space in the labyrinth zone, particularly in highlanders (LM, Pop., F_1,30_=12.8, P=0.001; O_2_, F_1,30_=3.71, P=0.06; Pop.xO_2_, F_1,30_=0.39, P=0.53). Significant interaction terms are shown in the bottom left of each boxplot (*P <0.05, †P<0.07). Different letters indicate significant (P<0.05) differences in post-hoc, pairwise comparisons among between group means. N = 8-11 implantation sites per group, each from unique dams.

Rodent placentas are organized into three functional layers: the decidua, the junctional zone and the labyrinth zone (reviewed in (22, 23)). The decidua is comprised predominantly of maternal tissues, and it is the region where fetal and maternal cell types interact to establish blood flow from maternal circulation to the placenta. The fetal compartments of the placenta include the junctional zone, which plays a key role in the production of hormones producing cell types responsible for vascular remodeling, and the labyrinth zone, where maternal and fetal circulatory systems are brought into close apposition to facilitate gas and nutrient exchange. Because each layer of the placenta performs distinct functions, the relative contribution of each to placental expansion present distinct functional hypotheses about how expansion protects fetal growth under maternal hypoxia. We therefore asked whether underlying variation in placental structure could explain fetal growth.

Using immunohistochemistry to differentiate layers along the midline of placentas collected from dams at mid-gestation (Fig. 2C), we found that placentas from highland dams gestating under hypoxia had larger labyrinth zones as a proportion of the fetal placenta (LM, P<0.04 for all contrasts; Table S5). The highlander labyrinth zones from hypoxic pregnancies also had a greater proportion of their volume allocated to blood space (Fig. 2E,F, Table S5), and the maternal canal (a prominent vascular structure in the placenta though which maternal blood returns to maternal circulation) was largest in placentas from highland dams gestating under hypoxia (Table S5).

These results point to vascular growth and organization within the labyrinth zone as key factors mediating fetal growth under chronic gestational hypoxia. Our data suggest that increases in blood delivery and the resultant opportunity for nutrient and gas exchange within the placenta contribute to fetal growth protection in highland deer mice. Laboratory strains of mice and rats gestating under hypoxia similarly expand labyrinth zone blood space, and this modification is associated with an increase in placental mass near term (21, 24, 25). In contrast, the villous portion of the human placenta, which is functionally analogous to the labyrinth zone (17), tends to be smaller at term in high elevation pregnancies (5, 7). Nonetheless, humans with highland ancestry display other structural changes to the vasculature in this compartment that may counteract the overall decrease in volume (5, 7). Thus, structural differences that modify nutrient and gas exchange appear universally relevant to fetal growth outcomes under chronic gestational hypoxia.

Given our experimental design, we cannot say whether placental mass would be elevated in lowlanders subjected to gestational hypoxia near term, which would replicate the apparent adaptive structural remodeling shown in other laboratory rodents (21, 24, 25). However, fetal growth restriction persists in these populations despite labyrinth zone expansion, demonstrating that these responses remain insufficient to prevent fetal growth restriction in some lowland lineages (24, 25). These limitations may stem from insufficiencies in the magnitude or timing of expansion in lowlanders. Alternatively, full protection of fetal growth may require additional adaptations.

### Shared genes involved in gestational hypoxia and high elevation adaptation in humans and deer mice

Hypoxia-induced changes in endocrine signaling and gene expression are also likely to be important contributors to the that development and function of the placenta, and subsequent protection of fetal growth. Because of its roles in hormone production and coordinating maternal vasculature development at the implantation site, the junctional zone of the placenta has been a major focus for understanding fetal growth during gestational hypoxia to-date (26–30). We were therefore interested in assessing the transcriptomic responses of each compartment (junctional zone/decidua and labyrinth zone) to hypoxia and in linking gene expression within each compartment to fetal growth trajectories.

We performed layer-enriched RNAseq on labyrinth zone and junctional zone/decidua tissues from lowland and highland dams sampled at mid-gestation (N = 19-20 implantation sites per treatment per population, Table 1; layer enrichment following (31)). After filtering, we detected expression of 14,345 genes in the labyrinth zone, and 14,000 genes in the junctional zone/decidua tissues (see Methods and Supp. Info.; Table S17). To identify genes that were relevant to fetal growth outcomes and adaptive evolution in response to environmental hypoxia, we focused on genes whose expression (i) correlated with fetal mass, (ii) differed with gestational treatment (hypoxia vs. normoxia), and/or (iii) differed between populations derived from different elevations. We first identified genes falling into these categories using a LMM framework (dream, (32)); this approach allowed us to account for non-independence among samples collected from the same dam. We balanced sample sizes across groups for fetal sex, however we did not find substantive effects of sex on transcriptome-wide expression among tissues, treatment (hypoxia or normoxia), or population (lowlander or highland) (see **Supporting Information** for details). Accordingly, fetal sex was not considered further.

We first examined expression within an *a priori* set of genes hypothesized to be relevant to high elevation adaptation, placental physiology, and fetal growth outcomes in humans (33–41). Placental evolution has likely involved recurrent co-option of common genetic pathways and neofunctionalization of gene duplications underlying similar developmental processes (42–44). To test for shared placental genetic responses to hypoxia in humans and mice, we evaluated an *a priori* gene list comprised of 212 genes and 7 receptor/ligand or gene families in humans, for which we were able to identify 253 genes in deer mice (orthologs and paralogs; Table S6). Of these, 204 were expressed in the deer mouse junctional zone/decidua and 208 were expressed within the deer mouse labyrinth zone. The expression of five genes (2.4%) in the junctional zone/decidua and 30 genes (14.4%) in the labyrinth zone were correlated with fetal growth outcomes (Fig. 3A). These proportions represent significant enrichment relative to proportion of genes correlated with fetal growth in the full dataset (Fisher’s Exact Test; junctional zone/decidua: P=3.5 E-8; labyrinth zone: P=0.04), suggesting that candidate genes shared between humans and deer mice are likely involved in similar core placental processes.

**Fig. 3.**
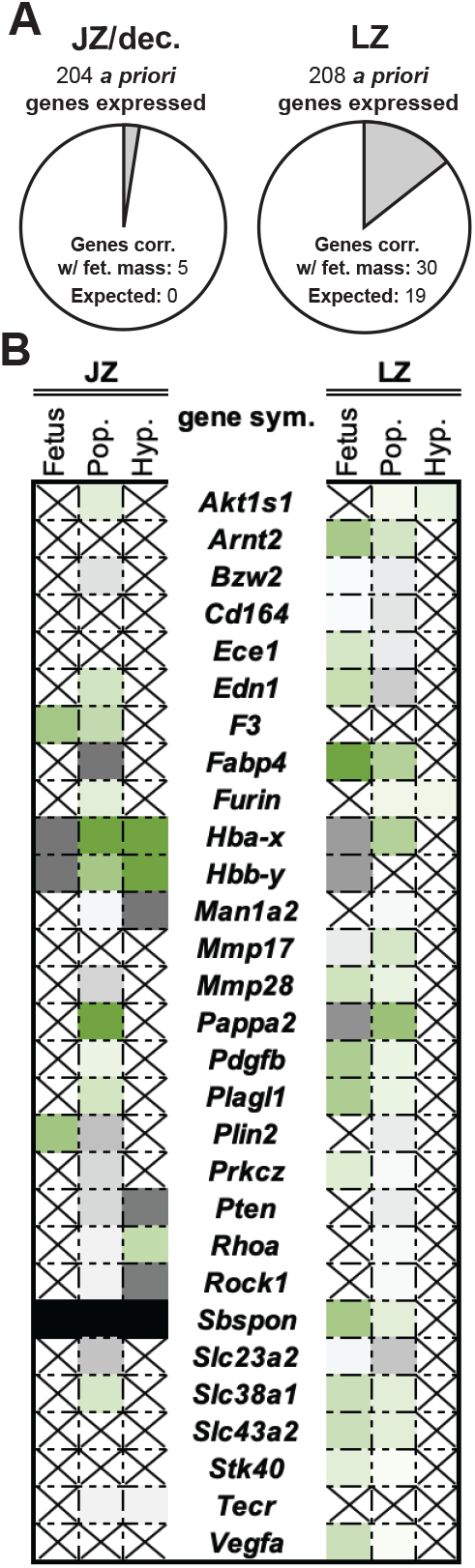
(A) Pie charts showing enrichment for genes significantly correlated with fetal mass. (B) Summary table showing all a *priori* genes for which at least two categories in one tissue were significant. For all columns, cells are filled in significant correlations such that green cells are positively correlated or up-regulated, and grey cells are negatively correlated or down-regulated. Intensity of the color (i.e., light vs. dark grey) indicates log-fold-change of the effect. Numeric values are provided in Table S6. Black (*Sbspon* in JZ) indicates absence of expression in that tissue. For the population (“Pop.”) column, up-regulation (i.e., green fill) indicates greater expression in lowland deer mice, and down-regulation (i.e., dark grey fill) indicates greater expression in highland deer mice. Crosses indicate no significant relationship.

Of the genes in our *a priori* list, nine genes in the junctional zone/decidua and 21 genes in the labyrinth zone were significant for at least two of our focal comparisons (correlation with fetal mass, population effect, oxygen treatment effect, or their interaction; Fig. 3B; Table S6). In the junctional zone/decidua, we found associations between fetal growth and the expression of fetal hemoglobins *Hba-x* and *Hbb-y*(oxygen transport), *Plin2* (trophoblast cell survival under hypoxia (45)), and *F3* (clotting, pre-eclampsia risk (46)). In the labyrinth zone, the *a priori* genes for which expression was associated with fetal growth were involved in diverse processes including nutrient transport (i.e., *Slc32a2, Slc38a1, Fabp4*; (47)), vascular growth and function (i.e., *Edn1* and *Plagl1*; (48–50)), and trophoblast cell differentiation or survival (i.e., *Pappa2, Prkcz*, and *Mmp28*; (51–54)). Two of the most prominent gene candidates in humans, *PRKAA1* and *EDNRA* (33), were not significant for multiple focal comparisons of interest in deer mice. However, *Prkaa1* expression in the labyrinth zone was significantly correlated with fetal growth, and *Ednra* expression in both the labyrinth and junctional zones was constitutively lower in highlanders. These genes may thus still be of relevance to placental or fetal development under hypoxia in the deer mouse, as they are in humans. Overall, broad concordance among genes that underlie fetal growth outcomes in humans and deer mice suggests that there are fundamental similarities in how chronic hypoxia shapes placental development, likely resulting in evolutionary convergence in how mammals have mediated the challenges of gestation at high elevations.

### Global transcription associated with fetal growth

A major limitation of *a priori* gene sets is that they are less likely to discover novel mechanisms. We therefore surveyed transcriptome-wide patterns of gene expression to identify additional genes associated with fetal growth in deer mice. We first used a gene-level expression analyses (dream (32)) to test for associations between expression and fetal growth outcomes. We found that expression of 1,354 genes in the labyrinth zone correlated with fetal growth; in striking contrast, fetal growth was correlated with the expression of only three genes in the junctional zone/decidua (Dataset 2). The relative absence of associations between gene expression in the junctional zone/decidua and fetal growth suggests that junctional zone and decidual cell types are not strongly tied to hypoxia-dependent variation in early fetal growth phenotypes in deer mice. The labyrinth zone, on the other hand, showed patterns of extensive gene expression associations, which are consistent with our histological data showing that labyrinth zone structure contributed to protection of fetal growth in highland deer mice.

### Hypoxia-dependent fetal growth and expression of fetal hemoglobins

Two of the three genes correlated with fetal growth in the junctional zone/decidua were also among those genes correlated with fetal growth in the labyrinth zone, and these genes (fetal hemoglobin subunits alpha and beta (*Hba-x* and *Hbb-y*) had large effect sizes in both compartments. Fetal hemoglobin genes have greater oxygen affinity than their adult counterparts and are critical for moving oxygen from the maternal circulation into fetal circulation (55). Greater expression of fetal hemoglobins could support greater oxygen uptake by fetal tissues, thereby protecting fetal growth under maternal hypoxia. However, we found the opposite pattern: in both the junctional zone/decidua and labyrinth zone, expression of *Hba-x* and *Hbb-y* was negatively correlated with fetal mass (i.e., greater expression of hemoglobins was associated with smaller fetuses). Overexpression of hemoglobins is associated with microvascular damage and endothelial toxicity (56, 57), which could have detrimental effects on fetal growth. However, for a given expression level of *Hba-x* and *Hbb-y*, highlander fetuses still tended to be larger than lowlander fetuses (Dataset 2). Thus, if hemoglobin expression does contribute to fetal growth restriction via endothelial damage, there must be additional factors that exacerbate hemoglobin-related endothelial damage in lowlanders and/or mechanisms by which highlanders are protected from hemoglobin-related endothelial toxicity.

### Angiogenic processes and fetal growth under hypoxia

We next focused on identifying the potential functions or processes by which changes in gene expression may be influencing fetal growth trajectories. With nearly 10% of expressed genes in the labyrinth zone associated with fetal growth, gene ontology (GO) analysis identified enrichment for a number of broad biological processes related to cell replication, division, and differentiation (Table S7). Among genes with expression positively correlated with fetal mass, we found enrichment for terms specifically linked to blood vessel growth and development (Table S8). In contrast, genes with expression that was negatively correlated with fetal mass remained enriched for broad cell division and replication terms (Table S9).

To resolve potential regulatory networks and the sets of co-expressed genes involved, we next applied an unsupervised network-based approach (WGCNA (58)) to our transcriptomic data. Unsupervised network-based approaches identify sets of genes with correlated patterns of expression (hereafter, gene modules) that can help identify regulatory mechanisms not apparent from gene-level tests of differential expression. The expression of these modules can be summarized using the first principle component of individual gene expression values (also known as the module eigengene E; (58)), and these eigengene values can then be associated with phenotypic outcomes. Using WGCNA, we constructed co-expression networks for each tissue separately; WGCNA identified 19 gene modules in the junctional zone/decidua and 20 modules within the labyrinth zone (See SI Methods for details). For ease of reporting results, we numbered modules consecutively by gene content such that the number of genes within the module increased as module number increased (i.e., the module M1 contained the fewest genes, and M20 contained the most genes; see Fig. 4).

**Fig. 4.**
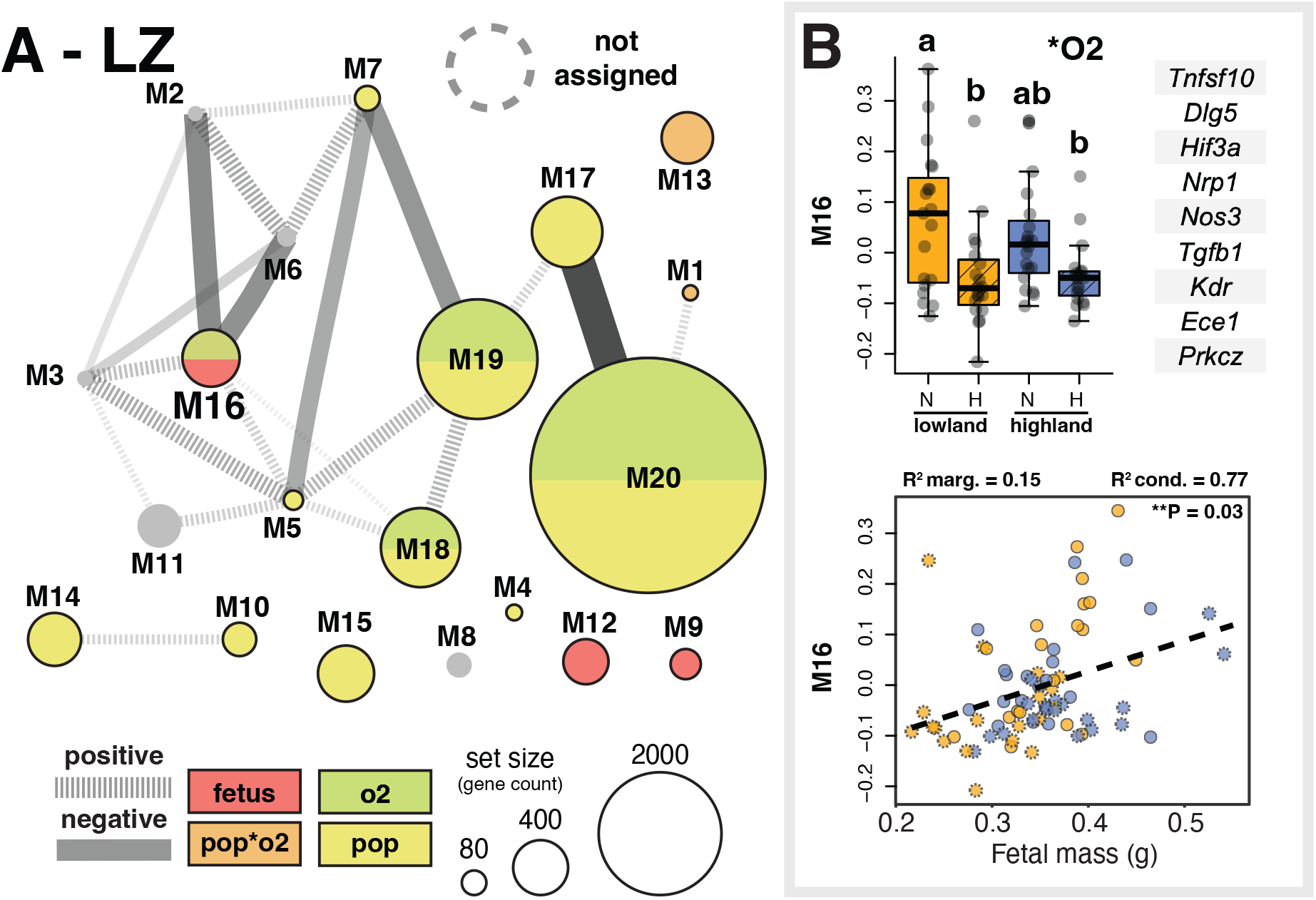
(A) WGCNA module network illustrating gene expression and module connectivity within the labyrinth zone [LZ]. Modules (ellipses) are scaled by the number of genes in each module. Edges connect modules with a Pearson correlation of >0.6. Width and opacity of edges corresponds to strength of the correlation such that wider and darker edges indicate stronger correlations. Solid edges indicate positive associations, and dashed lines indicated negative correlations. Modules are also colored by their association with outcomes or experimental treatments of interest (fetal mass, red; population-by-hypoxia interactions, orange; hypoxia, green; population, yellow). (B) Plots of M16 eigengenes (see panel A) showing differences among treatment groups (boxplot) and correlation with fetal mass (scatter plots, right). Genes of a *priori* interest included in this module are listed to the right of the boxplot. In the boxplot, significant main effects are indicated in the top right (*P<0.05), and different letters indicate significant (P<0.05) differences between group means in post-hoc pairwise comparisons using a Benjamini-Hochberg p-value adjustment. In scatterplot, dashed lines show the linear relationship between fetal mass and the module eigengene. Within the scatterplot, p-values for the relationship between fetal mass and module eigengene is shown in the top right, and pseudo-R-squared values for the LMM are shown above.

We identified 3 modules in the labyrinth zone whose expression patterns correlated with fetal mass (Fig. 4A, Table S10). As expected from our gene-level differential expression analyses, there were no modules in the junctional zone/decidua that correlated with fetal mass (Fig. S2, Table S11). In the labyrinth zone, expression patterns in two of the three modules that correlated with fetal growth (M9 and M12) did not differ by population or hypoxia treatment (Table S10), suggesting that these gene sets influenced fetal growth but were not involved in hypoxia-dependent growth outcomes. In contrast, gene module M16 (434 genes) was positively correlated with fetal growth and hypoxia-responsive (Fig. 4B), and M16 was enriched for gene functions tied to angiogenesis and blood vessel formation (Table S12). This module also included a number of angiogenic-associated genes identified in our *a priori* gene list (Fig. 4B). However, unlike patterns apparent in our gene-level differential expression analysis, M19 was hypoxia responsive, indicating that M19 is characterized by angiogenic genes that were also differentially-expressed in response to hypoxia. The positive correlation between this module and fetal growth along with the suppression of these genes in response to hypoxia (Fig. 4B) suggests that the down-regulation of these core vascular growth and angiogenic genes associated with fetal growth restriction in lowland deer mice. However, because highland expression follows similar patterns in the absence of fetal growth restriction, highlanders must also possess some adaptations that mediate these effects on fetal growth.

### Gene expression associated with adaptive protection of fetal growth in highlanders

In search of gene sets that might explain highlander protection of fetal growth, we next focused on genes and gene modules that were differentially expressed by both population and hypoxia, a pattern suggestive of evolutionary modification of the regulation of hypoxia-sensitive gene networks (59). In the labyrinth zone, we found 228 genes and five gene modules where expression was sensitive to hypoxia and differed between highlanders and lowlanders, either through interactive effects (125 genes, and modules M1 and M13), or additive effects (103 genes, and modules M18, M19, M20) (Fig. 4A, Fig. S2). Here, we focus on those genes and modules where we found evidence for interactive effects.

M1 contained only 30 genes, which were enriched for the broad term “organelle membrane” and included genes associated with endoplasmic reticulum and Golgi apparatus function (Table S12). The larger module, M13, contained 350 genes that tend to be up-regulated by lowlanders under hypoxia, but down-regulated by highlanders (Figure S2). M13 contained the genes *Fosb* and *Igf2*, which are closely connected to placental growth and growth factor regulation (60, 61). Specifically, house mice (*Mus musculus*) up-regulate expression of *Igf2* in response to moderate hypoxia (24), which should benefit fetal growth by promoting vascular growth and expansion in the labyrinth zone (62). However, we found that highland deer mice *down-regulate Igf2* expression in response to hypoxia (Dataset 2), suggesting that up-regulation of *Igf2* seen in lowland rodents incurs some costs (i.e., the hypoxia-dependent plasticity has been reversed by natural selection, (59)). Pathway-specific investigations linking transcript abundance, protein production and half-life, and functional outcomes like microvascular structure or fetal growth trajectory under hypoxia will be necessary to better understand the trade-offs governing hypoxia and *Igf2* interactions in the placenta.

M13 was also functionally enriched for RNA metabolism and transcription (Table S12). This functional enrichment was also apparent in our gene-level differential expression analyses; the 125 genes that were individually significant for the interaction term (i.e., Pop.xO_2_) were enriched for a number of transcription factor binding motifs and ribonuclear protein terms (Table S12). Thus, the regulation of RNA synthesis and processing likely contributes to placental adaptations to maternal hypoxia. These processes may influence fetal growth by interacting with vascular growth pathways that were otherwise inhibited by hypoxia. Indeed, transcriptional repression and decreases in protein production have been previously associated with hypoxic stress in the placenta (e.g., (63)), and thus one way in which highlanders may protect fetal growth is by preventing this suppression.

### The genetic and genomic basis of placental adaptations in highland deer mice

Previous population genomic scans focused on elevational adaptation in mammals (e.g., (2, 12, 64–67)) have lacked a detailed understanding of placenta-specific developmental responses to gestational hypoxia, including intermediate gene expression changes (reviewed in (68)). Our transcriptomic data thus uniquely positioned us to advance our understanding of genotype-to-phenotype connections in reproductive traits.

To ask whether local adaptation in high elevation deer mice has targeted genes or sets of genes relevant to fetal outcomes, we cross-referenced our placental gene sets with genes previously shown to be the targets of spatially-varying selection associated with elevation in deer mice (12, 15). We identified genes under positive selection in highland deer mice using two metrics from (15): a redundancy analysis and the Population Branch Statistic (PBS; (69)). The redundancy analysis was used to test for genotypic associations with elevation. The PBS is a measure of population differentiation similar to the population fixation index (F_ST_), but it uses an outgroup to polarize the direction of selection (described in more detail in (12, 69)). Using this approach, higher PBS values signify greater levels of differentiation in our high elevation population relative to two low elevation populations (12), suggestive of positive selection specific to the highland population (69). To determine the significance threshold for empirically-derived PBS values, (12) used the inferred demographic history of these three populations to simulate a neutral background distribution of PBS to account for neutral differentiation and minimize false positives due to genetic drift (see discussion in Supporting Information). Values above the 99.9^th^ percentile in the simulated distribution were considered significant outliers. In combination, these approaches identified 993 genes under selection in the high elevation Colorado-based population from which we sourced our experimental highland mice (Table S13).

We found that 626 of the genes bearing signatures of selection at high elevation were expressed in the labyrinth zone. Using this full set of expressed genes under selection as a background expectation, we found significant enrichment for genes under selection in our *a priori* gene list (Fisher’s Exact Test, P=0.002). Enrichment for positive selection targets in this gene list reinforces our previous findings that many of the genes involved in mediating the effects of gestational hypoxia on fetal growth are shared between humans and deer mice. While some of these genes coordinate hypoxia responses across many tissue types (e.g., *Angpt1, Epas1*), there were also genes within this set that have well-established and specific functions connected to placental development (e.g., matrix metalloproteinases (21, 54), and the vascular endothelial growth factor receptor *Flt1* (21, 70)). However, we did not find that targets of selection were over-represented among genes correlated with fetal mass or among genes that exhibited population-specific responses to hypoxia. The absence of overrepresentation among these gene sets suggests that the genes identified from our experimental manipulations are not each direct targets of local genetic adaptation. Instead, there are likely a small number of selection targets that shape transcriptome-wide patterns of expression (i.e., selection on a few regulatory factors) or that there are a few targets that have key functional impacts on their own.

Recently, (14) showed that large inversions in the genome are common across *P. maniculatus* populations and tend to harbor adaptive alleles that contribute to local adaptation in other deer mouse ecotypes. To test whether inversions might also contribute to adaptive gene expression in the highland deer mouse placenta, we used data from (14) to identify 14 inversions segregating in highland and/or lowland deer mouse populations. Eight of these polymorphic inversions showed large haplotype frequency differences (>0.6) between highland and eastern lowland populations, consistent with an association between the inversion haplotypes and adaptation to high elevation environments (Table S14). We then asked whether these inversions were enriched for positive selection candidates expressed in the labyrinth zone and/or for genes that were associated with outcomes of interest (e.g., fetal mass, Pop.xO_2_ interaction) in our differential expression analyses. Although these large inversions (ranging from 1.5 Mb to 43.8 Mb) contained relatively few genes expressed in the labyrinth zone (Range: 7 – 274 genes), all 8 inversions showing large allele frequency differences between highland and lowland deer mice were enriched for selection candidates expressed in the labyrinth zone (Hypergeometric tests; P<0.006 for all inversions; Table S15, Fig. S4). Notably, the 8 elevation-associated inversions contained over 1/3^rd^ of all selection candidates expressed in the LZ, but only 4% of all genes in the genome. We further found that the inversions on chromosomes 6, 7, and 15 were enriched for genes that were included in differential expression gene sets (e.g., Pop.xHypoxia, Fetus-assoc.) (Fig. S4).

Collectively, these results suggest that inversions are likely to play important roles in structuring the genomic architecture of local adaptation to high elevation in deer mice, and that at least some of these inversions contain genes that contribute to population-specific fetal growth trajectories under gestational hypoxia. Moreover, the lack of enrichment for positive selection targets broadly among differentially-expressed genes indicate that the causal drivers of genome-wide transcriptomic responses to hypoxia are likely determined by evolutionary changes across relatively few key genes. That is, many of the changes in gene expression that track variation in fetal mass may represent correlated outcomes of a small number of selection targets, rather than the result of selection targeting many of these genes individually. This inference should be considered preliminary since the genic targets of selection were identified from whole-exome data, which likely fails to capture differentiation of regulatory sequences that are not closely linked to sequenced exonic regions. Whole genome sequence data will be necessary to clarify relevant sequence variation in regulatory regions of genes that are functionally associated with fetal growth trajectories.

### A weighted-ranking approach to stratify genes in the labyrinth zone underlying fetal growth

Given the limitations inherent to the current genomic scan in deer mice, we were also interested in developing a quantitative ranking for our large set of differentially expressed genes to help nominate specific candidate genes for further functional study. To accomplish this, we calculated a simple aggregate rank for each gene that considered effect size and p-value for associations with fetal growth or differential expression between populations and hypoxia treatment, as well as whether genes were selection targets or *a priori* candidates (Fig. 5A; see SI Methods).

**Fig. 5.**
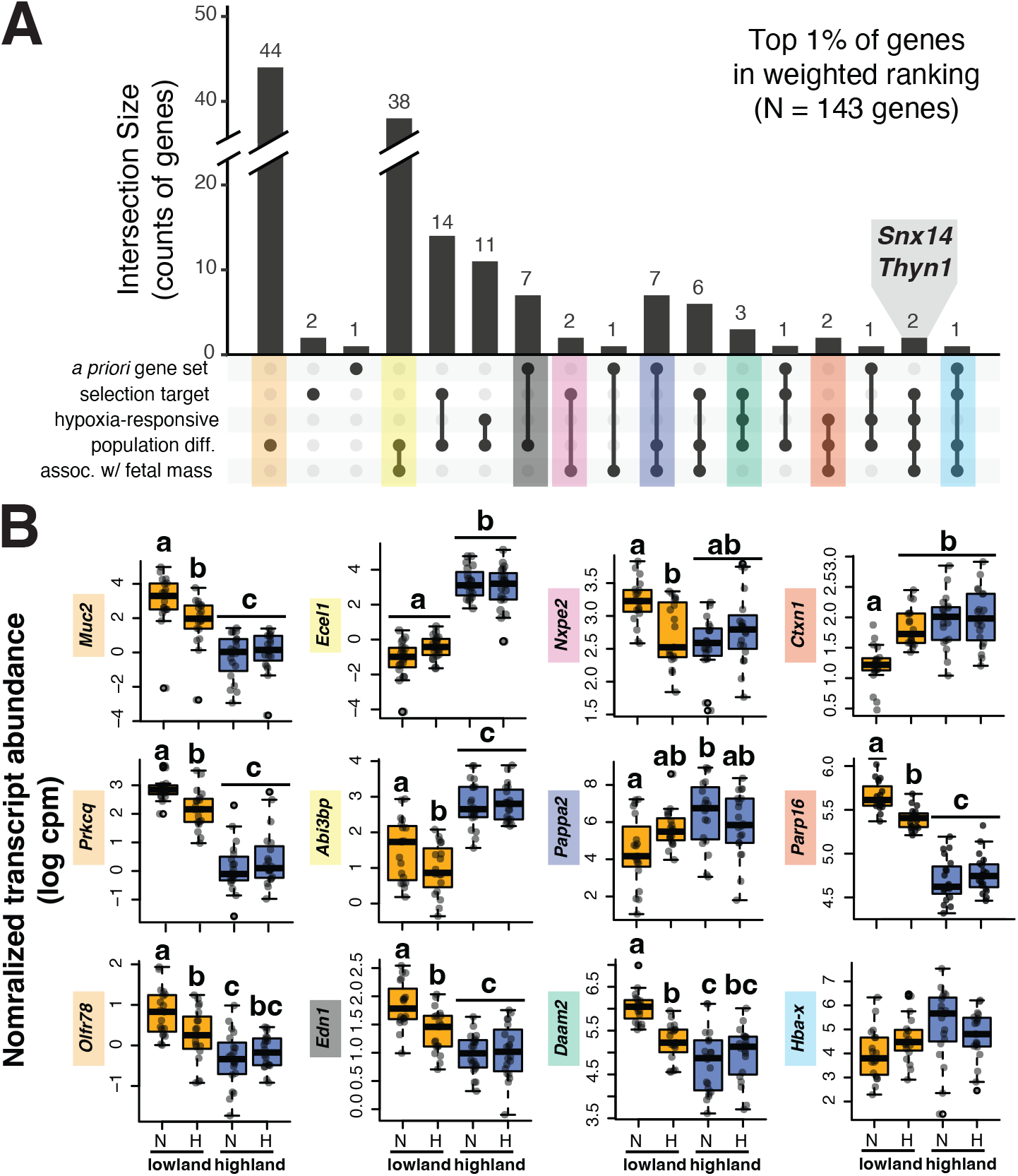
(A) UpSet plot illustrating intersection size based only on FDR-corrected statistical significance in a differential expression framework for all genes within the top 1% of weighted rankings (N = 143). Grey inset shows genes that were significant for 4 categories but that do not appear in panel [B]. (B) Expression differences among populations and gestational treatments for the top twelve (12) candidate genes associated with fetal growth and population-specific gene regulation based on a weighted ranking system approach (see text and methods for details). Each boxplot shows the normalized transcript abundance as log-transformed counts per million (log cpm) by population and treatment. The specific gene is indicated on the immediate Y axis label. Mean group differences were evaluated in a LMM framework (ee Methods). In boxplots, different letters indicate significant (P <0.05) differences between group means in post-hoc pairwise comparisons using a Benjamini-Hochberg p-value adjustment. Colors behind each gene name indicate which intersection set contains that gene.

This weighted ranking approach has the benefit of identifying genes that would not have been recognized through simple set-overlap approaches. Nonetheless, we found three genes within the top 1% of ranked genes that were also significantly associated with many of the gene sets of interest (Fig. 5A). These include a fetal hemoglobin (*Hba-x*, discussed above), and two more enigmatic genes, *Snx14* and *Thyn1* (Fig. 5, inset). Although both *Snx14* and *Thyn1* have been linked to adverse gestational outcomes (71, 72), these genes are expressed near-universally among placental cell types in humans and house mice (73–75), and thus their mechanism of action and cell type-specific importance remains unclear. Ultimately, experimental single-cell data from *Peromyscus* placentas and cell-type specific experimental work focused on these genes are likely necessary to ascertain their role in placental development.

Our top ranked gene (regardless of weighting) was *Muc2*,a gene belonging to the mucin family. Mucin proteins are secreted onto epithelial surfaces where they provide lubrication and chemical barriers and can perform cell signaling roles (76, 77), suggesting that the extracellular environment may be a key factor shaping population-specific effects of gestational hypoxia on fetal growth. Indeed, the top 1% of genes in our ranking were enriched for functions that are involved in extracellular matrix organization and collagen metabolism (Table S16). Although *Muc2* expression in humans is absent from the placenta, the placental production of other mucins is common, and dysregulated expression of mucins have been implicated in a variety of placental development and pathologies (78–81). More broadly, collagen dysregulation and its metabolism in the extracellular matrix have been linked to vascular defects in the labyrinth zone (82–84). Combined with our earlier analyses pointing to angiogenesis broadly as relevant to fetal growth outcomes, our weighted ranking analysis specifically suggests that processes involved in *building* vasculature (as opposed to regulatory signals that promote vasculogenesis) are likely key determinants of fetal growth trajectories.

The top 12 genes in our ranking are provided in Fig. 5B. Many of these genes have putative roles in fetal growth or placental hypoxia responses, which affirms that mechanisms well-studied in other model systems and humans are important in deer mouse placentation and fetal growth. For example, placental expression of *Edn1* and *Abi3bp* in the placenta are associated with hypoxia-related pregnancy complications in humans (21, 50, 85).

Perhaps most compelling, two of our top 12 genes (*Daam2* and *Pappa2*) are intimately associated with placental development and fetal growth. Hypoxia-dependent overexpression of *Daam2* has recently been implicated in fetal growth restriction in humans (86). Although *Daam2* expression was not directly correlated with fetal mass in our deer mice, down-regulation of expression (Fig. 5B) is consistent with protective effects for fetal growth. Thus, although the response to hypoxia differs between humans and deer mice, the role in shaping fetal growth may be similar. *Pappa2* has been a gene of interest in fetal growth and hypoxia research for over a decade and thus much is known about its potential role in hypoxia-related gestational complications. Overexpression of *Pappa2* has been associated with inhibition of trophoblast migration and pre-eclampsia development (87–89), and indeed we find a strong negative correlation between *Pappa2* expression in the labyrinth zone and fetal growth outcomes in deer mice (Dataset 2). However, *Pappa2* is constitutively expressed at high levels in the highlander labyrinth zone (Fig. 5B). Interestingly, Andean women resident at high elevations also display elevated serum concentrations of PAPPA2, despite its association with pre-eclampsia risk in this population (88). This could suggest that elevated expression of *Pappa2* is adaptive in other contexts that preclude placental adaptations to expression. Alternatively, post-transcriptional gene regulatory mechanisms, such as those identified our gene module analyses, may mediate the effects of high *Pappa2* transcript abundance in highland deer mice.

Beyond the data from Andeans, which did not assess adaptive function, neither *Pappa2* nor *Daam2* have previously been suggested as relevant to adaptation to high-elevation in humans or other mammals. Experimental work that demonstrates the functional relevance of these candidate genes (e.g., using genetic tools to manipulate expression in specific cell types, either *in vivo* or *in vitro*) is necessary to begin to understand their functional importance for hypoxia-dependent fetal growth restriction and adaptive processes that may modify those associations.

### Conclusions

Understanding the basis of reproductive adaptations to high elevation has the potential to yield important insights for fields of medicine, physiology, and evolutionary biology. However, progress in these areas has been hampered by the absence of an accessible study system. Our work shows that an established rodent model for adaptation to high elevation, *Peromyscus maniculatus*, can be used to understand the drivers of fetal growth trajectories in lowland and highland populations under hypoxia.

We found both structural and functional evidence that fetal growth under hypoxia is tied to the development and organization of the placental zone responsible for nutrient and gas exchange. Hypoxia-dependent suppression of gene expression related to angiogenesis and vascular growth in the placenta appears to be an ancestral response to hypoxia that persists in highland-adapted deer mice. As part of their adaptation to high elevation environments, highland deer mice have overcome these effects through (a) modification of pathways that ultimately promote expansion of the placental compartment responsible for gas and nutrient exchange, (b) alterations to hypoxia-sensitive expression of genes tied to the regulation of RNA transcription and processing, (c) sequence evolution in genes associated with to hypoxia-dependent fetal growth trajectories, and (d) structural features (i.e., inversions) in the genome that preserve associations among adaptive alleles.

We also showed that many of these genes relevant to adaptive phenotypes in deer mice are the focus of on-going work in humans or have understood roles in human placental development or responses to hypoxia. The deer mouse and human placenta thus likely share fundamental gene networks involved in mediating hypoxia responses. In addition to affirming deer mice as a potential translational model, these findings point to conserved or convergent gene regulatory patterns that shape adaptive evolution in divergent mammals.

Ultimately, the experiments and analyses presented here only scratch the surface of a complex physiological trait (fetal growth). Our results point towards several mechanisms that may contribute to population-differences in susceptibility to hypoxia-related fetal growth restriction. The links between differential gene expression and fetal growth trajectories may simply involve altered abundance of those same proteins, or complex changes in transcript half-life, RNA interference, or protein-protein interactions. Similarly, we expect cell-type within the placenta to provide critical context for furthering our mechanistic understanding of how genetic variants lead to altered placental development and fetal growth protection under maternal hypoxia.

Our findings thus establish a basic understanding of the genetic and physiological factors associated with gestational outcomes in a tractable rodent model that can be used to pursue a much broader set of questions. This model opens new avenues for exploring how mammalian reproduction adapts and evolves to meet fundamental challenges in ways that also inform research interested in clinical interventions or diagnostics that are important for maternal-fetal health.

## Data Availability

Phenotypic and histological data generated and analyzed as part of this study will be included in the published article online supporting files (Datasets S1 and S2). Raw reads from RNAseq datasets will be made available via SRA accession.

## Supporting Information Appendix (SI) included

### SI Datasets

Datasets S1 and S2 are provided with this manuscript.

## Materials and Methods

### Animal breeding, handling, and experimental design

All experimental procedures were carried out under IACUC-approved protocols at Univ. of Montana. Highland-adapted deer mice (*P. maniculatus*) were bred at Univ. of Montana from stock trapped at the summit of Mount Evans, CO. All highland-adapted deer mice were second generation offspring from wild-caught individuals (i.e., F2). Lowland deer mice were purchased from the Univ. of South Carolina stock center. Lowland deer mice (BW strain) are derived from a population trapped in Ann Arbor, MI. Further detail on genetic differentiation and divergence between these populations is provided in the **Supporting Information**. Pregnant dams were assigned to either hypobaric hypoxia or normobaric normoxia on day 1 of pregnancy. Animals assigned to hypobaric hypoxia were held under conditions mimicking 4300 m elevation in identical housing as the normobaric normoxia counterparts.

### Sample collection and handling

Placenta, fetal, and maternal tissue and blood samples were collected on day 18.5-19.5 of 23-24 day gestation, corresponding to Theiler Stage 23/24. For each litter, whole implantation sites were collected into isopentane chilled on dry ice for immunohistochemistry. The remaining sites were held in chilled phosphate-buffered saline for dissection. Fetuses and placentas were individually weighed before dissection and snap freezing. Samples chosen for sequencing were distributed roughly evenly across dams in each group while balancing for sex (Table 1, Table S15, Dataset 1). Details on maternal tissue handling and collection of other maternal trait data are provided in the Supporting Information.

### Placental histology

Frozen implantation sites (N = 1 per dam per experimental group) were cryosectioned at 10 um and midline sections (identified by the presence of the maternal canal) were slide mounted for immunohistochemistry. Following (90), sections were fixed with 4% paraformaldehyde, permeabilized using methanol, and blocked using 10% normal goat serum (Vector Laboratories S-1000). Sections were incubated over night with mouse anti-vimentin (Sigma-Aldrich V6630) followed by 1-h incubation with anti-mouse Alexafluor 568 (Invitrogen A11031). Sections were then incubated for 2-h with a pan-cytokeratin antibody conjugated to FITC (Sigma-Aldrich F3418) followed by DAPI to visualize nuclei. Immunostained sections were cover slipped with Fluoromount-G and stored at 4C until imaging on a Zeiss Laser-Scanning Microscope 880 at 10X. Quantification was performed using FIJI (ImageJ 2.0.0-rc-69/1.52p). Measures falling more than 3 SDs beyond the mean were excluded as erroneous.

### Statistical analyses

Comparisons among populations and treatments were carried out in R 4.0.5 using lm() (base R) or lmer()(91). Where relevant (see **Supporting Info.**), we included litter size as a predictor, and we included maternal ID as a random effect. We assessed significance of fixed effects and interactions within models using type III sum of squares in the car package(92), and we performed post-hoc tests within emmeans and lmerTest packages (93, 94) using a Benjamini-Hochberg correction for multiple comparisons. Full model results are provided in **Supporting Information and Tables**.

### RNAseq data generation and analysis

Tissue was homogenized in TriReagent (T9424, Sigma Life Sciences) using a Qiagen TissueLyser, and RNA was extracted using a hybrid TriReagent – RNeasy spin column method. Following TriReagent phase separation, the aqueous phase was used as input to an RNeasy column (Qiagen 74106), after which the manufacturer’s protocol was followed. Stranded, RNA libraries were then prepared by Oregon Health & Sciences University and sequenced using 150 bp PE Illumina NovaSeqS4. We generated an average of 50.7M paired-end raw reads for our junctional zone/decidual samples (Range: 27.1 - 67.1M) and an average of 53.8M paired-end raw reads for our labyrinth zone samples (Range: 31.3 - 78.8M). Data were trimmed for adaptor contamination and quality using Trimmomatic (95). Sequences were then aligned to the *Peromyscus maniculatus bairdii* genome (assembly HU_Pman_2.1.3) using HISAT2 (96). Read counts were determined using featureCounts in Subread (97), allowing for fractional counting of mapping reads. We also annotated and mapped reads to 182 placenta-specific genes from *Mus* that were not annotated within the *P. maniculatus* genome (see Supporting Information). After filtering, mapping, and feature assignment, our analysis included an average of 29.5M reads from our junctional zone/decidua samples (Range: 16.2 - 39.3M), and an average of 31.3M reads from our labyrinth zone samples (Range: 17.2 – 47.4M) (Table S15).

### GO-enrichment analyses

We cross-referenced *P. maniculatus* gene IDs with *Mus* gene IDs via Ensembl before running GO analyses. *P. maniculatus* genes without *Mus* orthologues could not be included in GO analyses, leaving us with 13,632 genes in the LZ and 13,269 genes in the JZ/Dec. for enrichment analyses.

### a *priori* dataset generation

We compiled *a priori* genes of interest from the literature, including genes hypothesized to be relevant to altitude adaptation and protection of fetal growth in humans (33, 35–39, 41) as well as genes with empirical evidence for differential expression among lowland and highland human populations in the placenta (34, 40). (34) and (40) focus on genes are differentially expressed between highlanders (Tibetans or Andeans) and lowlanders and hypoxia sensitivity. From (40), we included only the top 10% of genes that were differentially expressed between highlanders and lowlanders (appx. 100 genes).

### Inversion analysis

Using haplotype frequency data from (14), we identified chromosomal inversions segregating in highland (Colorado) and lowland (Nebraska and California) *P. maniculatus* populations. We then identified gene content within these inversions using genomic coordinates, also from (14). We tested for enrichment of our differentially expressed genes and genes under selection within inversions using a hypergeometric test (*phyper* function in R).

## ACKNOWLEDGMENTS

The authors thank MJ Soares for support and advisement early in project development. The authors also thank the UNVEIL Research Network and members of the Cheviron and Good labs for feedback and input across the duration of the study. This work was supported by the National Institutes of Health (R15 HD103925, ZAC and KW; F32HD100086, KW; R01HD094787, JMG) and the National Science Foundation (IOS-1755411, ZAC; OIA-1736249, ZAC and JMG; DBI-1907233, KW). This study included research conducted in the UM Genomics Core, supported by a grant from the M. J. Murdock Charitable Trust (to JMG) and using computational resources from the UM’s Griz Shared Computing Cluster, supported by the National Science Foundation (CC-2018112 and OAC-1925267, JMG co-PI). The authors thank OHSU for support in RNAseq data generation.

