## Supporting Information for "Adaptive structural and functional evolution of the placenta protects fetal growth in high elevation deer mice"

**This PDF file includes:** **Page**

Supporting Information Text … … … … … … 2
Fig. S1 - Plots of maternal trait data … … … … … 7
Fig. S2 - Junctional zone/decidua module network … … … 8
Fig. S3 - Eigengenes from the labyrinth zone … … … … 9
Fig. S4 - Inversion enrichment plots … … … … … 10
Table S1 - Statistical output for analyses of pup weight outcomes … 11
Table S2 - Statistical output for analyses of maternal physiology … … 12
Table S3 - Statistical output for associations between mat. traits and fetal mass 13
Table S4 - Statistical output for analyses of placental mass … … 14
Table S5 - Statistical output for analyses of placental structure … … 15
Table S6 - *A priori* gene set list and assoc. with tissue-specific gene expression 16
Table S7 - GO enrichment results for all DE genes correlated with fetal mass 26
Table S8 - GO enrichment results for DE genes positively corr. with fetal mass 28
Table S9 - GO enrichment results for DE genes negatively corr. with fetal mass 30
Table S10 - Labyrinth zone modules and associations with outcomes of interest 32
Table S11 - Junctional zone modules and associations with outcomes of interest 33
Table S12 - GO enrichment results for gene modules of interest in the LZ 34
Table S13 - Full list of genes under positive selection in highland deer mice 43
Table S14 - Inversions limits and haplotype frequencies across populations 80
Table S15 - Hypergeometric tests for gene set enrichment in inversions 80
Table S16 - GO enrichment results for top 1% of genes in weighted ranking 79
Table S17 – Read counts across filtering for all samples used in RNAseq 81
Legends for Dataset S1 and S2 … … … … … … 86
SI References … … … … … … … … 87

**Other supplementary materials for the manuscript include the following**

Dataset S1
Dataset S2

### Supporting Information Text

### Expanded Materials and Methods

**Animal breeding, handling, and experimental design:** All animals used in experiments were adults (>P90). Throughout experiments, animals were housed in static caging that was changed weekly. Ad libitum food and water were provided. Animals also received a small number of irradiated sunflower seeds for enrichment weekly.

Pregnancies were timed using the first post-partum estrous. Females were paired with males and weighed weekly to monitor pregnancy status until birth. Pups were weighed on the day of birth. Females remained paired with males and pups for 24 h after birth, at which point the male and pups were removed. Females were then assigned to treatment and singly housed until euthanasia. A subset of mothers were held in hypoxia until the birth of their second litter. These pups were weighed on the day of birth. All births in hypoxia occurred within the 23-24 day estimated gestational period for deer mice; no differences in gestational length between populations or under hypoxia were apparent.

**Maternal physiological traits:** On gestational day 18.5-19.5, females were euthanized using isoflurane overdose followed by rapid decapitation. Trunk blood was collected into capillary tubes for hematocrit quantification. Blood was also used to measure hemoglobin content using Hemocue devices. After processing fetal and placental tissues, we dissected out maternal tissues of interesting, including the lungs and heart. Maternal tissues were cleaned and weighed prior to freezing.

**Maternal consumption:** Food was weighed on day 1.5 and on the day of euthanasia. Food consumption was calculated as the difference between the two measures.

**Body composition:** Body composition (fat mass, lean mass, free body water, and total body water) was quantified with an EchoMRI quantitative magnetic resonance system (E26-281-BHlab, Houston, TX). Animals were measured on day 1.5 and immediately prior to euthanasia. For each measurement, animals were scanned three times and the scans were averaged. Fat measurements were corrected using a linear regression that fitted measured values of canola oil standards on the day of the scan their known masses (15.3, 5.0, 2.0, and 1.0 grams).

**Detailed maternal organ weight measurements:** After thawing, maternal hearts were dissected using fine spring forceps to separate right ventricle mass from left ventricle and septum. Dissected portions of the heart were individually massed to quantify right ventricle hypertrophy. To measure lung tissue mass and edema, lungs were dried in a 60°C drying oven and weighed every 24 h until constant weight.

**Quantification of the maternal canal diameter:** To quantify diameter of the maternal canal in placenta sections, we measured the diameter of the canal at the largest point of the canal. Diameter measurements were taken perpendicular to the sides of the canal. The measurement was taken three times per section and the average of these three measures was used for downstream analysis.

**Quantification of blood space in the labyrinth zone**: To quantify blood space within the labyrinth zone, we converted composite images containing all channels to a single 8-bit image (i.e., greyscale), and then used the threshold function in FIJI to divide pixels into two classes based on degree of fluorescence (either "fluorescently labeled" or "empty"). Pixels labeled as "empty" were counted as blood space. We then imposed a 0.25 $mm^2$ grid on each image and quantified the blood space within four grid squares distributed across the labyrinth zone for each image. These values were summed within each image for analysis. Because images were collected over multiple rounds of staining and imaging, thresholds were visually determined for each image by a single investigator blind to treatment or population, and we included staining round as a random effect in the model. Importantly, thresholds used in this analysis were still similar across images (range: 4-15, mode: 10), and we found no correlation between threshold and blood space calculated per image ($P > 0.7$).

**Statistical analyses of maternal, fetal, and placental mass traits:** Treatment- and population-specific differences in maternal and fetal traits were evaluated using linear mixed models in R(1): we used the *lm* function in base R or *lme4(2)* (when random effects were needed), assessed significance of fixed effects and interactions using type III sum of squares in the *car* package(3), and we performed post-hoc tests within *emmeans*(4) and *lmerTest*(5) packages. Model specifications and output are provided in Tables S1-S3.

In our initial analyses, we found that highland deer mice had larger litters, on average, relative to lowland deer mice both at birth and at mid-gestation (See Table S1 and Dataset 1). Due to our small number of replicates in hypoxic litters at birth, we could not robustly test for differences in litter size between hypoxia and normoxia at birth, however the litter sizes appear similar those collected under normoxia (see Dataset 1). Consistent with these observations, highlanders had larger litters on average at mid-gestation ($P < 1.8$ E -7; Table S1), but litter size was not altered by hypoxia in either population ($P > 0.12$; Table S1).

Litter size has a small but detectable negative effect on birthweight, and thus we included litter size in our birthweight analyses (Table S1). In contrast, litter size did not impact fetal mass at mid-gestation ($P = 0.22$, Table S1), though including litter size did change the significant effect of Population x Hypoxia to a marginal p-value ($P = 0.09$). This impact on the interaction term is due to population differences in litter size: when analyzing each population separately, we do not detect an effect of litter size in either population individually ($P > 0.36$ in both populations, Table S1), and we only detect an effect of hypoxia on pup mass in lowlanders (see Table S1). These analyses suggest that litter size is confounding variance assignment in the model containing both populations (because it covaries with population). We therefore excluded litter size from the model.

In testing for relationships between maternal traits and fetal growth outcomes, we used residuals of tissue and organ masses for fat mass, mass gain, overall mass, heart, right and left ventricle/septum, and lung tissue that were calculated using maternal lean mass on gestational day 1, thereby correcting for maternal body size at the start of the experiment.

We compared variance explained by these models with a model using only the experimental design (i.e., Population x Hypoxia) to ask whether maternal traits explained significant variance in fetal growth, which would suggest that the maternal trait provided better predictive value. For organ analyses, we compared models using AICc from the MuMIn package(6). In all cases, we compared models run on datasets that included identical observations (i.e., we subsetted observations with missing data when appropriate). While some traits did explain variance among dams, the power to explain fetal growth largely overlapped with the predictive power of the experimental design model which ignored trait variation (see Table S3).

To test whether maternal traits in combination predicted pup outcomes, we reduced variance of all maternal traits into major PCA axes using *prcomp* from base R. Because PCA cannot tolerate missing values and some trait data were missing from some dams (e.g., in cases where lung tissue was lost before drying), we first had to impute some missing values. In total, we imputed 12 values across our dataset (2.3% of values), which were spread across 9 traits, with no more than two imputed values per trait. Imputations were performed using *Amelia* (7). Five imputations were generated and then averaged to create the final imputation values. Within our PCA of maternal trait measures, the first two principal components summarized 62.7% of the variation in maternal trait data, and the first five principal components explained over 95% of the variation. We than repeated analyses asking whether PCA loadings from the first 5 dimensions (together explaining >95% of variance in the dataset) predicted average pup mass among dams.

**Fetal sex determination:** DNA samples from mid-gestation pups were digested using ProK in lysis buffer and extracted using the Qiagen DNeasy Blood & Tissue kit (69506) according to manufacturer instructions. Fetal sex was determine using a PCR assay. We used a two-part PCR assay for which ZFY primers amplified only in males, and LGL primers amplified in both sexes. Primer sequences are provided below. PCR amplification was performed using 95°C activation step (5 min) followed by 38 cycles as follows: 94°C denature (30 s), 60°C anneal (30 s), and 72°C extension (1 m 30 s), followed by a 4°C hold. Samples were moved into a 4°C fridge for short-term storage until they were run on a gel (1% agarose, 115 V for 60 m). Sexes were determined by presence/absence of bands on the gel.

| Primer Pair | Forward sequence | R sequence |
|---|---|---|
| LGL | AGACCTGATTCCAGGCAGTACCA | CTTGGCAAGGTGTTCAGGGT |
| ZFY | GGGCTAAGTAATTGGAGGGCATT | TGCATGTCATAGGAGTGTGATGAT |

**RNAseq data processing**: Data were cleaned of poly-G reads using fastp with flags to disable quality and length filtering as well as adaptor trimming (8). We then removed adaptors using BBtools bbduk with default settings (9). Reads were then quality filtered using the settings [ILLUMINACLIP:TruSeq3-PE.fa:2:30:10 LEADING:3 TRAILING:3 SLIDINGWINDOW:5:20 MINLEN:36] in Trimmomatic (10).

**Placenta-specific transcripts from *Mus***: Although mapping rates were high (>90%), a large number of reads fell outside of annotated regions of the genome (~40%). In an attempt to supplement the *P. maniculatus* annotation, which is based on *in silico* annotations confirmed with transcriptomic data that does not include the placenta, we looked for one-to-one orthologues of placenta-specific expressed transcripts in mice in the *P. maniculatus* genome using top blast results. If the best hit region identified by blast already contained or overlapped with an annotated gene in *Peromyscus,* it was dropped. Ultimately, 182 newly-annotated regions had sufficient transcript expression for analysis in our tissues. We were thus able to increase reads mapped to features by 1-2% across samples (average of 0.25M reads in junctional zone/decidual samples and 0.63M reads in labyrinth zone samples).

**RNAseq data analysis:** See Table S14 for sample-specific information on total reads and reads mapped to features used in this analysis. RNAseq data from one placenta (089, both junctional zone/decidua and labyrinth zone) was dropped from analysis based on separation from other data in PCA/MDS plots during QC. In our first analysis, we analyzed data using a simple differential expression framework using dream and limma/voom. We analyzed the full set of samples (both strains and treatments) in a single analysis. Samples with extreme differentiation from the dataset based on PCA and clustering approaches were excluded (one sample in each tissue type). Genes were filtered such that only genes with more than 0.5 cpm in 75% of samples were retained.

Overall, we found large effects of population on gene expression – 6,919 (49.4%) and 7,436 (51.8%) genes were differentially expressed between populations in the junctional zone/decidua and labyrinth zone tissues, respectively. For this reason, we included Population as predictive factor in all analyses to ensure that correlations were shared between strains even when overall expression differed.

To examine differential expression of genes correlated with fetal or placental outcomes, we first used a linear mixed model in dream as:

$$\sim \text{Phenotype} + \text{Population} + (1|\text{DamID})$$

In our second analysis, we used WGCNA to understand how coexpressed networks of genes explained phenotypic outcomes of interest (e.g., fetal and placental mass). For WGCNA analysis, counts were normalized using transcripts per million (TPM). We filtered TPM datasets using the set of genes include in the dream analysis so that analyses were performed on the same gene sets.

To generate signed hybrid networks using WGCNA, we used powers of 11 and 6, respectively, for the junctional zone and decidua and the labyrinth zone based on scale-free independence and mean connectivity values, as recommended in the WGCNA manual (11). Networks for each tissue were then constructed using default settings along with a merging threshold of 0.25, a maximum block size of 14000, and minimum module size of 20. We determined association between WGCNA modules and phenotypes or treatments of interest using linear mixed modeling and formal enrichment tests. First, we tested for significant associations between module eigengenes and phenotypes or treatments in a mixed model framework, including dam as a random effect. Modules that remained significantly associated with the outcome of interest after corrections for multiple testing across modules in each tissue (using Benjamini-Hochberg corrections) were considered associated with the outcome of interest regardless of enrichment testing. For modules that did not remain significant after correcting for multiple testing, we also looked for enrichment within that module for genes that were associated with the outcome of interest in the DE analysis relative to all expressed genes using a Fisher's exact test. Thus, modules could be considered significantly associated with outcomes of interest if either (a) the module was significant for outcome of interest after corrections for multiple testing or (b) the module was nominally significant for the outcome of interest AND enriched for genes DE for that same outcome. Marginal and conditional pseudo-R-squared values were calculated using MuMIn(6).

**Aggregate rank analysis:** Aggregate ranks were calculated using the following approach. First, all genes were assigned a rank (1-14,345) based on the effect size and adjusted p-value within each factor or outcome of interest (Population, hypoxia, the interaction of population and hypoxia treatment, and fetal mass), as shown in Eq. 1. By summing across these ranks (*Eq.* 2) and then examining those with the lowest summed rank score, genes that are marginally associated with many outcomes of interest can to outrank genes that may only be strongly associated with a single trait. However, we also wished to weight gene ranks based on *a priori* expectations about gene importance and what we knew about genes under selection in the deer mouse. We thus assigned weights of 1 or 0 for *a priori* and selection inclusion and weighted the summed rank based using this information as shown in Eq. 3

*Equation 1*

$$|logFC| \times (-log_{10}[p.val.])$$

*Equation 2*

$$AggRank = Rank_{fetus} + Rank_{pop.} + Rank_{hypoxia} + Rank_{popxhyp}$$

*Equation 3*

$$\frac{AggRank}{1 + a\,priori + selection}$$

Although weighting based on inclusion in gene sets under selection or in our *a priori* sets increased the ranking of those genes substantially, the majority of the top 1% of ranked genes (104 of 143 genes) remain in the top 1% of regardless of whether selection and *a priori* status were considered. Furthermore, the top 0.1% of genes after weighting all fell within the top 0.2% of genes prior to weighting; thus, the weighting elevated genes of known or likely importance without swamping the majority of the biological signal in our datasets.

**Evaluating the impact of fetal sex on phenotypic and gene expression responses to hypoxia:** To test for main and interactive effects of fetal sex on mass and placental outcomes, we used linear mixed models (as described above) and included an interaction term between sex and hypoxia. However, we did not find any significant effects of sex on fetal mass or placental mass, nor was there any evidence for an interactive effect of hypoxia and sex on these outcomes:

| Trait | Model | | Sum Sq | Mean Sq | NumDF | DenDF | F | Pr(>F) |
|---|---|---|---|---|---|---|---|---|
| Pup mass (mid-gestation) | Strain*O2 + Sex*O2 + (1\|Mom) | Population | 0.0068285 | 0.0068285 | 1 | 34.823 | 9.2585 | 0.004438 |
| | | O2 | 0.000044 | 0.000044 | 1 | 59.757 | 0.0597 | 0.80786 |
| | | Sex | 0.0020687 | 0.0010343 | 2 | 85.484 | 1.4024 | 0.251602 |
| | | Pop.:O2 | 0.0037012 | 0.0037012 | 1 | 34.823 | 5.0183 | 0.031564 |
| | | O2:Sex | 0.0021481 | 0.001074 | 2 | 85.484 | 1.4562 | 0.238837 |
| Placenta Mass | Strain*O2 + Sex*O2 + (1\|Mom) | Population | 0.00051055 | 0.00051055 | 1 | 32.183 | 2.4284 | 0.12893 |
| | | O2 | 0.00028852 | 0.00028852 | 1 | 99.454 | 1.3723 | 0.24421 |
| | | Sex | 0.00003671 | 0.00001835 | 2 | 99.47 | 0.0873 | 0.91648 |
| | | Pop.:O2 | 0.0014362 | 0.0014362 | 1 | 32.183 | 6.8312 | 0.01351 |
| | | O2:Sex | 0.00024836 | 0.00012418 | 2 | 99.47 | 0.5907 | 0.55589 |

We also asked whether there were sex differences in gene expression in the labyrinth zone or junctional zone/decidua. Consistent with the absence of sex-specific effects on phenotypic outcomes, we found few genes were differentially expressed between the fetal sexes (18 in the labyrinth zone, and two in the junctional zone/decidua). Including this variable did not substantially impact the number or set of genes included in other categories of interest. Furthermore, higher order models (i.e., including a three-way interact) also did not yield meaningful numbers of genes with sex-specific responses:

| Tissue | Model | Effect | DE genes for model incl. fetal sex term | DE genes for model w/o fetal sex (in main manuscript) |
|---|---|---|---|---|
| LZ | ~ population*O2 + Sex | Sex | 18 | - |
| | | population | 7419 | 7436 |
| | | Hypoxia | 218 | 254 |
| | | population*O2 | 129 | 126 |
| | ~ Fetus + Population + Sex | Sex | 19 | - |
| | | Fetus | 1297 | 1354 |
| | | population | 9311 | 9315 |
| | ~ population*O2*sex | O2*sex | 1 | - |
| | | population*O2*sex | 0 | - |
| | ~ Fetus*sex | Fetus*sex | 3 | - |
| JZ/DEC. | ~ population*O2 + Sex | Sex | 2 | - |
| | | population | 6939 | 6918 |
| | | Hypoxia | 108 | 101 |
| | | population*O2 | 371 | 359 |
| | ~ Fetus + Population + Sex | Sex | 2 | - |
| | | Fetus | 3 | 3 |
| | | population | 8249 | 8271 |
| | ~ population*O2*sex | O2*sex | 0 | - |
| | | population*O2*sex | 31 | - |
| | ~ Fetus*sex | Fetus*sex | 0 | - |

Because we find no effects of sex on phenotypic outcomes of interest **and** few genes show sex-specific patterns of expression or an interaction between sex and hypoxia, we believe sex is of relatively little importance to hypoxia responses that influence fetal growth trajectories during early-to-mid gestation in deer mice. we excluded fetal sex from all analyses and results presented in the main text. Fetal sex information is provided in Datasets 1 and 2

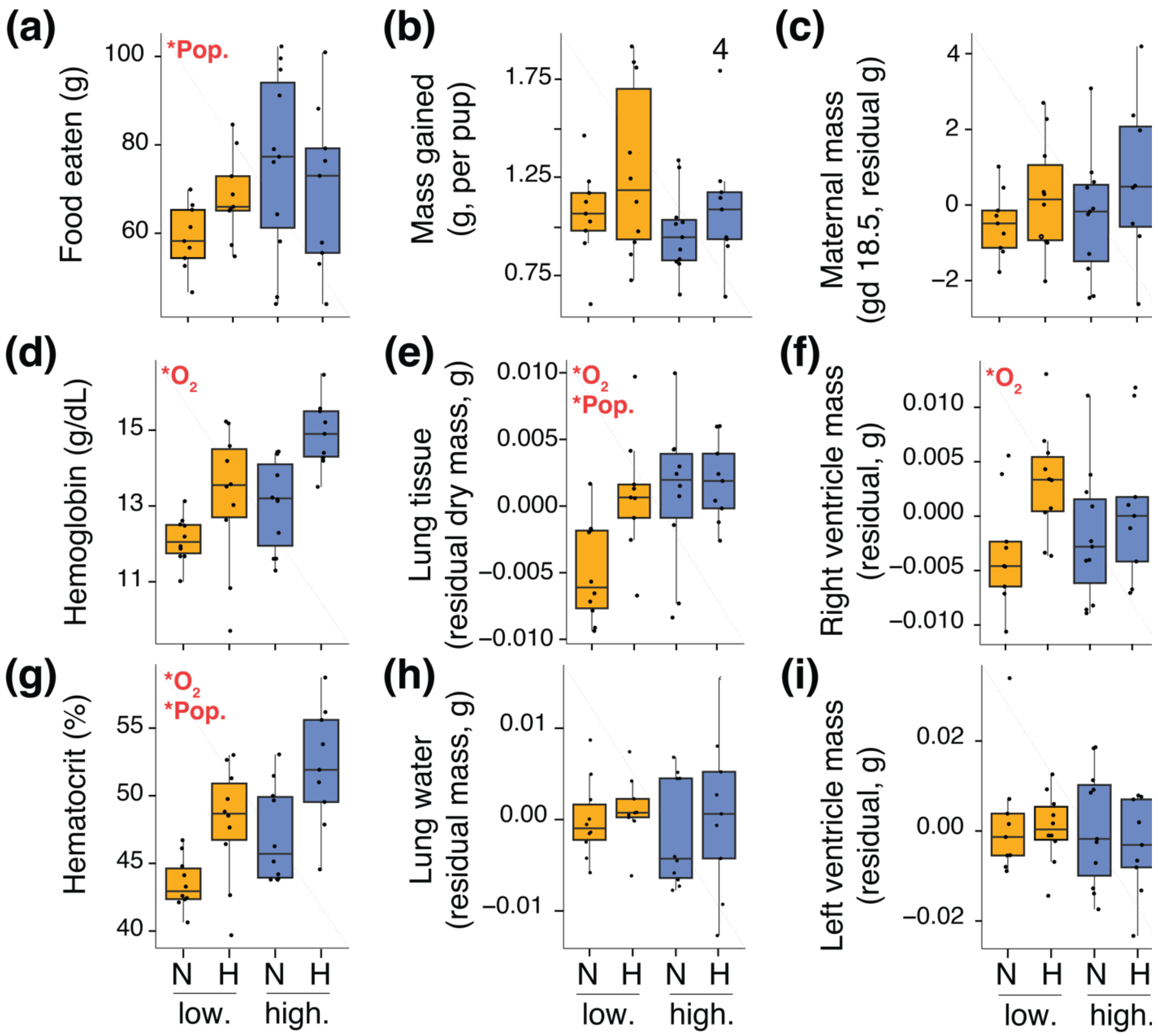

**Figure S1** Plots of maternal trait data. (a) Food eaten was calculated as the difference between food mass in the hopper on gestational day 1 and food in he hopper on gestational day 18.5 (at euthanasia). (b) Mass gained per pup was calculated as the difference between maternal mass on day 1 of gestation and day 18.5 (at euthanasia) divided by the number of pups *in utero*. (c) Residual maternal mass on d18.5 is expressed relative to maternal lean mass on day 1 of gestation and litter size. (d) Maternal hemoglobin at euthanasia on d18.5. (e) Residual dry mass of lung tissue was calculated relative to maternal lean mass on gestational day 1. (f) Residual mass of the right ventricle was calculated relative to maternal lean mass on gestational day 1. (g) Maternal hematocrit at euthanasia on d18.5. (h) Residual mass of water in lung tissue was calculated relative to dry lung tissue mass. (i) Residual mass of the left ventricle was calculated relative to maternal lean mass on gestational day 1. Differences among groups (lowland [low.], highland [high.], normoxia [N], and hypoxia [H]) were determined using linear models including population, hypoxia treatment, and the interaction between the two (see **Extended Materials and Methods, Statistical analyses of maternal, fetal, and placental mass traits**). In all panels, significant main effects are indicated in the top left of each boxplot. Effects were considered significant at $P < 0.05$.

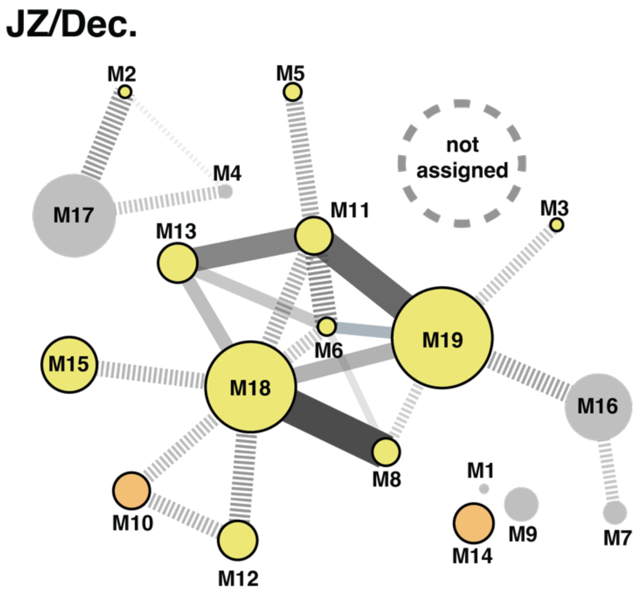

**Figure S2** WGCNA module network illustrating gene expression and module connectivity within the junctional zone/decidua. [JZ/Dec.]. Modules (ellipses) are scaled by the number of genes in each module. Correlations between modules of >0.6 R are shown by edges connecting modules. Width and opacity of edges corresponds to strength of the correlation such that wider and darker edges indicate stronger correlations. Solid edges indicate positive associations, and dashed lines indicated negative correlations. Modules are also colored by their association with outcomes or experimental treatments of interest (fetal mass, red; population-by-hypoxia interactions, orange; hypoxia, green; population, yellow).

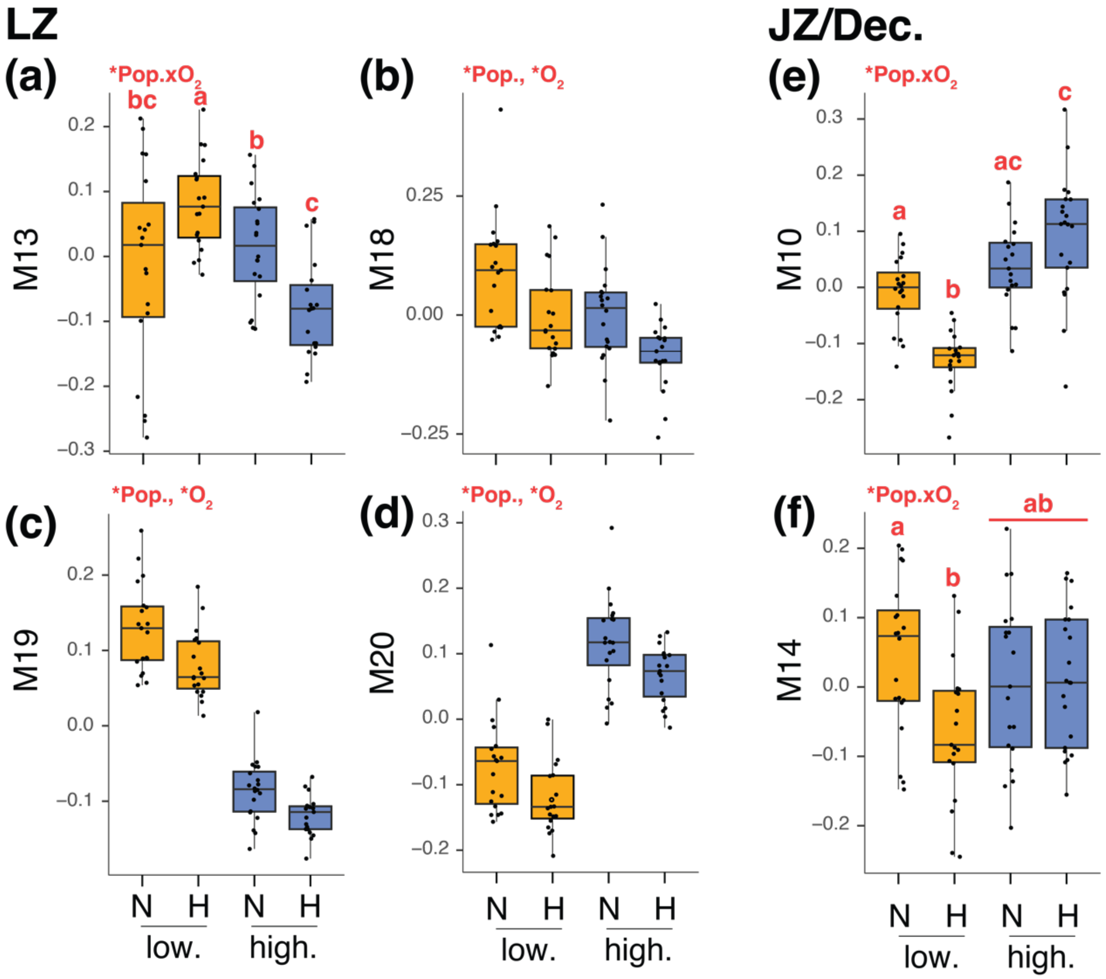

**Figure S3** Plots of eigengenes from the labyrinth zone (a-d) and junctional zone (e,f) modules differentially expressed by hypoxia and population growth (see Fig. 4A) showing differences among treatment groups. Differences among groups (lowland [low.], highland [high.], normoxia [N], and hypoxia [H]) were determined using linear mixed models including population, hypoxia treatment, and the interaction between the two as well as a random effect of dam ID (see **Extended Materials and Methods, Statistical analyses of maternal, fetal, and placental mass traits**). In all panels, presence of a significant interaction or individual main effects are indicated in the top left of each boxplot. Effects were considered significant at $P < 0.05$. For modules with significant interactions, letters indicate statisticalyl significant ($P < 0.05$) group means in a post-hoc tests where p-value adjustments for multiple comparisons were performed using the Benjamini-Hochberg method.

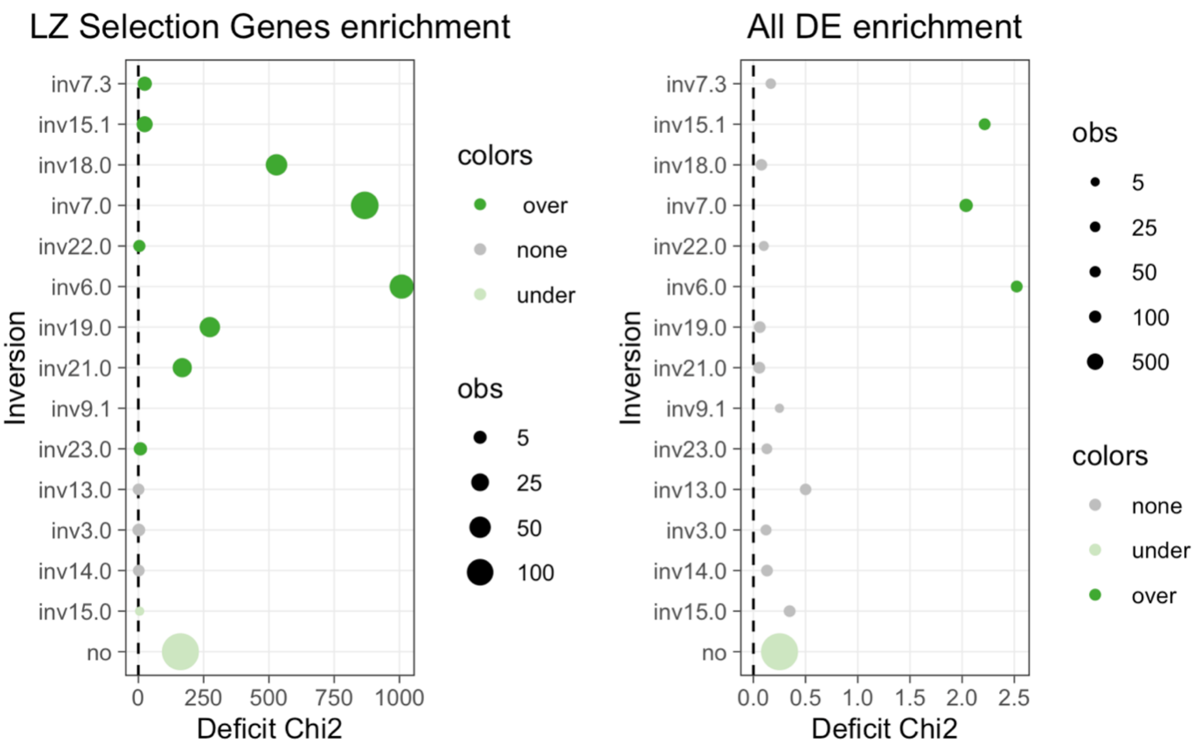

**Figure S4** Enrichment within inversions for gene sets of interest. (**Left**) Enrichment within inversions for targets of selection that are expressed in the labyrinth zone. (**Right**) Enrichment within inversions for genes differentially expressed in any gene set. In each plot, the number of genes within the observed inversion or whole genome (no) is indicated by the size of the circle in that row (larger circles correspond to larger gene sets). Circles falling farther to the right indicate greater significant in a hypergeometic tests (see Table S15). Inversions 7.3 and 15.1 (top) are associated with east/west geographic divides such that inversion frequency differs between highland deer mice and lowland deer mice to the east, whereas inversions 7.0 to 21.0 (below) display frequency differences with lowland deer mouse populations to both east and west. Inversions 9.1, 23.0, 13.0, 3.0, 14.0, and 15.0 do not vary in frequency among populations.

**Table S1** Summary table of output from linear mixed models evaluating effects of treatment (gestational hypoxia vs. normoxia), population (highland vs. lowland deer mice), and litter size on pup weight at birth and mid-gestation. Litter size differs between populations and it impacts birth weight, however it does not affect pup weight at mid-gestation and thus was excluded from mid-gestation models reported in the main text.

| Trait | Model | | Sum Sq | Mean Sq | NumDF | DenDF | F value | Pr(>F) |
|---|---|---|---|---|---|---|---|---|
| Litter size (Birth) | Strain + (1 \| Dam ID) | Population | 35.745 | 35.745 | 1 | 57.524 | 28.16 | 0.000 |
| Pup mass (birth, including litter size) | LitterSize + Treatment*Pop. + (1\|Pair.ID) | LitterSize | 0.088139 | 0.088139 | 1 | 54.479 | 5.5085 | 0.023 |
| | | O2 | 0.087327 | 0.087327 | 1 | 129.315 | 5.4577 | 0.021 |
| | | Population | 0.102182 | 0.102182 | 1 | 83.688 | 6.3861 | 0.013 |
| | | O2:Pop. | 0.114038 | 0.114038 | 1 | 145.688 | 7.1271 | 0.008 |
| Pup mass (birth; without litter size) | Treatment*Pop. + (1\|Pair.ID) | O2 | 0.17596 | 0.17596 | 1 | 157.05 | 10.9797 | 0.001 |
| | | Population | 0.039168 | 0.039168 | 1 | 100.1 | 2.444 | 0.121 |
| | | O2:Pop. | 0.084488 | 0.084488 | 1 | 157.05 | 5.2719 | 0.023 |

| Trait | Model | | Df | Sum Sq | Mean Sq | F value | Pr(>F) |
|---|---|---|---|---|---|---|---|
| Litter Size (Mid-gestation) | Population *Treatment | Population | 1 | 103.027 | 103.027 | 38.6188 | 0.000 |
| | | O2 | 1 | 6.622 | 6.622 | 2.4821 | 0.123 |
| | | O2:Pop. | 1 | 0.487 | 0.487 | 0.1827 | 0.671 |
| | | Residuals | 43 | 114.715 | 2.668 | | |

| Trait | Model | | Sum Sq | Mean Sq | NumDF | DenDF | F value | Pr(>F) |
|---|---|---|---|---|---|---|---|---|
| Pup mass (Mid-gestation) | LitterSize + Pop.*Treatment + (1\|Dam ID) | LitterSize | 0.00116014 | 0.00116014 | 1 | 34.526 | 1.5726 | 0.218 |
| | | Population | 0.00098016 | 0.00098016 | 1 | 33.561 | 1.3287 | 0.257 |
| | | O2 | 0.00155488 | 0.00155488 | 1 | 34.08 | 2.1077 | 0.156 |
| | | O2:Pop. | 0.00220149 | 0.00220149 | 1 | 33.94 | 2.9843 | 0.093 |
| Pup mass (Mid-gestation) | Pop.*Treatment + (1\|Dam ID) | Population | 0.0071627 | 0.0071627 | 1 | 35.131 | 9.7103 | 0.004 |
| | | O2 | 0.0010297 | 0.0010297 | 1 | 35.131 | 1.3959 | 0.245 |
| | | O2:Pop. | 0.0036452 | 0.0036452 | 1 | 35.131 | 4.9417 | 0.033 |
| Pup mass (Mid-gestation) Lowland only | LitterSize + Treatment + (1\|Dam ID) | LitterSize | 0.0011185 | 0.0011185 | 1 | 15.24 | 0.8827 | 0.362 |
| | | O2 | 0.0125277 | 0.0125277 | 1 | 14.733 | 9.8858 | 0.007 |
| Pup mass (Mid-gestation) Highland only | LitterSize + Treatment + (1\|Dam ID) | LitterSize | 0.00045445 | 0.00045445 | 1 | 17.159 | 0.818 | 0.378 |
| | | O2 | 0.00000292 | 0.00000292 | 1 | 16.916 | 0.0053 | 0.943 |

**Table S2** Summary table of statistical tests for effects of population and hypoxia effects on maternal physiology

| Trait | Model | | Sum Sq | Df | F value | Pr(>F) |
|---|---|---|---|---|---|---|
| Food consumed (g) | ~ Population*O2 + GD1_lean | Population | 380.9 | 1 | 1.5698 | 0.21931 |
| | | O2TRT | 382 | 1 | 1.5746 | 0.21863 |
| | | GD1.lean | 495.7 | 1 | 2.0431 | 0.16258 |
| | | Population:O2TRT | 746 | 1 | 3.0748 | 0.08909 |
| F.Mass_gain | ~ Population*O2 + GD1_lean + DissLitter.Size | Population | 0.001 | 1 | 0.0004 | 0.9832 |
| | | O2TRT | 1.653 | 1 | 0.6268 | 0.4344 |
| | | GD1.lean | 0.383 | 1 | 0.1453 | 0.7056 |
| | | DissLitter.Size | 55.407 | 1 | 21.0142 | 6.63E-05 |
| | | Population:O2TRT | 0.261 | 1 | 0.0991 | 0.755 |
| Hemoglobin (Hb) | ~ Population*O2 | Population | 4.41 | 1 | 2.9793 | 0.09291 |
| | | O2TRT | 6.38 | 1 | 4.3159 | 0.04495 |
| | | Population:O2TRT | 1.37 | 1 | 0.9276 | 0.34192 |
| Hematocrit (Hct) | ~ Population*O2 | Population | 65.1 | 1 | 4.8673 | 0.034026 |
| | | O2TRT | 102.8 | 1 | 7.6846 | 0.008863 |
| | | Population:O2TRT | 0.6 | 1 | 0.042 | 0.838745 |
| Heart mass (g) | ~ Population*O2 + GD1_lean | Population | 0 | 1 | 0.0001 | 0.991697 |
| | | O2TRT | 0.0007717 | 1 | 2.747 | 0.106914 |
| | | GD1.lean | 0.0035138 | 1 | 12.5075 | 0.001226 |
| | | Population:O2TRT | 0.0007672 | 1 | 2.7308 | 0.107913 |
| Left ventricle and septum mass (g) | ~ Population*O2 + GD1_lean | Population | 0.0000394 | 1 | 0.2939 | 0.5913794 |
| | | O2TRT | 0.0000076 | 1 | 0.057 | 0.8127574 |
| | | GD1.lean | 0.0021208 | 1 | 15.8071 | 0.0003603 |
| | | Population:O2TRT | 0.0000496 | 1 | 0.3697 | 0.5473296 |
| Right ventricle mass (g) | ~ Population*O2 + GD1_lean | Population | 0.00000064 | 1 | 0.0181 | 0.89382 |
| | | O2TRT | 0.00018801 | 1 | 5.3067 | 0.02768 |
| | | GD1.lean | 0.00006326 | 1 | 1.7855 | 0.19062 |
| | | Population:O2TRT | 0.0000357 | 1 | 1.0077 | 0.32277 |
| Lung (wet mass, g) | ~ Population*O2 + GD1_lean | Population | 0.0048187 | 1 | 17.9062 | 1.74E-04 |
| | | O2TRT | 0.003768 | 1 | 14.0021 | 0.0006955 |
| | | GD1.lean | 0.0029023 | 1 | 10.7849 | 0.002427 |
| | | Population:O2TRT | 0.0003859 | 1 | 1.434 | 0.2396468 |
| Lung tissue (dry mass, g) | ~ Population*O2 + GD1_lean | Population | 0.00033083 | 1 | 23.625 | 2.98E-05 |
| | | O2TRT | 0.00016774 | 1 | 11.9788 | 0.001547 |
| | | GD1.lean | 0.00011594 | 1 | 8.2797 | 0.007084 |
| | | Population:O2TRT | 0.00001624 | 1 | 1.1598 | 0.289568 |
| Lung edema (absolute, g) | ~ Population*O2 + GD1_lean | Population | 0.0026494 | 1 | 15.4945 | 4.19E-04 |
| | | O2TRT | 0.0019239 | 1 | 11.2513 | 0.0020596 |
| | | GD1.lean | 0.0018116 | 1 | 10.5944 | 0.0026804 |
| | | Population:O2TRT | 0.000175 | 1 | 1.0236 | 0.3192599 |
| Lung edema (by tissue mass) | ~ Population*O2 + GD1_lean + LungTissue | Population | 0.0000158 | 1 | 0.4374 | 5.13E-01 |
| | | O2TRT | 0.0000088 | 1 | 0.2453 | 0.6239 |
| | | GD1.lean | 0.0000643 | 1 | 1.7842 | 0.1914 |
| | | LungTissue | 0.0043554 | 1 | 120.9386 | 3.16E-12 |
| | | Population:O2TRT | 0.0000004 | 1 | 0.0119 | 0.9139 |

**Table S3** Summary table with output of statistical tests for association maternal traits with fetal mass

| Trait | Model | Effect | Sum Sq | Df | F value | Pr(>F) | Adj. R2 | DeltaAICc |
|---|---|---|---|---|---|---|---|---|
| Average Pup Mass | ~ Population*O2 | Population | 0.00024 | 1 | 0.0543 | 0.817066 | 0.295 | N/A |
| | | Hypoxia | 0.05421 | 1 | 12.2936 | 0.001237 | | |
| | | PopulationxHypoxia | 0.03802 | 1 | 8.6217 | 0.005758 | | |
| Average pup mass (g) | ~ GD1_residFat*Pop. + GD1_residFat*O2 | GD1_fatCORRRESID | 0.000169 | 1 | 0.0313 | 0.86072 | 0.136 | 11.58 |
| | | Population | 0.031247 | 1 | 5.7815 | 0.02179 | | |
| | | O2TRT | 0.016872 | 1 | 3.1218 | 0.08623 | | |
| | | GD1_fatCORRRESID:Pop. | 0.005037 | 1 | 0.932 | 0.34115 | | |
| | | GD1_fatCORRRESID:O2 | 0.007018 | 1 | 1.2986 | 0.26244 | | |
| | ~ GD18_residFat | GD18_fatCORRRESID | 0.00051 | 1 | 0.089 | 0.76728 | 0.085 | 13.92 |
| | | Population | 0.028862 | 1 | 5.0378 | 0.03142 | | |
| | | O2TRT | 0.018843 | 1 | 3.289 | 0.07858 | | |
| | | GD18_fatCORRRESID:Pop. | 0.000476 | 1 | 0.0831 | 0.77489 | | |
| | | GD18_fatCORRRESID:O2 | 0.000624 | 1 | 0.1089 | 0.74338 | | |
| | ~ GD18_residMass + DissLitter_Size | DissLitter_Size | 0.01501 | 1 | 3.173 | 0.08407 | 0.24 | 8.21 |
| | | F.Mass_GD18.5RESID | 0.011581 | 1 | 2.4481 | 0.12721 | | |
| | | Population | 0.017189 | 1 | 3.6337 | 0.06536 | | |
| | | O2TRT | 0.02694 | 1 | 5.6949 | 0.0229 | | |
| | | F.Mass_GD18.5RESID:Pop. | 0.015003 | 1 | 3.1715 | 0.08414 | | |
| | | F.Mass_GD18.5RESID:O2 | 0.002274 | 1 | 0.4807 | 0.49294 | | |
| | ~ Hb*Pop. + Hb*O2 | Hb | 0.008005 | 1 | 1.6016 | 0.214285 | 0.201 | 8.46 |
| | | Population | 0.013598 | 1 | 2.7206 | 0.108273 | | |
| | | O2TRT | 0.013399 | 1 | 2.6808 | 0.110786 | | |
| | | Hb:Population | 0.017276 | 1 | 3.4564 | 0.071679 | | |
| | | Hb:O2TRT | 0.009668 | 1 | 1.9342 | 0.173337 | | |
| | ~ Hct*Pop. + Hct*O2 | HctAv | 0.018391 | 1 | 3.4643 | 0.071635 | 0.172 | 9.64 |
| | | Population | 0.003371 | 1 | 0.635 | 0.431206 | | |
| | | O2TRT | 0.014383 | 1 | 2.7094 | 0.109254 | | |
| | | HctAv:Population | 0.005799 | 1 | 1.0923 | 0.30356 | | |
| | | HctAv:O2TRT | 0.012538 | 1 | 2.3618 | 0.13387 | | |
| | ~ HeartRESID*Pop. + HeartRESID*O3 | HeartRESID | 0.00023 | 1 | 0.0435 | 0.83608 | 0.165 | 9.88 |
| | | Population | 0.01717 | 1 | 3.2147 | 0.08215 | | |
| | | O2TRT | 0.01064 | 1 | 1.9923 | 0.16746 | | |
| | | HeartRESID:Pop. | 0.00199 | 1 | 0.3723 | 0.54591 | | |
| | | HeartRESID:O2TRT | 0.00935 | 1 | 1.7514 | 0.1948 | | |
| | ~ LV.SeptRESID*Pop. + LV.SeptRESID*O3 | LV.SeptRESID | 0.00014 | 1 | 0.0271 | 0.87031 | 0.205 | 7.96 |
| | | Population | 0.02086 | 1 | 4.1042 | 0.05092 | | |
| | | O2TRT | 0.01695 | 1 | 3.3338 | 0.07693 | | |
| | | LV.SeptRESID:Pop. | 0.00397 | 1 | 0.7809 | 0.38327 | | |
| | | LV.SeptRESID:O2 | 0.02476 | 1 | 4.8716 | 0.03437 | | |
| | ~ RV.RESID*Pop. + RV.RESID*O4 | RVRESID | 0.00871 | 1 | 1.5824 | 0.21724 | 0.14 | 11.05 |
| | | Population | 0.02779 | 1 | 5.0491 | 0.03145 | | |
| | | O2TRT | 0.00444 | 1 | 0.8064 | 0.37569 | | |
| | | RVRESID:Population | 0.0014 | 1 | 0.2553 | 0.61674 | | |
| | | RVRESID:O2TRT | 0.00042 | 1 | 0.0755 | 0.78526 | | |
| | ~ LungTissue.RESID*Pop. + LungTissue.RESID*O2 | LungTissueRESID | 0.01246 | 1 | 2.5642 | 0.11913 | 0.25 | 5.17 |
| | | Population | 0.02325 | 1 | 4.7849 | 0.03613 | | |
| | | O2TRT | 0.01182 | 1 | 2.4332 | 0.12863 | | |
| | | LungTissRESID:Pop | 0.02913 | 1 | 5.995 | 0.02001 | | |
| | | LungTissRESID:O2 | 0.01385 | 1 | 2.8497 | 0.10112 | | |
| | ~ PCAd1 + PCAd2 + PCAd3 + PCAd4 + PCAd5 | PCAd1 | 0.0049 | 1 | 0.8005 | 0.37725 | 0.031 | 16.209 |
| | | PCAd2 | 0.001 | 1 | 0.1611 | 0.6907 | | |
| | | PCAd3 | 0.0001 | 1 | 0.0135 | 0.90807 | | |
| | | PCAd4 | 0.0192 | 1 | 3.1668 | 0.08409 | | |
| | | PCAd5 | 0.0127 | 1 | 2.0927 | 0.15716 | | |

**Table S4** Summary table with output of statistical tests for effects of population and hypoxia on placental mass and association between placental mass and fetal mass.

| Trait | Model | Effect | Sum Sq | Mean Sq | NumDF | DenDF | F value | Pr(>F) |
|---|---|---|---|---|---|---|---|---|
| Placenta mass | ~ Population*Hypoxia | population | 0.00057004 | 0.00057004 | 1 | 33.522 | 2.7061 | 0.109309 |
| | | O2 | 0.00173449 | 0.00173449 | 1 | 33.522 | 8.2341 | 0.007067 |
| | | population:O2 | 0.00161169 | 0.00161169 | 1 | 33.522 | 7.6512 | 0.00916 |
| Fetal mass | ~ PlacentaMass + population*O2 | Placenta | 0.00624 | 0.00624 | 1 | 92.483 | 8.8801 | 0.003683 |
| | | O2 | 0.0000051 | 0.0000051 | 1 | 103.769 | 0.0073 | 0.932082 |
| | | population | 0.0011368 | 0.0011368 | 1 | 105.958 | 1.6178 | 0.206188 |
| | | Placenta:O2 | 0.0001245 | 0.0001245 | 1 | 87.475 | 0.1772 | 0.674866 |
| | | Placenta:pop. | 0.0000553 | 0.0000553 | 1 | 92.492 | 0.0787 | 0.779754 |
| Fetal mass, lowland only | ~ PlacentaMass*O2 | Placenta | 0.00250246 | 0.00250246 | 1 | 32.997 | 1.9519 | 0.1717 |
| | | O2 | 0.00084258 | 0.00084258 | 1 | 33.543 | 0.6572 | 0.4233 |
| | | Placenta:O2 | 0.00003985 | 0.00003985 | 1 | 32.997 | 0.0311 | 0.8611 |
| Fetal mass, highland only | ~ PlacentaMass*O2 | Placenta | 0.00251784 | 0.00251784 | 1 | 59.354 | 4.9054 | 0.03062 |
| | | O2 | 0.00034964 | 0.00034964 | 1 | 74.957 | 0.6812 | 0.4118 |
| | | Placenta:O2 | 0.00029267 | 0.00029267 | 1 | 59.354 | 0.5702 | 0.45317 |

**Table S5** Summary table with output of statistical tests for effects of population and hypoxia effects on placental structure.†

| Trait | Model | fixed effect | Df | Sum Sq | Mean Sq | F value | Pr(>F) |
|---|---|---|---|---|---|---|---|
| Labyrinth zone area (absolute) | ~ population*O2 | population | 1 | 91.86 | 91.857 | 3.8087 | 0.05979 |
| | | Treatment | 1 | 106.79 | 106.792 | 4.4279 | 0.0433 |
| | | population:Treatment | 1 | 13.24 | 13.243 | 0.5491 | 0.46408 |
| | | **contrast** | **estimate** | **SE** | **df** | **t.ratio** | **p.value** |
| | | BW 1N - ME 1N | -2.04 | 2.26 | 32 | -0.904 | 0.4472 |
| | | BW 1N - BW 2H | -2.165 | 2.39 | 32 | -0.907 | 0.4472 |
| | | BW 1N - ME 2H | -6.639 | 2.32 | 32 | -2.868 | 0.0435 |
| | | ME 1N - BW 2H | -0.125 | 2.33 | 32 | -0.054 | 0.9575 |
| | | ME 1N - ME 2H | -4.599 | 2.26 | 32 | -2.038 | 0.1399 |
| | | BW 2H - ME 2H | -4.474 | 2.39 | 32 | -1.875 | 0.1399 |
| Labyrinth zone area (relative) | ~ fetal placenta area + population*O2 | **fixed effect** | **Df** | **Sum Sq** | **Mean Sq** | **F value** | **Pr(>F)** |
| | | fetal placenta | 1 | 804.39 | 804.39 | 257.5642 | 2.87E-16 |
| | | population | 1 | 19.37 | 19.37 | 6.2026 | 0.01852 |
| | | Treatment | 1 | 11.03 | 11.03 | 3.5327 | 0.06992 |
| | | population:Treatment | 1 | 9.39 | 9.39 | 3.0081 | 0.09311 |
| | | **contrast** | **estimate** | **SE** | **df** | **t.ratio** | **p.value** |
| | | BW 1N - ME 1N | -0.6235 | 0.818 | 30 | -0.762 | 0.6386 |
| | | BW 1N - BW 2H | -0.0929 | 0.87 | 30 | -0.107 | 0.9157 |
| | | BW 1N - ME 2H | -2.7977 | 0.891 | 30 | -3.139 | 0.0145 |
| | | ME 1N - BW 2H | 0.5305 | 0.839 | 30 | 0.632 | 0.6386 |
| | | ME 1N - ME 2H | -2.1742 | 0.85 | 30 | -2.557 | 0.0317 |
| | | BW 2H - ME 2H | -2.7048 | 0.889 | 30 | -3.043 | 0.0145 |
| Maternal canal diameter | ~ population*O2 | **fixed effect** | **Df** | **Sum Sq** | **Mean Sq** | **F value** | **Pr(>F)** |
| | | population | 1 | 58946 | 58946 | 7.6645 | 0.00942 |
| | | Treatment | 1 | 37039 | 37039 | 4.8159 | 0.03582 |
| | | population:Treatment | 1 | 2553 | 2553 | 0.3319 | 0.56871 |

| Trait | Model | fixed effect | Sum Sq | Mean Sq | NumDF | DenDF | F value | Pr(>F) |
|---|---|---|---|---|---|---|---|---|
| Bloodspace | ~ population*O2 + (1\|Round) | population | 0.047528 | 0.047528 | 1 | 30.729 | 12.8204 | 0.001164 |
| | | Treatment | 0.013755 | 0.013755 | 1 | 29.68 | 3.7103 | 0.063702 |
| | | population:Treatment | 0.001458 | 0.001458 | 1 | 29.25 | 0.3932 | 0.53551 |

**Table S6** Cross-referenced a priori gene list and log FC association between gene of interest and outcome (Fetal mass, Population, Hypoxia, and the interaction of Population and Hypoxia). Significance was assessed after BH correction for multiple comparisons at P-adj. < 0.05.

| human_geneID | source | mus genename | Pman ensemblID | JZ/DEC | | | | LZ | | | |
|---|---|---|---|---|---|---|---|---|---|---|---|
| | | | | Fetus_lgFC | Pop_lgFC | O2_lgFC | PopxO2_lgFC | Fetus_lgFC | Pop_lgFC | O2_lgFC | PopxO2_lgFC |
| ADAM19 | Gundling 2018 | Adam19 | ENSPEMG00000011893 | NS | -0.8225366 | NS | NS | NS | -0.4789854 | NS | NS |
| ADAMTSL4 | Gundling 2018 | Adamtsl4 | ENSPEMG00000008970 | NS | 0.52455087 | NS | NS | NS | NS | NS | NS |
| Adra1a | Moore 2004 | Arda1a | ENSPEMG00000021686 | not expressed | | | | not expressed | | | |
| Agt | Tissot van Patot, 2012 | Agt | ENSPEMG00000012794 | NS | -0.4964402 | NS | NS | NS | NS | NS | NS |
| AHSP | Gundling 2018 | Ahsp | ENSPEMG00000025167 | not expressed | | | | NS | NS | NS | NS |
| ALDH3B2 | Gundling 2018 | No ensembl ID | | NA | | | | | | | |
| ALOX5AP | Gundling 2018 | Alox5ap | ENSPEMG00000000576 | NS | -0.8922434 | NS | NS | NS | -1.0791945 | NS | NS |
| Ang1 | Zamudio, 2003 | Angpt1 | ENSPEMG00000013024 | not expressed | | | | NS | 0.50168176 | NS | NS |
| Ang2 | Zamudio, 2003 | Angpt2 | ENSPEMG00000012005 | NS | -0.629526 | NS | NS | NS | -1.331118 | NS | NS |
| ANXA1 | Wu et al 2022 | Anxa1 | ENSPEMG00000014478 | NS | -0.3376898 | NS | NS | NS | NS | NS | NS |
| ANXA4 | Wu et al 2022 | Anxa4 | ENSPEMG00000016652 | NS | -0.459921 | NS | NS | NS | NS | NS | NS |
| AOC1 | Wu et al 2022 | Aoc1 | ENSPEMG00000013298 | NS | 1.97050698 | NS | NS | NS | 0.95155407 | NS | NS |
| Arnt | Moore 2004 | Arnt | ENSPEMG00000029682 | NS | NS | NS | NS | NS | -0.2428542 | NS | NS |
| Arnt2 | Moore 2004 | Arnt2 | ENSPEMG00000024032 | NS | NS | NS | NS | 3.24847703 | 0.78349697 | NS | NS |
| Arnt3 | Moore 2004 | Arntl | ENSPEMG00000011047 | NS | NS | NS | NS | NS | NS | NS | NS |
| Atf4 | Vaughan 2020 | Atf4 | ENSMUSG00000042406 | NS | NS | NS | NS | NS | -0.1942357 | NS | NS |
| ATP2C2 | Gundling 2018 | Atp2c2 | ENSPEMG00000014405 | not expressed | | | | NS | NS | NS | NS |
| ATP6V1B1 | Gundling 2018 | Atp6v1b1 | ENSPEMG00000017795 | not expressed | | | | not expressed | | | |
| BTBD16 | Gundling 2018 | Btbd16 | ENSPEMG00000021258 | not expressed | | | | not expressed | | | |
| BZW2 | Wu et al 2022 | Bzw2 | ENSPEMG00000019563 | NS | -0.520953 | NS | NS | -0.8230768 | -0.3517147 | NS | NS |
| Ca4 | Gundling 2018 | Car4 | ENSPEMG00000010937 | NS | 1.28654282 | NS | NS | NS | NS | NS | NS |
| CALM1 | Wu et al 2022 | Calm1 | ENSPEMG00000021298 | NS | NS | NS | NS | NS | -0.271081 | NS | NS |
| CANX | Wu et al 2022 | Canx | ENSPEMG00000005531 | NS | -0.192152 | NS | NS | NS | NS | NS | NS |

| | | | | | | | | | | | |
|---|---|---|---|---|---|---|---|---|---|---|---|
| CBLB | Gundling 2018 | Cblb | ENSPEMG00000013659 | NS | NS | NS | NS | NS | -0.5213263 | NS | NS |
| CCL2 | Gundling 2018 | Ccl2 | ENSPEMG00000024087 | NS | -2.0730159 | NS | NS | NS | -1.3541015 | NS | NS |
| CCL3L3 | Gundling 2018 | No ensembl ID | | NA | | | | | | | |
| CD163 | Gundling 2018 | Cd163 | ENSPEMG00000023074 | NS | 1.24447984 | NS | NS | NS | 2.56377925 | NS | NS |
| CD164 | Wu et al 2022 | Cd164 | ENSPEMG00000025622 | NS | NS | NS | NS | -0.9383056 | -0.4633873 | NS | NS |
| CD46 | Wu et al 2022 | Cd46 | ENSPEMG00000013673 | NS | NS | NS | NS | NS | NS | NS | NS |
| CD59 | Wu et al 2022 | No ensembl ID | | NA | | | | | | | |
| Cdh1 | Bigham 2014 | Cdh1 | ENSPEMG00000011674 | NS | NS | NS | NS | NS | -0.7574266 | NS | NS |
| CEBPB | Wu et al 2022 | No ensembl ID | | NA | | | | | | | |
| CLIC3 | Wu et al 2022 | Clic3 | ENSPEMG00000010344 | not expressed | | | | not expressed | | | |
| Cops5 | Moore 2004 | Cops5 | ENSPEMG00000017987 | NS | NS | NS | NS | NS | NS | NS | NS |
| COX7A1 | Gundling 2018 | Cox7a1 | ENSPEMG00000002515 | not expressed | | | | NS | 1.47426207 | NS | NS |
| Cpvl | Gundling 2018 | Cpvl | ENSPEMG00000018828 | not expressed | | | | not expressed | | | |
| CRIM1 | Wu et al 2022 | Crim1 | ENSPEMG00000015940 | NS | NS | NS | NS | NS | NS | NS | NS |
| CSH2 | Wu et al 2022 | | ENSPEMG00000005403 | NS | NS | NS | NS | 3.17279481 | NS | NS | NS |
| CSHL1 | Wu et al 2022 | No ensembl ID | | NA | | | | | | | |
| CTGF | Wu et al 2022 | Ccn2 | ENSPEMG00000023835 | NS | -0.6536032 | NS | NS | NS | NS | NS | NS |
| CTSC | Gundling 2018 | Ctsc | ENSPEMG00000023091 | NS | -0.7711359 | NS | NS | NS | 0.43910718 | NS | NS |
| Cul2 | Moore 2004 | Cul2 | ENSPEMG00000019784 | NS | -0.1603923 | NS | NS | NS | NS | NS | NS |
| CYR61 | Wu et al 2022 | No ensembl ID | | NA | | | | | | | |
| DAPK1 | Wu et al 2022 | Dapk1 | ENSPEMG00000022627 | NS | -1.0899296 | NS | NS | NS | 1.22978027 | NS | NS |
| DDB1 | Wu et al 2022 | Ddb1 | ENSPEMG00000012241 | NS | NS | NS | NS | NS | -0.1682354 | NS | NS |
| DLG5 | Wu et al 2022 | Dlg5 | ENSPEMG00000000028 | NS | NS | NS | NS | NS | -0.2046704 | NS | NS |
| DNAAF1 | Gundling 2018 | Dnaaf1 | ENSPEMG00000000781 ¬† | not expressed | | | | not expressed | | | |
| DSTYK | Gundling 2018 | Dstyk | ENSPEMG00000022259 | NS | NS | NS | NS | NS | 0.20740463 | NS | NS |
| DYSF | Gundling 2018 | Dysf | ENSPEMG00000008372 | NS | NS | NS | NS | NS | -0.4111289 | NS | NS |
| EBI3 | Wu et al 2022 | Ebi3 | ENSPEMG00000017922 | NS | NS | NS | NS | not expressed | | | |
| Ece | Moore 2004 | Ece1 | ENSPEMG00000017410 | NS | NS | NS | NS | 1.53622273 | -0.3266592 | NS | NS |

| | | | | | | | | | | | |
|---|---|---|---|---|---|---|---|---|---|---|---|
| Edn1 | Tissot van Patot, 2012; Moore 2004 | Edn1 | ENSPEMG00000026888 | NS | 0.80194402 | NS | NS | 2.18884702 | -0.8675983 | NS | NS |
| Ednra | Bigham 2014 | Ednra | ENSPEMG00000013386 | NS | -0.5369974 | NS | NS | NS | -0.5759391 | NS | NS |
| EFHD1 | Wu et al 2022 | Efhd1 | ENSPEMG00000009702 | NS | NS | NS | NS | NS | NS | NS | NS |
| EGFL7 | Wu et al 2022 | Egfl7 | ENSPEMG00000013587 | NS | 0.81052648 | NS | NS | NS | 0.42014854 | NS | NS |
| Egln1 | Bigham 2014, Moore 2004 | Egln1 | ENSPEMG00000003824 | NS | NS | NS | NS | NS | NS | NS | NS |
| Egln2 | Moore 2004 | Egln2 | ENSPEMG00000015706 | NS | 0.3036563 | NS | NS | NS | 0.27760658 | NS | NS |
| Egln3 | Moore 2004 | Egln3 | ENSPEMG00000005061 | NS | -1.4403053 | NS | NS | NS | -2.4329227 | NS | NS |
| EGR1 | Gundling 2018 | Egr1 | ENSPEMG00000027631 | NS | 1.42946415 | NS | NS | NS | NS | NS | NS |
| eIF2a | Moore 2004 | Eif2a | ENSPEMG00000015613 | NS | -0.2520316 | NS | NS | -0.7835603 | NS | NS | NS |
| EIF4H | Wu et al 2022 | Eif4h | ENSPEMG00000018422 | NS | NS | NS | NS | NS | NS | NS | NS |
| Epas1 | Bigham 2014 | Epas1 | ENSPEMG00000020696 | NS | NS | NS | NS | NS | 0.10924167 | NS | NS |
| Epo | Moore 2004 | Epo | ENSPEMG00000011939 | not expressed | | | | NS | NS | NS | NS |
| Epor | Vaughan 2020 | Epor | ENSPEMG00000020463 | NS | NS | NS | NS | NS | NS | NS | NS |
| ESAM | Wu et al 2022 | Esam | ENSPEMG00000012151 | NS | 1.68728502 | NS | NS | NS | 0.21107067 | NS | NS |
| EZR | Wu et al 2022 | Ezr | ENSPEMG00000008621 | NS | 0.71654113 | NS | NS | NS | 0.14704178 | NS | NS |
| F13A1 | Gundling 2018 | F13a1 | ENSPEMG00000016989 | NS | NS | NS | NS | NS | NS | NS | NS |
| F3 | Wu et al 2022 | F3 | ENSPEMG00000005387 | 3.4029882 | 1.08660024 | NS | NS | NS | NS | NS | NS |
| FABP4 | Gundling 2018 | Fabp4 | ENSPEMG00000008716 | NS | -2.4550407 | NS | NS | 5.64323084 | 1.33696906 | NS | NS |
| FAM129B | Wu et al 2022 | No ensembl ID | | NA | | | | | | | |
| FAM46C | Gundling 2018 | No ensembl ID | | NA | | | | | | | |
| FBLN1 | Wu et al 2022 | Fbln1 | ENSPEMG00000022785 | NS | 1.63467653 | NS | NS | NS | -0.5205827 | NS | NS |
| FBN2 | Wu et al 2022 | Fbn2 | ENSPEMG00000007637 | NS | -0.9440439 | NS | NS | NS | -1.2133785 | NS | NS |
| FKBP2 | Wu et al 2022 | No ensembl ID | | NA | | | | | | | |
| Flt1 | Zamudio, 2003 | Flt1 | ENSPEMG00000019000 | NS | 0.87349664 | NS | NS | NS | NS | NS | NS |
| FOS | Gundling 2018 | Fos | ENSPEMG00000025669 | NS | NS | NS | NS | NS | NS | NS | NS |
| FOSB | Gundling 2018 | Fosb | ENSPEMG00000016150 | NS | 0.88515885 | NS | NS | NS | NS | NS | NS |
| FURIN | Wu et al 2022 | Furin | ENSPEMG00000010067 | NS | 0.51586819 | NS | NS | NS | 0.16422074 | 0.26631813 | NS |
| GABARAPL1 | Wu et al 2022 | Gabarapl1 | ENSPEMG00000024648 | NS | NS | NS | NS | NS | NS | NS | NS |

| | | | | | | | | | | | |
|---|---|---|---|---|---|---|---|---|---|---|---|
| GDF15 | Wu et al 2022 | No ensembl ID | | NA | | | | | | | |
| GH2 | Wu et al 2022 | No ensembl ID | | NA | | | | | | | |
| GLDN | Gundling 2018 | Gldn | ENSPEMG00000020076 | not expressed | | | | not expressed | | | |
| Hemoglobin genes | Bigham 2014 | Hba-x | ENSPEMG00000011750 | -17.68449 | 2.65410363 | 2.19815501 | NS | -12.472519 | 1.29098705 | NS | NS |
| | Bigham 2014 | Hba-a1 | ENSPEMG00000024319 | NS | 1.70281384 | NS | NS | NS | 0.62173814 | NS | NS |
| | Bigham 2014 | Hbb-y | ENSPEMG00000027150 | -17.58836 | 1.61121469 | 1.89507689 | NS | -12.066972 | NS | NS | NS |
| | Bigham 2014 | Hba-a1 | ENSPEMG00000028557 | NS | 1.34208097 | NS | NS | NS | NS | NS | NS |
| | Bigham 2014 | Hba-a1 | ENSPEMG00000030709 | NS | 1.83844459 | NS | NS | NS | 1.0571866 | NS | NS |
| | Bigham 2014 | Hbb-bt | ENSPEMG00000030902 | NS | 1.38201485 | NS | NS | NS | 2.18382499 | NS | NS |
| | Bigham 2014 | Hbb-bt | ENSPEMG00000031168 | NS | 0.80794476 | NS | NS | NS | NS | NS | NS |
| GPC3 | Wu et al 2022 | Gpc3 | ENSPEMG00000010163 | NS | -2.8509047 | NS | NS | NS | NS | NS | NS |
| GRN | Wu et al 2022 | Grn | ENSPEMG00000013392 | NS | NS | NS | NS | NS | NS | NS | NS |
| GUCA2A | Gundling 2018 | Guca2a | ENSPEMG00000029117 | NS | NS | NS | NS | NS | NS | NS | NS |
| GYPE | Gundling 2018 | No ensembl ID | | NA | | | | | | | |
| Hif1a | Zamudio, 2003 | Hif1a | ENSPEMG00000021582 | NS | -0.6019963 | NS | NS | NS | 0.14663198 | NS | NS |
| Hif3a | Moore 2004 | Hif3a | ENSPEMG00000017981 | NS | 1.24430516 | NS | NS | 4.91545497 | NS | NS | NS |
| HMGB3 | Wu et al 2022 | No ensembl ID | | NA | | | | | | | |
| HSD17B1 | Wu et al 2022 | Hsd17b1 | ENSPEMG00000010252 | not expressed | | | | not expressed | | | |
| HSPB1 | Wu et al 2022 | Hspb1 | ENSPEMG00000011657 | NS | 1.50044454 | NS | NS | -1.355685 | NS | NS | NS |
| HTRA1 | Wu et al 2022 | Htra1 | ENSPEMG00000022401 | NS | -0.4785226 | NS | NS | NS | -0.8676658 | NS | NS |
| IER3 | Gundling 2018 | Ier3 | ENSPEMG00000030288 | NS | NS | NS | NS | NS | NS | NS | NS |
| IFI30 | Wu et al 2022 | Ifi30 | ENSPEMG00000016976 | NS | -0.7024334 | NS | NS | NS | NS | NS | NS |
| IFIT1-related genes | Gundling 2018 | Ifit1 | ENSPEMG00000000359 | NS | NS | NS | NS | NS | 0.657087 | NS | NS |
| | Gundling 2018 | Ifit1bl1 | ENSPEMG00000014943 | NS | NS | NS | NS | NS | NS | NS | NS |
| | Gundling 2018 | Ifit1 | ENSPEMG00000024487 | NS | NS | NS | NS | NS | 0.63936366 | NS | NS |
| | Gundling 2018 | Ifit1 | ENSPEMG00000025189 | NS | NS | NS | NS | NS | NS | NS | NS |
| | Gundling 2018 | Ifit1 | ENSPEMG00000027303 | not expressed | | | | NS | NS | NS | NS |
| | Gundling 2018 | Ifit1 | ENSPEMG00000029363 | NS | NS | NS | NS | NS | 2.07022977 | NS | NS |

| | | | | | | | | | | | |
|---|---|---|---|---|---|---|---|---|---|---|---|
| Igf2 | Moore 2004 | Igf2 | ENSPEMG00000024839 | NS | NS | NS | NS | NS | -0.3067369 | NS | NS |
| Igf2-related genes | Moore 2004 | Igf2bp1 | ENSPEMG00000021974 | NS | NS | NS | NS | NS | 0.15089657 | NS | NS |
| | Moore 2004 | Igf2bp3 | ENSPEMG00000023450 | NS | NS | NS | NS | NS | NS | NS | NS |
| Igfbp1 | Moore 2004 | Igfbp1 | ENSPEMG00000013560 | not expressed | | | | not expressed | | | |
| Igfbp2 | Moore 2004 | Igfbp2 | ENSPEMG00000018165 | NS | 3.1382841 | NS | NS | NS | 2.24565778 | NS | NS |
| Igfbp3 | Moore 2004 | Igfbp3 | ENSPEMG00000031297 | NS | -1.1071741 | NS | NS | NS | NS | NS | NS |
| Il1 | Moore 2004 | Il1r2 | ENSPEMG00000005448 | NS | NS | NS | NS | NS | -1.0346341 | NS | NS |
| | Moore 2004 | Il1r1 | ENSPEMG00000015111 | NS | NS | NS | NS | NS | -1.0659218 | NS | NS |
| Il1b | Moore 2011 | Il1b | ENSPEMG00000011618 | NS | 2.42338494 | NS | NS | NS | 0.83245063 | NS | NS |
| Il6 | Bigham 2014, Tissot van Patot 2012 | Il6ra | ENSPEMG00000012201 | NS | NS | NS | NS | NS | -0.750268 | NS | NS |
| INHBA | Wu et al 2022 | Inhba | ENSPEMG00000030253 | NS | 0.85076983 | NS | NS | NS | NS | NS | NS |
| ISM2 | Wu et al 2022 | Ism2 | ENSPEMG00000016990 | NS | 1.46801871 | NS | NS | NS | 0.65691693 | NS | NS |
| JUN | Gundling 2018 | Jun | ENSPEMG00000014041 | NS | NS | NS | NS | NS | NS | NS | NS |
| Kcnma1 | Bigham 2014 | Kcnma1 | ENSPEMG00000021200 | not expressed | | | | not expressed | | | |
| Kdr | Moore 2004 | Kdr | ENSPEMG00000018642 | NS | 0.54384724 | NS | NS | 1.34972382 | NS | NS | NS |
| KISS1 | Wu et al 2022 | #N/A | ENSPEMG00000027240 | not expressed | | | | not expressed | | | |
| KLF6 | Wu et al 2022 | Klf6 | ENSPEMG00000008003 | NS | NS | NS | NS | NS | -0.2905701 | NS | NS |
| LAP3 | Gundling 2018 | Lap3 | ENSPEMG00000020769 | NS | -0.5861462 | NS | NS | NS | -0.2810665 | NS | NS |
| LAPTM4A | Wu et al 2022 | Laptm4a | ENSPEMG00000015789 | NS | NS | NS | NS | NS | -0.1801203 | NS | NS |
| LAPTM5 | Gundling 2018 | Laptm5 | ENSPEMG00000016483 | NS | -1.2271455 | NS | NS | NS | -1.5196219 | NS | NS |
| LCOR | Gundling 2018 | Lcor | ENSPEMG00000018115 | NS | NS | NS | NS | NS | NS | NS | NS |
| LGALS13 | Wu et al 2022 | No ensembl ID | | NA | | | | | | | |
| LGALS14 | Wu et al 2022 | No ensembl ID | | NA | | | | | | | |
| LIMCH1 | Wu et al 2022 | Limch1 | ENSPEMG00000025597 | NS | NS | NS | NS | NS | NS | NS | NS |
| Lypd3 | Gundling 2018 | Lypd3 | ENSPEMG00000021920 | NS | -1.4070846 | NS | NS | NS | -1.6102333 | NS | NS |
| LYVE1 | Gundling 2018 | Lyve1 | ENSPEMG00000023759 | NS | -1.0420121 | NS | NS | NS | -0.8697743 | NS | NS |
| MAFF | Wu et al 2022 | Maff | ENSPEMG00000029295 | NS | 0.57952116 | NS | NS | NS | NS | NS | NS |

| | | | | | | | | | | | |
|---|---|---|---|---|---|---|---|---|---|---|---|
| MAN1A2 | Wu et al 2022 | Man1a2 | ENSPEMG00000007670 | NS | -0.1934116 | -0.2392107 | 0.368862377 | NS | -0.1753753 | NS | NS |
| MGAT1 | Wu et al 2022 | Mgat1 | ENSPEMG00000005968 | NS | NS | NS | NS | NS | -0.299101 | NS | NS |
| MLLT1 | Wu et al 2022 | Mllt1 | ENSPEMG00000020708 | NS | 0.11379057 | NS | NS | NS | NS | NS | NS |
| Matrix metalloproteinases | Tissot van Patot, 2012 | Mmp23 | ENSPEMG00000000624 | NS | 0.60254527 | NS | NS | NS | NS | NS | NS |
| | Tissot van Patot, 2012 | Mmp11 | ENSPEMG00000011798 | NS | NS | NS | NS | NS | NS | NS | NS |
| | Tissot van Patot, 2012 | Mmp15 | ENSPEMG00000012320 | NS | 0.90843045 | NS | NS | NS | 0.63342802 | NS | NS |
| | Tissot van Patot, 2012 | Mmp14 | ENSPEMG00000014840 | 2.4033782 | NS | NS | NS | NS | -0.3072212 | NS | NS |
| | Tissot van Patot, 2012 | Mmp19 | ENSPEMG00000015034 | NS | NS | NS | NS | NS | NS | NS | NS |
| | Tissot van Patot, 2012 | Mmp17 | ENSPEMG00000018027 | NS | NS | NS | NS | -2.666939 | 0.71979095 | NS | NS |
| | Tissot van Patot, 2012 | Mmp28 | ENSPEMG00000019080 | NS | -0.7306343 | NS | NS | 1.78262306 | 0.41040894 | NS | NS |
| | Tissot van Patot, 2012 | Mmp12 | ENSPEMG00000020371 | NS | 1.86088741 | NS | NS | NS | NS | NS | NS |
| | Tissot van Patot, 2012 | Mmp1a | ENSPEMG00000020690 | NS | NS | NS | NS | NS | NS | NS | NS |
| | Tissot van Patot, 2012 | Mmp10 | ENSPEMG00000021262 | NS | 1.90291399 | NS | NS | NS | -1.0799097 | NS | NS |
| | Tissot van Patot, 2012 | Mmp2 | ENSPEMG00000023486 | NS | NS | NS | NS | NS | NS | NS | NS |
| | Tissot van Patot, 2012 | Mmp9 | ENSPEMG00000023917 | NS | NS | NS | NS | NS | NS | NS | NS |
| MS4A6A | Gundling 2018 | No ensembl ID | | NA | | | | | | | |
| NAALAD2 | Gundling 2018 | Naalad2 | ENSPEMG00000018071 | not expressed | | | | NS | -1.9610327 | NS | NS |
| Naprt1 | Gundling 2018 | Naprt | ENSPEMG00000001062 | NS | -1.0471317 | NS | NS | NS | -0.3329518 | NS | NS |
| NNAT | Gundling 2018 | Nnat | ENSPEMG00000027306 | NS | -1.1604662 | NS | NS | NS | -1.9424291 | NS | NS |
| Nos2 | Bigham 2014, Moore 2004 | Nos2 | ENSPEMG00000004104 | NS | NS | NS | NS | NS | NS | NS | NS |
| Nos3 | Moore 2004 | Nos3 | ENSPEMG00000018547 | NS | 0.38613617 | NS | NS | 2.11484654 | NS | NS | NS |
| NR4A1 | Wu et al 2022 | Nr4a1 | ENSPEMG00000003687 | NS | NS | NS | NS | NS | -0.3762735 | NS | NS |
| Nrp1 | Moore 2004 | Nrp1 | ENSPEMG00000017771 | NS | -0.4487875 | NS | NS | NS | NS | NS | NS |
| Nrp2 | Moore 2004 | Nrp2 | ENSPEMG00000020944 | NS | -0.543768 | NS | NS | NS | -0.5629475 | NS | NS |
| NUCB2 | Wu et al 2022 | Nucb2 | ENSPEMG00000022928 | NS | -0.7597733 | NS | NS | NS | -0.4335078 | NS | NS |
| OLFML3 | Gundling 2018 | Olfml3 | ENSPEMG00000010380 | NS | 1.54267035 | NS | NS | NS | NS | NS | NS |

| | | | | | | | | | | | |
|---|---|---|---|---|---|---|---|---|---|---|---|
| P13K-AKT pathway | Tissot van Patot, 2012 | Pdk1 | ENSPEMG00000002841 | NS | -0.745318 | NS | NS | NS | NS | NS | NS |
| | Tissot van Patot, 2012 | Akt1s1 | ENSPEMG00000010956 | NS | 0.5512375 | NS | NS | NS | 0.26724498 | 0.33099523 | NS |
| | Tissot van Patot, 2012 | Mtor | ENSPEMG00000013439 | NS | NS | NS | NS | NS | -0.1452651 | NS | NS |
| | Tissot van Patot, 2012 | Akt1 | ENSPEMG00000013949 | NS | NS | NS | NS | NS | NS | NS | NS |
| | Tissot van Patot, 2012 | Pik3ca | ENSPEMG00000018399 | NS | NS | NS | NS | NS | -0.096351 | NS | NS |
| | Tissot van Patot, 2012 | Pten | ENSPEMG00000030393 | NS | -0.5323137 | -0.2714034 | 0.462501868 | NS | -0.2975688 | NS | NS |
| P3h2 | Gundling 2018 | P3h2 | ENSPEMG00000016547 | NS | 1.10233403 | NS | NS | NS | 1.3036785 | NS | NS |
| PAM | Wu et al 2022 | Pam | ENSPEMG00000008719 | NS | NS | NS | NS | NS | -0.464709 | NS | NS |
| PAPPA2 | Wu et al 2022 | Pappa2 | ENSPEMG00000019822 | NS | 2.69213379 | NS | NS | -13.857731 | 1.80104785 | NS | NS |
| Pdgfb | Moore 2004 | Pdgfb | ENSPEMG00000010524 | NS | 0.3653755 | NS | NS | 3.03309883 | 0.36085651 | NS | NS |
| PDPN | Gundling 2018 | Pdpn | ENSPEMG00000007524 | NS | NS | NS | NS | NS | NS | NS | NS |
| Pgf | Zamudio, 2003 | Pgf | ENSPEMG00000017060 | NS | -0.7198726 | NS | NS | NS | -1.2111639 | NS | NS |
| PGRMC2 | Wu et al 2022 | Pgrmc2 | ENSPEMG00000027295 | NS | NS | NS | NS | NS | -0.2859011 | NS | NS |
| PLA2G2A | Gundling 2018 | Novel gene | ENSPEMG00000012135 | NS | NS | NS | NS | not expressed | not expressed | not expressed | not expressed |
| PLAC1 | Wu et al 2022 | Plac1 | ENSPEMG00000022795 | NS | NS | NS | NS | -2.6260725 | NS | NS | NS |
| PLAGL1 | Wu et al 2022 | Plagl1 | ENSPEMG00000031010 | NS | 0.80789965 | NS | NS | 2.74718682 | 0.31727569 | NS | NS |
| PLIN2 | Wu et al 2022 | Plin2 | ENSPEMG00000020242 | 3.5125541 | -1.0972353 | NS | NS | NS | -0.3756259 | NS | NS |
| Polr2a | Bigham 2014 | Polr2a | ENSPEMG00000012521 | NS | NS | NS | NS | NS | -0.2368257 | NS | NS |
| Prkaa1 | Bigham 2014 | Prkaa1 | ENSPEMG00000000574 | NS | NS | NS | NS | -0.8904459 | NS | NS | NS |
| PRKCZ | Wu et al 2022 | Prkcz | ENSPEMG00000009397 | NS | -0.6689883 | NS | NS | 1.22377832 | -0.1628035 | NS | NS |
| PROCR | Gundling 2018 | Procr | ENSPEMG00000022438 | NS | NS | NS | NS | NS | NS | NS | NS |
| PSG1 | Wu et al 2022 | No ensembl ID | | NA | | | | | | | |
| PSG3 | Wu et al 2022 | No ensembl ID | | NA | | | | | | | |
| PSG7 | Wu et al 2022 | No ensembl ID | | NA | | | | | | | |
| PSG8 | Wu et al 2022 | No ensembl ID | | NA | | | | | | | |
| Psmc3-related | Bigham 2014 | Psmc3ip | ENSPEMG00000017694 | NS | NS | NS | NS | NS | 0.65858506 | NS | NS |
| PTGDS | Wu et al 2022 | Ptgds | ENSPEMG00000008581 ¬† | not expressed | | | | not expressed | | | |

| | | | | | | | | | | | |
|---|---|---|---|---|---|---|---|---|---|---|---|
| QSOX1 | Gundling 2018 | Qsox1 | ENSPEMG00000021181 | NS | 0.75672738 | NS | NS | NS | NS | NS | NS |
| RAB11FIP5 | Gundling 2018 | Rab11fip5 | ENSPEMG00000000746 | NS | -0.8345316 | NS | NS | NS | 0.66464097 | NS | NS |
| RBBP6 | Wu et al 2022 | Rbbp6 | ENSPEMG00000025348 | NS | NS | NS | NS | NS | -0.4049111 | NS | NS |
| Rbx1 | Moore 2004 | Rbx1 | ENSPEMG00000015808 | NS | NS | NS | NS | NS | NS | NS | NS |
| RDH13 | Wu et al 2022 | Rdh13 | ENSPEMG00000002264 | NS | -0.4284824 | NS | NS | NS | NS | NS | NS |
| RGS1 | Gundling 2018 | Rgs1 | ENSPEMG00000012482 | NS | NS | NS | NS | not expressed | | | |
| RHBDF2 | Gundling 2018 | Rhbdf2 | ENSPEMG00000011677 | NS | -0.7688178 | NS | NS | NS | -0.7178716 | NS | NS |
| Rhoa | Tissot van Patot, 2012 | Rhoa | ENSPEMG00000020093 | NS | -0.2869575 | NS | 0.349169899 | NS | -0.1034324 | NS | NS |
| RHOBTB1 | Gundling 2018 | Rhobtb1 | ENSPEMG00000022032 | NS | -0.6850366 | NS | NS | NS | NS | NS | NS |
| RHOBTB3 | Wu et al 2022 | Rhobtb3 | ENSPEMG00000011890 | NS | -0.7903548 | NS | NS | NS | -0.4236149 | NS | NS |
| Rock1 | Tissot van Patot, 2012 | Rock1 | ENSPEMG00000021896 | NS | -0.3077339 | -0.2598457 | 0.378004691 | NS | -0.1549989 | NS | NS |
| Rock2 | Tissot van Patot, 2012 | Rock2 | ENSPEMG00000020257 | NS | NS | NS | 0.521032923 | NS | 0.31603227 | NS | NS |
| Rps23 | Gundling 2018 | Rps23 | ENSPEMG00000020618 | NS | NS | NS | NS | NS | NS | NS | NS |
| Rps26 | Gundling 2018 | Rps26 | ENSPEMG00000004286 | NS | NS | NS | NS | NS | 0.25410281 | NS | NS |
| S100A4 | Wu et al 2022 | S100a4 | ENSPEMG00000028277 | NS | NS | NS | NS | NS | NS | NS | NS |
| Satb1 | Bigham 2014 | Satb1 | ENSPEMG00000019744 | NS | 1.14776761 | NS | NS | NS | -0.4749187 | NS | NS |
| SBSPON | Gundling 2018 | Sbspon | ENSPEMG00000017540 | not expressed | | | | 3.29347561 | 0.50643734 | NS | NS |
| SECISBP2L | Gundling 2018 | Secisbp2l | ENSPEMG00000010577 | NS | NS | NS | NS | NS | NS | NS | NS |
| SERPINE2 | Wu et al 2022 | Serpine2 | ENSPEMG00000015844 | NS | -0.5887871 | NS | NS | NS | -0.473417 | NS | NS |
| SLC23A2 | Gundling 2018 | Slc23a2 | ENSPEMG00000024117 | NS | -1.0013891 | NS | NS | -1.4364626 | -0.9856395 | NS | NS |
| Slc2a1 | Zamudio, 2003 | Slc2a1 | ENSPEMG00000023507 | NS | NS | NS | NS | NS | NS | NS | NS |
| Slc38a1 | Vaughan 2020 | Slc38a1 | ENSPEMG00000027302 | NS | 0.72798054 | NS | NS | 1.95859687 | 0.50076685 | NS | NS |
| Slc38a2 | Vaughan 2020 | Slc38a2 | ENSPEMG00000016717 | NS | NS | NS | NS | NS | -0.4726687 | NS | NS |
| Slc38a3 | Vaughan 2020 | Slc38a3 | ENSPEMG00000000409 | not expressed | | | | 2.55275012 | NS | NS | NS |
| SLC43A2 | Wu et al 2022 | Slc43a2 | ENSPEMG00000003908 | NS | NS | NS | NS | 1.82267596 | 0.48851938 | NS | NS |
| SLC44A2 | Wu et al 2022 | Slc44a2 | ENSPEMG00000008320 | NS | NS | NS | NS | NS | NS | NS | NS |
| SLC4A1 | Gundling 2018 | Slc4a1 | ENSPEMG00000001535 | NS | NS | NS | NS | NS | NS | NS | NS |

| | | | | | | | | | | | |
|---|---|---|---|---|---|---|---|---|---|---|---|
| SNORD13 | Gundling 2018 | Snord13 | ENSPEMG00000004317 | not expressed | | | | not expressed | | | |
| SNORD3A | Wu et al 2022 | No ensembl ID | | NA | NA | NA | NA | NA | NA | NA | NA |
| SPINT2 | Wu et al 2022 | Spint2 | ENSPEMG00000018523 | NS | NS | NS | NS | NS | 0.22560085 | NS | NS |
| SPTLC3 | Wu et al 2022 | #N/A | ENSPEMG00000003941 | not expressed | | | | not expressed | | | |
| ST3GAL4 | Wu et al 2022 | St3gal4 | ENSPEMG00000015869 | NS | -1.3678374 | NS | NS | -1.6858307 | NS | NS | NS |
| Stc1 | Gundling 2018 | Stc1 | ENSPEMG00000027424 | NS | NS | NS | NS | not expressed | | | |
| STK40 | Gundling 2018 | Stk40 | ENSPEMG00000010968 | NS | NS | NS | NS | 0.91682883 | 0.15051276 | NS | NS |
| SULT2B1 | Gundling 2018 | Sult2b1 | ENSPEMG00000010051 | not expressed | | | | not expressed | | | |
| TAC3 | Wu et al 2022 | Tac2 | ENSPEMG00000007115 | not expressed | | | | not expressed | | | |
| TECR | Wu et al 2022 | Tecr | ENSPEMG00000008519 | NS | -0.1704561 | NS | -0.275710631 | NS | NS | NS | NS |
| TFPI | Wu et al 2022 | Tfpi | ENSPEMG00000029900 | NS | NS | NS | NS | NS | -0.5383295 | NS | NS |
| TFPI2 | Wu et al 2022 | Tfpi2 | ENSPEMG00000010362 | NS | 1.78698854 | NS | NS | NS | 0.5280382 | NS | NS |
| Tgfa | Bigham 2014, Moore 2004 | Tgfa | ENSPEMG00000018354 | NS | 2.25156824 | NS | NS | not expressed | | | |
| Tgfb and related genes | Tissot van Patot, 2012 | Tgfb2 | ENSPEMG00000014434 | NS | 1.41298198 | NS | NS | NS | -0.5988658 | NS | NS |
| | Tissot van Patot, 2012 | Tgfb1 | ENSPEMG00000016409 | NS | NS | NS | NS | NS | NS | NS | NS |
| | Tissot van Patot, 2012 | Tgfbr2 | ENSPEMG00000019375 | NS | -0.38462 | NS | NS | NS | NS | NS | NS |
| | Tissot van Patot, 2012 | Tgfbr3 | ENSPEMG00000022511 | NS | NS | NS | NS | NS | -0.57291 | NS | NS |
| | Tissot van Patot, 2012 | Tgfb3 | ENSPEMG00000023520 | NS | NS | NS | NS | NS | -0.5712436 | NS | NS |
| | Tissot van Patot, 2012 | Tgfbr1 | ENSPEMG00000027439 | NS | NS | NS | NS | NS | NS | NS | NS |
| TGM2 | Wu et al 2022 | Tgm2 | ENSPEMG00000022951 | NS | NS | NS | NS | NS | -0.5816611 | NS | NS |
| Th | Moore 2004 | Th | ENSPEMG00000018997 | NS | NS | NS | NS | NS | NS | NS | NS |
| Thnsl2 | Gundling 2018 | Thnsl2 | ENSPEMG00000014795 | not expressed | | | | NS | 2.08031582 | NS | NS |
| TIMM10 | Gundling 2018 | Timm10 | ENSPEMG00000028146 | NS | -0.288029 | NS | NS | NS | NS | NS | NS |
| TIMP2 | Wu et al 2022 | Timp2 | ENSPEMG00000026701 | NS | -0.5675464 | NS | NS | NS | -0.8405131 | NS | NS |
| TMED10 | Wu et al 2022 | Tmed10 | ENSPEMG00000014812 | NS | 1.23541508 | NS | NS | NS | 0.98116748 | NS | NS |
| TMEM54 | Wu et al 2022 | Tmem54 | ENSPEMG00000018200 | not expressed | | | | not expressed | | | |
| Tnc | Bigham 2014 | Tnc | ENSPEMG00000012234 | NS | NS | NS | NS | NS | NS | NS | NS |

| | | | | | | | | | | | |
|---|---|---|---|---|---|---|---|---|---|---|---|
| Tnfa | Moore 2004 | Tnf | ENSPEMG00000017232 | NS | NS | NS | NS | not expressed | | | |
| TNFSF10 | Tissot van Patot, 2012 | Tnfsf10 | ENSPEMG00000014291 | NS | 1.9441123 | NS | NS | NS | NS | NS | NS |
| TNFSF6 | Tissot van Patot, 2012 | Fasl | ENSPEMG00000018529 | not expressed | | | | not expressed | | | |
| TPM1 | Wu et al 2022 | Tpm1 | ENSPEMG00000020869 | NS | 0.70900506 | NS | NS | NS | 0.47038994 | NS | NS |
| Trf | Moore 2004 | Trf | ENSPEMG00000003714 | NS | NS | NS | NS | NS | NS | NS | NS |
| TSPAN33 | Gundling 2018 | Tspan33 | ENSPEMG00000020688 | NS | NS | NS | NS | NS | NS | NS | NS |
| TUSC3 | Wu et al 2022 | TUSC3 | ENSPEMG00000021629 | NS | 1.11337029 | NS | NS | NS | 0.3232212 | NS | NS |
| TYROBP | Gundling 2018 | Tyrobp | ENSPEMG00000004463 | NS | -0.8268989 | NS | NS | NS | -0.6057127 | NS | NS |
| Vegfa | Zamudio, 2003 | Vegfa | ENSPEMG00000023496 | NS | NS | NS | NS | 1.48643966 | 0.27316095 | NS | NS |
| Vegfb | Zamudio, 2003 | Vegfb | ENSPEMG00000012050 | NS | 0.75775532 | NS | NS | NS | NS | NS | NS |
| Vhl | Moore 2004 | Vhl | ENSPEMG00000014902 | NS | NS | NS | NS | NS | NS | NS | NS |
| Vsir | Gundling 2018 | Vsir | ENSPEMG00000010452 | NS | 0.80274736 | NS | NS | NS | NS | NS | NS |

**Table S7** GO enrichment results for genes in the LZ for which expression was correlated with fetal mass

| p_value | Term size | Query size | Intersection size | precision | recall | term_id | source | term_name |
|---|---|---|---|---|---|---|---|---|
| 1.45E-04 | 6 | 1297 | 6 | 0.00462606 | 1 | CORUM:122 | CORUM | MCM complex |
| 6.09E-21 | 744 | 1297 | 166 | 0.12798766 | 0.2231183 | GO:0000278 | GO:BP | mitotic cell cycle |
| 8.95E-21 | 628 | 1297 | 148 | 0.11410948 | 0.2356688 | GO:1903047 | GO:BP | mitotic cell cycle process |
| 8.39E-20 | 984 | 1297 | 198 | 0.15265998 | 0.2012195 | GO:0022402 | GO:BP | cell cycle process |
| 6.46E-18 | 1417 | 1297 | 251 | 0.19352352 | 0.1771348 | GO:0007049 | GO:BP | cell cycle |
| 1.66E-17 | 511 | 1297 | 123 | 0.09483423 | 0.2407045 | GO:0051301 | GO:BP | cell division |
| 3.65E-16 | 180 | 1297 | 63 | 0.04857363 | 0.35 | GO:0000819 | GO:BP | sister chromatid segregation |
| 2.36E-15 | 249 | 1297 | 75 | 0.05782575 | 0.3012048 | GO:0140014 | GO:BP | mitotic nuclear division |
| 4.48E-15 | 246 | 1297 | 74 | 0.05705474 | 0.300813 | GO:0098813 | GO:BP | nuclear chromosome segregation |
| 5.89E-14 | 362 | 1297 | 92 | 0.07093292 | 0.2541436 | GO:0000280 | GO:BP | nuclear division |
| 6.99E-14 | 303 | 1297 | 82 | 0.06322282 | 0.2706271 | GO:0007059 | GO:BP | chromosome segregation |
| 1.62E-13 | 154 | 1297 | 54 | 0.04163454 | 0.3506494 | GO:0000070 | GO:BP | mitotic sister chromatid segregation |
| 7.31E-12 | 1134 | 1297 | 196 | 0.15111796 | 0.1728395 | GO:0007010 | GO:BP | cytoskeleton organization |
| 7.85E-12 | 408 | 1297 | 95 | 0.07324595 | 0.2328431 | GO:0048285 | GO:BP | organelle fission |
| 2.21E-09 | 7597 | 1297 | 876 | 0.67540478 | 0.1153087 | GO:0050789 | GO:BP | regulation of biological process |
| 3.02E-09 | 7270 | 1297 | 844 | 0.65073246 | 0.1160935 | GO:0050794 | GO:BP | regulation of cellular process |
| 1.90E-08 | 8071 | 1297 | 916 | 0.70624518 | 0.1134928 | GO:0065007 | GO:BP | biological regulation |
| 2.69E-08 | 505 | 1297 | 101 | 0.07787201 | 0.2 | GO:0000226 | GO:BP | microtubule cytoskeleton organization |
| 3.34E-08 | 1919 | 1297 | 278 | 0.21434079 | 0.1448671 | GO:0051128 | GO:BP | regulation of cellular component organization |
| 1.68E-07 | 872 | 1297 | 148 | 0.11410948 | 0.1697248 | GO:0051726 | GO:BP | regulation of cell cycle |
| 2.08E-07 | 4778 | 1297 | 585 | 0.45104086 | 0.1224362 | GO:0051716 | GO:BP | cellular response to stimulus |
| 3.02E-07 | 576 | 1297 | 108 | 0.08326908 | 0.1875 | GO:0010564 | GO:BP | regulation of cell cycle process |
| 4.89E-07 | 4042 | 1297 | 506 | 0.39013107 | 0.1251856 | GO:0048856 | GO:BP | anatomical structure development |
| 5.99E-07 | 133 | 1297 | 40 | 0.0308404 | 0.3007519 | GO:1902850 | GO:BP | microtubule cytoskeleton organization involved in mitosis |
| 7.19E-07 | 842 | 1297 | 142 | 0.10948342 | 0.1686461 | GO:0051276 | GO:BP | chromosome organization |
| 8.57E-07 | 3482 | 1297 | 445 | 0.34309946 | 0.1278001 | GO:0007275 | GO:BP | multicellular organism development |
| 1.10E-06 | 29 | 1297 | 17 | 0.01310717 | 0.5862069 | GO:0006270 | GO:BP | DNA replication initiation |
| 1.75E-06 | 971 | 1297 | 157 | 0.12104857 | 0.161689 | GO:0033043 | GO:BP | regulation of organelle organization |
| 2.15E-06 | 4691 | 1297 | 570 | 0.43947571 | 0.1215093 | GO:0016043 | GO:BP | cellular component organization |
| 2.29E-06 | 167 | 1297 | 45 | 0.03469545 | 0.2694611 | GO:0007051 | GO:BP | spindle organization |
| 2.94E-06 | 3130 | 1297 | 404 | 0.31148805 | 0.1290735 | GO:0048731 | GO:BP | system development |
| 3.62E-06 | 3061 | 1297 | 396 | 0.30531997 | 0.1293695 | GO:0006996 | GO:BP | organelle organization |
| 5.64E-06 | 51 | 1297 | 22 | 0.01696222 | 0.4313725 | GO:0007062 | GO:BP | sister chromatid cohesion |
| 5.89E-06 | 4370 | 1297 | 534 | 0.41171935 | 0.1221968 | GO:0032502 | GO:BP | developmental process |
| 7.83E-06 | 5626 | 1297 | 662 | 0.51040864 | 0.117668 | GO:0050896 | GO:BP | response to stimulus |
| 1.75E-05 | 108 | 1297 | 33 | 0.02544333 | 0.3055556 | GO:0007052 | GO:BP | mitotic spindle organization |

| | | | | | | | | |
|---|---|---|---|---|---|---|---|---|
| 1.81E-05 | 208 | 1297 | 50 | 0.0385505 | 0.2403846 | GO:0090068 | GO:BP | positive regulation of cell cycle process |
| 5.37E-05 | 690 | 1297 | 116 | 0.08943716 | 0.1681159 | GO:0007017 | GO:BP | microtubule-based process |
| 7.13E-05 | 1961 | 1297 | 267 | 0.20585968 | 0.136155 | GO:0035556 | GO:BP | intracellular signal transduction |
| 1.22E-04 | 4868 | 1297 | 577 | 0.44487278 | 0.1185292 | GO:0071840 | GO:BP | cellular component organization or biogenesis |
| 1.34E-04 | 2027 | 1297 | 273 | 0.21048574 | 0.1346818 | GO:0009653 | GO:BP | anatomical structure morphogenesis |
| 1.60E-04 | 3903 | 1297 | 476 | 0.36700077 | 0.1219575 | GO:0023052 | GO:BP | signaling |
| 1.78E-04 | 3963 | 1297 | 482 | 0.37162683 | 0.121625 | GO:0007154 | GO:BP | cell communication |
| 2.27E-04 | 1852 | 1297 | 252 | 0.19429453 | 0.1360691 | GO:0050793 | GO:BP | regulation of developmental process |
| 2.33E-04 | 185 | 1297 | 44 | 0.03392444 | 0.2378378 | GO:0007160 | GO:BP | cell-matrix adhesion |
| 3.21E-04 | 4614 | 1297 | 548 | 0.42251349 | 0.118769 | GO:0032501 | GO:BP | multicellular organismal process |
| 3.41E-04 | 132 | 1297 | 35 | 0.02698535 | 0.2651515 | GO:0006261 | GO:BP | DNA-templated DNA replication |
| 3.61E-04 | 379 | 1297 | 72 | 0.05551272 | 0.1899736 | GO:0007264 | GO:BP | small GTPase mediated signal transduction |
| 3.69E-04 | 3594 | 1297 | 441 | 0.34001542 | 0.1227045 | GO:0007165 | GO:BP | signal transduction |
| 6.44E-04 | 458 | 1297 | 82 | 0.06322282 | 0.1790393 | GO:0051493 | GO:BP | regulation of cytoskeleton organization |
| 6.84E-04 | 251 | 1297 | 53 | 0.04086353 | 0.2111554 | GO:0006260 | GO:BP | DNA replication |
| 7.51E-04 | 286 | 1297 | 58 | 0.04471858 | 0.2027972 | GO:0045787 | GO:BP | positive regulation of cell cycle |
| 8.05E-04 | 167 | 1297 | 40 | 0.0308404 | 0.239521 | GO:0046578 | GO:BP | regulation of Ras protein signal transduction |
| 1.85E-03 | 600 | 1297 | 99 | 0.07632999 | 0.165 | GO:0051129 | GO:BP | negative regulation of cellular component organization |
| 2.08E-03 | 891 | 1297 | 135 | 0.10408635 | 0.1515152 | GO:0043085 | GO:BP | positive regulation of catalytic activity |
| 2.83E-03 | 2909 | 1297 | 362 | 0.27910563 | 0.1244414 | GO:0030154 | GO:BP | cell differentiation |
| 3.22E-03 | 2931 | 1297 | 364 | 0.28064765 | 0.1241897 | GO:0048869 | GO:BP | cellular developmental process |
| 3.44E-03 | 202 | 1297 | 44 | 0.03392444 | 0.2178218 | GO:0051056 | GO:BP | regulation of small GTPase mediated signal transduction |
| 3.85E-03 | 91 | 1297 | 26 | 0.02004626 | 0.2857143 | GO:0051304 | GO:BP | chromosome separation |
| 4.17E-03 | 395 | 1297 | 71 | 0.05474171 | 0.1797468 | GO:0007346 | GO:BP | regulation of mitotic cell cycle |
| 4.41E-03 | 158 | 1297 | 37 | 0.02852737 | 0.2341772 | GO:1903046 | GO:BP | meiotic cell cycle process |
| 4.66E-03 | 294 | 1297 | 57 | 0.04394757 | 0.1938776 | GO:0007265 | GO:BP | Ras protein signal transduction |
| 5.20E-03 | 36 | 1297 | 15 | 0.01156515 | 0.4166667 | GO:0008608 | GO:BP | attachment of spindle microtubules to kinetochore |
| 5.66E-03 | 87 | 1297 | 25 | 0.01927525 | 0.2873563 | GO:0051983 | GO:BP | regulation of chromosome segregation |
| 6.14E-03 | 141 | 1297 | 34 | 0.02621434 | 0.2411348 | GO:0051302 | GO:BP | regulation of cell division |
| 7.31E-03 | 142 | 1297 | 34 | 0.02621434 | 0.2394366 | GO:0140013 | GO:BP | meiotic nuclear division |
| 7.56E-03 | 1233 | 1297 | 173 | 0.13338473 | 0.1403082 | GO:0006468 | GO:BP | protein phosphorylation |
| 7.77E-03 | 2227 | 1297 | 285 | 0.21973786 | 0.1279749 | GO:0022607 | GO:BP | cellular component assembly |
| 7.88E-03 | 912 | 1297 | 135 | 0.10408635 | 0.1480263 | GO:0048646 | GO:BP | anatomical structure formation involved in morphogenesis |
| 8.38E-03 | 1478 | 1297 | 201 | 0.15497301 | 0.1359946 | GO:0016310 | GO:BP | phosphorylation |
| 1.00E-02 | 1204 | 1297 | 169 | 0.13030069 | 0.1403654 | GO:0044093 | GO:BP | positive regulation of molecular function |

**Table S8** GO enrichment results for genes in the LZ significantly positively correlated with fetal growth outcomes

| p_value | Term size | Query size | Intersection size | precision | recall | term_id | source | term_name |
|---|---|---|---|---|---|---|---|---|
| 3.07E-11 | 2027 | 750 | 197 | 0.26266667 | 0.09718796 | GO:0009653 | GO:BP | anatomical structure morphogenesis |
| 2.06E-10 | 3903 | 750 | 319 | 0.42533333 | 0.081732 | GO:0023052 | GO:BP | signaling |
| 3.25E-10 | 3963 | 750 | 322 | 0.42933333 | 0.08125158 | GO:0007154 | GO:BP | cell communication |
| 2.93E-09 | 961 | 750 | 111 | 0.148 | 0.11550468 | GO:0007155 | GO:BP | cell adhesion |
| 8.66E-09 | 3594 | 750 | 293 | 0.39066667 | 0.08152476 | GO:0007165 | GO:BP | signal transduction |
| 1.67E-07 | 1140 | 750 | 120 | 0.16 | 0.10526316 | GO:0016477 | GO:BP | cell migration |
| 4.01E-07 | 4042 | 750 | 314 | 0.41866667 | 0.07768431 | GO:0048856 | GO:BP | anatomical structure development |
| 5.40E-07 | 185 | 750 | 36 | 0.048 | 0.19459459 | GO:0007160 | GO:BP | cell-matrix adhesion |
| 6.92E-07 | 518 | 750 | 68 | 0.09066667 | 0.13127413 | GO:0048514 | GO:BP | blood vessel morphogenesis |
| 9.20E-07 | 735 | 750 | 86 | 0.11466667 | 0.1170068 | GO:0035239 | GO:BP | tube morphogenesis |
| 9.53E-07 | 4477 | 750 | 339 | 0.452 | 0.07572035 | GO:0051179 | GO:BP | localization |
| 1.05E-06 | 912 | 750 | 100 | 0.13333333 | 0.10964912 | GO:0048646 | GO:BP | anatomical structure formation involved in morphogenesis |
| 1.10E-06 | 535 | 750 | 69 | 0.092 | 0.12897196 | GO:0034330 | GO:BP | cell junction organization |
| 1.39E-06 | 3130 | 750 | 254 | 0.33866667 | 0.08115016 | GO:0048731 | GO:BP | system development |
| 1.40E-06 | 904 | 750 | 99 | 0.132 | 0.10951327 | GO:0035295 | GO:BP | tube development |
| 1.97E-06 | 4370 | 750 | 331 | 0.44133333 | 0.07574371 | GO:0032502 | GO:BP | developmental process |
| 2.20E-06 | 167 | 750 | 33 | 0.044 | 0.19760479 | GO:0046578 | GO:BP | regulation of Ras protein signal transduction |
| 2.78E-06 | 1340 | 750 | 131 | 0.17466667 | 0.09776119 | GO:0040011 | GO:BP | locomotion |
| 3.37E-06 | 754 | 750 | 86 | 0.11466667 | 0.11405836 | GO:0022603 | GO:BP | regulation of anatomical structure morphogenesis |
| 3.52E-06 | 1236 | 750 | 123 | 0.164 | 0.09951456 | GO:0048870 | GO:BP | cell motility |
| 3.52E-06 | 1236 | 750 | 123 | 0.164 | 0.09951456 | GO:0051674 | GO:BP | localization of cell |
| 3.98E-06 | 622 | 750 | 75 | 0.1 | 0.12057878 | GO:0001944 | GO:BP | vasculature development |
| 4.50E-06 | 4778 | 750 | 354 | 0.472 | 0.07408958 | GO:0051716 | GO:BP | cellular response to stimulus |
| 4.89E-06 | 3482 | 750 | 274 | 0.36533333 | 0.07869041 | GO:0007275 | GO:BP | multicellular organism development |
| 5.15E-06 | 439 | 750 | 59 | 0.07866667 | 0.13439636 | GO:0001525 | GO:BP | angiogenesis |
| 5.18E-06 | 8071 | 750 | 541 | 0.72133333 | 0.06703011 | GO:0065007 | GO:BP | biological regulation |
| 7.17E-06 | 594 | 750 | 72 | 0.096 | 0.12121212 | GO:0001568 | GO:BP | blood vessel development |
| 9.44E-06 | 379 | 750 | 53 | 0.07066667 | 0.13984169 | GO:0007264 | GO:BP | small GTPase mediated signal transduction |
| 1.21E-05 | 1852 | 750 | 165 | 0.22 | 0.08909287 | GO:0050793 | GO:BP | regulation of developmental process |
| 1.58E-05 | 1530 | 750 | 142 | 0.18933333 | 0.09281046 | GO:0006928 | GO:BP | movement of cell or subcellular component |
| 1.92E-05 | 4614 | 750 | 341 | 0.45466667 | 0.0739055 | GO:0032501 | GO:BP | multicellular organismal process |
| 2.09E-05 | 301 | 750 | 45 | 0.06 | 0.14950166 | GO:0031589 | GO:BP | cell-substrate adhesion |
| 2.58E-05 | 202 | 750 | 35 | 0.04666667 | 0.17326733 | GO:0051056 | GO:BP | regulation of small GTPase mediated signal transduction |
| 3.04E-05 | 294 | 750 | 44 | 0.05866667 | 0.14965986 | GO:0007265 | GO:BP | Ras protein signal transduction |
| 3.13E-05 | 5626 | 750 | 400 | 0.53333333 | 0.07109847 | GO:0050896 | GO:BP | response to stimulus |

| | | | | | | | | |
|---|---|---|---|---|---|---|---|---|
| 3.28E-05 | 48 | 750 | 16 | 0.02133333 | 0.33333333 | GO:0046580 | GO:BP | negative regulation of Ras protein signal transduction |
| 3.71E-05 | 921 | 750 | 96 | 0.128 | 0.10423453 | GO:0072359 | GO:BP | circulatory system development |
| 4.15E-05 | 582 | 750 | 69 | 0.092 | 0.1185567 | GO:0030155 | GO:BP | regulation of cell adhesion |
| 5.71E-05 | 300 | 750 | 44 | 0.05866667 | 0.14666667 | GO:0034329 | GO:BP | cell junction assembly |
| 7.24E-05 | 639 | 750 | 73 | 0.09733333 | 0.114241 | GO:0030029 | GO:BP | actin filament-based process |
| 8.60E-05 | 1961 | 750 | 169 | 0.22533333 | 0.08618052 | GO:0035556 | GO:BP | intracellular signal transduction |
| 1.01E-04 | 7597 | 750 | 509 | 0.67866667 | 0.06700013 | GO:0050789 | GO:BP | regulation of biological process |
| 1.53E-04 | 7270 | 750 | 490 | 0.65333333 | 0.06740028 | GO:0050794 | GO:BP | regulation of cellular process |
| 1.67E-04 | 2444 | 750 | 200 | 0.26666667 | 0.08183306 | GO:0023051 | GO:BP | regulation of signaling |
| 1.74E-04 | 2099 | 750 | 177 | 0.236 | 0.08432587 | GO:0032879 | GO:BP | regulation of localization |
| 2.22E-04 | 54 | 750 | 16 | 0.02133333 | 0.2962963 | GO:0051058 | GO:BP | negative regulation of small GTPase mediated signal transduction |
| 2.97E-04 | 2909 | 750 | 229 | 0.30533333 | 0.07872121 | GO:0030154 | GO:BP | cell differentiation |
| 3.17E-04 | 526 | 750 | 62 | 0.08266667 | 0.11787072 | GO:0071363 | GO:BP | cellular response to growth factor stimulus |
| 3.60E-04 | 2931 | 750 | 230 | 0.30666667 | 0.07847151 | GO:0048869 | GO:BP | cellular developmental process |
| 4.78E-04 | 581 | 750 | 66 | 0.088 | 0.11359725 | GO:0030036 | GO:BP | actin cytoskeleton organization |
| 5.20E-04 | 825 | 750 | 85 | 0.11333333 | 0.1030303 | GO:0000902 | GO:BP | cell morphogenesis |
| 6.39E-04 | 2167 | 750 | 179 | 0.23866667 | 0.08260268 | GO:0009966 | GO:BP | regulation of signal transduction |
| 6.56E-04 | 464 | 750 | 56 | 0.07466667 | 0.12068966 | GO:0030335 | GO:BP | positive regulation of cell migration |
| 7.45E-04 | 1977 | 750 | 166 | 0.22133333 | 0.0839656 | GO:0051239 | GO:BP | regulation of multicellular organismal process |
| 7.60E-04 | 702 | 750 | 75 | 0.1 | 0.10683761 | GO:0007167 | GO:BP | enzyme-linked receptor protein signaling pathway |
| 7.90E-04 | 846 | 750 | 86 | 0.11466667 | 0.10165485 | GO:0051270 | GO:BP | regulation of cellular component movement |
| 8.56E-04 | 2434 | 750 | 196 | 0.26133333 | 0.08052588 | GO:0010646 | GO:BP | regulation of cell communication |
| 1.06E-03 | 812 | 750 | 83 | 0.11066667 | 0.10221675 | GO:0040012 | GO:BP | regulation of locomotion |
| 1.08E-03 | 365 | 750 | 47 | 0.06266667 | 0.12876712 | GO:0003013 | GO:BP | circulatory system process |
| 1.08E-03 | 747 | 750 | 78 | 0.104 | 0.10441767 | GO:0030334 | GO:BP | regulation of cell migration |
| 1.15E-03 | 545 | 750 | 62 | 0.08266667 | 0.11376147 | GO:0070848 | GO:BP | response to growth factor |
| 1.33E-03 | 2525 | 750 | 201 | 0.268 | 0.07960396 | GO:0048513 | GO:BP | animal organ development |
| 1.36E-03 | 1073 | 750 | 102 | 0.136 | 0.09506058 | GO:2000026 | GO:BP | regulation of multicellular organismal development |
| 1.52E-03 | 2793 | 750 | 218 | 0.29066667 | 0.07805227 | GO:0048583 | GO:BP | regulation of response to stimulus |
| 1.91E-03 | 479 | 750 | 56 | 0.07466667 | 0.11691023 | GO:2000147 | GO:BP | positive regulation of cell motility |
| 2.05E-03 | 785 | 750 | 80 | 0.10666667 | 0.10191083 | GO:2000145 | GO:BP | regulation of cell motility |
| 2.12E-03 | 103 | 750 | 21 | 0.028 | 0.2038835 | GO:0001952 | GO:BP | regulation of cell-matrix adhesion |
| 2.20E-03 | 1015 | 750 | 97 | 0.12933333 | 0.0955665 | GO:0051094 | GO:BP | positive regulation of developmental process |
| 2.47E-03 | 42 | 750 | 13 | 0.01733333 | 0.30952381 | GO:0048010 | GO:BP | vascular endothelial growth factor receptor signaling pathway |
| 2.67E-03 | 1836 | 750 | 154 | 0.20533333 | 0.083878 | GO:0007166 | GO:BP | cell surface receptor signaling pathway |
| 3.66E-03 | 89 | 750 | 19 | 0.02533333 | 0.21348315 | GO:0150115 | GO:BP | cell-substrate junction organization |

**Table S9** GO enrichment results for genes in the LZ significantly negatively correlated with fetal growth outcomes

| p_value | Term size | Query Size | Intersection size | precision | recall | term_id | source | term_name |
|---|---|---|---|---|---|---|---|---|
| 3.97E-07 | 6 | 547 | 6 | 0.01096892 | 1 | CORUM:122 | CORUM | MCM complex |
| 8.93E-04 | 14 | 547 | 6 | 0.01096892 | 0.42857143 | CORUM:38 | CORUM | 20S proteasome |
| 4.85E-41 | 1417 | 547 | 178 | 0.32541133 | 0.1256175 | GO:0007049 | GO:BP | cell cycle |
| 1.01E-38 | 984 | 547 | 143 | 0.26142596 | 0.1453252 | GO:0022402 | GO:BP | cell cycle process |
| 2.41E-35 | 744 | 547 | 119 | 0.21755027 | 0.15994624 | GO:0000278 | GO:BP | mitotic cell cycle |
| 2.63E-35 | 628 | 547 | 109 | 0.19926874 | 0.17356688 | GO:1903047 | GO:BP | mitotic cell cycle process |
| 9.64E-35 | 303 | 547 | 76 | 0.13893967 | 0.25082508 | GO:0007059 | GO:BP | chromosome segregation |
| 1.31E-34 | 246 | 547 | 69 | 0.1261426 | 0.2804878 | GO:0098813 | GO:BP | nuclear chromosome segregation |
| 2.41E-33 | 180 | 547 | 59 | 0.10786106 | 0.32777778 | GO:0000819 | GO:BP | sister chromatid segregation |
| 1.37E-32 | 842 | 547 | 123 | 0.22486289 | 0.14608076 | GO:0051276 | GO:BP | chromosome organization |
| 4.04E-31 | 511 | 547 | 93 | 0.17001828 | 0.18199609 | GO:0051301 | GO:BP | cell division |
| 7.04E-30 | 362 | 547 | 77 | 0.14076782 | 0.21270718 | GO:0000280 | GO:BP | nuclear division |
| 1.46E-27 | 154 | 547 | 50 | 0.09140768 | 0.32467532 | GO:0000070 | GO:BP | mitotic sister chromatid segregation |
| 1.71E-27 | 249 | 547 | 62 | 0.11334552 | 0.24899598 | GO:0140014 | GO:BP | mitotic nuclear division |
| 5.95E-27 | 408 | 547 | 78 | 0.14259598 | 0.19117647 | GO:0048285 | GO:BP | organelle fission |
| 2.48E-18 | 872 | 547 | 102 | 0.18647166 | 0.11697248 | GO:0051726 | GO:BP | regulation of cell cycle |
| 1.77E-16 | 835 | 547 | 96 | 0.17550274 | 0.11497006 | GO:0006259 | GO:BP | DNA metabolic process |
| 2.37E-16 | 576 | 547 | 77 | 0.14076782 | 0.13368056 | GO:0010564 | GO:BP | regulation of cell cycle process |
| 3.92E-16 | 3061 | 547 | 222 | 0.40585009 | 0.07252532 | GO:0006996 | GO:BP | organelle organization |
| 2.18E-15 | 251 | 547 | 48 | 0.08775137 | 0.19123506 | GO:0006260 | GO:BP | DNA replication |
| 1.54E-14 | 715 | 547 | 84 | 0.1535649 | 0.11748252 | GO:0006974 | GO:BP | cellular response to DNA damage stimulus |
| 9.00E-14 | 51 | 547 | 22 | 0.04021938 | 0.43137255 | GO:0007062 | GO:BP | sister chromatid cohesion |
| 1.24E-13 | 505 | 547 | 67 | 0.12248629 | 0.13267327 | GO:0000226 | GO:BP | microtubule cytoskeleton organization |
| 1.89E-13 | 167 | 547 | 37 | 0.06764168 | 0.22155689 | GO:0007051 | GO:BP | spindle organization |
| 2.13E-13 | 132 | 547 | 33 | 0.06032907 | 0.25 | GO:0006261 | GO:BP | DNA-templated DNA replication |
| 1.39E-12 | 158 | 547 | 35 | 0.06398537 | 0.22151899 | GO:1903046 | GO:BP | meiotic cell cycle process |
| 1.70E-12 | 465 | 547 | 62 | 0.11334552 | 0.13333333 | GO:0006281 | GO:BP | DNA repair |
| 3.15E-12 | 201 | 547 | 39 | 0.07129799 | 0.19402985 | GO:0051321 | GO:BP | meiotic cell cycle |
| 1.05E-11 | 208 | 547 | 39 | 0.07129799 | 0.1875 | GO:0090068 | GO:BP | positive regulation of cell cycle process |
| 1.63E-11 | 142 | 547 | 32 | 0.05850091 | 0.22535211 | GO:0140013 | GO:BP | meiotic nuclear division |
| 1.65E-11 | 29 | 547 | 16 | 0.02925046 | 0.55172414 | GO:0006270 | GO:BP | DNA replication initiation |
| 7.86E-11 | 189 | 547 | 36 | 0.06581353 | 0.19047619 | GO:0033044 | GO:BP | regulation of chromosome organization |
| 1.10E-10 | 971 | 547 | 93 | 0.17001828 | 0.09577755 | GO:0033043 | GO:BP | regulation of organelle organization |
| 8.06E-10 | 690 | 547 | 73 | 0.13345521 | 0.1057971 | GO:0007017 | GO:BP | microtubule-based process |
| 8.24E-10 | 133 | 547 | 29 | 0.05301645 | 0.21804511 | GO:1902850 | GO:BP | microtubule cytoskeleton organization involved in mitosis |

| | | | | | | | | |
|---|---|---|---|---|---|---|---|---|
| 1.25E-09 | 286 | 547 | 43 | 0.0786106 | 0.15034965 | GO:0045787 | GO:BP | positive regulation of cell cycle |
| 1.25E-09 | 91 | 547 | 24 | 0.04387569 | 0.26373626 | GO:0051304 | GO:BP | chromosome separation |
| 1.29E-09 | 108 | 547 | 26 | 0.04753199 | 0.24074074 | GO:0007052 | GO:BP | mitotic spindle organization |
| 3.77E-09 | 87 | 547 | 23 | 0.04204753 | 0.26436782 | GO:0051983 | GO:BP | regulation of chromosome segregation |
| 4.92E-09 | 105 | 547 | 25 | 0.04570384 | 0.23809524 | GO:0051225 | GO:BP | spindle assembly |
| 2.43E-08 | 36 | 547 | 15 | 0.0274223 | 0.41666667 | GO:0008608 | GO:BP | attachment of spindle microtubules to kinetochore |
| 2.47E-08 | 86 | 547 | 22 | 0.04021938 | 0.25581395 | GO:0032508 | GO:BP | DNA duplex unwinding |
| 3.97E-08 | 368 | 547 | 47 | 0.08592322 | 0.12771739 | GO:0044772 | GO:BP | mitotic cell cycle phase transition |
| 5.60E-08 | 98 | 547 | 23 | 0.04204753 | 0.23469388 | GO:0071103 | GO:BP | DNA conformation change |
| 6.63E-08 | 90 | 547 | 22 | 0.04021938 | 0.24444444 | GO:0032392 | GO:BP | DNA geometric change |
| 1.55E-07 | 1134 | 547 | 95 | 0.17367459 | 0.08377425 | GO:0007010 | GO:BP | cytoskeleton organization |
| 3.07E-07 | 71 | 547 | 19 | 0.03473492 | 0.26760563 | GO:1905818 | GO:BP | regulation of chromosome separation |
| 3.71E-07 | 448 | 547 | 51 | 0.09323583 | 0.11383929 | GO:0044770 | GO:BP | cell cycle phase transition |
| 3.75E-07 | 325 | 547 | 42 | 0.07678245 | 0.12923077 | GO:0140694 | GO:BP | non-membrane-bounded organelle assembly |
| 6.16E-07 | 100 | 547 | 22 | 0.04021938 | 0.22 | GO:0061982 | GO:BP | meiosis I cell cycle process |
| 6.79E-07 | 4691 | 547 | 268 | 0.48994516 | 0.05713068 | GO:0016043 | GO:BP | cellular component organization |
| 6.98E-07 | 140 | 547 | 26 | 0.04753199 | 0.18571429 | GO:0044839 | GO:BP | cell cycle G2/M phase transition |
| 1.42E-06 | 77 | 547 | 19 | 0.03473492 | 0.24675325 | GO:0045132 | GO:BP | meiotic chromosome segregation |
| 1.58E-06 | 61 | 547 | 17 | 0.03107861 | 0.27868852 | GO:0090307 | GO:BP | mitotic spindle assembly |
| 1.83E-06 | 96 | 547 | 21 | 0.03839122 | 0.21875 | GO:0007127 | GO:BP | meiosis I |
| 1.88E-06 | 4868 | 547 | 274 | 0.50091408 | 0.05628595 | GO:0071840 | GO:BP | cellular component organization or biogenesis |
| 4.42E-06 | 141 | 547 | 25 | 0.04570384 | 0.17730496 | GO:0051302 | GO:BP | regulation of cell division |
| 5.12E-06 | 121 | 547 | 23 | 0.04204753 | 0.19008264 | GO:0007098 | GO:BP | centrosome cycle |
| 5.69E-06 | 132 | 547 | 24 | 0.04387569 | 0.18181818 | GO:0031023 | GO:BP | microtubule organizing center organization |
| 7.30E-06 | 4271 | 547 | 245 | 0.44789762 | 0.05736362 | GO:0046483 | GO:BP | heterocycle metabolic process |
| 7.56E-06 | 4451 | 547 | 253 | 0.46252285 | 0.05684116 | GO:1901360 | GO:BP | organic cyclic compound metabolic process |
| 7.66E-06 | 4162 | 547 | 240 | 0.43875686 | 0.05766458 | GO:0006139 | GO:BP | nucleobase-containing compound metabolic process |
| 1.17E-05 | 126 | 547 | 23 | 0.04204753 | 0.18253968 | GO:0000086 | GO:BP | G2/M transition of mitotic cell cycle |
| 1.18E-05 | 4313 | 547 | 246 | 0.44972578 | 0.05703687 | GO:0006725 | GO:BP | cellular aromatic compound metabolic process |
| 1.27E-05 | 3809 | 547 | 223 | 0.40767824 | 0.05854555 | GO:0090304 | GO:BP | nucleic acid metabolic process |
| 1.52E-05 | 26 | 547 | 11 | 0.02010969 | 0.42307692 | GO:0007064 | GO:BP | mitotic sister chromatid cohesion |
| 1.60E-05 | 395 | 547 | 44 | 0.08043876 | 0.11139241 | GO:0007346 | GO:BP | regulation of mitotic cell cycle |
| 1.97E-05 | 21 | 547 | 10 | 0.01828154 | 0.47619048 | GO:0006268 | GO:BP | DNA unwinding involved in DNA replication |
| 3.38E-05 | 34 | 547 | 12 | 0.02193784 | 0.35294118 | GO:0044786 | GO:BP | cell cycle DNA replication |
| 4.91E-05 | 42 | 547 | 13 | 0.023766 | 0.30952381 | GO:0033048 | GO:BP | negative regulation of mitotic sister chromatid segregation |
| 4.91E-05 | 42 | 547 | 13 | 0.023766 | 0.30952381 | GO:0033046 | GO:BP | negative regulation of sister chromatid segregation |

**Table S10** Labyrinth zone module associations with outcomes of interest.

| Module color | Module number | sig.fet.count | sig.o2.count | sig.ixn.count | sig.pop | sig.select | Total genes in module | fet_assoc_raw | fet_assoc_BH | popByO2_raw | popByO2_BH | O2_raw | O2_BH | Pop_raw | Pop_BH |
|---|---|---|---|---|---|---|---|---|---|---|---|---|---|---|---|
| green | M16 | 145 | 12 | 0 | 118 | 18 | 434 | 0.0270299 | 0.13514952 | 0.52960397 | 0.96961455 | 0.00492789 | 0.02217551 | 0.5472533 | 0.7577353 |
| magenta | M12 | 57 | 0 | 0 | 44 | 37 | 271 | 0.00525011 | 0.0525011 | 0.89319209 | 0.96961455 | 0.35334851 | 0.45430523 | 0.2034246 | 0.3051368 |
| tan | M9 | 45 | 1 | 0 | 10 | 7 | 124 | 0.00070785 | 0.014157 | 0.63672295 | 0.96961455 | 0.65987866 | 0.69869505 | 0.7484502 | 0.7895793 |
| royalblue | M1 | 0 | 0 | 1 | 9 | 1 | 30 | 0.68671239 | 0.81175218 | 0.00416986 | 0.04169863 | NA | NA | NA | NA |
| pink | M13 | 7 | 80 | 54 | 88 | 5 | 350 | 0.53524528 | 0.72341921 | 0.00126111 | 0.0252221 | NA | NA | NA | NA |
| brown | M18 | 51 | 3 | 3 | 633 | 50 | 830 | 0.86802185 | 0.88603834 | 0.98029706 | 0.98029706 | 0.00159302 | 0.01433718 | 0.00105737 | 0.00237909 |
| blue | M19 | 116 | 24 | 10 | 1765 | 92 | 1899 | 0.35041029 | 0.64738704 | 0.52720211 | 0.96961455 | 0.00139839 | 0.01433718 | 3.41E-17 | 6.138E-16 |
| turquoise | M20 | 830 | 119 | 57 | 3581 | 314 | 7318 | 0.88603834 | 0.88603834 | 0.92113383 | 0.96961455 | 0.00321001 | 0.01926004 | 6.48E-13 | 3.888E-12 |
| Green yellow | M10 | 1 | 0 | 0 | 75 | 8 | 148 | 0.22936658 | 0.50970352 | 0.8606001 | 0.96961455 | 0.81193855 | 0.81193855 | 0.00198186 | 0.00396371 |
| black | M14 | 12 | 0 | 0 | 188 | 20 | 373 | 0.20749482 | 0.50970352 | 0.89118076 | 0.96961455 | 0.13590521 | 0.22239034 | 0.00069426 | 0.00205709 |
| red | M15 | 10 | 0 | 0 | 114 | 14 | 418 | 0.36739108 | 0.64738704 | 0.82472655 | 0.96961455 | 0.4140176 | 0.49682112 | 0.00571366 | 0.01028459 |
| yellow | M17 | 13 | 8 | 0 | 534 | 27 | 658 | 0.16819196 | 0.50970352 | 0.4223419 | 0.96961455 | 0.09425099 | 0.18850198 | 7.74E-14 | 6.966E-13 |
| grey60 | M4 | 1 | 0 | 0 | 13 | 0 | 33 | 0.54256441 | 0.72341921 | 0.1384882 | 0.69244099 | 0.06953716 | 0.16515621 | 0.00079998 | 0.00205709 |
| lightcyan | M5 | 1 | 0 | 0 | 46 | 2 | 49 | 0.84527493 | 0.88603834 | 0.20772642 | 0.83090568 | 0.11033629 | 0.19860532 | 0.000048 | 0.0001728 |
| cyan | M7 | 9 | 1 | 0 | 16 | 2 | 82 | 0.0514653 | 0.20586121 | 0.70358729 | 0.96961455 | 0.21657773 | 0.3248666 | 6.58E-07 | 2.961E-06 |
| purple | M11 | 36 | 1 | 0 | 59 | 18 | 254 | 0.38843222 | 0.64738704 | 0.32984418 | 0.96961455 | 0.59695598 | 0.67157547 | 0.1338884 | 0.2190902 |
| lightyellow | M2 | 4 | 0 | 0 | 5 | **3** | 31 | 0.01380221 | 0.09201471 | 0.56442579 | 0.96961455 | 0.23475487 | 0.3250452 | 0.7535029 | 0.7895793 |
| lightgreen | M3 | 0 | 0 | 0 | 0 | 4 | 32 | 0.54229614 | 0.72341921 | 0.64314936 | 0.96961455 | 0.04701401 | 0.14104204 | 0.7747577 | 0.7895793 |
| midnightblue | M6 | 2 | 0 | 0 | 0 | 3 | 50 | 0.18142533 | 0.50970352 | 0.89106469 | 0.96961455 | 0.02122468 | 0.07640884 | 0.7895793 | 0.7895793 |
| salmon | M8 | 0 | 0 | 0 | 18 | 1 | 85 | 0.68998936 | 0.81175218 | 0.12961567 | 0.69244099 | 0.07340276 | 0.16515621 | 0.6216042 | 0.7895793 |
| **DE_gene_set** | - | **1354** | **254** | **125** | 7436 | 654 | **14345** | | | | | | | | |

**Table S11** Junctional zone module associations with outcomes of interest.

| Module_color | Module_number | sig.fet.count | sig.o2.count | sig.ixn.count | sig.pop | Total genes in module | fet_assoc_raw | fet_assoc_BH | popByO2_raw | popByO2_BH | O2_raw | O2_BH | Pop_raw | Pop_BH |
|---|---|---|---|---|---|---|---|---|---|---|---|---|---|---|
| purple | 10 | | 12 | 63 | 51 | 319 | 0.117015082 | 0.7031672 | 0.00103713 | 0.01970539 | 0.09241614 | 0.4389766 | 2.18E-06 | 5.92E-06 |
| red | 14 | 0 | 12 | 28 | 111 | 375 | 0.478023018 | 0.7031672 | 0.0488421 | 0.31565741 | 0.04877364 | 0.4389766 | 0.51814835 | 5.47E-01 |
| lightgreen | 2 | 0 | 0 | 0 | 25 | 36 | 0.60083858 | 0.7227875 | 0.17684485 | 0.34975471 | 0.23155243 | 0.4620514 | 0.00277895 | 4.80E-03 |
| grey60 | 3 | 0 | 0 | 0 | 30 | 38 | 0.481114371 | 0.7031672 | 0.04984064 | 0.31565741 | 0.23470344 | 0.4620514 | 0.00071283 | 1.35E-03 |
| midnightblue | 5 | 0 | 0 | 0 | 23 | 73 | 0.164196508 | 0.7031672 | 0.99702909 | 0.99702909 | 0.67897113 | 0.8062782 | 0.00370625 | 5.87E-03 |
| cyan | 6 | 0 | 0 | 1 | 40 | 77 | 0.271228258 | 0.7031672 | 0.1992765 | 0.34975471 | 0.01038135 | 0.1972456 | 2.46E-09 | 1.17E-08 |
| tan | 8 | 0 | 0 | 1 | 172 | 182 | 0.704725603 | 0.7227875 | 0.20058817 | 0.34975471 | 0.26750342 | 0.4620514 | 1.58E-13 | 1.50E-12 |
| magenta | 11 | 0 | 0 | 0 | 197 | 335 | 0.18206349 | 0.7031672 | 0.72093804 | 0.85611393 | 0.9868545 | 0.9868545 | 9.93E-09 | 3.77E-08 |
| pink | 12 | 0 | 5 | 2 | 135 | 365 | 0.187173129 | 0.7031672 | 0.82258226 | 0.91935664 | 0.0793018 | 0.4389766 | 1.84E-05 | 4.37E-05 |
| black | 13 | 0 | 0 | 8 | 229 | 371 | 0.352499373 | 0.7031672 | 0.10921814 | 0.34379371 | 0.60237619 | 0.7630098 | 3.08E-08 | 9.75E-08 |
| green | 15 | 0 | 0 | 2 | 272 | 733 | 0.664091086 | 0.7227875 | 0.51056545 | 0.74621103 | 0.59091166 | 0.7630098 | 0.0001012 | 2.14E-04 |
| blue | 18 | 1 | 7 | 21 | 1717 | 1961 | 0.361754734 | 0.7031672 | 0.91279204 | 0.96350271 | 0.25266134 | 0.4620514 | 3.18E-20 | 6.04E-19 |
| turquoise | 19 | 0 | 35 | 72 | 2191 | 2419 | 0.374043083 | 0.7031672 | 0.20248957 | 0.34975471 | 0.16610455 | 0.4620514 | 4.09E-12 | 2.59E-11 |
| lightyellow | 1 | 0 | 0 | 0 | 5 | 23 | 0.059114443 | 0.7031672 | 0.55107866 | 0.74789247 | 0.12587568 | 0.4620514 | 0.09049003 | 1.23E-01 |
| lightcyan | 4 | 0 | 0 | 0 | 9 | 46 | 0.630910158 | 0.7227875 | 0.32435609 | 0.51356381 | 0.92927327 | 0.9832319 | 0.630943 | 6.31E-01 |
| salmon | 7 | 0 | 1 | 7 | 29 | 132 | 0.582313025 | 0.7227875 | 0.08512568 | 0.34379371 | 0.24692965 | 0.4620514 | 0.19524524 | 2.32E-01 |
| greenyellow | 9 | 0 | 0 | 2 | 48 | 280 | 0.722787463 | 0.7227875 | 0.12666084 | 0.34379371 | 0.93148285 | 0.9832319 | 0.05869246 | 8.58E-02 |
| yellow | 16 | 0 | 2 | 17 | 422 | 1078 | 0.471240627 | 0.7031672 | 0.61203448 | 0.77524368 | 0.36137732 | 0.5721808 | 0.11090744 | 1.40E-01 |
| brown | 17 | 0 | 0 | 31 | 493 | 1654 | 0.384125915 | 0.7031672 | 0.09833217 | 0.34379371 | 0.45282248 | 0.6618175 | 0.4304333 | 4.81E-01 |
| **DE_gene_set** | - | **3** | 101 | 359 | 6918 | **14000** | | | | | | | | |

**Table S12** GO enrichment results for gene modules of interest in the LZ

| Outcome Assoc. | Mod. | p_value | Term size | Query size | Intersection size | precision | recall | term_id | source | term_name |
|---|---|---|---|---|---|---|---|---|---|---|
| Fetal mass | M16 | 2.55E-14 | 735 | 415 | 72 | 0.17349398 | 0.09795918 | GO:0035239 | GO:BP | tube morphogenesis |
| Fetal mass | M16 | 8.61E-14 | 594 | 415 | 63 | 0.15180723 | 0.10606061 | GO:0001568 | GO:BP | blood vessel development |
| Fetal mass | M16 | 8.37E-13 | 622 | 415 | 63 | 0.15180723 | 0.10128617 | GO:0001944 | GO:BP | vasculature development |
| Fetal mass | M16 | 2.85E-12 | 518 | 415 | 56 | 0.13493976 | 0.10810811 | GO:0048514 | GO:BP | blood vessel morphogenesis |
| Fetal mass | M16 | 3.76E-12 | 921 | 415 | 78 | 0.18795181 | 0.08469055 | GO:0072359 | GO:BP | circulatory system development |
| Fetal mass | M16 | 7.48E-12 | 2027 | 415 | 128 | 0.30843373 | 0.06314751 | GO:0009653 | GO:BP | anatomical structure morphogenesis |
| Fetal mass | M16 | 1.42E-11 | 904 | 415 | 76 | 0.18313253 | 0.0840708 | GO:0035295 | GO:BP | tube development |
| Fetal mass | M16 | 4.44E-10 | 3903 | 415 | 195 | 0.46987952 | 0.04996157 | GO:0023052 | GO:BP | signaling |
| Fetal mass | M16 | 6.99E-10 | 1836 | 415 | 115 | 0.27710843 | 0.06263617 | GO:0007166 | GO:BP | cell surface receptor signaling pathway |
| Fetal mass | M16 | 8.98E-10 | 3594 | 415 | 183 | 0.44096386 | 0.0509182 | GO:0007165 | GO:BP | signal transduction |
| Fetal mass | M16 | 4.00E-09 | 3130 | 415 | 164 | 0.39518072 | 0.05239617 | GO:0048731 | GO:BP | system development |
| Fetal mass | M16 | 9.59E-09 | 3963 | 415 | 193 | 0.46506024 | 0.04870048 | GO:0007154 | GO:BP | cell communication |
| Fetal mass | M16 | 7.72E-08 | 1423 | 415 | 92 | 0.22168675 | 0.06465214 | GO:0009888 | GO:BP | tissue development |
| Fetal mass | M16 | 3.99E-07 | 912 | 415 | 67 | 0.16144578 | 0.07346491 | GO:0048646 | GO:BP | anatomical structure formation involved in morphogenesis |
| Fetal mass | M16 | 5.05E-07 | 1977 | 415 | 113 | 0.27228916 | 0.05715731 | GO:0051239 | GO:BP | regulation of multicellular organismal process |
| Fetal mass | M16 | 5.82E-07 | 3482 | 415 | 170 | 0.40963855 | 0.04882252 | GO:0007275 | GO:BP | multicellular organism development |
| Fetal mass | M16 | 6.64E-07 | 1073 | 415 | 74 | 0.17831325 | 0.06896552 | GO:2000026 | GO:BP | regulation of multicellular organismal development |
| Fetal mass | M16 | 8.94E-07 | 439 | 415 | 42 | 0.10120482 | 0.09567198 | GO:0001525 | GO:BP | angiogenesis |
| Fetal mass | M16 | 1.02E-06 | 2931 | 415 | 149 | 0.35903614 | 0.05083589 | GO:0048869 | GO:BP | cellular developmental process |
| Fetal mass | M16 | 1.03E-06 | 869 | 415 | 64 | 0.15421687 | 0.07364787 | GO:0060429 | GO:BP | epithelium development |
| Fetal mass | M16 | 1.04E-06 | 1852 | 415 | 107 | 0.25783133 | 0.05777538 | GO:0050793 | GO:BP | regulation of developmental process |
| Fetal mass | M16 | 1.20E-06 | 77 | 415 | 17 | 0.04096386 | 0.22077922 | GO:0001570 | GO:BP | vasculogenesis |
| Fetal mass | M16 | 1.55E-06 | 4614 | 415 | 208 | 0.50120482 | 0.04508019 | GO:0032501 | GO:BP | multicellular organismal process |

| | | | | | | | | | | |
|---|---|---|---|---|---|---|---|---|---|---|
| Fetal mass | M16 | 2.30E-06 | 2909 | 415 | 147 | 0.35421687 | 0.05053283 | GO:0030154 | GO:BP | cell differentiation |
| Fetal mass | M16 | 2.34E-06 | 681 | 415 | 54 | 0.13012048 | 0.07929515 | GO:0051093 | GO:BP | negative regulation of developmental process |
| Fetal mass | M16 | 8.22E-06 | 1340 | 415 | 83 | 0.2 | 0.0619403 | GO:0040011 | GO:BP | locomotion |
| Fetal mass | M16 | 1.23E-05 | 2444 | 415 | 127 | 0.3060241 | 0.05196399 | GO:0023051 | GO:BP | regulation of signaling |
| Fetal mass | M16 | 1.25E-05 | 1236 | 415 | 78 | 0.18795181 | 0.0631068 | GO:0048870 | GO:BP | cell motility |
| Fetal mass | M16 | 1.25E-05 | 1236 | 415 | 78 | 0.18795181 | 0.0631068 | GO:0051674 | GO:BP | localization of cell |
| Fetal mass | M16 | 1.38E-05 | 4778 | 415 | 210 | 0.5060241 | 0.04395144 | GO:0051716 | GO:BP | cellular response to stimulus |
| Fetal mass | M16 | 1.50E-05 | 2167 | 415 | 116 | 0.27951807 | 0.05353023 | GO:0009966 | GO:BP | regulation of signal transduction |
| Fetal mass | M16 | 1.88E-05 | 2434 | 415 | 126 | 0.30361446 | 0.05176664 | GO:0010646 | GO:BP | regulation of cell communication |
| Fetal mass | M16 | 2.11E-05 | 958 | 415 | 65 | 0.15662651 | 0.06784969 | GO:0009790 | GO:BP | embryo development |
| Fetal mass | M16 | 2.39E-05 | 4042 | 415 | 184 | 0.44337349 | 0.04552202 | GO:0048856 | GO:BP | anatomical structure development |
| Fetal mass | M16 | 2.44E-05 | 1140 | 415 | 73 | 0.17590361 | 0.06403509 | GO:0016477 | GO:BP | cell migration |
| Fetal mass | M16 | 2.84E-05 | 4370 | 415 | 195 | 0.46987952 | 0.04462243 | GO:0032502 | GO:BP | developmental process |
| Fetal mass | M16 | 4.22E-05 | 5626 | 415 | 236 | 0.5686747 | 0.0419481 | GO:0050896 | GO:BP | response to stimulus |
| Fetal mass | M16 | 5.00E-05 | 2525 | 415 | 128 | 0.30843373 | 0.05069307 | GO:0048513 | GO:BP | animal organ development |
| Fetal mass | M16 | 7.51E-05 | 751 | 415 | 54 | 0.13012048 | 0.07190413 | GO:0009887 | GO:BP | animal organ morphogenesis |
| Fetal mass | M16 | 7.93E-05 | 1172 | 415 | 73 | 0.17590361 | 0.06228669 | GO:0045595 | GO:BP | regulation of cell differentiation |
| Fetal mass | M16 | 8.54E-05 | 1015 | 415 | 66 | 0.15903614 | 0.06502463 | GO:0051094 | GO:BP | positive regulation of developmental process |
| Fetal mass | M16 | 1.04E-04 | 114 | 415 | 18 | 0.04337349 | 0.15789474 | GO:0003158 | GO:BP | endothelium development |
| Fetal mass | M16 | 1.87E-04 | 1530 | 415 | 87 | 0.20963855 | 0.05686275 | GO:0006928 | GO:BP | movement of cell or subcellular component |
| Fetal mass | M16 | 2.23E-04 | 489 | 415 | 40 | 0.09638554 | 0.08179959 | GO:0045596 | GO:BP | negative regulation of cell differentiation |
| Fetal mass | M16 | 2.42E-04 | 452 | 415 | 38 | 0.09156627 | 0.0840708 | GO:0030855 | GO:BP | epithelial cell differentiation |
| Fetal mass | M16 | 2.72E-04 | 108 | 415 | 17 | 0.04096386 | 0.15740741 | GO:0030856 | GO:BP | regulation of epithelial cell differentiation |
| Fetal mass | M16 | 2.93E-04 | 362 | 415 | 33 | 0.07951807 | 0.09116022 | GO:0048568 | GO:BP | embryonic organ development |
| Fetal mass | M16 | 3.34E-04 | 961 | 415 | 62 | 0.14939759 | 0.06451613 | GO:0007155 | GO:BP | cell adhesion |
| Fetal mass | M16 | 3.38E-04 | 401 | 415 | 35 | 0.08433735 | 0.0872818 | GO:1905114 | GO:BP | cell surface receptor signaling pathway involved in cell-cell signaling |
| Fetal mass | M16 | 4.14E-04 | 747 | 415 | 52 | 0.1253012 | 0.06961178 | GO:0030334 | GO:BP | regulation of cell migration |
| Fetal mass | M16 | 4.18E-04 | 812 | 415 | 55 | 0.13253012 | 0.06773399 | GO:0040012 | GO:BP | regulation of locomotion |

| | | | | | | | | | | |
|---|---|---|---|---|---|---|---|---|---|---|
| Fetal mass | M16 | 4.21E-04 | 167 | 415 | 21 | 0.05060241 | 0.1257485 | GO:0046578 | GO:BP | regulation of Ras protein signal transduction |
| Fetal mass | M16 | 6.47E-04 | 202 | 415 | 23 | 0.05542169 | 0.11386139 | GO:0051056 | GO:BP | regulation of small GTPase mediated signal transduction |
| Fetal mass | M16 | 6.94E-04 | 695 | 415 | 49 | 0.11807229 | 0.0705036 | GO:0048534 | GO:BP | hematopoietic or lymphoid organ development |
| Fetal mass | M16 | 8.24E-04 | 785 | 415 | 53 | 0.12771084 | 0.06751592 | GO:2000145 | GO:BP | regulation of cell motility |
| Fetal mass | M16 | 8.56E-04 | 37 | 415 | 10 | 0.02409639 | 0.27027027 | GO:0030857 | GO:BP | negative regulation of epithelial cell differentiation |
| Fetal mass | M16 | 1.58E-03 | 526 | 415 | 40 | 0.09638554 | 0.07604563 | GO:0071363 | GO:BP | cellular response to growth factor stimulus |
| Fetal mass | M16 | 1.61E-03 | 846 | 415 | 55 | 0.13253012 | 0.06501182 | GO:0051270 | GO:BP | regulation of cellular component movement |
| Fetal mass | M16 | 1.66E-03 | 1961 | 415 | 101 | 0.24337349 | 0.05150433 | GO:0035556 | GO:BP | intracellular signal transduction |
| Fetal mass | M16 | 2.34E-03 | 702 | 415 | 48 | 0.11566265 | 0.06837607 | GO:0007167 | GO:BP | enzyme-linked receptor protein signaling pathway |
| Fetal mass | M16 | 2.37E-03 | 1619 | 415 | 87 | 0.20963855 | 0.05373687 | GO:0048468 | GO:BP | cell development |
| Fetal mass | M16 | 2.45E-03 | 98 | 415 | 15 | 0.03614458 | 0.15306122 | GO:0045446 | GO:BP | endothelial cell differentiation |
| Fetal mass | M16 | 2.49E-03 | 1017 | 415 | 62 | 0.14939759 | 0.06096362 | GO:0007267 | GO:BP | cell-cell signaling |
| Fetal mass | M16 | 2.49E-03 | 515 | 415 | 39 | 0.0939759 | 0.07572816 | GO:0048729 | GO:BP | tissue morphogenesis |
| Fetal mass | M16 | 2.60E-03 | 341 | 415 | 30 | 0.07228916 | 0.08797654 | GO:0016055 | GO:BP | Wnt signaling pathway |
| Fetal mass | M16 | 2.76E-03 | 342 | 415 | 30 | 0.07228916 | 0.0877193 | GO:0198738 | GO:BP | cell-cell signaling by wnt |
| Fetal mass | M16 | 3.25E-03 | 1095 | 415 | 65 | 0.15662651 | 0.05936073 | GO:0051240 | GO:BP | positive regulation of multicellular organismal process |
| Fetal mass | M16 | 3.47E-03 | 2793 | 415 | 131 | 0.31566265 | 0.04690297 | GO:0048583 | GO:BP | regulation of response to stimulus |
| Fetal mass | M16 | 3.56E-03 | 1145 | 415 | 67 | 0.16144578 | 0.05851528 | GO:0009967 | GO:BP | positive regulation of signal transduction |
| Fetal mass | M16 | 3.95E-03 | 545 | 415 | 40 | 0.09638554 | 0.0733945 | GO:0070848 | GO:BP | response to growth factor |
| Fetal mass | M16 | 4.89E-03 | 742 | 415 | 49 | 0.11807229 | 0.06603774 | GO:0002520 | GO:BP | immune system development |
| Pop-by-hypoxia | M1 | 0.01101926 | 3 | 24 | 1 | 0.04166667 | 0.33333333 | CORUM:6221 | CORUM | Ccm2-Krit1-Ccm3 complex |
| Pop-by-hypoxia | M1 | 0.01857411 | 2493 | 24 | 13 | 0.54166667 | 0.0052146 | GO:0031090 | GO:CC | organelle membrane |
| Pop-by-hypoxia | M13 | 9.66E-05 | 2563 | 320 | 104 | 0.325 | 0.04057745 | GO:0051252 | GO:BP | regulation of RNA metabolic process |
| Pop-by-hypoxia | M13 | 1.40E-04 | 2821 | 320 | 111 | 0.346875 | 0.03934775 | GO:0019219 | GO:BP | regulation of nucleobase-containing compound metabolic process |
| Pop-by-hypoxia | M13 | 3.67E-04 | 1736 | 320 | 77 | 0.240625 | 0.04435484 | GO:0006366 | GO:BP | transcription by RNA polymerase II |
| Pop-by-hypoxia | M13 | 4.48E-04 | 1681 | 320 | 75 | 0.234375 | 0.0446163 | GO:0006357 | GO:BP | regulation of transcription by RNA polymerase II |
| Pop-by-hypoxia | M13 | 1.43E-03 | 4034 | 320 | 141 | 0.440625 | 0.0349529 | GO:0051171 | GO:BP | regulation of nitrogen compound metabolic process |

| | | | | | | | | | | |
|---|---|---|---|---|---|---|---|---|---|---|
| Pop-by-hypoxia | M13 | 2.05E-03 | 2718 | 320 | 104 | 0.325 | 0.03826343 | GO:0010556 | GO:BP | regulation of macromolecule biosynthetic process |
| Pop-by-hypoxia | M13 | 2.25E-03 | 2413 | 320 | 95 | 0.296875 | 0.03937008 | GO:0006351 | GO:BP | transcription |
| Pop-by-hypoxia | M13 | 2.29E-03 | 2414 | 320 | 95 | 0.296875 | 0.03935377 | GO:0097659 | GO:BP | nucleic acid-templated transcription |
| Pop-by-hypoxia | M13 | 2.51E-03 | 3369 | 320 | 122 | 0.38125 | 0.03621253 | GO:0010468 | GO:BP | regulation of gene expression |
| Pop-by-hypoxia | M13 | 2.89E-03 | 2324 | 320 | 92 | 0.2875 | 0.03958692 | GO:0006355 | GO:BP | regulation of transcription |
| Pop-by-hypoxia | M13 | 2.95E-03 | 2325 | 320 | 92 | 0.2875 | 0.03956989 | GO:1903506 | GO:BP | regulation of nucleic acid-templated transcription |
| Pop-by-hypoxia | M13 | 2.99E-03 | 2844 | 320 | 107 | 0.334375 | 0.03762307 | GO:0031326 | GO:BP | regulation of cellular biosynthetic process |
| Pop-by-hypoxia | M13 | 3.11E-03 | 2430 | 320 | 95 | 0.296875 | 0.03909465 | GO:0032774 | GO:BP | RNA biosynthetic process |
| Pop-by-hypoxia | M13 | 3.12E-03 | 4159 | 320 | 143 | 0.446875 | 0.03438327 | GO:0080090 | GO:BP | regulation of primary metabolic process |
| Pop-by-hypoxia | M13 | 3.38E-03 | 2332 | 320 | 92 | 0.2875 | 0.03945111 | GO:2001141 | GO:BP | regulation of RNA biosynthetic process |
| Pop-by-hypoxia | M13 | 3.95E-03 | 3324 | 320 | 120 | 0.375 | 0.03610108 | GO:0016070 | GO:BP | RNA metabolic process |
| Pop-by-hypoxia | M13 | 4.78E-03 | 2906 | 320 | 108 | 0.3375 | 0.03716449 | GO:0009889 | GO:BP | regulation of biosynthetic process |
| Pop-by-hypoxia | M13 | 4.81E-03 | 3336 | 320 | 120 | 0.375 | 0.03597122 | GO:0044271 | GO:BP | cellular nitrogen compound biosynthetic process |
| Pop-by-hypoxia | M13 | 1.15E-02 | 3354 | 320 | 119 | 0.371875 | 0.03548002 | GO:0009059 | GO:BP | macromolecule biosynthetic process |
| Pop-by-hypoxia | M13 | 1.19E-02 | 2889 | 320 | 106 | 0.33125 | 0.0366909 | GO:1901362 | GO:BP | organic cyclic compound biosynthetic process |
| Pop-by-hypoxia | M13 | 1.29E-02 | 4369 | 320 | 146 | 0.45625 | 0.03341726 | GO:0010467 | GO:BP | gene expression |
| Pop-by-hypoxia | M13 | 1.39E-02 | 2792 | 320 | 103 | 0.321875 | 0.03689112 | GO:0018130 | GO:BP | heterocycle biosynthetic process |
| Pop-by-hypoxia | M13 | 1.69E-02 | 2733 | 320 | 101 | 0.315625 | 0.03695573 | GO:0034654 | GO:BP | nucleobase-containing compound biosynthetic process |
| Pop-by-hypoxia | M13 | 1.88E-02 | 4396 | 320 | 146 | 0.45625 | 0.03321201 | GO:0060255 | GO:BP | regulation of macromolecule metabolic process |
| Pop-by-hypoxia | M13 | 2.53E-02 | 2793 | 320 | 102 | 0.31875 | 0.03651987 | GO:0019438 | GO:BP | aromatic compound biosynthetic process |
| Pop-by-hypoxia | M13 | 3.59E-02 | 4290 | 320 | 142 | 0.44375 | 0.03310023 | GO:0031323 | GO:BP | regulation of cellular metabolic process |
| Pop-by-hypoxia | M13 | 5.33E-03 | 5667 | 320 | 177 | 0.553125 | 0.03123346 | GO:0005634 | GO:CC | nucleus |
| Pop-by-hypoxia | M13 | 2.40E-04 | 668 | 320 | 39 | 0.121875 | 0.05838323 | GO:0000987 | GO:MF | cis-regulatory region sequence-specific DNA binding |
| Pop-by-hypoxia | M13 | 2.89E-04 | 978 | 320 | 50 | 0.15625 | 0.05112474 | GO:0043565 | GO:MF | sequence-specific DNA binding |

| | | | | | | | | | | |
|---|---|---|---|---|---|---|---|---|---|---|
| Pop-by-hypoxia | M13 | 3.31E-04 | 650 | 320 | 38 | 0.11875 | 0.05846154 | GO:0000978 | GO:MF | RNA polymerase II cis-regulatory region sequence-specific DNA binding |
| Pop-by-hypoxia | M13 | 1.03E-03 | 1140 | 320 | 54 | 0.16875 | 0.04736842 | GO:0140110 | GO:MF | transcription regulator activity |
| Pop-by-hypoxia | M13 | 1.05E-03 | 1575 | 320 | 68 | 0.2125 | 0.0431746 | GO:0003677 | GO:MF | DNA binding |
| Pop-by-hypoxia | M13 | 3.58E-03 | 746 | 320 | 39 | 0.121875 | 0.05227882 | GO:0003700 | GO:MF | DNA-binding transcription factor activity |
| Pop-by-hypoxia | M13 | 6.84E-03 | 767 | 320 | 39 | 0.121875 | 0.05084746 | GO:0000977 | GO:MF | RNA polymerase II transcription regulatory region sequence-specific DNA binding |
| Pop-by-hypoxia | M13 | 8.31E-03 | 890 | 320 | 43 | 0.134375 | 0.04831461 | GO:1990837 | GO:MF | sequence-specific double-stranded DNA binding |
| Pop-by-hypoxia | M13 | 1.16E-02 | 699 | 320 | 36 | 0.1125 | 0.05150215 | GO:0000981 | GO:MF | DNA-binding transcription factor activity |
| Pop-by-hypoxia | M13 | 1.18E-02 | 844 | 320 | 41 | 0.128125 | 0.0485782 | GO:0000976 | GO:MF | transcription cis-regulatory region binding |
| Pop-by-hypoxia | M13 | 1.32E-02 | 848 | 320 | 41 | 0.128125 | 0.04834906 | GO:0001067 | GO:MF | transcription regulatory region nucleic acid binding |
| Pop-by-hypoxia | M13 | 1.34E-02 | 40 | 320 | 7 | 0.021875 | 0.175 | MIRNA:mmu-miR-125b-5p | MIRNA | mmu-miR-125b-5p |
| Pop-by-hypoxia | M13 | 2.71E-04 | 14 | 320 | 6 | 0.01875 | 0.42857143 | REAC:R-MMU-9670095 | REAC | Inhibition of DNA recombination at telomere |
| Pop-by-hypoxia | M13 | 4.42E-04 | 15 | 320 | 6 | 0.01875 | 0.4 | REAC:R-MMU-110330 | REAC | Recognition and association of DNA glycosylase with site containing an affected purine |
| Pop-by-hypoxia | M13 | 1.05E-03 | 17 | 320 | 6 | 0.01875 | 0.35294118 | REAC:R-MMU-73927 | REAC | Depurination |
| Pop-by-hypoxia | M13 | 1.05E-03 | 17 | 320 | 6 | 0.01875 | 0.35294118 | REAC:R-MMU-110331 | REAC | Cleavage of the damaged purine |
| Pop-by-hypoxia | M13 | 5.70E-03 | 22 | 320 | 6 | 0.01875 | 0.27272727 | REAC:R-MMU-73929 | REAC | Base-Excision Repair |
| Pop-by-hypoxia | M13 | 1.16E-02 | 36 | 320 | 7 | 0.021875 | 0.19444444 | REAC:R-MMU-73772 | REAC | RNA Polymerase I Promoter Escape |
| Pop-by-hypoxia | M13 | 1.40E-02 | 37 | 320 | 7 | 0.021875 | 0.18918919 | REAC:R-MMU-5250913 | REAC | Positive epigenetic regulation of rRNA expression |
| Pop-by-hypoxia | M13 | 1.40E-02 | 37 | 320 | 7 | 0.021875 | 0.18918919 | REAC:R-MMU-5250924 | REAC | B-WICH complex positively regulates rRNA expression |
| Pop-by-hypoxia | M13 | 2.74E-02 | 55 | 320 | 8 | 0.025 | 0.14545455 | REAC:R-MMU-212165 | REAC | Epigenetic regulation of gene expression |
| Pop-by-hypoxia | M13 | 3.40E-02 | 10 | 320 | 4 | 0.0125 | 0.4 | REAC:R-MMU-427413 | REAC | NoRC negatively regulates rRNA expression |
| Pop-by-hypoxia | M13 | 1.97E-07 | 9684 | 320 | 284 | 0.8875 | 0.02932672 | TF:M00716_1 | TF | Factor: ZF5; motif: GSGCGCGR; match class: 1 |
| Pop-by-hypoxia | M13 | 9.97E-07 | 9199 | 320 | 273 | 0.853125 | 0.02967714 | TF:M01240_1 | TF | Factor: BEN; motif: CAGCGRNV; match class: 1 |
| Pop-by-hypoxia | M13 | 3.03E-06 | 4743 | 320 | 169 | 0.528125 | 0.03563146 | TF:M00245 | TF | Factor: Egr-3; motif: NTGCGTGGGCGK |
| Pop-by-hypoxia | M13 | 3.90E-05 | 5360 | 320 | 181 | 0.565625 | 0.03376866 | TF:M00322 | TF | Factor: c-Myc:Max; motif: GCCAYGYGSN |

| | | | | | | | | | | |
|---|---|---|---|---|---|---|---|---|---|---|
| Pop-by-hypoxia | M13 | 4.11E-05 | 8496 | 320 | 254 | 0.79375 | 0.02989642 | TF:M00803 | TF | Factor: E2F; motif: GGCGSG |
| Pop-by-hypoxia | M13 | 4.23E-05 | 2363 | 320 | 99 | 0.309375 | 0.0418959 | TF:M00428_1 | TF | Factor: E2F-1; motif: NKTSSCGC; match class: 1 |
| Pop-by-hypoxia | M13 | 5.28E-05 | 10043 | 320 | 285 | 0.890625 | 0.02837797 | TF:M04662_1 | TF | Factor: FOXN4; motif: NNWANNCGWMCGCGTCNNNNMT; match class: 1 |
| Pop-by-hypoxia | M13 | 5.36E-05 | 5499 | 320 | 184 | 0.575 | 0.03346063 | TF:M05547 | TF | Factor: ZAC; motif: KGGGCCR |
| Pop-by-hypoxia | M13 | 6.97E-05 | 5838 | 320 | 192 | 0.6 | 0.03288798 | TF:M00803_1 | TF | Factor: E2F; motif: GGCGSG; match class: 1 |
| Pop-by-hypoxia | M13 | 2.18E-04 | 888 | 320 | 49 | 0.153125 | 0.05518018 | TF:M02914_1 | TF | Factor: SP4; motif: NNWAGGCGTGNCNNN; match class: 1 |
| Pop-by-hypoxia | M13 | 2.43E-04 | 11134 | 320 | 303 | 0.946875 | 0.02721394 | TF:M00716 | TF | Factor: ZF5; motif: GSGCGCGR |
| Pop-by-hypoxia | M13 | 2.50E-04 | 8197 | 320 | 245 | 0.765625 | 0.02988898 | TF:M07991_1 | TF | Factor: Hes1; motif: NNCACGYGNN; match class: 1 |
| Pop-by-hypoxia | M13 | 4.17E-04 | 5153 | 320 | 172 | 0.5375 | 0.03337861 | TF:M08870 | TF | Factor: DEC1; motif: NCNCACRTGNSC |
| Pop-by-hypoxia | M13 | 7.06E-04 | 6195 | 320 | 197 | 0.615625 | 0.03179984 | TF:M00428 | TF | Factor: E2F-1; motif: NKTSSCGC |
| Pop-by-hypoxia | M13 | 8.81E-04 | 1073 | 320 | 54 | 0.16875 | 0.05032619 | TF:M10200_1 | TF | Factor: CREB1; motif: NRRTGACGTMA; match class: 1 |
| Pop-by-hypoxia | M13 | 9.10E-04 | 4467 | 320 | 153 | 0.478125 | 0.03425118 | TF:M02012 | TF | Factor: HIF-1alpha; motif: NCACGT |
| Pop-by-hypoxia | M13 | 1.73E-03 | 2716 | 320 | 104 | 0.325 | 0.03829161 | TF:M07260_1 | TF | Factor: Ikaros; motif: TGGGAGN; match class: 1 |
| Pop-by-hypoxia | M13 | 1.73E-03 | 4895 | 320 | 163 | 0.509375 | 0.03329928 | TF:M08005 | TF | Factor: Tcfl5; motif: NNCNCGNGNN |
| Pop-by-hypoxia | M13 | 1.73E-03 | 4895 | 320 | 163 | 0.509375 | 0.03329928 | TF:M08005_1 | TF | Factor: Tcfl5; motif: NNCNCGNGNN; match class: 1 |
| Pop-by-hypoxia | M13 | 2.16E-03 | 4031 | 320 | 140 | 0.4375 | 0.03473084 | TF:M08885 | TF | Factor: HAIRYLIKE; motif: NNNNCANGTG |
| Pop-by-hypoxia | M13 | 2.50E-03 | 7124 | 320 | 217 | 0.678125 | 0.03046042 | TF:M07250 | TF | Factor: E2F-1; motif: NNNSSCGCSAANN |
| Pop-by-hypoxia | M13 | 3.30E-03 | 11516 | 320 | 307 | 0.959375 | 0.02665856 | TF:M04662 | TF | Factor: FOXN4; motif: NNWANNCGWMCGCGTCNNNNMT |
| Fetal mass | M12 | 4.91E-02 | 3 | 263 | 2 | 0.007604563 | 0.66666667 | CORUM:6912 | CORUM | Cbl-Frs2-Grb2 complex |
| Fetal mass | M12 | 1.50E-05 | 2821 | 263 | 98 | 0.372623574 | 0.03473945 | GO:0019219 | GO:BP | regulation of nucleobase-containing compound metabolic process |
| Fetal mass | M12 | 1.76E-05 | 842 | 263 | 44 | 0.16730038 | 0.05225653 | GO:0051276 | GO:BP | chromosome organization |
| Fetal mass | M12 | 5.85E-05 | 4290 | 263 | 130 | 0.494296578 | 0.03030303 | GO:0031323 | GO:BP | regulation of cellular metabolic process |
| Fetal mass | M12 | 7.34E-05 | 4396 | 263 | 132 | 0.501901141 | 0.0300273 | GO:0060255 | GO:BP | regulation of macromolecule metabolic process |
| Fetal mass | M12 | 7.94E-05 | 468 | 263 | 30 | 0.114068441 | 0.06410256 | GO:0006325 | GO:BP | chromatin organization |
| Fetal mass | M12 | 1.30E-04 | 4159 | 263 | 126 | 0.479087452 | 0.03029574 | GO:0080090 | GO:BP | regulation of primary metabolic process |

| | | | | | | | | | | |
|---|---|---|---|---|---|---|---|---|---|---|
| Fetal mass | M12 | 1.52E-04 | 4769 | 263 | 139 | 0.52851711 | 0.02914657 | GO:0019222 | GO:BP | regulation of metabolic process |
| Fetal mass | M12 | 1.53E-04 | 4034 | 263 | 123 | 0.467680608 | 0.03049083 | GO:0051171 | GO:BP | regulation of nitrogen compound metabolic process |
| Fetal mass | M12 | 6.02E-04 | 3809 | 263 | 116 | 0.441064639 | 0.03045419 | GO:0090304 | GO:BP | nucleic acid metabolic process |
| Fetal mass | M12 | 6.68E-04 | 2572 | 263 | 87 | 0.330798479 | 0.03382582 | GO:0031325 | GO:BP | positive regulation of cellular metabolic process |
| Fetal mass | M12 | 1.08E-03 | 3061 | 263 | 98 | 0.372623574 | 0.03201568 | GO:0006996 | GO:BP | organelle organization |
| Fetal mass | M12 | 1.18E-03 | 2563 | 263 | 86 | 0.326996198 | 0.03355443 | GO:0051252 | GO:BP | regulation of RNA metabolic process |
| Fetal mass | M12 | 1.59E-03 | 2416 | 263 | 82 | 0.311787072 | 0.0339404 | GO:0051173 | GO:BP | positive regulation of nitrogen compound metabolic process |
| Fetal mass | M12 | 1.70E-03 | 6873 | 263 | 178 | 0.676806084 | 0.02589844 | GO:0043170 | GO:BP | macromolecule metabolic process |
| Fetal mass | M12 | 2.34E-03 | 3369 | 263 | 104 | 0.395437262 | 0.03086969 | GO:0010468 | GO:BP | regulation of gene expression |
| Fetal mass | M12 | 3.37E-03 | 1552 | 263 | 59 | 0.224334601 | 0.03801546 | GO:0045935 | GO:BP | positive regulation of nucleobase-containing compound metabolic process |
| Fetal mass | M12 | 3.98E-03 | 2718 | 263 | 88 | 0.33460076 | 0.03237675 | GO:0010556 | GO:BP | regulation of macromolecule biosynthetic process |
| Fetal mass | M12 | 7.11E-03 | 4162 | 263 | 120 | 0.456273764 | 0.02883229 | GO:0006139 | GO:BP | nucleobase-containing compound metabolic process |
| Fetal mass | M12 | 1.04E-02 | 2906 | 263 | 91 | 0.346007605 | 0.03131452 | GO:0009889 | GO:BP | regulation of biosynthetic process |
| Fetal mass | M12 | 1.11E-02 | 2324 | 263 | 77 | 0.292775665 | 0.03313253 | GO:0006355 | GO:BP | regulation of transcription |
| Fetal mass | M12 | 1.13E-02 | 2325 | 263 | 77 | 0.292775665 | 0.03311828 | GO:1903506 | GO:BP | regulation of nucleic acid-templated transcription |
| Fetal mass | M12 | 1.26E-02 | 2332 | 263 | 77 | 0.292775665 | 0.03301887 | GO:2001141 | GO:BP | regulation of RNA biosynthetic process |
| Fetal mass | M12 | 1.48E-02 | 2844 | 263 | 89 | 0.338403042 | 0.03129395 | GO:0031326 | GO:BP | regulation of cellular biosynthetic process |
| Fetal mass | M12 | 1.63E-02 | 4271 | 263 | 121 | 0.460076046 | 0.0283306 | GO:0046483 | GO:BP | heterocycle metabolic process |
| Fetal mass | M12 | 1.87E-02 | 7597 | 263 | 188 | 0.714828897 | 0.02474661 | GO:0050789 | GO:BP | regulation of biological process |
| Fetal mass | M12 | 1.94E-02 | 1372 | 263 | 52 | 0.197718631 | 0.03790087 | GO:0051254 | GO:BP | positive regulation of RNA metabolic process |
| Fetal mass | M12 | 2.10E-02 | 1681 | 263 | 60 | 0.228136882 | 0.03569304 | GO:0006357 | GO:BP | regulation of transcription by RNA polymerase II |
| Fetal mass | M12 | 2.19E-02 | 2658 | 263 | 84 | 0.319391635 | 0.03160271 | GO:0010604 | GO:BP | positive regulation of macromolecule metabolic process |
| Fetal mass | M12 | 2.77E-02 | 4313 | 263 | 121 | 0.460076046 | 0.02805472 | GO:0006725 | GO:BP | cellular aromatic compound metabolic process |
| Fetal mass | M12 | 2.83E-02 | 1736 | 263 | 61 | 0.231939163 | 0.03513825 | GO:0006366 | GO:BP | transcription by RNA polymerase II |
| Fetal mass | M12 | 2.93E-02 | 124 | 263 | 12 | 0.045627376 | 0.09677419 | GO:0031497 | GO:BP | chromatin assembly |
| Fetal mass | M12 | 3.03E-02 | 7270 | 263 | 181 | 0.688212928 | 0.02489684 | GO:0050794 | GO:BP | regulation of cellular process |
| Fetal mass | M12 | 3.13E-02 | 2895 | 263 | 89 | 0.338403042 | 0.03074266 | GO:0009893 | GO:BP | positive regulation of metabolic process |
| Fetal mass | M12 | 4.34E-02 | 3354 | 263 | 99 | 0.376425856 | 0.02951699 | GO:0009059 | GO:BP | macromolecule biosynthetic process |

| | | | | | | | | | | |
|---|---|---|---|---|---|---|---|---|---|---|
| Fetal mass | M12 | 4.60E-02 | 2413 | 263 | 77 | 0.292775665 | 0.03191048 | GO:0006351 | GO:BP | transcription |
| Fetal mass | M12 | 4.65E-02 | 249 | 263 | 17 | 0.064638783 | 0.06827309 | GO:0018105 | GO:BP | peptidyl-serine phosphorylation |
| Fetal mass | M12 | 4.67E-02 | 2414 | 263 | 77 | 0.292775665 | 0.03189727 | GO:0097659 | GO:BP | nucleic acid-templated transcription |
| Fetal mass | M12 | 8.53E-07 | 5667 | 263 | 162 | 0.615969582 | 0.02858655 | GO:0005634 | GO:CC | nucleus |
| Fetal mass | M12 | 3.91E-04 | 3126 | 263 | 98 | 0.372623574 | 0.03134997 | GO:0005654 | GO:CC | nucleoplasm |
| Fetal mass | M12 | 1.31E-03 | 10504 | 263 | 239 | 0.908745247 | 0.02275324 | GO:0005622 | GO:CC | intracellular anatomical structure |
| Fetal mass | M12 | 1.45E-03 | 3565 | 263 | 106 | 0.403041825 | 0.02973352 | GO:0031981 | GO:CC | nuclear lumen |
| Fetal mass | M12 | 2.39E-03 | 1007 | 263 | 42 | 0.159695817 | 0.04170804 | GO:0005694 | GO:CC | chromosome |
| Fetal mass | M12 | 2.29E-02 | 197 | 263 | 14 | 0.053231939 | 0.07106599 | GO:0036464 | GO:CC | cytoplasmic ribonucleoprotein granule |
| Fetal mass | M12 | 3.19E-02 | 9660 | 263 | 221 | 0.840304183 | 0.02287785 | GO:0043226 | GO:CC | organelle |
| Fetal mass | M12 | 4.11E-02 | 9510 | 263 | 218 | 0.828897338 | 0.02292324 | GO:0043229 | GO:CC | intracellular organelle |
| Fetal mass | M12 | 3.27E-05 | 1575 | 263 | 63 | 0.239543726 | 0.04 | GO:0003677 | GO:MF | DNA binding |
| Fetal mass | M12 | 4.24E-04 | 2484 | 263 | 83 | 0.315589354 | 0.03341385 | GO:0003676 | GO:MF | nucleic acid binding |
| Fetal mass | M12 | 2.20E-03 | 33 | 263 | 7 | 0.02661597 | 0.21212121 | GO:0140658 | GO:MF | ATP-dependent chromatin remodeler activity |
| Fetal mass | M12 | 2.92E-03 | 7380 | 263 | 184 | 0.699619772 | 0.02493225 | GO:0005515 | GO:MF | protein binding |
| Fetal mass | M12 | 1.27E-02 | 1140 | 263 | 44 | 0.16730038 | 0.03859649 | GO:0140110 | GO:MF | transcription regulator activity |
| Fetal mass | M12 | 1.49E-02 | 9925 | 263 | 227 | 0.863117871 | 0.02287154 | GO:0005488 | GO:MF | binding |
| Fetal mass | M12 | 1.65E-02 | 4326 | 263 | 119 | 0.452471483 | 0.02750809 | GO:0043167 | GO:MF | ion binding |
| Fetal mass | M12 | 2.50E-02 | 358 | 263 | 20 | 0.076045627 | 0.05586592 | GO:0004674 | GO:MF | protein serine/threonine kinase activity |
| Fetal mass | M12 | 2.63E-02 | 4089 | 263 | 113 | 0.429657795 | 0.02763512 | GO:1901363 | GO:MF | heterocyclic compound binding |
| Fetal mass | M12 | 2.72E-02 | 4138 | 263 | 114 | 0.433460076 | 0.02754954 | GO:0097159 | GO:MF | organic cyclic compound binding |
| Fetal mass | M12 | 3.82E-02 | 2911 | 263 | 86 | 0.326996198 | 0.02954311 | GO:0046872 | GO:MF | metal ion binding |
| Fetal mass | M12 | 4.76E-02 | 2972 | 263 | 87 | 0.330798479 | 0.02927322 | GO:0043169 | GO:MF | cation binding |
| Fetal mass | M12 | 6.90E-05 | 522 | 263 | 32 | 0.121673004 | 0.06130268 | HP:0000729 | HP | Autistic behavior |
| Fetal mass | M12 | 7.52E-05 | 1575 | 263 | 64 | 0.243346008 | 0.04063492 | HP:0000163 | HP | Abnormal oral cavity morphology |
| Fetal mass | M12 | 7.52E-05 | 1575 | 263 | 64 | 0.243346008 | 0.04063492 | HP:0031816 | HP | Abnormal oral morphology |

| | | | | | | | | | | |
|---|---|---|---|---|---|---|---|---|---|---|
| Fetal mass | M12 | 8.54E-05 | 890 | 263 | 44 | 0.16730038 | 0.0494382 | HP:0000290 | HP | Abnormality of the forehead |
| Fetal mass | M12 | 1.36E-04 | 840 | 263 | 42 | 0.159695817 | 0.05 | HP:0000174 | HP | Abnormal palate morphology |
| Fetal mass | M12 | 3.18E-04 | 1636 | 263 | 64 | 0.243346008 | 0.0391198 | HP:0000153 | HP | Abnormality of the mouth |
| Fetal mass | M12 | 4.72E-04 | 1843 | 263 | 69 | 0.262357414 | 0.03743896 | HP:0012373 | HP | Abnormal eye physiology |
| Fetal mass | M12 | 7.01E-04 | 577 | 263 | 32 | 0.121673004 | 0.05545927 | HP:0000218 | HP | High palate |
| Fetal mass | M12 | 1.15E-03 | 809 | 263 | 39 | 0.148288973 | 0.04820766 | HP:0000486 | HP | Strabismus |
| Fetal mass | M12 | 1.20E-03 | 843 | 263 | 40 | 0.152091255 | 0.04744958 | HP:0000549 | HP | Abnormal conjugate eye movement |
| Fetal mass | M12 | 1.28E-03 | 878 | 263 | 41 | 0.155893536 | 0.04669704 | HP:0000811 | HP | Abnormal external genitalia |
| Fetal mass | M12 | 1.36E-03 | 847 | 263 | 40 | 0.152091255 | 0.0472255 | HP:0000032 | HP | Abnormality of male external genitalia |
| Fetal mass | M12 | 1.71E-03 | 921 | 263 | 42 | 0.159695817 | 0.04560261 | HP:0001999 | HP | Abnormal facial shape |
| Fetal mass | M9 | 0.02560552 | 706 | 119 | 18 | 0.1512605 | 0.02549575 | GO:0005815 | GO:CC | microtubule organizing center |
| Fetal mass | M9 | 0.03078101 | 2 | 119 | 2 | 0.01680672 | 1 | GO:0071540 | GO:CC | eukaryotic translation initiation factor 3 complex |
| Fetal mass | M9 | 0.03330231 | 1776 | 119 | 32 | 0.26890756 | 0.01801802 | GO:0005856 | GO:CC | cytoskeleton |

**Table S13** Full list of genes under positive selection in highland deer mice as determined using a two-way outlier analysis (PBS and RDA; see text for further info). For each gene, the window location and SNP locations for which PBS and/or RDA values were calculated are provided. The PBS value and percentile within the simulated distribution for each window is shown. Those genes that are expressed in the labyrinth zone (LZ) and junctional zone/decidua (JZ/Dec.) are indicated as "Y" (expressed) or blank (not expressed).

| gene_symbol_pman | gene_mus | Window.Location | SNP.Location | PBS.ME.max | PBS.ME.max.perc | RDA.max | Expressed in LZ | Expressed in JZ/Dec. |
|---|---|---|---|---|---|---|---|---|
| A1bg | A1bg | NW_006501430.1:827501-832500 | NW_006501430.1:830147 | 0.283534348 | 1 | 0.544211747 | | |
| Abcc2 | Abcc2 | NW_006501599.1:505001-510000 | NW_006501599.1:507577 | 0.416975332 | 1 | 0.781229238 | Y | Y |
| Abhd18 | Abhd18 | NW_006501366.1:1787501-1792500 | NW_006501366.1:1791309 | 0.447479165 | 1 | 0.389537347 | Y | Y |
| Abhd3 | Abhd3 | NW_006501454.1:1802501-1807500 | NW_006501454.1:1807118 | 0.161528781 | 1 | 0.537948711 | Y | Y |
| Abhd6 | Abhd6 | NW_006501835.1:890001-895000 | NW_006501835.1:870359 | 0.112540066 | 0.999058879 | 0.61024541 | Y | Y |
| Acad8 | Acad8 | NW_006501112.1:4662501-4667500 | NW_006501112.1:4672768 | 0.295518056 | 1 | 0.531793339 | Y | Y |
| Acox2 | Acox2 | NW_006501835.1:612501-617500 | NW_006501835.1:596073 | 0.168225569 | 1 | 0.500623836 | Y | |
| Actl6a | Actl6a | NW_006501046.1:3985001-3990000 | NW_006501046.1:3987074 | 0.554147494 | 1 | 0.689379431 | Y | Y |
| Actrt3 | Actrt3 | NW_006501046.1:6697501-6702500 | NW_006501046.1:6696749 | 0.337193517 | 1 | 0.693346058 | | |
| Adamts15 | Adamts15 | NW_006501112.1:610001-615000 | NW_006501112.1:616056 | 0.206150374 | 1 | 0.561964136 | Y | Y |
| Adamts8 | Adamts8 | NW_006501112.1:545001-550000 | NW_006501112.1:548485 | 0.160311353 | 1 | 0.625616191 | Y | Y |
| Adcy8 | Adcy8 | NW_006501177.1:1012501-1017500 | NW_006501177.1:1007397 | 0.149273963 | 0.99982354 | 0.475886294 | | Y |
| Adgrf4 | Adgrf4 | NW_006502238.1:12501-17500 | NW_006502238.1:13128 | 0.123079934 | 0.999529439 | 0.409842765 | Y | Y |
| Adgrf5 | Adgrf5 | NW_006501862.1:90001-95000 | NW_006501862.1:58249 | 0.160029046 | 1 | 0.451704616 | Y | Y |
| Adgrg4 | Adgrg4 | NW_006501595.1:1870001-1875000 | NW_006501595.1:1896788 | 0.802841375 | 1 | 0.55210245 | | |
| Aff2 | Aff2 | NW_006501704.1:907501-912500 | NW_006501704.1:910760 | 1.787702494 | 1 | 0.790210629 | | |
| Agap2 | Agap2 | NW_006501066.1:687501-692500 | NW_006501066.1:704083 | 0.309007493 | 1 | 0.601906845 | Y | Y |
| Agtr1 | Agtr1b | NW_006501722.1:142501-147500 | NW_006501722.1:146500 | 0.652593445 | 1 | 0.701479994 | | |
| Alg5 | Alg5 | NW_006501125.1:3627501-3632500 | NW_006501125.1:3616033 | 0.345572487 | 1 | 0.449335771 | Y | Y |
| Alkbh8 | Alkbh8 | NW_006501598.1:197501-202500 | NW_006501598.1:218046 | 0.913662232 | 1 | 0.806348448 | Y | Y |
| Alms1 | Alms1 | NW_006501567.1:2145001-2150000 | NW_006501567.1:2148838 | 0.177684152 | 1 | 0.63672935 | Y | Y |
| Angel2 | Angel2 | NW_006501054.1:6077501-6082500 | NW_006501054.1:6078729 | 0.23293253 | 1 | 0.618214668 | Y | Y |

| | | | | | | | | |
|---|---|---|---|---|---|---|---|---|
| Angpt1 | Angpt1 | NW_006501143.1:3235001-3240000 | NW_006501143.1:3032740 | 0.991325304 | 1 | 0.735421979 | Y | |
| Ankrd13c | Ankrd13c | NW_006501088.1:1047501-1052500 | NW_006501088.1:1040042 | 0.300427139 | 1 | 0.418468889 | Y | Y |
| Ankrd50 | Ankrd50 | NW_006501581.1:97501-102500 | NW_006501581.1:110030 | 0.283508806 | 1 | 0.645152852 | Y | Y |
| Ankrd52 | Ankrd52 | NW_006501362.1:622501-627500 | NW_006501362.1:619441 | 0.215985113 | 1 | 0.785898537 | Y | Y |
| Ankrd60 | Ankrd60 | NW_006501107.1:445001-450000 | NW_006501107.1:444514 | 0.55071668 | 1 | 0.793872895 | | |
| Anln | Anln | NW_006501211.1:2210001-2215000 | NW_006501211.1:2261651 | 0.367013259 | 1 | 0.593534918 | Y | Y |
| Antxr1 | Antxr1 | NW_006501269.1:702501-707500 | NW_006501269.1:752668 | 0.132136106 | 0.99976472 | 0.36817265 | Y | Y |
| Anxa4 | Anxa4 | NW_006501269.1:1187501-1192500 | NW_006501269.1:1192350 | 0.332279742 | 1 | 0.50933156 | Y | Y |
| Anxa5 | Anxa5 | NW_006501046.1:312501-317500 | NW_006501046.1:325989 | 0.217165005 | 1 | 0.462586101 | Y | Y |
| Anxa7 | Anxa7 | NW_006501274.1:3835001-3840000 | NW_006501274.1:3850903 | 0.397774998 | 1 | 0.559275198 | Y | Y |
| Ap3m1 | Ap3m1 | NW_006501274.1:3235001-3240000 | NW_006501274.1:3259207 | 0.210650409 | 1 | 0.64466678 | Y | Y |
| Apbb3 | Sra1 | NW_006502100.1:15001-20000 | NW_006502100.1:12046 | 0.364435967 | 1 | 0.724587483 | Y | Y |
| Aplp2 | Aplp2 | NW_006501112.1:277501-282500 | NW_006501112.1:308703 | 0.521052622 | 1 | 0.542845654 | Y | Y |
| Arcn1 | Arcn1 | NW_006501694.1:347501-352500 | NW_006501694.1:350706 | 0.23706625 | 1 | 0.759150047 | Y | Y |
| Arhgap32 | Arhgap32 | NW_006501788.1:525001-530000 | NW_006501788.1:538884 | 0.342528757 | 1 | 0.781062003 | Y | Y |
| Arhgap42 | Arhgap42 | NW_006501245.1:4177501-4182500 | NW_006501245.1:4391964 | 0.607013142 | 1 | 0.698487129 | Y | Y |
| Arhgap9 | Arhgap9 | NW_006501066.1:445001-450000 | NW_006501066.1:447949 | 0.172793847 | 1 | 0.564196651 | Y | Y |
| Arhgef12 | Arhgef12 | NW_006501191.1:3165001-3170000 | NW_006501191.1:3087808 | 0.828966735 | 1 | 0.694212796 | Y | Y |
| Arhgef26 | Arhgef26 | NW_006501060.1:1377501-1382500 | NW_006501060.1:1392060 | 0.761627502 | 1 | 0.64130005 | Y | Y |
| Armc1 | Armc1 | NW_006501084.1:1307501-1312500 | NW_006501084.1:1311989 | 0.233403986 | 1 | 0.743757672 | Y | Y |
| As3mt | As3mt | NW_006501231.1:4935001-4940000 | NW_006501231.1:4910401 | 0.131206092 | 0.99976472 | 0.317258662 | Y | Y |
| Asb15 | Asb15 | NW_006501405.1:1650001-1655000 | NW_006501405.1:1664587 | 0.372962038 | 1 | 0.674067693 | | |
| Asb3 | Asb3 | NW_006502047.1:187501-192500 | NW_006502047.1:31847 | 0.117658599 | 0.999352979 | 0.386048358 | Y | Y |
| Asb4 | Asb4 | NW_006501346.1:622501-627500 | NW_006501346.1:636886 | 0.345294824 | 1 | 0.39073801 | Y | |
| Ash1l | Ash1l | NW_006501110.1:3235001-3240000 | NW_006501110.1:3260376 | 0.116603557 | 0.999235339 | 0.410840366 | Y | Y |
| Asxl3 | Asxl3 | NW_006501038.1:8445001-8450000 | NW_006501038.1:8526801 | 0.219670478 | 1 | 0.608499362 | Y | |
| Atad2 | Atad2 | NW_006501508.1:1967501-1972500 | NW_006501508.1:1989277 | 0.37163254 | 1 | 0.340704091 | Y | Y |
| Atp10a | Atp10a | NW_006501596.1:605001-610000 | NW_006501596.1:567825 | 0.200116458 | 1 | 0.442724936 | Y | Y |

| | | | | | | | | |
|---|---|---|---|---|---|---|---|---|
| Atp11b | Atp11b | NW_006501046.1:922501-927500 | NW_006501046.1:858297 | 0.340132528 | 1 | 0.453247699 | Y | Y |
| Atp5b | Atp5b | NW_006501362.1:987501-992500 | NW_006501362.1:993426 | 0.144260904 | 0.99982354 | 0.417859531 | Y | Y |
| Atp6ap1 | Atp6ap1 | NW_006501630.1:362501-367500 | NW_006501630.1:365361 | 0.449552584 | 1 | 0.502156682 | Y | Y |
| Atp6v0a1 | Atp6v0a1 | NW_006501663.1:1152501-1157500 | NW_006501663.1:1185082 | 0.152487172 | 0.99988236 | 0.332833855 | Y | Y |
| Atp6v0d1 | Atp6v0d1 | NW_006501344.1:765001-770000 | NW_006501344.1:803568 | 0.310370468 | 1 | 0.409785802 | Y | Y |
| Avil | Avil | NW_006501066.1:755001-760000 | NW_006501066.1:759527 | 0.162383153 | 1 | 0.58184901 | Y | Y |
| Azin1 | Azin1 | NW_006501220.1:2565001-2570000 | NW_006501220.1:2569686 | 0.757033614 | 1 | 0.769726181 | Y | Y |
| B3gnt9 | B3gnt9 | NW_006501344.1:452501-457500 | NW_006501344.1:457724 | 0.126624338 | 0.99964708 | 0.521336444 | Y | Y |
| B4galnt1 | B4galnt1 | NW_006501066.1:592501-597500 | NW_006501066.1:600801 | 0.148916153 | 0.99982354 | 0.514829004 | Y | Y |
| B4galt6 | B4galt6 | NW_006501038.1:6680001-6685000 | NW_006501038.1:6637722 | 0.763784502 | 1 | 0.699318518 | Y | Y |
| Baalc | AC164883.2 | NW_006501220.1:2945001-2950000 | NW_006501220.1:2949943 | 0.11675271 | 0.999235339 | 0.547547007 | | |
| Baz2a | Baz2a | NW_006501362.1:947501-952500 | NW_006501362.1:963714 | 0.429382584 | 1 | 0.774005678 | Y | Y |
| Bbs12 | Bbs12 | NW_006501581.1:1252501-1257500 | NW_006501581.1:1249094 | 0.222355402 | 1 | 0.74411241 | Y | Y |
| Bbs9 | Bbs9 | NW_006501574.1:3140001-3145000 | NW_006501574.1:2834945 | 0.344522595 | 1 | 0.483025221 | Y | Y |
| Bcl9l | Bcl9l | NW_006501694.1:85001-90000 | NW_006501694.1:97164 | 0.430050743 | 1 | 0.526903766 | Y | Y |
| Bend6 | Bend6 | NW_006503250.1:20001-25000 | NW_006503250.1:37305 | 0.264316822 | 1 | 0.498279504 | Y | Y |
| Birc2 | Birc2 | NW_006501245.1:3167501-3172500 | NW_006501245.1:3153430 | 0.400799008 | 1 | 0.775478144 | Y | Y |
| Blcap | Blcap | NW_006501257.1:620001-625000 | NW_006501257.1:625056 | 0.956193548 | 1 | 0.780196031 | Y | Y |
| Bmper | Bmper | NW_006501574.1:2337501-2342500 | NW_006501574.1:2348977 | 0.456196448 | 1 | 0.666483314 | Y | Y |
| Brwd1 | Brwd1 | NW_006501195.1:2487501-2492500 | NW_006501195.1:2468018 | 0.411595541 | 1 | 0.703091653 | Y | Y |
| Bsg | Bsg | NW_006501136.1:2387501-2392500 | NW_006501136.1:2392644 | 0.234039228 | 1 | 0.417213519 | Y | Y |
| Bud13 | Bud13 | NW_006501163.1:3185001-3190000 | NW_006501163.1:3186863 | 1.057862959 | 1 | 0.802969457 | Y | Y |
| C1qtnf5 | C1qtnf5 | NW_006501652.1:320001-325000 | NW_006501652.1:323436 | 0.269861157 | 1 | 0.505239949 | Y | Y |
| C2cd2l | C2cd2l | NW_006501652.1:532501-537500 | NW_006501652.1:528491 | 0.362749401 | 1 | 0.39524996 | Y | Y |
| Cacna1c | Cacna1c | NW_006501314.1:1657501-1662500 | NW_006501314.1:1689316 | 0.144522885 | 0.99982354 | 0.285267577 | Y | Y |
| Cacna2d1 | Cacna2d1 | NW_006501045.1:3835001-3840000 | NW_006501045.1:3464920 | 0.117138805 | 0.999294159 | 0.28924877 | Y | Y |
| Cacul1 | Cacul1 | NW_006501138.1:6127501-6132500 | NW_006501138.1:6112659 | 0.172970282 | 1 | 0.34239157 | Y | Y |
| Cadm1 | Cadm1 | NW_006501163.1:1602501-1607500 | NW_006501163.1:1552848 | 1.04073232 | 1 | 0.831360416 | Y | Y |

| | | | | | | | | |
|---|---|---|---|---|---|---|---|---|
| Cadps2 | Cadps2 | NW_006501405.1:462501-467500 | NW_006501405.1:408217 | 0.932137711 | 1 | 0.394916249 | Y | Y |
| Camk4 | Camk4 | NW_006501133.1:2342501-2347500 | NW_006501133.1:2559740 | 0.122696515 | 0.999470619 | 0.57378047 | | |
| Cand1 | Cand1 | NW_006501066.1:8457501-8462500 | NW_006501066.1:8458521 | 0.134922939 | 0.99982354 | 0.481610394 | Y | Y |
| Casd1 | Casd1 | NW_006501550.1:225001-230000 | NW_006501550.1:202587 | 0.789135003 | 1 | 0.578530138 | Y | Y |
| Casp1 | Casp1 | NW_006501245.1:1017501-1022500 | NW_006501245.1:1019405 | 0.554444468 | 1 | 0.805627591 | Y | Y |
| Cbfb | Cbfb | NW_006501344.1:412501-417500 | NW_006501344.1:417230 | 0.162120221 | 1 | 0.71520474 | Y | Y |
| Cbl | Cbl | NW_006501652.1:372501-377500 | NW_006501652.1:369077 | 0.528075851 | 1 | 0.487141528 | Y | Y |
| Cbln4 | Cbln4 | NW_006501290.1:1257501-1262500 | NW_006501290.1:1262398 | 0.140970275 | 0.99982354 | 0.449012259 | | |
| Ccdc15 | Ccdc15 | NW_006501547.1:905001-910000 | NW_006501547.1:820696 | 0.584510562 | 1 | 0.603423893 | Y | Y |
| Ccdc151 | Ccdc151 | NW_006501211.1:1827501-1832500 | NW_006501211.1:1830300 | 0.145493954 | 0.99982354 | 0.307693443 | | |
| Ccdc18 | Ccdc18 | NW_006501309.1:665001-670000 | NW_006501309.1:629267 | 0.213031265 | 1 | 0.353114605 | Y | Y |
| Ccna1 | Ccna1 | NW_006501125.1:3312501-3317500 | NW_006501125.1:3309688 | 0.359245255 | 1 | 0.746873768 | | |
| Ccna2 | Ccna2 | NW_006501581.1:1947501-1952500 | NW_006501581.1:1940669 | 0.391407731 | 1 | 0.303911538 | Y | Y |
| Cd180 | Cd180 | NW_006501218.1:4225001-4230000 | NW_006501218.1:4232567 | 0.137253308 | 0.99982354 | 0.719827809 | Y | Y |
| Cdh16 | Cdh16 | NW_006501139.1:3257501-3262500 | NW_006501139.1:3256802 | 0.221550261 | 1 | 0.599433912 | Y | |
| Cdh2 | Cdh2 | NW_006501225.1:1105001-1110000 | NW_006501225.1:1139714 | 0.308043426 | 1 | 0.749421089 | Y | |
| Cdh20 | Cdh20 | NW_006501753.1:1212501-1217500 | NW_006501753.1:1202425 | 0.148644438 | 0.99982354 | 0.603745695 | Y | |
| Cdh7 | Cdh7 | NW_006501661.1:1017501-1022500 | NW_006501661.1:1068508 | 0.313402858 | 1 | 0.716609283 | | |
| Cdk2 | Cdk2 | NW_006501362.1:300001-305000 | NW_006501362.1:306400 | 0.679034109 | 1 | 0.583536416 | Y | Y |
| Cdk8 | Cdk8 | NW_006501160.1:4107501-4112500 | NW_006501160.1:4080528 | 0.439972117 | 1 | 0.636284667 | Y | Y |
| Cdon | Cdon | NW_006501164.1:3130001-3135000 | NW_006501164.1:3152487 | 0.328397672 | 1 | 0.665666111 | Y | Y |
| Cep126 | Cep126 | NW_006501245.1:3415001-3420000 | NW_006501245.1:3436602 | 0.705925677 | 1 | 0.786868462 | Y | Y |
| Cep162 | Cep162 | NW_006501243.1:1292501-1297500 | NW_006501243.1:1328123 | 0.11905658 | 0.999352979 | 0.642562754 | Y | Y |
| Cep57 | Cep57 | NW_006501653.1:1297501-1302500 | NW_006501653.1:1293312 | 0.135407154 | 0.99982354 | 0.817593948 | Y | Y |
| Ces3 | Ces3a | NW_006501344.1:310001-315000 | NW_006501344.1:316914 | 0.269562307 | 1 | 0.571316646 | | |
| Cetn1 | Cetn1 | NW_006501756.1:1052501-1057500 | NW_006501756.1:1054621 | 0.172977304 | 1 | 0.259586789 | | |
| Cfap43 | Cfap43 | NW_006501231.1:3850001-3855000 | NW_006501231.1:3856193 | 0.321605525 | 1 | 0.446691488 | Y | Y |
| Cfap70 | Cfap70 | NW_006501274.1:3897501-3902500 | NW_006501274.1:3874820 | 0.188113659 | 1 | 0.444420927 | Y | Y |

| | | | | | | | | |
|---|---|---|---|---|---|---|---|---|
| Chek1 | Chek1 | NW_006501547.1:210001-215000 | NW_006501547.1:232000 | 0.132574224 | 0.99976472 | 0.580604739 | Y | Y |
| Chordc1 | Chordc1 | NW_006501646.1:1160001-1165000 | NW_006501646.1:1168717 | 0.281940408 | 1 | 0.747473775 | Y | Y |
| Cirbp | Cirbp | NW_006501136.1:1945001-1950000 | NW_006501136.1:1928330 | 0.112402937 | 0.999058879 | 0.460516885 | Y | Y |
| Cldn11 | Cldn11 | NW_006501046.1:7337501-7342500 | NW_006501046.1:7341401 | 0.267575422 | 1 | 0.742295373 | | |
| Clec16a | Clec16a | NW_006501248.1:4077501-4082500 | NW_006501248.1:3968274 | 0.126589036 | 0.99964708 | 0.363134148 | Y | Y |
| Clmp | Clmp | NW_006501191.1:762501-767500 | NW_006501191.1:767350 | 0.274697005 | 1 | 0.767933938 | Y | Y |
| Clnk | Clnk | NW_006502417.1:460001-465000 | NW_006502417.1:528101 | 0.740473188 | 1 | 0.392657241 | | Y |
| Clrn1 | Clrn1 | NW_006501814.1:402501-407500 | NW_006501814.1:369155 | 0.327984011 | 1 | 0.517707242 | | |
| Cmtm4 | Cmtm4 | NW_006501139.1:2992501-2997500 | NW_006501139.1:2992802 | 0.218694204 | 1 | 0.465255375 | Y | Y |
| Cntn5 | Cntn5 | NW_006501115.1:2977501-2982500 | NW_006501115.1:3229229 | 0.619205345 | 1 | 0.780666994 | | |
| Col12a1 | Col12a1 | NW_006501170.1:2170001-2175000 | NW_006501170.1:2162749 | 0.111722036 | 0.999000059 | 0.691761536 | Y | Y |
| Col14a1 | Col14a1 | NW_006501212.1:1762501-1767500 | NW_006501212.1:1691887 | 0.205275658 | 1 | 0.627423804 | Y | Y |
| Col1a2 | Col1a2 | NW_006501550.1:327501-332500 | NW_006501550.1:343862 | 0.880665904 | 1 | 0.554390488 | Y | Y |
| Col5a3 | Col5a3 | NW_006501353.1:902501-907500 | NW_006501353.1:909195 | 0.593223682 | 1 | 0.739961519 | Y | Y |
| Col9a1 | Col9a1 | NW_006501541.1:490001-495000 | NW_006501541.1:491134 | 0.137647848 | 0.99982354 | 0.450700909 | Y | |
| Colec10 | Colec10 | NW_006501212.1:2745001-2750000 | NW_006501212.1:2748388 | 0.212231829 | 1 | 0.518399688 | | |
| Colec12 | Colec12 | NW_006501756.1:815001-820000 | NW_006501756.1:828278 | 0.874898631 | 1 | 0.65829217 | Y | Y |
| Coq10a | Coq10a | NW_006501362.1:642501-647500 | NW_006501362.1:650068 | 0.552250887 | 1 | 0.677369249 | Y | Y |
| Cpa3 | Cpa3 | NW_006501722.1:272501-277500 | NW_006501722.1:283951 | 1.380931067 | 1 | 0.485607045 | | |
| Cpa4 | Cpa4 | NW_006501154.1:2877501-2882500 | NW_006501154.1:2892333 | 0.223658767 | 1 | 0.358912306 | | |
| Cpb1 | Cpb1 | NW_006501722.1:225001-230000 | NW_006501722.1:221496 | 1.052168822 | 1 | 0.735663502 | | |
| Cped1 | Cped1 | NW_006501785.1:637501-642500 | NW_006501785.1:506415 | 0.335709942 | 1 | 0.547127665 | Y | Y |
| Cpm | Cpm | NW_006501066.1:9872501-9877500 | NW_006501066.1:9859764 | 0.304863822 | 1 | 0.596243631 | Y | Y |
| Cpsf6 | Cpsf6 | NW_006501066.1:10150001-10155000 | NW_006501066.1:1017290 4 | 1.135731748 | 1 | 0.851877345 | Y | Y |
| Cr1l | Cr1l | NW_006501343.1:1617501-1622500 | NW_006501343.1:1603065 | 0.282701125 | 1 | 0.529680433 | Y | Y |
| Cramp1 | Cramp1l | NW_006501050.1:9825001-9830000 | NW_006501050.1:9829423 | 0.135212272 | 0.99982354 | 0.351648145 | Y | Y |
| Crip3 | Crip3 | NW_006501519.1:102501-107500 | NW_006501519.1:108271 | 0.122537911 | 0.999470619 | 0.475772636 | Y | |
| Crtam | Crtam | NW_006501191.1:957501-962500 | NW_006501191.1:940020 | 0.24638577 | 1 | 0.550873352 | | |

| | | | | | | | | |
|---|---|---|---|---|---|---|---|---|
| Cs | Cs | NW_006501362.1:657501-662500 | NW_006501362.1:651971 | 0.519378457 | 1 | 0.807122991 | | |
| Csmd3 | Csmd3 | NW_006501689.1:1497501-1502500 | NW_006501689.1:1082955 | 0.421715376 | 1 | 0.479065676 | | |
| Csn1s1 | NA | NW_006501573.1:70001-75000 | NW_006501573.1:62191 | 0.177875347 | 1 | 0.512709222 | | |
| Csn2 | Csn2 | NW_006501573.1:85001-90000 | NW_006501573.1:89817 | 0.148643935 | 0.99982354 | 0.698092571 | | |
| Cth | Cth | NW_006501088.1:1152501-1157500 | NW_006501088.1:1155292 | 0.129331163 | 0.99976472 | 0.426250711 | Y | Y |
| Ctsz | Ctsz | NW_006501107.1:1315001-1320000 | NW_006501107.1:1306134 | 1.094955139 | 1 | 0.417207041 | Y | Y |
| Cul5 | Cul5 | NW_006501057.1:350001-355000 | NW_006501057.1:356324 | 0.140863815 | 0.99982354 | 0.560104421 | Y | Y |
| Cul7 | Cul7 | NW_006501519.1:345001-350000 | NW_006501519.1:355697 | 0.219518742 | 1 | 0.460226805 | Y | Y |
| Cul9 | Cul9 | NW_006501519.1:205001-210000 | NW_006501519.1:209768 | 0.133702186 | 0.99982354 | 0.42079352 | Y | Y |
| Cwf19l1 | Cwf19l1 | NW_006501394.1:117501-122500 | NW_006501394.1:104893 | 0.142934591 | 0.99982354 | 0.318823423 | Y | Y |
| Cwf19l2 | Cwf19l2 | NW_006501598.1:270001-275000 | NW_006501598.1:334183 | 1.324226603 | 1 | 0.77744985 | Y | Y |
| Cxcl17 | NA | NW_006501390.1:1162501-1167500 | NW_006501390.1:1166833 | 0.112690186 | 0.999058879 | 0.295024443 | | |
| Cxcr5 | Cxcr5 | NW_006501694.1:100001-105000 | NW_006501694.1:103398 | 0.219234066 | 1 | 0.70400015 | | |
| Daam2 | Daam2 | NW_006501778.1:570001-575000 | NW_006501778.1:612718 | 0.20774485 | 1 | 0.431010509 | Y | Y |
| Dcaf13 | Dcaf13 | NW_006501220.1:3130001-3135000 | NW_006501220.1:3153194 | 0.281623346 | 1 | 0.523889862 | Y | Y |
| Dclk1 | Dclk1 | NW_006501125.1:3102501-3107500 | NW_006501125.1:2818003 | 0.52859024 | 1 | 0.71593618 | Y | Y |
| Dctn2 | Dctn2 | NW_006501066.1:485001-490000 | NW_006501066.1:485459 | 0.185811886 | 1 | 0.689450438 | Y | Y |
| Dcun1d1 | Dcun1d1 | NW_006501046.1:822501-827500 | NW_006501046.1:824235 | 0.504172984 | 1 | 0.702541285 | Y | Y |
| Ddit3 | Ddit3 | NW_006501066.1:472501-477500 | NW_006501066.1:473897 | 0.213492115 | 1 | 0.589091553 | Y | Y |
| Deptor | Deptor | NW_006501212.1:2052501-2057500 | NW_006501212.1:1950559 | 0.239143644 | 1 | 0.397999177 | Y | Y |
| Derl1 | Derl1 | NW_006501508.1:2250001-2255000 | NW_006501508.1:2258165 | 0.661075161 | 1 | 0.680122016 | Y | Y |
| Dgka | Dgka | NW_006501362.1:270001-275000 | NW_006501362.1:288315 | 0.413109304 | 1 | 0.798731323 | Y | Y |
| Dgkd | Dgkd | NW_006501258.1:3585001-3590000 | NW_006501258.1:3589108 | 0.138033497 | 0.99982354 | 0.704278557 | Y | Y |
| Dhdh | Dhdh | NW_006501325.1:2172501-2177500 | NW_006501325.1:2176489 | 0.123386422 | 0.999529439 | 0.419868181 | | |
| Dhx36 | Dhx36 | NW_006501060.1:1412501-1417500 | NW_006501060.1:1447593 | 0.862143868 | 1 | 0.670869333 | Y | Y |
| Diaph2 | Diaph2 | NW_006501330.1:327501-332500 | NW_006501330.1:80611 | 0.59681109 | 1 | 0.673828995 | Y | Y |
| Dido1 | Dido1 | NW_006501107.1:4755001-4760000 | NW_006501107.1:4759763 | 0.81757208 | 1 | 0.720651402 | Y | Y |
| Dlg5 | Dlg5 | NW_006501274.1:110001-115000 | NW_006501274.1:140340 | 0.431302879 | 1 | 0.387918413 | Y | Y |

| | | | | | | | | |
|---|---|---|---|---|---|---|---|---|
| Dmd | Dmd | NW_006501372.1:1072501-1077500 | NW_006501372.1:2462553 | 0.261286825 | 1 | 0.288793287 | Y | Y |
| Dnah12 | Dnah12 | NW_006501199.1:305001-310000 | NW_006501199.1:313297 | 0.207690493 | 1 | 0.257603282 | Y | |
| Dnajc14 | Dnajc14 | NW_006501362.1:182501-187500 | NW_006501362.1:187018 | 0.467237064 | 1 | 0.78586217 | Y | Y |
| Dnajc19 | Dnajc19 | NW_006501046.1:2635001-2640000 | NW_006501046.1:2636589 | 0.118128793 | 0.999352979 | 0.444503278 | | |
| Dnm2 | Dnm2 | NW_006501353.1:197501-202500 | NW_006501353.1:104557 | 0.35169325 | 1 | 0.734616888 | Y | Y |
| Dopey2 | Dopey2 | NW_006501195.1:172501-177500 | NW_006501195.1:178788 | 0.219856538 | 1 | 0.450498109 | | |
| Dpagt1 | Dpagt1 | NW_006501652.1:545001-550000 | NW_006501652.1:552127 | 0.255305448 | 1 | 0.472459483 | Y | Y |
| Dppa5 | Dppa5a | NW_006501895.1:185001-190000 | NW_006501895.1:189428 | 0.126681773 | 0.99964708 | 0.366082562 | | |
| Dpy19l1 | Dpy19l1 | NW_006501574.1:1430001-1435000 | NW_006501574.1:1435136 | 0.436186574 | 1 | 0.514531131 | Y | Y |
| Dpy19l2 | Dpy19l2 | NW_006501574.1:1295001-1300000 | NW_006501574.1:1298463 | 0.42732805 | 1 | 0.805619988 | | |
| Drd2 | Drd2 | NW_006501057.1:4870001-4875000 | NW_006501057.1:4861212 | 0.689518127 | 1 | 0.83332974 | | |
| Dsc1 | Dsc1 | NW_006501038.1:6145001-6150000 | NW_006501038.1:6142587 | 0.179828412 | 1 | 0.371668771 | | |
| Dsg3 | Dsg3 | NW_006501038.1:6487501-6492500 | NW_006501038.1:6502501 | 0.21595135 | 1 | 0.515728391 | | |
| Dsg4 | Dsg4 | NW_006501038.1:6410001-6415000 | NW_006501038.1:6432270 | 0.203309105 | 1 | 0.492972467 | | |
| Dst | Dst | NW_006502403.1:105001-110000 | NW_006502403.1:144069 | 0.282310167 | 1 | 0.601276498 | Y | Y |
| Dtna | Dtna | NW_006501796.1:592501-597500 | NW_006501796.1:593929 | 0.258422785 | 1 | 0.679213754 | Y | Y |
| Dtx3 | Dtx3 | NW_006501066.1:567501-572500 | NW_006501066.1:570557 | 0.228683484 | 1 | 0.497049366 | Y | Y |
| Dusp13 | Dusp13 | NW_006501274.1:2495001-2500000 | NW_006501274.1:2485633 | 0.455200701 | 1 | 0.611252156 | | |
| Dync1h1 | Dync1h1 | NW_006501607.1:950001-955000 | NW_006501607.1:981158 | 0.126089516 | 0.99958826 | 0.363206018 | Y | Y |
| Dync2h1 | Dync2h1 | NW_006501245.1:2427501-2432500 | NW_006501245.1:2399127 | 1.26810771 | 1 | 0.807863389 | Y | Y |
| Dyrk1a | Dyrk1a | NW_006501195.1:1097501-1102500 | NW_006501195.1:1134502 | 0.199863745 | 1 | 0.4052805 | Y | Y |
| Dyrk2 | Dyrk2 | NW_006501066.1:8807501-8812500 | NW_006501066.1:8812797 | 0.351878718 | 1 | 0.758474814 | Y | Y |
| Dysf | Dysf | NW_006501567.1:617501-622500 | NW_006501567.1:630824 | 0.164397725 | 1 | 0.589240167 | Y | Y |
| E2f4 | E2f4 | NW_006501344.1:505001-510000 | NW_006501344.1:510240 | 0.176267005 | 1 | 0.474461317 | Y | Y |
| E2f5 | E2f5 | NW_006501833.1:360001-365000 | NW_006501833.1:373006 | 0.343843658 | 1 | 0.474118039 | Y | Y |
| Edil3 | Edil3 | NW_006501180.1:1480001-1485000 | NW_006501180.1:1573523 | 0.197985556 | 1 | 0.449551272 | | Y |
| Edn3 | Edn3 | NW_006501107.1:1652501-1657500 | NW_006501107.1:1660223 | 1.070804909 | 1 | 0.65619666 | | |
| Eepd1 | Eepd1 | NW_006501574.1:307501-312500 | NW_006501574.1:297585 | 0.142815958 | 0.99982354 | 0.57997629 | Y | Y |

| | | | | | | | | |
|---|---|---|---|---|---|---|---|---|
| Eif2a | Eif2a | NW_006501814.1:32501-37500 | NW_006501814.1:33538 | 0.477722687 | 1 | 0.698861122 | Y | Y |
| Eif5a2 | Eif5a2 | NW_006501046.1:7882501-7887500 | NW_006501046.1:7875139 | 0.114983118 | 0.999235339 | 0.560537741 | Y | Y |
| Elf2 | Elf2 | NW_006501125.1:7227501-7232500 | NW_006501125.1:7228819 | 0.289850302 | 1 | 0.725393325 | Y | Y |
| Endod1 | Endod1 | NW_006501653.1:1845001-1850000 | NW_006501653.1:1846158 | 0.188095959 | 1 | 0.553728734 | | |
| Eno4 | Eno4 | NW_006501138.1:4492501-4497500 | NW_006501138.1:4505792 | 0.18839433 | 1 | 0.356974876 | Y | Y |
| Enpp2 | Enpp2 | NW_006501212.1:2400001-2405000 | NW_006501212.1:2410594 | 0.159462484 | 0.99994118 | 0.484052365 | Y | Y |
| Epas1 | Epas1 | NW_006501333.1:1627501-1632500 | NW_006501333.1:1625647 | 0.189730874 | 1 | 0.345037294 | Y | Y |
| Epb41l4a | Epb41l4a | NW_006501133.1:1750001-1755000 | NW_006501133.1:1755693 | 0.307231117 | 1 | 0.275052298 | Y | Y |
| Erap1 | Erap1 | NW_006501154.1:145001-150000 | NW_006501154.1:136507 | 0.347990164 | 1 | 0.774191423 | Y | Y |
| Erbb3 | Erbb3 | NW_006501362.1:427501-432500 | NW_006501362.1:447504 | 0.567655072 | 1 | 0.68460433 | Y | Y |
| Erbin | Erbin | NW_006501218.1:5267501-5272500 | NW_006501218.1:5270808 | 0.155655625 | 0.99994118 | 0.71665766 | Y | Y |
| Ercc4 | Ercc4 | NW_006501248.1:1460001-1465000 | NW_006501248.1:1460222 | 0.119948373 | 0.999352979 | 0.433897842 | Y | Y |
| Ercc6l2 | Ercc6l2 | NW_006501203.1:4605001-4610000 | NW_006501203.1:4601465 | 0.137059091 | 0.99982354 | 0.524200725 | Y | Y |
| Erg | Erg | NW_006501195.1:1837501-1842500 | NW_006501195.1:1921600 | 0.259462282 | 1 | 0.375484391 | Y | Y |
| Erich6 | Erich6 | NW_006501814.1:90001-95000 | NW_006501814.1:92566 | 0.594974217 | 1 | 0.641049636 | | |
| Esam | Esam | NW_006502995.1:52501-57500 | NW_006502995.1:59400 | 0.356471555 | 1 | 0.650160645 | Y | Y |
| Esyt1 | Esyt1 | NW_006501362.1:490001-495000 | NW_006501362.1:493447 | 0.254179771 | 1 | 0.751539903 | Y | Y |
| Exoc6b | Exoc6b | NW_006501567.1:1457501-1462500 | NW_006501567.1:1061086 | 1.251373948 | 1 | 0.687467007 | Y | Y |
| Exosc8 | Exosc8 | NW_006501125.1:3662501-3667500 | NW_006501125.1:3670022 | 0.632724059 | 1 | 0.530124362 | Y | Y |
| F2rl1 | F2rl1 | NW_006501306.1:92501-97500 | NW_006501306.1:85659 | 0.192498884 | 1 | 0.698852754 | Y | Y |
| Fam118b | Fam118b | NW_006501164.1:2855001-2860000 | NW_006501164.1:2853432 | 0.29280578 | 1 | 0.651225343 | Y | Y |
| Fam132a | Fam132a | NW_006501627.1:360001-365000 | NW_006501627.1:362803 | 0.152755148 | 0.99988236 | 0.289947227 | | |
| Fam132b | Fam132b | NW_006501416.1:3452501-3457500 | NW_006501416.1:3451382 | 0.195176477 | 1 | 0.323972302 | | |
| Fam135a | Fam135a | NW_006501541.1:297501-302500 | NW_006501541.1:272506 | 0.220267622 | 1 | 0.493529723 | Y | Y |
| Fam13b | Fam13b | NW_006501133.1:1050001-1055000 | NW_006501133.1:1092764 | 0.244964163 | 1 | 0.387716918 | Y | Y |
| Fam149b1 | Fam149b | NW_006501274.1:3955001-3960000 | NW_006501274.1:3936657 | 0.17758953 | 1 | 0.330430132 | Y | Y |
| Fam177a1 | Fam177a | NW_006501287.1:1220001-1225000 | NW_006501287.1:1214097 | 0.305118944 | 1 | 0.415439681 | Y | Y |
| Fam19a2 | Fam19a2 | NW_006501066.1:3925001-3930000 | NW_006501066.1:3927525 | 0.248597547 | 1 | 0.513105211 | | |

| | | | | | | | | |
|---|---|---|---|---|---|---|---|---|
| Fam3c | Fam3c | NW_006501785.1:437501-442500 | NW_006501785.1:456160 | 0.118805791 | 0.999352979 | 0.296224391 | Y | Y |
| Fam76b | Fam76b | NW_006501653.1:1317501-1322500 | NW_006501653.1:1334471 | 0.268466418 | 1 | 0.80433015 | Y | Y |
| Fam84b | Fam84b | NW_006501430.1:712501-717500 | NW_006501430.1:712366 | 0.225684687 | 1 | 0.464400534 | | |
| Fam91a1 | Fam91a1 | NW_006501508.1:1610001-1615000 | NW_006501508.1:1604675 | 0.183256364 | 1 | 0.651972528 | Y | Y |
| Faxc | Faxc | NW_006501152.1:4792501-4797500 | NW_006501152.1:4795275 | 0.183720778 | 1 | 0.594605798 | Y | Y |
| Fbxo11 | Fbxo11 | NW_006501333.1:417501-422500 | NW_006501333.1:419184 | 0.116658812 | 0.999235339 | 0.327558286 | Y | Y |
| Fbxo32 | Fbxo32 | NW_006501508.1:1890001-1895000 | NW_006501508.1:1897995 | 0.378021486 | 1 | 0.470744683 | Y | Y |
| Fer1l6 | Fer1l6 | NW_006501508.1:1432501-1437500 | NW_006501508.1:1402344 | 0.311956454 | 1 | 0.392250037 | | |
| Fgf2 | Fgf2 | NW_006501581.1:1170001-1175000 | NW_006501581.1:1155329 | 0.590887429 | 1 | 0.643338931 | | |
| Fli1 | Fli1 | NW_006501164.1:90001-95000 | NW_006501164.1:91264 | 0.123047578 | 0.999529439 | 0.591351461 | Y | Y |
| Flnb | Flnb | NW_006501835.1:1050001-1055000 | NW_006501835.1:983071 | 0.126646918 | 0.99964708 | 0.351951922 | Y | Y |
| Flt1 | Flt1 | NW_006501160.1:2612501-2617500 | NW_006501160.1:2736720 | 0.178013788 | 1 | 0.659207571 | Y | Y |
| Fmr1 | Fmr1 | NW_006506577.1:32501-37500 | NW_006506577.1:22266 | 0.355829436 | 1 | 0.590172381 | Y | Y |
| Fndc3b | Fndc3b | NW_006501046.1:9307501-9312500 | NW_006501046.1:9344832 | 0.178720713 | 1 | 0.635680067 | Y | Y |
| Foxo1 | Foxo1 | NW_006501125.1:6085001-6090000 | NW_006501125.1:6085691 | 0.172168587 | 1 | 0.67115353 | | |
| Foxp2 | Foxp2 | NW_006501259.1:3872501-3877500 | NW_006501259.1:3922867 | 0.132269604 | 0.99976472 | 0.366630941 | | |
| Frem2 | Frem2 | NW_006501125.1:4852501-4857500 | NW_006501125.1:4748263 | 0.302126625 | 1 | 0.715364129 | Y | |
| Frmd7 | Frmd7 | NW_006501915.1:870001-875000 | NW_006501915.1:876058 | 0.208392989 | 1 | 0.305727866 | | |
| Fry | Fry | NW_006501358.1:1347501-1352500 | NW_006501358.1:1445200 | 0.1409603 | 0.99982354 | 0.426699136 | Y | Y |
| Fsd1l | Fsd1l | NW_006501898.1:1410001-1415000 | NW_006501898.1:1425771 | 0.27107581 | 1 | 0.318795804 | Y | Y |
| Fut11 | Fut11 | NW_006501274.1:3602501-3607500 | NW_006501274.1:3598367 | 0.170328206 | 1 | 0.642169486 | Y | Y |
| Fxr1 | Fxr1 | NW_006501046.1:2650001-2655000 | NW_006501046.1:2633664 | 0.994381512 | 1 | 0.512174469 | Y | Y |
| Gabra3 | Gabra3 | NW_006501587.1:800001-805000 | NW_006501587.1:804307 | 0.724794454 | 1 | 0.566176406 | | |
| Gabrq | Gabrq | NW_006501587.1:1297501-1302500 | NW_006501587.1:1300279 | 1.529624818 | 1 | 0.656950653 | Y | Y |
| Galnt1 | Galnt1 | NW_006501173.1:3967501-3972500 | NW_006501173.1:3987075 | 0.144869284 | 0.99982354 | 0.387655967 | Y | Y |
| Gatad1 | Gatad1 | NW_006501550.1:2135001-2140000 | NW_006501550.1:2138577 | 0.127382816 | 0.9997059 | 0.44032185 | Y | Y |
| Gdf11 | Gdf11 | NW_006501362.1:115001-120000 | NW_006501362.1:123904 | 0.55312591 | 1 | 0.856954715 | Y | Y |
| Gfm2 | Gfm2 | NW_006501306.1:1802501-1807500 | NW_006501306.1:1796619 | 0.129451896 | 0.99976472 | 0.520606074 | Y | Y |

| | | | | | | | | |
|---|---|---|---|---|---|---|---|---|
| Gfpt1 | Gfpt1 | NW_006501269.1:815001-820000 | NW_006501269.1:802786 | 0.204971402 | 1 | 0.548213269 | Y | Y |
| Gfra3 | Gfra3 | NW_006501133.1:842501-847500 | NW_006501133.1:848356 | 0.174227475 | 1 | 0.428623225 | Y | |
| Glp2r | Glp2r | NW_006501067.1:2697501-2702500 | NW_006501067.1:2694960 | 0.132417549 | 0.99976472 | 0.319214976 | | |
| Gls2 | Gls2 | NW_006501362.1:860001-865000 | NW_006501362.1:842330 | 0.383969241 | 1 | 0.56190089 | Y | Y |
| Gltscr1l | Gltscr1l | NW_006501519.1:547501-552500 | NW_006501519.1:582306 | 0.139435648 | 0.99982354 | 0.623754902 | | |
| Gmcl1 | Gmcl1 | NW_006501269.1:1202501-1207500 | NW_006501269.1:1234513 | 0.255501566 | 1 | 0.29753839 | Y | Y |
| Gmps | Gmps | NW_006501060.1:2765001-2770000 | NW_006501060.1:2794934 | 0.793456815 | 1 | 0.608929039 | Y | Y |
| Gnb1 | Gnb1 | NW_006501627.1:772501-777500 | NW_006501627.1:776432 | 0.120258022 | 0.999470619 | 0.385105691 | Y | Y |
| Gnb4 | Gnb4 | NW_006501046.1:4145001-4150000 | NW_006501046.1:4146588 | 0.201579311 | 1 | 0.699791252 | Y | Y |
| Gns | Gns | NW_006501066.1:6250001-6255000 | NW_006501066.1:6221356 | 0.153510216 | 0.99988236 | 0.673534195 | Y | Y |
| Golga3 | Golga3 | NW_006501256.1:780001-785000 | NW_006501256.1:813976 | 0.187593432 | 1 | 0.382079519 | Y | Y |
| Gpn1 | Gpn1 | NW_006501275.1:950001-955000 | NW_006501275.1:938128 | 0.219445013 | 1 | 0.34011684 | Y | Y |
| Gpr149 | Gpr149 | NW_006501060.1:1530001-1535000 | NW_006501060.1:1468207 | 0.448011177 | 1 | 0.678880048 | | |
| Gpr182 | Gpr182 | NW_006503545.1:22501-27500 | NW_006503545.1:26699 | 0.181607913 | 1 | 0.672707251 | Y | |
| Gpr75 | Gpr75 | NW_006502047.1:262501-267500 | NW_006502047.1:267066 | 0.158599162 | 0.99994118 | 0.30809317 | | |
| Gpr87 | Gpr87 | NW_006501814.1:735001-740000 | NW_006501814.1:737703 | 0.483427412 | 1 | 0.676410538 | | |
| Gramd1b | Gramd1b | NW_006501191.1:247501-252500 | NW_006501191.1:247049 | 0.183732097 | 1 | 0.818115872 | Y | Y |
| Greb1l | Greb1l | NW_006501756.1:172501-177500 | NW_006501756.1:114783 | 0.312873976 | 1 | 0.647050506 | Y | Y |
| Grhl2 | Grhl2 | NW_006501220.1:1327501-1332500 | NW_006501220.1:1330840 | 0.209221404 | 1 | 0.587025973 | Y | Y |
| Gria4 | Gria4 | NW_006501245.1:37501-42500 | NW_006501245.1:40269 | 1.218582023 | 1 | 0.837457744 | | |
| Grik2 | Grik2 | NW_006501244.1:4105001-4110000 | NW_006501244.1:3766054 | 0.300255369 | 1 | 0.333793084 | Y | |
| Grip1 | Grip1 | NW_006501066.1:7597501-7602500 | NW_006501066.1:7601858 | 0.422684665 | 1 | 0.691183908 | Y | Y |
| Gspt1 | Gspt1 | NW_006501248.1:3372501-3377500 | NW_006501248.1:3402025 | 0.122789321 | 0.999470619 | 0.438598519 | Y | Y |
| Gucy1a2 | Gucy1a2 | NW_006501598.1:930001-935000 | NW_006501598.1:669362 | 0.768713555 | 1 | 0.826930211 | Y | |
| Gyg1 | Gyg | NW_006501722.1:402501-407500 | NW_006501722.1:406377 | 0.445765578 | 1 | 0.69644259 | Y | Y |
| Has2 | Has2 | NW_006501212.1:615001-620000 | NW_006501212.1:616119 | 0.212193664 | 1 | 0.729590363 | | Y |
| Helb | Helb | NW_006501066.1:7580001-7585000 | NW_006501066.1:7592453 | 0.183474838 | 1 | 0.510698833 | Y | Y |
| Hells | Hells | NW_006501115.1:107501-112500 | NW_006501115.1:94188 | 0.142021475 | 0.99982354 | 0.645507395 | Y | Y |

| | | | | | | | | |
|---|---|---|---|---|---|---|---|---|
| Hepacam | Hepacam | NW_006501547.1:922501-927500 | NW_006501547.1:945727 | 0.21371366 | 1 | 0.667677609 | | |
| Hepacam2 | Hepacam2 | NW_006501550.1:1297501-1302500 | NW_006501550.1:1338692 | 0.47218644 | 1 | 0.676420832 | | |
| Hinfp | Hinfp | NW_006501652.1:507501-512500 | NW_006501652.1:512478 | 0.337582442 | 1 | 0.389873022 | Y | Y |
| Hmbs | Hmbs | NW_006501652.1:560001-565000 | NW_006501652.1:555503 | 0.279722291 | 1 | 0.380197579 | Y | Y |
| Hps3 | Hps3 | NW_006501722.1:610001-615000 | NW_006501722.1:590083 | 1.573383306 | 1 | 0.737313798 | Y | Y |
| Hs6st1 | Hs6st1 | NW_006501361.1:435001-440000 | NW_006501361.1:438475 | 0.163051748 | 1 | 0.641095331 | Y | Y |
| Hsd11b1 | Hsd11b1 | NW_006501393.1:297501-302500 | NW_006501393.1:303334 | 0.432518578 | 1 | 0.316974718 | Y | Y |
| Hsd17b3 | Hsd17b3 | NW_006501203.1:4855001-4860000 | NW_006501203.1:4861410 | 0.452290391 | 1 | 0.685226291 | | |
| Hsf4 | Hsf4 | NW_006501344.1:470001-475000 | NW_006501344.1:480088 | 0.170484497 | 1 | 0.409782052 | Y | Y |
| Hspa4l | Hspa4l | NW_006501366.1:1650001-1655000 | NW_006501366.1:1682358 | 1.090963123 | 1 | 0.736019345 | Y | Y |
| Hspa8 | Hspa8 | NW_006501191.1:782501-787500 | NW_006501191.1:782912 | 0.169824012 | 1 | 0.680773135 | Y | Y |
| Hspa9 | Hspa9 | NW_006501133.1:560001-565000 | NW_006501133.1:576189 | 0.310072252 | 1 | 0.463932297 | Y | Y |
| Hsph1 | Hsph1 | NW_006501160.1:640001-645000 | NW_006501160.1:650251 | 0.124349607 | 0.999529439 | 0.459641543 | Y | Y |
| Htr1b | Htr1b | NW_006501170.1:300001-305000 | NW_006501170.1:301883 | 0.150458722 | 0.99982354 | 0.402275081 | Y | Y |
| Htr3a | Htr3a | NW_006501163.1:297501-302500 | NW_006501163.1:308731 | 0.598371905 | 1 | 0.733549602 | | |
| Hyls1 | Hyls1 | NW_006501164.1:3210001-3215000 | NW_006501164.1:3209490 | 0.416713357 | 1 | 0.421971291 | Y | Y |
| Hyou1 | Hyou1 | NW_006501652.1:605001-610000 | NW_006501652.1:617427 | 0.250064981 | 1 | 0.550971061 | Y | Y |
| Icam1 | Icam1 | NW_006501353.1:615001-620000 | NW_006501353.1:620294 | 0.290207121 | 1 | 0.437569804 | Y | Y |
| Icam4 | Icam4 | NW_006501353.1:615001-620000 | NW_006501353.1:618073 | 0.290207121 | 1 | 0.838355717 | | |
| Ifng | NA | NW_006501066.1:9240001-9245000 | NW_006501066.1:9245050 | 0.244088333 | 1 | 0.722863498 | | |
| Igsf10 | Igsf10 | NW_006501814.1:892501-897500 | NW_006501814.1:885044 | 0.683571263 | 1 | 0.717261365 | Y | Y |
| Igsf9b | Igsf9b | NW_006501112.1:4307501-4312500 | NW_006501112.1:4293294 | 0.263539536 | 1 | 0.837856448 | Y | Y |
| Ikzf4 | Ikzf4 | NW_006501362.1:360001-365000 | NW_006501362.1:373300 | 0.55775898 | 1 | 0.853702717 | Y | Y |
| Il18 | NA | NW_006501057.1:3625001-3630000 | NW_006501057.1:3621595 | 0.141224159 | 0.99982354 | 0.48262676 | Y | Y |
| Il23a | Il23a | NW_006501362.1:710001-715000 | NW_006501362.1:709268 | 0.17907092 | 1 | 0.514585882 | Y | Y |
| Ilf3 | Ilf3 | NW_006501353.1:227501-232500 | NW_006501353.1:216427 | 0.378087279 | 1 | 0.599929292 | Y | Y |
| Irak3 | Irak3 | NW_006501066.1:7482501-7487500 | NW_006501066.1:7528095 | 0.231830736 | 1 | 0.675044341 | Y | Y |
| Itga7 | Itga7 | NW_006501362.1:47501-52500 | NW_006501362.1:64209 | 0.565290246 | 1 | 0.336242404 | Y | Y |

| | | | | | | | | |
|---|---|---|---|---|---|---|---|---|
| Itgbl1 | Itgbl1 | NW_006501145.1:6057501-6062500 | NW_006501145.1:6167950 | 0.475778863 | 1 | 0.275052298 | | |
| Iws1 | Iws1 | NW_006501133.1:3410001-3415000 | NW_006501133.1:3402783 | 0.173793254 | 1 | 0.677235818 | Y | Y |
| Jade1 | Jade1 | NW_006501366.1:2467501-2472500 | NW_006501366.1:2407917 | 0.372640128 | 1 | 0.7101761 | Y | Y |
| Jam3 | Jam3 | NW_006501112.1:4525001-4530000 | NW_006501112.1:4532454 | 0.2892468 | 1 | 0.469298084 | Y | Y |
| Kank2 | Kank2 | NW_006501211.1:1587501-1592500 | NW_006501211.1:1578578 | 0.309676985 | 1 | 0.791650307 | Y | Y |
| Kat2b | Kat2b | NW_006501911.1:140001-145000 | NW_006501911.1:148082 | 0.386930142 | 1 | 0.713092456 | | |
| Kat6b | Kat6b | NW_006501274.1:2617501-2622500 | NW_006501274.1:2574615 | 0.686385896 | 1 | 0.709936939 | Y | Y |
| Kbtbd3 | Kbtbd3 | NW_006503040.1:70001-75000 | NW_006503040.1:74068 | 0.555222143 | 1 | 0.775812271 | Y | Y |
| Kcnh8 | Kcnh8 | NW_006501707.1:290001-295000 | NW_006501707.1:98121 | 0.368045772 | 1 | 0.352650197 | | |
| Kcnj1 | Kcnj1 | NW_006501164.1:50001-55000 | NW_006501164.1:58143 | 0.315416055 | 1 | 0.791548072 | | |
| Kcnj5 | Kcnj5 | NW_006501788.1:602501-607500 | NW_006501788.1:600155 | 0.246245834 | 1 | 0.492877757 | | |
| Kcnj6 | Kcnj6 | NW_006501195.1:1295001-1300000 | NW_006501195.1:1207168 | 0.321421572 | 1 | 0.639371148 | | |
| Kcnma1 | Kcnma1 | NW_006501274.1:965001-970000 | NW_006501274.1:976440 | 0.434716598 | 1 | 0.797004414 | | |
| Kcnq3 | Kcnq3 | NW_006501177.1:2225001-2230000 | NW_006501177.1:2176552 | 0.149815941 | 0.99982354 | 0.331623192 | | |
| Kcnv1 | Kcnv1 | NW_006501143.1:742501-747500 | NW_006501143.1:745510 | 0.121984544 | 0.999470619 | 0.516851632 | | |
| Kctd1 | Kctd1 | NW_006501225.1:2622501-2627500 | NW_006501225.1:2647599 | 0.219293 | 1 | 0.687004054 | Y | Y |
| Kdm3b | Kdm3b | NW_006501133.1:685001-690000 | NW_006501133.1:682748 | 0.307116717 | 1 | 0.399138769 | Y | Y |
| Kdm6a | Kdm6a | NW_006502276.1:197501-202500 | NW_006502276.1:201351 | 0.130110538 | 0.99976472 | 0.742559191 | Y | Y |
| Kiaa0196 | E430025E21Rik | NW_006501508.1:650001-655000 | NW_006501508.1:667404 | 0.589715908 | 1 | 0.758006264 | | |
| Kiaa0430 | Marf1 | NW_006501248.1:370001-375000 | NW_006501248.1:391782 | 0.114986597 | 0.999235339 | 0.39924592 | Y | Y |
| Kiaa0895 | 9530077C05Rik | NW_006501211.1:2322501-2327500 | NW_006501211.1:2325273 | 0.915428202 | 1 | 0.809870077 | Y | Y |
| Kiaa1109 | 4932438A13Rik | NW_006501581.1:1645001-1650000 | NW_006501581.1:1611466 | 0.682295111 | 1 | 0.739409069 | Y | Y |
| Kiaa1217 | Etl4 | NW_006501144.1:2865001-2870000 | NW_006501144.1:2934468 | 0.185911488 | 1 | 0.28924877 | Y | Y |
| Kiaa1328 | AW554918 | NW_006501048.1:5807501-5812500 | NW_006501048.1:5676525 | 0.117161246 | 0.999294159 | 0.613885219 | Y | Y |
| Kiaa2022 | C77370 | NW_006501568.1:1072501-1077500 | NW_006501568.1:1076209 | 0.219797436 | 1 | 0.564260462 | | |
| Kidins220 | Kidins220 | NW_006501185.1:327501-332500 | NW_006501185.1:351273 | 0.953773427 | 1 | 0.77414946 | Y | Y |
| Kif24 | Kif24 | NW_006501318.1:422501-427500 | NW_006501318.1:399092 | 0.1277294 | 0.9997059 | 0.379212202 | Y | Y |
| Kif5a | Kif5a | NW_006501066.1:525001-530000 | NW_006501066.1:542055 | 0.135016435 | 0.99982354 | 0.3059163 | Y | Y |

| | | | | | | | | |
|---|---|---|---|---|---|---|---|---|
| Kif6 | Kif6 | NW_006501778.1:307501-312500 | NW_006501778.1:416014 | 0.364314799 | 1 | 0.499432628 | | Y |
| Kl | Kl | NW_006501358.1:1942501-1947500 | NW_006501358.1:1984460 | 0.32154046 | 1 | 0.46258063 | | |
| Klc4 | Klc4 | NW_006501519.1:342501-347500 | NW_006501519.1:344415 | 0.232869632 | 1 | 0.580674142 | Y | Y |
| Klf10 | Klf10 | NW_006501220.1:2382501-2387500 | NW_006501220.1:2382676 | 0.150110462 | 0.99982354 | 0.513973991 | Y | Y |
| Klhdc3 | Klhdc3 | NW_006501519.1:375001-380000 | NW_006501519.1:372409 | 0.194626334 | 1 | 0.600687149 | Y | Y |
| Kmt2a | Kmt2a | NW_006501694.1:475001-480000 | NW_006501694.1:481688 | 1.186195412 | 1 | 0.831735656 | Y | Y |
| Lama3 | Lama3 | NW_006501225.1:5115001-5120000 | NW_006501225.1:5136099 | 0.23092404 | 1 | 0.459332125 | Y | Y |
| Lama5 | Lama5 | NW_006501107.1:4227501-4232500 | NW_006501107.1:4248060 | 0.170338412 | 1 | 0.510407914 | Y | Y |
| Large | Large | NW_006501868.1:235001-240000 | NW_006501868.1:247853 | 0.155240445 | 0.99994118 | 0.3914754 | | |
| Lhfp | Lhfp | NW_006501125.1:5342501-5347500 | NW_006501125.1:5140991 | 0.300672759 | 1 | 0.621516424 | Y | Y |
| LOC102902765 | Vmn1r224 | NW_006501759.1:477501-482500 | NW_006501759.1:479511 | 0.162885555 | 1 | 0.508460126 | | |
| LOC102902789 | Cpq | NW_006501126.1:2445001-2450000 | NW_006501126.1:2447846 | 0.198477547 | 1 | 0.476931349 | Y | Y |
| LOC102903073 | Zfp160 | NW_006501916.1:32501-37500 | NW_006501916.1:22959 | 0.137719809 | 0.99982354 | 0.534776054 | Y | Y |
| LOC102903383 | Vmn1r237 | NW_006501916.1:482501-487500 | NW_006501916.1:485943 | 0.246606935 | 1 | 0.472811018 | | |
| LOC102903517 | Cox7a2 | NW_006501170.1:2075001-2080000 | NW_006501170.1:2074418 | 0.136779404 | 0.99982354 | 0.704367315 | Y | Y |
| LOC102903712 | Hba-x | NW_006501146.1:1102501-1107500 | NW_006501146.1:1104977 | 0.138457921 | 0.99982354 | 0.433139861 | Y | Y |
| LOC102903802 | Rdh16 | NW_006502830.1:15001-20000 | NW_006502830.1:24042 | 0.127354146 | 0.9997059 | 0.490303342 | | |
| LOC102903992 | Fpr-rs3 | NW_006501759.1:877501-882500 | NW_006501759.1:880359 | 0.286153002 | 1 | 0.561689933 | | |
| LOC102903994 | Naip5 | NW_006501779.1:10001-15000 | NW_006501779.1:18259 | 0.117907897 | 0.999352979 | 0.586799476 | | |
| LOC102904031 | F8a | NW_006512344.1:1132254 | NW_006512344.1:4877 | 0.145512691 | 0.99982354 | 0.523172067 | Y | Y |
| LOC102904307 | Vmn2r110 | NW_006501759.1:922501-927500 | NW_006501759.1:928939 | 0.937841497 | 1 | 0.537213053 | | |
| LOC102904434 | Gm4924 | NW_006502780.1:217501-222500 | NW_006502780.1:186187 | 0.153275463 | 0.99988236 | 0.46798113 | | |
| LOC102904812 | Cyp27b1 | NW_006501066.1:727501-732500 | NW_006501066.1:729655 | 0.161118812 | 1 | 0.507478131 | | Y |
| LOC102904934 | Tcaf1 | NW_006502289.1:825001-830000 | NW_006502289.1:827544 | 0.486910488 | 1 | 0.705756657 | Y | Y |
| LOC102905396 | Ces2c | NW_006501344.1:97501-102500 | NW_006501344.1:108576 | 0.660396291 | 1 | 0.638429407 | | |
| LOC102905460 | Zik1 | NW_006502084.1:832501-837500 | NW_006502084.1:834886 | 0.500438315 | 1 | 0.751319914 | Y | Y |
| LOC102905605 | Olfr1272 | NW_006501262.1:2007501-2012500 | NW_006501262.1:2012703 | 0.115903846 | 0.999235339 | 0.450412376 | | |
| LOC102905707 | Cd33 | NW_006501325.1:222501-227500 | NW_006501325.1:221275 | 0.126755826 | 0.99964708 | 0.268817519 | | |

| | | | | | | | | |
|---|---|---|---|---|---|---|---|---|
| LOC102905782 | Rgs22 | NW_006501126.1:25001-30000 | NW_006501126.1:60716 | 0.443112374 | 1 | 0.566450903 | | |
| LOC102905923 | Ces2e | NW_006501344.1:152501-157500 | NW_006501344.1:151752 | 0.353585853 | 1 | 0.459837283 | | Y |
| LOC102906152 | Zfp558 | NW_006501646.1:1312501-1317500 | NW_006501646.1:1302946 | 0.266169814 | 1 | 0.662846248 | Y | Y |
| LOC102906218 | Ctsr | NW_006501203.1:3187501-3192500 | NW_006501203.1:3194090 | 0.200110549 | 1 | 0.463223152 | Y | Y |
| LOC102906541 | Ces2b | NW_006501344.1:235001-240000 | NW_006501344.1:244202 | 1.101791111 | 1 | 0.686060706 | Y | Y |
| LOC102906561 | Zfp950 | NW_006501046.1:7501-12500 | NW_006501046.1:9905 | 0.26683573 | 1 | 0.589729316 | | |
| LOC102906846 | Cyp4a32 | NW_006501336.1:1537501-1542500 | NW_006501336.1:1632360 | 0.191577372 | 1 | 0.722580103 | | |
| LOC102906935 | Mmp1a | NW_006501245.1:2847501-2852500 | NW_006501245.1:2841407 | 1.298571885 | 1 | 0.595715748 | Y | Y |
| LOC102907006 | Vmn1r75 | NW_006502084.1:75001-80000 | NW_006502084.1:79596 | 0.131933676 | 0.99976472 | 0.502897926 | | |
| LOC102907220 | Olfr382 | NW_006502910.1:57501-62500 | NW_006502910.1:60973 | 0.113485295 | 0.999117699 | 0.442855853 | | |
| LOC102907251 | Ces2h | NW_006501344.1:262501-267500 | NW_006501344.1:262819 | 0.112028804 | 0.999058879 | 0.563104414 | | |
| LOC102907276 | Qrfpr | NW_006501046.1:572501-577500 | NW_006501046.1:587750 | 0.370804282 | 1 | 0.354837599 | | |
| LOC102907317 | Cpq | NW_006501126.1:2755001-2760000 | NW_006501126.1:2792806 | 0.812718348 | 1 | 0.420260775 | Y | Y |
| LOC102907513 | Olfr1447 | NW_006501881.1:1012501-1017500 | NW_006501881.1:1016072 | 0.11906025 | 0.999352979 | 0.34021569 | | |
| LOC102907544 | Vmn1r67 | NW_006504762.1:1132254 | NW_006504762.1:2935 | 0.280728479 | 1 | 0.678909671 | | |
| LOC102907590 | D3Ertd254e | NW_006501046.1:607501-612500 | NW_006501046.1:612349 | 0.157605428 | 0.99994118 | 0.76294182 | | |
| LOC102907619 | Olfr780 | NW_006502037.1:320001-325000 | NW_006502037.1:323998 | 0.194834557 | 1 | 0.412109205 | | |
| LOC102907678 | Rps26 | NW_006501362.1:377501-382500 | NW_006501362.1:381994 | 0.177620965 | 1 | 0.431103031 | Y | Y |
| LOC102907711 | Vmn2r107 | NW_006501916.1:140001-145000 | NW_006501916.1:168466 | 0.138800452 | 0.99982354 | 0.655822967 | | |
| LOC102907993 | Olfr24 | NW_006501646.1:1675001-1680000 | NW_006501646.1:1677956 | 0.120613769 | 0.999470619 | 0.555217748 | | |
| LOC102908027 | Ces2c | NW_006503433.1:10001-15000 | NW_006503433.1:15405 | 0.180831157 | 1 | 0.605490316 | | |
| LOC102908268 | Msra | NW_006501437.1:102501-107500 | NW_006501437.1:106593 | 0.443095499 | 1 | 0.66905269 | Y | Y |
| LOC102908477 | Birc3 | NW_006501245.1:3177501-3182500 | NW_006501245.1:3185628 | 0.429281368 | 1 | 0.816118189 | Y | Y |
| LOC102908633 | Vmn2r61 | NW_006501916.1:315001-320000 | NW_006501916.1:281382 | 0.258025805 | 1 | 0.502671926 | | |
| LOC102908952 | Vmn2r120 | NW_006502106.1:242501-247500 | NW_006502106.1:213366 | 0.146642054 | 0.99982354 | 0.499141263 | | |
| LOC102909014 | Ankhd1 | NW_006501530.1:1170001-1175000 | NW_006501530.1:1235474 | 0.635572892 | 1 | 0.286857414 | Y | Y |
| LOC102909057 | Slco1a4 | NW_006501111.1:585001-590000 | NW_006501111.1:575267 | 0.142107744 | 0.99982354 | 0.743671208 | | |
| LOC102909107 | NA | NW_006501325.1:885001-890000 | NW_006501325.1:884298 | 0.138305303 | 0.99982354 | 0.603311369 | | |

| | | | | | | | | |
|---|---|---|---|---|---|---|---|---|
| LOC102909279 | Nat8 | NW_006502671.1:72501-77500 | NW_006502671.1:75363 | 0.113977617 | 0.999117699 | 0.495418941 | | |
| LOC102909369 | Slco1a4 | NW_006501111.1:522501-527500 | NW_006501111.1:539929 | 0.180496722 | 1 | 0.58739124 | | |
| LOC102909407 | Gm4847 | NW_006501228.1:1777501-1782500 | NW_006501228.1:1789646 | 0.240153044 | 1 | 0.486121774 | | |
| LOC102909417 | D230025D16Rik | NW_006501344.1:445001-450000 | NW_006501344.1:453990 | 0.271444247 | 1 | 0.399482959 | Y | Y |
| LOC102909592 | Cyp2b10 | NW_006503307.1:45001-50000 | NW_006503307.1:32847 | 0.134577766 | 0.99982354 | 0.358392585 | | |
| LOC102909690 | Rdh7 | NW_006502313.1:97501-102500 | NW_006502313.1:102370 | 0.480848015 | 1 | 0.594747908 | | |
| LOC102909735 | Akr1c14 | NW_006501386.1:92501-97500 | NW_006501386.1:97818 | 0.682244362 | 1 | 0.568879812 | Y | |
| LOC102909864 | Ubtfl1 | NW_006501646.1:1235001-1240000 | NW_006501646.1:1240624 | 0.483376685 | 1 | 0.712901924 | | |
| LOC102909871 | Olfr1336 | NW_006501712.1:680001-685000 | NW_006501712.1:685382 | 0.149741411 | 0.99982354 | 0.361656312 | | |
| LOC102909979 | Zfp94 | NW_006501692.1:740001-745000 | NW_006501692.1:746967 | 0.199293734 | 1 | 0.643322643 | Y | Y |
| LOC102910012 | Ctnna1 | NW_006501133.1:245001-250000 | NW_006501133.1:240279 | 0.227607199 | 1 | 0.694424769 | Y | Y |
| LOC102910196 | NA | NW_006501065.1:12347501-12352500 | NW_006501065.1:12351693 | 0.188398133 | 1 | 0.369507738 | | |
| LOC102910510 | Ubtfl1 | NW_006501646.1:1252501-1257500 | NW_006501646.1:1255798 | 0.414568293 | 1 | 0.820507219 | | |
| LOC102910758 | Cyp2c66 | NW_006502502.1:42501-47500 | NW_006502502.1:47421 | 0.200549766 | 1 | 0.53343809 | | |
| LOC102910861 | Zfp934 | NW_006503132.1:77501-82500 | NW_006503132.1:80120 | 0.153157377 | 0.99988236 | 0.678386452 | | |
| LOC102910947 | Slc7a12 | NW_006501833.1:220001-225000 | NW_006501833.1:249945 | 0.288142453 | 1 | 0.554782403 | | |
| LOC102910975 | Kdm3b | NW_006501133.1:735001-740000 | NW_006501133.1:758111 | 0.548818327 | 1 | 0.357646034 | Y | Y |
| LOC102911074 | Sult2a6 | NW_006502290.1:302501-307500 | NW_006502290.1:303670 | 0.306307124 | 1 | 0.793532134 | | |
| LOC102911142 | Zbtb26 | NW_006501060.1:1460001-1465000 | NW_006501060.1:1464363 | 0.220346385 | 1 | 0.493045322 | Y | Y |
| LOC102911167 | Zcchc3 | NW_006502109.1:307501-312500 | NW_006502109.1:311715 | 0.22871569 | 1 | 0.772642316 | Y | Y |
| LOC102911278 | Olfr25 | NW_006502317.1:37501-42500 | NW_006502317.1:40533 | 0.206143787 | 1 | 0.420586743 | | |
| LOC102911575 | Olfr25 | NW_006502317.1:67501-72500 | NW_006502317.1:70226 | 0.182802249 | 1 | 0.346713704 | | |
| LOC102911621 | Cbr2 | NW_006501378.1:1355001-1360000 | NW_006501378.1:1357273 | 0.297575125 | 1 | 0.662358701 | | |
| LOC102911850 | Dsc3 | NW_006501038.1:6010001-6015000 | NW_006501038.1:6016917 | 0.117705451 | 0.999352979 | 0.309424182 | Y | |
| LOC102912008 | Zcchc3 | NW_006502109.1:465001-470000 | NW_006502109.1:469616 | 0.349432105 | 1 | 0.788243712 | Y | Y |
| LOC102912016 | Cntnap5b | NW_006502503.1:297501-302500 | NW_006502503.1:91016 | 0.390540416 | 1 | 0.508280793 | | |
| LOC102912036 | Itgb2l | NW_006501195.1:2925001-2930000 | NW_006501195.1:2922798 | 0.232387316 | 1 | 0.391911873 | | |
| LOC102912117 | Olfr906 | NW_006502558.1:25001-30000 | NW_006502558.1:28294 | 0.127320934 | 0.9997059 | 0.713934857 | | |

| | | | | | | | | |
|---|---|---|---|---|---|---|---|---|
| LOC102912198 | BC048403 | NW_006501066.1:5845001-5850000 | NW_006501066.1:5835093 | 0.116634197 | 0.999235339 | 0.526587985 | Y | Y |
| LOC102912308 | Olfr914 | NW_006501638.1:47501-52500 | NW_006501638.1:52537 | 0.428137961 | 1 | 0.514202665 | | |
| LOC102912376 | Kcnk16 | NW_006501274.1:4052501-4057500 | NW_006501274.1:4050138 | 0.150482904 | 0.99982354 | 0.513368667 | Y | |
| LOC102912415 | Zim1 | NW_006501712.1:872501-877500 | NW_006501712.1:876667 | 0.34808835 | 1 | 0.472181291 | Y | Y |
| LOC102912625 | Olfr921 | NW_006501638.1:130001-135000 | NW_006501638.1:132359 | 0.26677093 | 1 | 0.572968912 | | |
| LOC102912747 | Cyp3a13 | NW_006502145.1:130001-135000 | NW_006502145.1:119127 | 0.15813965 | 0.99994118 | 0.370339523 | Y | Y |
| LOC102912792 | Hao2 | NW_006501296.1:117501-122500 | NW_006501296.1:127688 | 0.130449824 | 0.99976472 | 0.689821838 | | |
| LOC102912857 | Zcchc3 | NW_006502109.1:355001-360000 | NW_006502109.1:357733 | 0.200000288 | 1 | 0.658156338 | Y | Y |
| LOC102912934 | Olfr922 | NW_006501638.1:165001-170000 | NW_006501638.1:167630 | 0.238883398 | 1 | 0.770181399 | | |
| LOC102912971 | NA | NW_006502381.1:270001-275000 | NW_006502381.1:272820 | 0.154066578 | 0.99994118 | 0.369002391 | | |
| LOC102913033 | E130311K13Rik | NW_006501060.1:2697501-2702500 | NW_006501060.1:2678413 | 0.390809149 | 1 | 0.560497095 | Y | Y |
| LOC102913167 | Zcchc3 | NW_006502109.1:402501-407500 | NW_006502109.1:405149 | 0.815293976 | 1 | 0.515826925 | Y | Y |
| LOC102913247 | Olfr923 | NW_006501638.1:175001-180000 | NW_006501638.1:177451 | 0.245191946 | 1 | 0.752837881 | | |
| LOC102913558 | H2afx | NW_006501652.1:552501-557500 | NW_006501652.1:554852 | 0.202318731 | 1 | 0.683808169 | | |
| LOC102913583 | NA | NW_006502084.1:670001-675000 | NW_006502084.1:675062 | 0.209406964 | 1 | 0.630121433 | | |
| LOC102913629 | Hsd3b6 | NW_006501296.1:252501-257500 | NW_006501296.1:250698 | 0.327681818 | 1 | 0.732457644 | | |
| LOC102913816 | Nxpe4 | NW_006501163.1:800001-805000 | NW_006501163.1:799308 | 0.552920872 | 1 | 0.737064545 | Y | Y |
| LOC102913869 | Olfr933 | NW_006501638.1:197501-202500 | NW_006501638.1:199993 | 0.187002819 | 1 | 0.825405188 | | |
| LOC102913898 | Vmn1r70 | NW_006502084.1:715001-720000 | NW_006502084.1:718933 | 0.340177717 | 1 | 0.750724365 | | |
| LOC102913983 | Vmn2r1 | NW_006501060.1:2832501-2837500 | NW_006501060.1:2822306 | 0.499475139 | 1 | 0.696445751 | | |
| LOC102914090 | C130079G13Rik | NW_006501814.1:1102501-1107500 | NW_006501814.1:1070476 | 0.806309483 | 1 | 0.525906739 | | |
| LOC102914280 | Vmn2r1 | NW_006501060.1:2892501-2897500 | NW_006501060.1:2892987 | 0.679538038 | 1 | 0.389578453 | | |
| LOC102914398 | C130079G13Rik | NW_006501814.1:1127501-1132500 | NW_006501814.1:1131140 | 0.719723532 | 1 | 0.688627852 | | |
| LOC102914497 | Olfr934 | NW_006501638.1:222501-227500 | NW_006501638.1:225759 | 0.15907956 | 0.99994118 | 0.755646078 | | |
| LOC102914701 | Gm8298 | NW_006501814.1:1227501-1232500 | NW_006501814.1:1255421 | 0.961491632 | 1 | 0.467633898 | | |
| LOC102914718 | NA | NW_006502236.1:290001-295000 | NW_006502236.1:291831 | 0.125914546 | 0.99958826 | 0.488653835 | | |
| LOC102914774 | 4933412E24Rik | NW_006501430.1:7501-12500 | NW_006501430.1:11760 | 0.272123719 | 1 | 0.724227684 | | |
| LOC102914793 | Olfr26 | NW_006501638.1:235001-240000 | NW_006501638.1:239720 | 0.123080493 | 0.999529439 | 0.600293763 | | |

| | | | | | | | | |
|---|---|---|---|---|---|---|---|---|
| LOC102914946 | Olfr888 | NW_006502865.1:47501-52500 | NW_006502865.1:50057 | 0.169795756 | 1 | 0.792240637 | | |
| LOC102915078 | Cyp2c29 | NW_006501320.1:1137501-1142500 | NW_006501320.1:1050697 | 0.123106106 | 0.999529439 | 0.313827586 | | |
| LOC102915109 | Olfr935 | NW_006501638.1:265001-270000 | NW_006501638.1:269298 | 0.268542068 | 1 | 0.698326784 | | |
| LOC102915169 | Olfr985 | NW_006501191.1:60001-65000 | NW_006501191.1:63636 | 0.176875733 | 1 | 0.640384126 | | |
| LOC102915274 | Rgs22 | NW_006501220.1:47501-52500 | NW_006501220.1:40770 | 0.242141977 | 1 | 0.441832691 | | |
| LOC102915326 | Aadacl2 | NW_006501814.1:1312501-1317500 | NW_006501814.1:1332304 | 0.392832165 | 1 | 0.675987565 | | |
| LOC102915382 | Cyb5r4 | NW_006501243.1:1500001-1505000 | NW_006501243.1:1435373 | 0.176635446 | 1 | 0.481374891 | Y | Y |
| LOC102915530 | Nlrp1b | NW_006501749.1:150001-155000 | NW_006501749.1:129142 | 0.299613404 | 1 | 0.28924877 | | |
| LOC102915553 | Cyp3a11 | NW_006502320.1:15001-20000 | NW_006502320.1:32957 | 0.1365994 | 0.99982354 | 0.370855033 | | |
| LOC102915596 | Zfp846 | NW_006501353.1:1085001-1090000 | NW_006501353.1:1088196 | 0.220270116 | 1 | 0.541126198 | Y | Y |
| LOC102915783 | Olfr986 | NW_006501191.1:125001-130000 | NW_006501191.1:129782 | 0.637612143 | 1 | 0.768448296 | | |
| LOC102915842 | Cyp2b10 | NW_006501851.1:582501-587500 | NW_006501851.1:593056 | 0.191965024 | 1 | 0.750244265 | | |
| LOC102915911 | Zfp266 | NW_006501353.1:1172501-1177500 | NW_006501353.1:1178032 | 0.126958381 | 0.99964708 | 0.762924178 | Y | Y |
| LOC102916061 | Mep1a | NW_006501862.1:30001-35000 | NW_006501862.1:40954 | 0.20231701 | 1 | 0.384941616 | | |
| LOC102916097 | Zfp809 | NW_006501211.1:2095001-2100000 | NW_006501211.1:2102994 | 0.18284806 | 1 | 0.721334429 | | |
| LOC102916219 | Gm43638 | NW_006501246.1:165001-170000 | NW_006501246.1:161829 | 0.236579537 | 1 | 0.545370463 | | |
| LOC102916229 | Zfp26 | NW_006501353.1:1285001-1290000 | NW_006501353.1:1284273 | 0.574924064 | 1 | 0.802271958 | Y | Y |
| LOC102916294 | Glb1l2 | NW_006501112.1:4845001-4850000 | NW_006501112.1:4873479 | 0.268384672 | 1 | 0.449729307 | Y | Y |
| LOC102916391 | Vmn1r68 | NW_006502084.1:935001-940000 | NW_006502084.1:936840 | 0.548997399 | 1 | 0.566540293 | | |
| LOC102916413 | Zfp810 | NW_006501211.1:2147501-2152500 | NW_006501211.1:2152586 | 0.247478306 | 1 | 0.627046358 | Y | Y |
| LOC102916444 | Imp4 | NW_006501040.1:17417501-17422500 | NW_006501040.1:17420335 | 0.111955999 | 0.999058879 | 0.4427679 | Y | Y |
| LOC102916678 | Olfr959 | NW_006501638.1:457501-462500 | NW_006501638.1:461442 | 0.307330325 | 1 | 0.79595906 | | |
| LOC102916719 | Olfr91 | NW_006503095.1:72501-77500 | NW_006503095.1:75425 | 0.158144675 | 0.99994118 | 0.452020965 | | |
| LOC102916932 | Gm14685 | NW_006502593.1:80001-85000 | NW_006502593.1:83938 | 0.383817453 | 1 | 0.596807705 | | |
| LOC102916970 | Ugt1a6a | NW_006501416.1:35001-40000 | NW_006501416.1:38741 | 0.193628139 | 1 | 0.549847438 | | |
| LOC102917074 | Cetn4 | NW_006501581.1:1257501-1262500 | NW_006501581.1:1258711 | 0.140549449 | 0.99982354 | 0.28924877 | | |
| LOC102917265 | Olfr871 | NW_006501353.1:1395001-1400000 | NW_006501353.1:1399695 | 0.458534424 | 1 | 0.661833393 | | |
| LOC102917596 | Olfr963 | NW_006501638.1:537501-542500 | NW_006501638.1:540033 | 0.193322667 | 1 | 0.451744485 | | |

| | | | | | | | | |
|---|---|---|---|---|---|---|---|---|
| LOC102917650 | Jhy | NW_006501191.1:867501-872500 | NW_006501191.1:887098 | 0.278310883 | 1 | 0.600029171 | | |
| LOC102917876 | Olfr873 | NW_006501353.1:1457501-1462500 | NW_006501353.1:1461549 | 0.195237732 | 1 | 0.677216929 | | |
| LOC102917904 | Olfr149 | NW_006501638.1:580001-585000 | NW_006501638.1:582141 | 0.147206875 | 0.99982354 | 0.542804897 | | |
| LOC102917993 | Vmn2r77 | NW_006501516.1:232501-237500 | NW_006501516.1:251128 | 0.135227655 | 0.99982354 | 0.363467468 | | |
| LOC102918038 | Gm1110 | NW_006501112.1:4700001-4705000 | NW_006501112.1:4729164 | 0.680428997 | 1 | 0.606182173 | | |
| LOC102918218 | Olfr965 | NW_006501638.1:637501-642500 | NW_006501638.1:642589 | 0.152994225 | 0.99988236 | 0.432502475 | | |
| LOC102918219 | Dhrs7 | NW_006501649.1:47501-52500 | NW_006501649.1:51161 | 0.132654413 | 0.99976472 | 0.306566694 | Y | Y |
| LOC102918499 | Olfr872 | NW_006501353.1:1550001-1555000 | NW_006501353.1:1553675 | 0.199104498 | 1 | 0.840886019 | | |
| LOC102918677 | Vmn1r67 | NW_006504337.1:22501-27500 | NW_006504337.1:25926 | 0.130983446 | 0.99976472 | 0.75414446 | | |
| LOC102918796 | Ugt2b5 | NW_006501246.1:370001-375000 | NW_006501246.1:373218 | 0.222953905 | 1 | 0.716275109 | | |
| LOC102918868 | Zfp808 | NW_006502722.1:110001-115000 | NW_006502722.1:112089 | 0.668923706 | 1 | 0.369355889 | Y | Y |
| LOC102919113 | Olfr862 | NW_006501353.1:1632501-1637500 | NW_006501353.1:1635567 | 0.127149005 | 0.99964708 | 0.704717834 | | |
| LOC102919204 | Bysl | NW_006501434.1:7501-12500 | NW_006501434.1:10989 | 0.123751883 | 0.999529439 | 0.391690469 | Y | Y |
| LOC102919257 | Adgrb3 | NW_006501113.1:867501-872500 | NW_006501113.1:899212 | 0.327491643 | 1 | 0.606361783 | | |
| LOC102919359 | Ces2c | NW_006502043.1:17501-22500 | NW_006502043.1:26137 | 0.121882848 | 0.999470619 | 0.678491967 | | |
| LOC102919750 | Olfr974 | NW_006501638.1:732501-737500 | NW_006501638.1:736112 | 0.267563641 | 1 | 0.833602528 | | |
| LOC102920059 | Olfr975 | NW_006501638.1:745001-750000 | NW_006501638.1:747870 | 0.186269972 | 1 | 0.509359422 | | |
| LOC102920188 | Ces2a | NW_006502043.1:80001-85000 | NW_006502043.1:88560 | 0.369289791 | 1 | 0.638658814 | | |
| LOC102920196 | Olfr1214 | NW_006502232.1:280001-285000 | NW_006502232.1:284399 | 0.130646497 | 0.99976472 | 0.453490058 | | |
| LOC102920371 | Olfr976 | NW_006501638.1:750001-755000 | NW_006501638.1:753416 | 0.136564519 | 0.99982354 | 0.521839596 | | |
| LOC102920506 | Olfr1214 | NW_006502232.1:337501-342500 | NW_006502232.1:342765 | 0.326877341 | 1 | 0.640454758 | | |
| LOC102920548 | Slc22a22 | NW_006501212.1:70001-75000 | NW_006501212.1:92607 | 0.463578881 | 1 | 0.757599721 | | |
| LOC102920571 | Hyal6 | NW_006501405.1:1825001-1830000 | NW_006501405.1:1829648 | 0.367462424 | 1 | 0.660638228 | | |
| LOC102920647 | Olfr853 | NW_006501353.1:1780001-1785000 | NW_006501353.1:1780020 | 0.312698877 | 1 | 0.732851822 | | |
| LOC102920693 | Mageb16 | NW_006501947.1:455001-460000 | NW_006501947.1:459403 | 0.555036423 | 1 | 0.542282533 | Y | Y |
| LOC102920805 | Olfr160 | NW_006502270.1:5001-10000 | NW_006502270.1:8881 | 0.126778625 | 0.99964708 | 0.665895683 | | |
| LOC102920815 | NA | NW_006501139.1:2952501-2957500 | NW_006501139.1:2972506 | 0.211582824 | 1 | 0.456800874 | | |
| LOC102920988 | Olfr978 | NW_006501638.1:792501-797500 | NW_006501638.1:796873 | 0.159620615 | 0.99994118 | 0.660358649 | | |

| | | | | | | | | |
|---|---|---|---|---|---|---|---|---|
| LOC102921127 | Olfr160 | NW_006502270.1:30001-35000 | NW_006502270.1:33973 | 0.28623274 | 1 | 0.699586736 | | |
| LOC102921168 | C87436 | NW_006501269.1:1505001-1510000 | NW_006501269.1:1510215 | 0.135637529 | 0.99982354 | 0.408804023 | Y | Y |
| LOC102921183 | Spam1 | NW_006501405.1:1955001-1960000 | NW_006501405.1:1954042 | 0.112915757 | 0.999058879 | 0.709728611 | | |
| LOC102921292 | Olfr979 | NW_006501638.1:800001-805000 | NW_006501638.1:805070 | 0.207038232 | 1 | 0.620904465 | | |
| LOC102921874 | Olfr843 | NW_006501353.1:1972501-1977500 | NW_006501353.1:1974628 | 0.275650341 | 1 | 0.543297757 | | |
| LOC102921878 | BC048546 | NW_006501387.1:212501-217500 | NW_006501387.1:212681 | 0.123403579 | 0.999529439 | 0.621483954 | | |
| LOC102922071 | Olfr982 | NW_006501191.1:5001-10000 | NW_006501191.1:7035 | 0.144802821 | 0.99982354 | 0.506782978 | | |
| LOC102922475 | Gm4924 | NW_006501210.1:425001-430000 | NW_006501210.1:430058 | 0.117869025 | 0.999352979 | 0.538057249 | | |
| LOC102922490 | Hsd17b6 | NW_006501362.1:1110001-1115000 | NW_006501362.1:1114039 | 0.469806576 | 1 | 0.711435713 | | |
| LOC102922517 | Olfr919 | NW_006501638.1:65001-70000 | NW_006501638.1:68548 | 0.335460013 | 1 | 0.516333603 | | |
| LOC102922543 | NA | NW_006501958.1:1037501-1042500 | NW_006501958.1:1039823 | 0.226173099 | 1 | 0.641364248 | | |
| LOC102922568 | Olfr147 | NW_006503562.1:7501-12500 | NW_006503562.1:13084 | 0.843184914 | 1 | 0.464851291 | | |
| LOC102922906 | Aldh1a1 | NW_006501292.1:3420001-3425000 | NW_006501292.1:3435199 | 0.122752049 | 0.999470619 | 0.557704095 | Y | Y |
| LOC102922983 | Olfr878 | NW_006502270.1:197501-202500 | NW_006502270.1:199700 | 0.289544753 | 1 | 0.742772305 | | |
| LOC102923286 | NA | NW_006501970.1:485001-490000 | NW_006501970.1:469846 | 0.285106515 | 1 | 0.461609676 | | |
| LOC102923362 | Lyz2 | NW_006501066.1:10220001-10225000 | NW_006501066.1:10223016 | 0.177136603 | 1 | 0.644756396 | Y | Y |
| LOC102923471 | Usp51 | NW_006501855.1:170001-175000 | NW_006501855.1:174610 | 0.939068939 | 1 | 0.666523343 | | |
| LOC102923525 | Gpr165 | NW_006501190.1:5030001-5035000 | NW_006501190.1:5031532 | 1.012425206 | 1 | 0.495701316 | | |
| LOC102923679 | Slc22a19 | NW_006501905.1:80001-85000 | NW_006501905.1:122000 | 0.230000695 | 1 | 0.353412534 | | |
| LOC102923707 | Vmn1r70 | NW_006501169.1:2050001-2055000 | NW_006501169.1:2052887 | 0.195095714 | 1 | 0.476039049 | | |
| LOC102924224 | Olfr145 | NW_006502270.1:177501-182500 | NW_006502270.1:180811 | 0.169890961 | 1 | 0.627970492 | | |
| LOC102924536 | Vmn2r56 | NW_006502295.1:435001-440000 | NW_006502295.1:427352 | 0.292801593 | 1 | 0.663029659 | | |
| LOC102924668 | Zfp560 | NW_006501353.1:1357501-1362500 | NW_006501353.1:1359384 | 0.164577089 | 1 | 0.44573088 | Y | Y |
| LOC102924700 | Olfr959 | NW_006501638.1:247501-252500 | NW_006501638.1:251835 | 0.153539291 | 0.99988236 | 0.509550442 | | |
| LOC102924798 | NA | NW_006501520.1:165001-170000 | NW_006501520.1:169910 | 0.264450784 | 1 | 0.65803976 | | |
| LOC102924848 | Vmn2r54 | NW_006502295.1:450001-455000 | NW_006502295.1:452865 | 0.140124941 | 0.99982354 | 0.697073752 | | |
| LOC102924917 | 1700122O11Rik | NW_006501738.1:365001-370000 | NW_006501738.1:368170 | 0.185308185 | 1 | 0.631741816 | | |
| LOC102924933 | Gm10310 | NW_006502126.1:377501-382500 | NW_006502126.1:380032 | 0.138226425 | 0.99982354 | 0.31061649 | | |

| | | | | | | | | |
|---|---|---|---|---|---|---|---|---|
| LOC102925027 | Vmn2r58 | NW_006501797.1:292501-297500 | NW_006501797.1:295482 | 0.180736064 | 1 | 0.414604151 | | |
| LOC102925569 | Ces2g | NW_006501139.1:3310001-3315000 | NW_006501139.1:3315386 | 0.23697505 | 1 | 0.605551075 | | |
| LOC102925729 | Zfp870 | NW_006501554.1:885001-890000 | NW_006501554.1:888357 | 0.282665997 | 1 | 0.533509426 | Y | Y |
| LOC102925951 | Olfr955 | NW_006501638.1:387501-392500 | NW_006501638.1:390926 | 0.17877953 | 1 | 0.735694941 | | |
| LOC102926234 | Olfr851 | NW_006501353.1:1810001-1815000 | NW_006501353.1:1812756 | 0.301882633 | 1 | 0.506503508 | | |
| LOC102926342 | Vmn2r60 | NW_006501520.1:1580001-1585000 | NW_006501520.1:1584967 | 0.139783191 | 0.99982354 | 0.61255975 | | |
| LOC102926469 | Zfp990 | NW_006501759.1:217501-222500 | NW_006501759.1:219829 | 0.187583065 | 1 | 0.3684571 | | |
| LOC102926599 | Ces2a | NW_006501139.1:3410001-3415000 | NW_006501139.1:3413285 | 0.235042744 | 1 | 0.418236947 | | |
| LOC102926603 | Vmn1r70 | NW_006503051.1:25001-30000 | NW_006503051.1:30021 | 0.38929334 | 1 | 0.701485984 | | |
| LOC102926642 | 1810026J23Rik | NW_006501353.1:5001-10000 | NW_006501353.1:8427 | 0.421794648 | 1 | 0.739450565 | | |
| LOC102926817 | Runx2 | NW_006502977.1:52501-57500 | NW_006502977.1:56706 | 0.133664204 | 0.99982354 | 0.557543418 | Y | Y |
| LOC102926827 | NA | NW_006501169.1:2400001-2405000 | NW_006501169.1:2402068 | 0.11349674 | 0.999117699 | 0.640432218 | | |
| LOC102926846 | Olfr869 | NW_006501353.1:1855001-1860000 | NW_006501353.1:1854445 | 0.434424807 | 1 | 0.719425485 | | |
| LOC102926876 | AW551984 | NW_006501638.1:482501-487500 | NW_006501638.1:471827 | 0.900472978 | 1 | 0.764602041 | Y | Y |
| LOC102926914 | Gm4924 | NW_006503596.1:15001-20000 | NW_006503596.1:21640 | 0.171710965 | 1 | 0.446775217 | | |
| LOC102926916 | Kirrel3 | NW_006501164.1:2692501-2697500 | NW_006501164.1:2703288 | 0.222709629 | 1 | 0.473815551 | Y | |
| LOC102926946 | Gm13124 | NW_006501437.1:167501-172500 | NW_006501437.1:163566 | 0.220304769 | 1 | 0.533756909 | | |
| LOC102927177 | Vwa5a | NW_006501638.1:597501-602500 | NW_006501638.1:614454 | 0.616220093 | 1 | 0.753613986 | Y | Y |
| LOC102927215 | Vmn1r70 | NW_006503051.1:97501-102500 | NW_006503051.1:100959 | 0.158157683 | 0.99994118 | 0.453316452 | | |
| LOC102927216 | Olfr243 | NW_006501149.1:257501-262500 | NW_006501149.1:261504 | 0.206157878 | 1 | 0.646335454 | | |
| LOC102927249 | Gm13177 | NW_006501437.1:232501-237500 | NW_006501437.1:232002 | 0.581042151 | 1 | 0.719143622 | | |
| LOC102927285 | Zfp53 | NW_006501856.1:462501-467500 | NW_006501856.1:461857 | 0.116958361 | 0.999235339 | 0.438131809 | | |
| LOC102927424 | Gsdmc | NW_006503966.1:2501-7500 | NW_006503966.1:10975 | 0.210720974 | 1 | 0.782684169 | | |
| LOC102927726 | Olfr902 | NW_006503971.1:22501-27500 | NW_006503971.1:26915 | 0.261682961 | 1 | 0.776025459 | | |
| LOC102927860 | Gm438 | NW_006501437.1:372501-377500 | NW_006501437.1:372897 | 0.114631943 | 0.999235339 | 0.6779687 | | |
| LOC102928024 | Olfr917 | NW_006503971.1:30001-35000 | NW_006503971.1:33824 | 0.222081518 | 1 | 0.61944891 | | |
| LOC102928062 | Acrv1 | NW_006501547.1:197501-202500 | NW_006501547.1:199625 | 0.187605138 | 1 | 0.466252554 | | |
| LOC102928123 | Olfr68 | NW_006501149.1:292501-297500 | NW_006501149.1:294211 | 0.156891987 | 0.99994118 | 0.75675273 | | |

| | | | | | | | | |
|---|---|---|---|---|---|---|---|---|
| LOC102928318 | Olfr918 | NW_006503971.1:7501-12500 | NW_006503971.1:12127 | 0.111991417 | 0.999058879 | 0.559902541 | | |
| LOC102928385 | Tas2r124 | NW_006501958.1:972501-977500 | NW_006501958.1:977911 | 0.14255272 | 0.99982354 | 0.540198165 | | |
| LOC102928410 | Olfr67 | NW_006501149.1:297501-302500 | NW_006501149.1:301121 | 0.185971526 | 1 | 0.751841235 | | |
| LOC102928660 | Slc35e2 | NW_006501627.1:712501-717500 | NW_006501627.1:715277 | 0.223537981 | 1 | 0.368349103 | Y | Y |
| LOC102928670 | 4930578C19Rik | NW_006501734.1:67501-72500 | NW_006501734.1:126598 | 0.194486172 | 1 | 0.744615864 | | |
| LOC102928879 | Vmn1r224 | NW_006501759.1:452501-457500 | NW_006501759.1:456590 | 0.160066757 | 1 | 0.532124603 | | |
| LOC102928913 | NA | NW_006501139.1:2970001-2975000 | NW_006501139.1:2972513 | 0.131451375 | 0.99976472 | 0.353039949 | | |
| LOC102928917 | Hba-a2 | NW_006501146.1:1092501-1097500 | NW_006501146.1:1098464 | 0.278145765 | 1 | 0.28924877 | | |
| LOC107399310 | Zfp35 | NW_006501173.1:3700001-3705000 | NW_006501173.1:3700648 | 0.133743084 | 0.99982354 | 0.625638061 | | |
| LOC107400260 | NA | NW_006502671.1:72501-77500 | NW_006502671.1:75363 | 0.113977617 | 0.999117699 | 0.614436876 | | |
| LOC107400365 | Vmn1r185 | NW_006503051.1:167501-172500 | NW_006503051.1:169717 | 0.36199702 | 1 | 0.636641153 | | |
| LOC107400753 | NA | NW_006501066.1:9787501-9792500 | NW_006501066.1:9768462 | 0.125084793 | 0.999529439 | 0.649996405 | | |
| LOC107401125 | NA | NW_006501107.1:4217501-4222500 | NW_006501107.1:4219927 | 0.124020396 | 0.999529439 | 0.445200955 | | |
| LOC107401168 | Adgrb3 | NW_006501113.1:1670001-1675000 | NW_006501113.1:1674647 | 0.297645718 | 1 | 0.723469875 | | |
| LOC107401289 | NA | NW_006501125.1:4847501-4852500 | NW_006501125.1:4823936 | 0.151883025 | 0.99988236 | 0.551285775 | | |
| LOC107401295 | NA | NW_006501126.1:707501-712500 | NW_006501126.1:713929 | 0.238039561 | 1 | 0.432772484 | | |
| LOC107401301 | NA | NW_006501126.1:2312501-2317500 | NW_006501126.1:2313353 | 0.778159182 | 1 | 0.781006023 | | |
| LOC107401388 | NA | NW_006501139.1:2777501-2782500 | NW_006501139.1:2780021 | 0.187836258 | 1 | 0.381491952 | | |
| LOC107401777 | NA | NW_006501046.1:2012501-2017500 | NW_006501046.1:1972842 | 0.245905307 | 1 | 0.710319819 | | |
| LOC107401828 | NA | NW_006501046.1:7092501-7097500 | NW_006501046.1:7086776 | 0.190352881 | 1 | 0.391549703 | | |
| LOC107401851 | NA | NW_006501046.1:9452501-9457500 | NW_006501046.1:9457631 | 0.651171869 | 1 | 0.715364129 | | |
| LOC107402053 | NA | NW_006501245.1:3180001-3185000 | NW_006501245.1:3180867 | 0.306593285 | 1 | 0.414231442 | | |
| LOC107402055 | NA | NW_006501245.1:4172501-4177500 | NW_006501245.1:4172445 | 0.474961847 | 1 | 0.806016703 | | |
| LOC107402495 | NA | NW_006501362.1:940001-945000 | NW_006501362.1:946017 | 0.298522511 | 1 | 0.587586492 | | |
| LOC107402497 | NA | NW_006501362.1:1110001-1115000 | NW_006501362.1:1114039 | 0.469806576 | 1 | 0.711435713 | | |
| LOC107402634 | NA | NW_006501405.1:1657501-1662500 | NW_006501405.1:1664971 | 0.124667131 | 0.999529439 | 0.674067693 | | |
| LOC107402972 | NA | NW_006501054.1:6080001-6085000 | NW_006501054.1:6085613 | 0.213980428 | 1 | 0.649939562 | | |
| LOC107402977 | NA | NW_006501550.1:2395001-2400000 | NW_006501550.1:2380568 | 0.121391012 | 0.999470619 | 0.59291121 | | |

| | | | | | | | | |
|---|---|---|---|---|---|---|---|---|
| LOC107403033 | NA | NW_006501574.1:3282501-3287500 | NW_006501574.1:3277488 | 0.241434756 | 1 | 0.273443517 | | |
| Lrfn2 | Lrfn2 | NW_006502066.1:22501-27500 | NW_006502066.1:24601 | 0.177614621 | 1 | 0.507910446 | | |
| Lrig3 | Lrig3 | NW_006501066.1:1690001-1695000 | NW_006501066.1:1705869 | 0.226626267 | 1 | 0.501185869 | Y | Y |
| Lrp1 | Lrp1 | NW_006501066.1:220001-225000 | NW_006501066.1:222959 | 0.193062933 | 1 | 0.43059591 | Y | Y |
| Lrrc31 | NA | NW_006501046.1:6802501-6807500 | NW_006501046.1:6787523 | 0.373412561 | 1 | 0.588889385 | | |
| Lrrc34 | Lrrc34 | NW_006501046.1:6755001-6760000 | NW_006501046.1:6728325 | 0.427519564 | 1 | 0.715364129 | | |
| Lrrc7 | Lrrc7 | NW_006501088.1:877501-882500 | NW_006501088.1:946510 | 0.194880493 | 1 | 0.396321264 | | |
| Lrrcc1 | Lrrcc1 | NW_006501833.1:335001-340000 | NW_006501833.1:345481 | 0.149529404 | 0.99982354 | 0.763132752 | Y | Y |
| Lrrd1 | Lrrd1 | NW_006501550.1:2395001-2400000 | NW_006501550.1:2380568 | 0.121391012 | 0.999470619 | 0.59291121 | | |
| Lrrtm2 | Lrrtm2 | NW_006501133.1:277501-282500 | NW_006501133.1:282699 | 0.114489765 | 0.999176519 | 0.427345923 | | |
| Lypd8 | Lypd8 | NW_006501276.1:4377501-4382500 | NW_006501276.1:4381050 | 0.174539186 | 1 | 0.427249687 | | |
| Lztfl1 | Lztfl1 | NW_006501517.1:512501-517500 | NW_006501517.1:535487 | 0.286768501 | 1 | 0.350342379 | Y | Y |
| Mab21l1 | Mab21l1 | NW_006501125.1:2550001-2555000 | NW_006501125.1:2555065 | 0.387391558 | 1 | 0.714120219 | | |
| Maml2 | Maml2 | NW_006501653.1:1090001-1095000 | NW_006501653.1:1093328 | 0.460370069 | 1 | 0.842691616 | Y | Y |
| Maml3 | Maml3 | NW_006501125.1:6772501-6777500 | NW_006501125.1:6341432 | 0.283110571 | 1 | 0.539308495 | Y | Y |
| Map3k2 | Map3k2 | NW_006501133.1:3265001-3270000 | NW_006501133.1:3260890 | 0.21625614 | 1 | 0.379902885 | Y | Y |
| March6 | #N/A | NW_006501168.1:2567501-2572500 | NW_006501168.1:2568743 | 0.149842971 | 0.99982354 | 0.774964552 | | |
| March9 | #N/A | NW_006501066.1:727501-732500 | NW_006501066.1:727381 | 0.161118812 | 1 | 0.697435506 | | |
| Mars | Mars | NW_006501066.1:465001-470000 | NW_006501066.1:471348 | 0.11475716 | 0.999235339 | 0.291284696 | | |
| Mast4 | Mast4 | NW_006501218.1:4277501-4282500 | NW_006501218.1:4262041 | 0.177625573 | 1 | 0.821347047 | Y | Y |
| Matn2 | Matn2 | NW_006501126.1:1550001-1555000 | NW_006501126.1:1629803 | 0.186638816 | 1 | 0.33450033 | Y | Y |
| Mbd6 | Mbd6 | NW_006501066.1:482501-487500 | NW_006501066.1:485459 | 0.163531252 | 1 | 0.689450438 | Y | Y |
| Mbnl1 | Mbnl1 | NW_006502570.1:477501-482500 | NW_006502570.1:481034 | 0.61937453 | 1 | 0.696806226 | Y | Y |
| Mboat2 | Mboat2 | NW_006501185.1:237501-242500 | NW_006501185.1:244793 | 0.611922481 | 1 | 0.660734253 | Y | Y |
| Mc3r | Mc3r | NW_006501290.1:1047501-1052500 | NW_006501290.1:1051282 | 0.334363545 | 1 | 0.70885305 | | |
| Mcam | Mcam | NW_006501652.1:355001-360000 | NW_006501652.1:357778 | 0.510852519 | 1 | 0.774863906 | Y | Y |
| Mdm1 | Mdm1 | NW_006501066.1:9375001-9380000 | NW_006501066.1:9378450 | 0.310804298 | 1 | 0.508756349 | Y | Y |
| Mea1 | Mea1 | NW_006501519.1:377501-382500 | NW_006501519.1:381719 | 0.121315575 | 0.999470619 | 0.454554757 | Y | Y |

| | | | | | | | | |
|---|---|---|---|---|---|---|---|---|
| Mecom | Mecom | NW_006501046.1:6060001-6065000 | NW_006501046.1:6062504 | 0.242495213 | 1 | 0.73509225 | Y | Y |
| Med12l | Med12l | NW_006501814.1:835001-840000 | NW_006501814.1:872041 | 1.031543477 | 1 | 0.661037733 | Y | Y |
| Med20 | Gm20517 | NW_006502218.1:155001-160000 | NW_006502218.1:149141 | 0.201155032 | 1 | 0.388771409 | | |
| Mef2c | Mef2c | NW_006501392.1:1362501-1367500 | NW_006501392.1:1363288 | 0.172945683 | 1 | 0.772627812 | Y | Y |
| Megf8 | Megf8 | NW_006501390.1:1232501-1237500 | NW_006501390.1:1207516 | 0.111889003 | 0.999058879 | 0.451741288 | Y | Y |
| Mfn1 | Mfn1 | NW_006501046.1:4170001-4175000 | NW_006501046.1:4151447 | 0.616528013 | 1 | 0.457058346 | Y | Y |
| Mfrp | Mfrp | NW_006501652.1:315001-320000 | NW_006501652.1:318423 | 0.339737972 | 1 | 0.505239949 | | |
| Mfsd8 | Mfsd8 | NW_006501366.1:1717501-1722500 | NW_006501366.1:1712701 | 0.625552341 | 1 | 0.704231783 | Y | Y |
| Mgarp | Mgarp | NW_006501125.1:7072501-7077500 | NW_006501125.1:7077545 | 0.445191305 | 1 | 0.387093256 | | |
| Mib1 | Mib1 | NW_006501454.1:1690001-1695000 | NW_006501454.1:1612583 | 0.648057129 | 1 | 0.721905626 | Y | Y |
| Mkrn3 | Mkrn3 | NW_006502754.1:182501-187500 | NW_006502754.1:187202 | 0.510283357 | 1 | 0.612273971 | Y | Y |
| Mme | Mme | NW_006501060.1:2105001-2110000 | NW_006501060.1:2148909 | 0.972024362 | 1 | 0.692966081 | Y | Y |
| Mmp10 | Mmp10 | NW_006501245.1:2857501-2862500 | NW_006501245.1:2855981 | 0.570660507 | 1 | 0.837640448 | Y | Y |
| Mmp12 | Mmp12 | NW_006501245.1:2792501-2797500 | NW_006501245.1:2784285 | 0.247088546 | 1 | 0.80802549 | Y | Y |
| Mmp3 | Mmp3 | NW_006501245.1:2820001-2825000 | NW_006501245.1:2821241 | 0.171015674 | 1 | 0.341192334 | | Y |
| Mmp8 | Mmp8 | NW_006501245.1:2885001-2890000 | NW_006501245.1:2897584 | 0.413698943 | 1 | 0.690416551 | | |
| Mon2 | Mon2 | NW_006501066.1:4617501-4622500 | NW_006501066.1:4639634 | 0.315041041 | 1 | 0.678907554 | Y | Y |
| Morc3 | Morc3 | NW_006501195.1:225001-230000 | NW_006501195.1:242376 | 0.239599611 | 1 | 0.329939784 | Y | Y |
| Mrpl2 | Mrpl2 | NW_006501519.1:342501-347500 | NW_006501519.1:344415 | 0.232869632 | 1 | 0.580674142 | Y | Y |
| Mrpl47 | Mrpl47 | NW_006501046.1:3960001-3965000 | NW_006501046.1:3972127 | 0.276859886 | 1 | 0.604157429 | Y | Y |
| Msh6 | Msh6 | NW_006501333.1:417501-422500 | NW_006501333.1:426319 | 0.116658812 | 0.999235339 | 0.582247572 | Y | Y |
| Msrb3 | Msrb3 | NW_006501066.1:6845001-6850000 | NW_006501066.1:6848860 | 0.205958443 | 1 | 0.768373003 | Y | Y |
| Mss51 | Mss51 | NW_006501274.1:3825001-3830000 | NW_006501274.1:3813323 | 0.312392589 | 1 | 0.558393634 | Y | Y |
| Mtbp | Mtbp | NW_006501212.1:1602501-1607500 | NW_006501212.1:1573943 | 0.30418169 | 1 | 0.321611836 | Y | Y |
| Mtdh | Mtdh | NW_006501126.1:1880001-1885000 | NW_006501126.1:1885074 | 0.450091185 | 1 | 0.727100588 | Y | Y |
| Mtmr2 | Mtmr2 | NW_006501653.1:1265001-1270000 | NW_006501653.1:1292374 | 0.265040239 | 1 | 0.805533557 | Y | Y |
| Mto1 | Mto1 | NW_006501895.1:52501-57500 | NW_006501895.1:63252 | 0.124726482 | 0.999529439 | 0.419315772 | Y | Y |
| Mtss1 | Mtss1 | NW_006501508.1:1082501-1087500 | NW_006501508.1:1085061 | 0.276067222 | 1 | 0.481047524 | Y | Y |

| | | | | | | | | |
|---|---|---|---|---|---|---|---|---|
| Myc | Myc | NW_006501430.1:1797501-1802500 | NW_006501430.1:1800574 | 0.223925045 | 1 | 0.364101535 | Y | Y |
| Mycbp2 | Mycbp2 | NW_006501134.1:2235001-2240000 | NW_006501134.1:2400028 | 0.113963382 | 0.999117699 | 0.346904611 | Y | Y |
| Mycn | Mycn | NW_006501114.1:1107501-1112500 | NW_006501114.1:1111736 | 0.118622373 | 0.999352979 | 0.552834245 | Y | |
| Mynn | Mynn | NW_006501046.1:6715001-6720000 | NW_006501046.1:6709618 | 0.551947044 | 1 | 0.682654804 | Y | Y |
| Myo7b | Myo7b | NW_006501133.1:3542501-3547500 | NW_006501133.1:3501616 | 0.1698972 | 1 | 0.405605866 | Y | Y |
| Myoz1 | Myoz1 | NW_006501274.1:3652501-3657500 | NW_006501274.1:3657411 | 0.135219139 | 0.99982354 | 0.337742722 | | |
| N4bp2l2 | N4bp2l2 | NW_006501358.1:1590001-1595000 | NW_006501358.1:1590479 | 0.14863871 | 0.99982354 | 0.493150342 | Y | Y |
| Naa15 | Naa15 | NW_006501125.1:6992501-6997500 | NW_006501125.1:6982139 | 0.853985021 | 1 | 0.694450259 | Y | Y |
| Naaladl2 | Naaladl2 | NW_006501046.1:12240001-12245000 | NW_006501046.1:12123208 | 0.531432067 | 1 | 0.658171628 | | Y |
| Nab2 | Nab2 | NW_006501066.1:82501-87500 | NW_006501066.1:86991 | 0.142335176 | 0.99982354 | 0.797535785 | Y | Y |
| Naca | Naca | NW_006501362.1:1030001-1035000 | NW_006501362.1:1041755 | 0.122099336 | 0.999470619 | 0.561547271 | Y | Y |
| Nbea | Nbea | NW_006501125.1:2675001-2680000 | NW_006501125.1:2333157 | 0.830039427 | 1 | 0.649027623 | Y | Y |
| Nbeal1 | Nbeal1 | NW_006501089.1:6885001-6890000 | NW_006501089.1:6882682 | 0.148827642 | 0.99982354 | 0.296224391 | Y | Y |
| Ncald | Ncald | NW_006501220.1:1475001-1480000 | NW_006501220.1:1474908 | 0.135833193 | 0.99982354 | 0.568259686 | Y | Y |
| Ncam1 | Ncam1 | NW_006501057.1:4725001-4730000 | NW_006501057.1:4733662 | 1.057721127 | 1 | 0.781067186 | Y | Y |
| Ncapd3 | Ncapd3 | NW_006501112.1:4580001-4585000 | NW_006501112.1:4536143 | 0.915802727 | 1 | 0.820716461 | Y | Y |
| Nceh1 | Nceh1 | NW_006501046.1:9562501-9567500 | NW_006501046.1:9542319 | 0.28576527 | 1 | 0.560161209 | Y | Y |
| Ndn | Ndn | NW_006502754.1:255001-260000 | NW_006502754.1:259176 | 0.433771912 | 1 | 0.650515731 | Y | Y |
| Ndufa4l2 | Ndufa4l2 | NW_006501066.1:245001-250000 | NW_006501066.1:244257 | 0.327986966 | 1 | 0.533727283 | Y | Y |
| Ndufa5 | NA | NW_006501405.1:1600001-1605000 | NW_006501405.1:1618460 | 0.131986369 | 0.99976472 | 0.636122267 | Y | Y |
| Ndufb5 | Ndufb5 | NW_006501046.1:3935001-3940000 | NW_006501046.1:3938532 | 0.621364987 | 1 | 0.322141269 | Y | Y |
| Ndufc1 | Ndufc1 | NW_006501125.1:7060001-7065000 | NW_006501125.1:7059777 | 0.410102011 | 1 | 0.501647893 | Y | Y |
| Nectin1 | Nectin1 | NW_006501191.1:3965001-3970000 | NW_006501191.1:3968806 | 0.347767101 | 1 | 0.545405231 | Y | Y |
| Nelfcd | Nelfcd | NW_006501107.1:1295001-1300000 | NW_006501107.1:1306667 | 0.619333628 | 1 | 0.417207041 | Y | Y |
| Nemp1 | Nemp1 | NW_006501066.1:40001-45000 | NW_006501066.1:38647 | 0.164630958 | 1 | 0.731055889 | Y | Y |
| Nfatc2 | Nfatc2 | NW_006501450.1:670001-675000 | NW_006501450.1:565237 | 0.210352716 | 1 | 0.286971912 | Y | Y |
| Nfya | Nfya | NW_006502214.1:185001-190000 | NW_006502214.1:193203 | 0.195088384 | 1 | 0.487821055 | Y | Y |
| Ngef | Ngef | NW_006501258.1:3140001-3145000 | NW_006501258.1:3211837 | 0.114911914 | 0.999235339 | 0.3708065 | Y | Y |

| | | | | | | | | |
|---|---|---|---|---|---|---|---|---|
| Nhlrc3 | Nhlrc3 | NW_006501125.1:4952501-4957500 | NW_006501125.1:4966702 | 0.410476464 | 1 | 0.614164504 | Y | Y |
| Nhsl2 | Nhsl2 | NW_006502024.1:610001-615000 | NW_006502024.1:605507 | 0.250214065 | 1 | 0.495641801 | Y | Y |
| Nid2 | Nid2 | NW_006502672.1:145001-150000 | NW_006502672.1:162423 | 0.752806252 | 1 | 0.597007196 | Y | Y |
| Nin | Nin | NW_006501293.1:2960001-2965000 | NW_006501293.1:2857863 | 0.237812212 | 1 | 0.339094746 | Y | Y |
| Nipal2 | Nipal2 | NW_006501126.1:1375001-1380000 | NW_006501126.1:1222358 | 0.157885378 | 0.99994118 | 0.367836315 | Y | Y |
| Nlgn1 | Nlgn1 | NW_006501046.1:11272501-11277500 | NW_006501046.1:11281042 | 0.959664391 | 1 | 0.699772717 | | |
| Nlrx1 | Nlrx1 | NW_006501652.1:480001-485000 | NW_006501652.1:481616 | 0.328220649 | 1 | 0.725997051 | Y | Y |
| Nnat | Nnat | NW_006501257.1:617501-622500 | NW_006501257.1:620180 | 1.036746015 | 1 | 0.3430331 | Y | Y |
| Nnmt | Nnmt | NW_006501163.1:600001-605000 | NW_006501163.1:601121 | 0.855380182 | 1 | 0.460193801 | | Y |
| Noct | Noct | NW_006501125.1:7230001-7235000 | NW_006501125.1:7234856 | 0.55250643 | 1 | 0.705734316 | Y | Y |
| Nol3 | Nol3 | NW_006501344.1:480001-485000 | NW_006501344.1:480088 | 0.132186193 | 0.99976472 | 0.409782052 | Y | Y |
| Nol4 | Nol4 | NW_006501038.1:8802501-8807500 | NW_006501038.1:8675081 | 0.44995445 | 1 | 0.457592416 | | |
| Nov | Nov | NW_006501212.1:2492501-2497500 | NW_006501212.1:2488617 | 0.223154565 | 1 | 0.556301958 | | |
| Nox4 | Nox4 | NW_006501965.1:32501-37500 | NW_006501965.1:146727 | 0.130341322 | 0.99976472 | 0.326760414 | Y | Y |
| Npepl1 | Npepl1 | NW_006501107.1:992501-997500 | NW_006501107.1:994625 | 0.579394715 | 1 | 0.790971321 | Y | Y |
| Nr3c1 | Nr3c1 | NW_006501119.1:1045001-1050000 | NW_006501119.1:1034979 | 0.445301564 | 1 | 0.614097228 | Y | Y |
| Nrep | NA | NW_006501133.1:2115001-2120000 | NW_006501133.1:2116816 | 0.143762591 | 0.99982354 | 0.516657731 | Y | Y |
| Nsmaf | Nsmaf | NW_006501440.1:370001-375000 | NW_006501440.1:357942 | 0.184967326 | 1 | 0.324843125 | Y | Y |
| Nsmce2 | Nsmce2 | NW_006501508.1:470001-475000 | NW_006501508.1:411135 | 0.27844419 | 1 | 0.784704459 | Y | Y |
| Ntm | Ntm | NW_006501112.1:2625001-2630000 | NW_006501112.1:2643502 | 0.30093302 | 1 | 0.576396386 | | |
| Nudt6 | Nudt6 | NW_006501581.1:1152501-1157500 | NW_006501581.1:1141835 | 0.192801634 | 1 | 0.655018903 | Y | Y |
| Nxf1 | Nxf1 | NW_006501905.1:312501-317500 | NW_006501905.1:321275 | 0.112532413 | 0.999058879 | 0.4589359 | Y | Y |
| Nxpe2 | Nxpe2 | NW_006501163.1:1027501-1032500 | NW_006501163.1:1022505 | 0.431825321 | 1 | 0.613614032 | Y | Y |
| Ocln | Ocln | NW_006501218.1:2160001-2165000 | NW_006501218.1:2148661 | 0.140669457 | 0.99982354 | 0.810895565 | Y | Y |
| Odf1 | Odf1 | NW_006501220.1:2292501-2297500 | NW_006501220.1:2286949 | 0.153134251 | 0.99988236 | 0.456945464 | | |
| Olah | NA | NW_006501315.1:1445001-1450000 | NW_006501315.1:1428307 | 0.269909185 | 1 | 0.304113376 | | |
| Olfm2 | Olfm2 | NW_006501353.1:1005001-1010000 | NW_006501353.1:1008738 | 0.112044754 | 0.999058879 | 0.699707154 | Y | Y |
| Olfml2a | Olfml2a | NW_006501634.1:335001-340000 | NW_006501634.1:319011 | 0.415319126 | 1 | 0.335630662 | Y | Y |

| | | | | | | | | |
|---|---|---|---|---|---|---|---|---|
| Opcml | Opcml | NW_006501112.1:2735001-2740000 | NW_006501112.1:2716124 | 0.189766779 | 1 | 0.828248356 | | |
| Oprk1 | Oprk1 | NW_006501700.1:242501-247500 | NW_006501700.1:234010 | 0.180179136 | 1 | 0.620016273 | | |
| Orc4 | Orc4 | NW_006501077.1:7162501-7167500 | NW_006501077.1:7173594 | 0.113218101 | 0.999058879 | 0.338811114 | Y | Y |
| Ormdl2 | Ormdl2 | NW_006501362.1:180001-185000 | NW_006501362.1:185641 | 0.661052646 | 1 | 0.565264583 | Y | Y |
| Osbpl2 | Osbpl2 | NW_006501107.1:4197501-4202500 | NW_006501107.1:4229106 | 0.863815889 | 1 | 0.445200955 | Y | Y |
| P2ry1 | P2ry1 | NW_006501060.1:105001-110000 | NW_006501060.1:103701 | 0.265236247 | 1 | 0.6096692 | Y | Y |
| P2ry12 | P2ry12 | NW_006501814.1:827501-832500 | NW_006501814.1:788673 | 0.467496828 | 1 | 0.719936459 | | Y |
| P2ry13 | P2ry13 | NW_006501814.1:770001-775000 | NW_006501814.1:771748 | 0.404924202 | 1 | 0.715364129 | | Y |
| P2ry14 | P2ry14 | NW_006501814.1:670001-675000 | NW_006501814.1:651909 | 0.631302538 | 1 | 0.667167679 | Y | Y |
| Pabpc4l | Pabpc4l | NW_006502486.1:557501-562500 | NW_006502486.1:562217 | 0.333968053 | 1 | 0.682442641 | Y | |
| Paip2 | Paip2 | NW_006501530.1:157501-162500 | NW_006501530.1:157654 | 0.179772707 | 1 | 0.370322624 | Y | Y |
| Pan2 | Pan2 | NW_006501362.1:685001-690000 | NW_006501362.1:689734 | 0.469909771 | 1 | 0.73597768 | Y | Y |
| Panx3 | Panx3 | NW_006502520.1:50001-55000 | NW_006502520.1:61448 | 0.460596259 | 1 | 0.3876829 | | |
| Pcdh10 | Pcdh10 | NW_006501799.1:1007501-1012500 | NW_006501799.1:1081845 | 0.16089383 | 1 | 0.691579313 | | |
| Pcdh18 | Pcdh18 | NW_006501125.1:8607501-8612500 | NW_006501125.1:8601278 | 0.37503697 | 1 | 0.592892777 | Y | |
| Pck1 | Pck1 | NW_006501290.1:25001-30000 | NW_006501290.1:29199 | 0.291489563 | 1 | 0.491902597 | | |
| Pcsk1 | Pcsk1 | NW_006501335.1:4732501-4737500 | NW_006501335.1:4742944 | 0.214969122 | 1 | 0.739025645 | Y | Y |
| Pdcl | Pdcl | NW_006501634.1:2025001-2030000 | NW_006501634.1:2036358 | 0.169544852 | 1 | 0.448621347 | Y | Y |
| Pdk4 | Pdk4 | NW_006501346.1:682501-687500 | NW_006501346.1:685871 | 0.156210191 | 0.99994118 | 0.38379809 | Y | Y |
| Pdzd3 | Pdzd3 | NW_006501652.1:460001-465000 | NW_006501652.1:463232 | 0.260215442 | 1 | 0.437647046 | Y | |
| Peg10 | Peg10 | NW_006501550.1:85001-90000 | NW_006501550.1:86837 | 0.693585819 | 1 | 0.474034659 | Y | Y |
| Peg3 | Peg3 | NW_006501712.1:890001-895000 | NW_006501712.1:889216 | 0.967167996 | 1 | 0.780894028 | Y | Y |
| Pex1 | Pex1 | NW_006501550.1:2107501-2112500 | NW_006501550.1:2093531 | 0.111923711 | 0.999058879 | 0.530369814 | Y | Y |
| Pex5l | Pex5l | NW_006501046.1:3587501-3592500 | NW_006501046.1:3738619 | 0.786956795 | 1 | 0.655433861 | | |
| Pgr | Pgr | NW_006501245.1:4145001-4150000 | NW_006501245.1:4149562 | 0.709332432 | 1 | 0.789861329 | Y | Y |
| Pgrmc2 | Pgrmc2 | NW_006501366.1:1942501-1947500 | NW_006501366.1:1943353 | 0.186480408 | 1 | 0.45586145 | Y | Y |
| Phc3 | Phc3 | NW_006501046.1:7092501-7097500 | NW_006501046.1:7071924 | 0.190352881 | 1 | 0.632921679 | Y | Y |
| Phf14 | Phf14 | NW_006501259.1:845001-850000 | NW_006501259.1:809289 | 0.405614656 | 1 | 0.405043201 | Y | Y |

| | | | | | | | | |
|---|---|---|---|---|---|---|---|---|
| Phf20 | Phf20 | NW_006501396.1:1405001-1410000 | NW_006501396.1:1422482 | 0.148628646 | 0.99982354 | 0.310214183 | Y | Y |
| Phf3 | Phf3 | NW_006501626.1:197501-202500 | NW_006501626.1:194050 | 0.187606876 | 1 | 0.545801724 | Y | Y |
| Phldb1 | Phldb1 | NW_006501694.1:292501-297500 | NW_006501694.1:294178 | 0.39135596 | 1 | 0.77839712 | Y | Y |
| Pid1 | Pid1 | NW_006501258.1:252501-257500 | NW_006501258.1:134346 | 0.131532677 | 0.99976472 | 0.585733456 | | |
| Pik3ca | Pik3ca | NW_006501046.1:4280001-4285000 | NW_006501046.1:4270988 | 0.391716388 | 1 | 0.686348701 | Y | Y |
| Pip4k2c | Pip4k2c | NW_006501066.1:567501-572500 | NW_006501066.1:566278 | 0.228683484 | 1 | 0.625348737 | Y | Y |
| Pkhd1 | Pkhd1 | NW_006501981.1:320001-325000 | NW_006501981.1:118275 | 0.13121442 | 0.99976472 | 0.359448123 | | |
| Pkhd1l1 | Pkhd1l1 | NW_006501143.1:1267501-1272500 | NW_006501143.1:1258551 | 0.458913042 | 1 | 0.631361485 | Y | Y |
| Pknox2 | Pknox2 | NW_006501547.1:430001-435000 | NW_006501547.1:431753 | 0.190665523 | 1 | 0.520236284 | Y | Y |
| Pla2g15 | Pla2g15 | NW_006501344.1:1475001-1480000 | NW_006501344.1:1483816 | 0.158619334 | 0.99994118 | 0.433044812 | Y | Y |
| Plch1 | Plch1 | NW_006501060.1:2525001-2530000 | NW_006501060.1:2440272 | 1.007467574 | 1 | 0.70415092 | | |
| Plcl2 | Plcl2 | NW_006501270.1:520001-525000 | NW_006501270.1:523938 | 0.208746651 | 1 | 0.432356322 | Y | Y |
| Pld1 | Pld1 | NW_006501046.1:8625001-8630000 | NW_006501046.1:8571041 | 0.725632678 | 1 | 0.717410536 | Y | Y |
| Plekha5 | Plekha5 | NW_006501111.1:1882501-1887500 | NW_006501111.1:1933873 | 0.261028005 | 1 | 0.352650197 | Y | Y |
| Plk4 | Plk4 | NW_006501366.1:1702501-1707500 | NW_006501366.1:1711879 | 0.371016887 | 1 | 0.707715783 | Y | Y |
| Polh | Polh | NW_006501277.1:137501-142500 | NW_006501277.1:149288 | 0.176747756 | 1 | 0.429014069 | Y | Y |
| Polk | Polk | NW_006501306.1:1152501-1157500 | NW_006501306.1:1122737 | 0.165202469 | 1 | 0.433494474 | Y | Y |
| Pon1 | Pon1 | NW_006501346.1:400001-405000 | NW_006501346.1:391868 | 0.557330294 | 1 | 0.321594824 | | |
| Pon2 | Pon2 | NW_006501346.1:482501-487500 | NW_006501346.1:485947 | 0.17950141 | 1 | 0.617268354 | Y | Y |
| Postn | Postn | NW_006501125.1:4032501-4037500 | NW_006501125.1:4011588 | 0.584930902 | 1 | 0.634910514 | Y | Y |
| Pou2f3 | Pou2f3 | NW_006501191.1:3262501-3267500 | NW_006501191.1:3251063 | 0.269830981 | 1 | 0.554663367 | | |
| Ppp1r9a | Ppp1r9a | NW_006501346.1:320001-325000 | NW_006501346.1:386618 | 0.532762601 | 1 | 0.559748761 | Y | Y |
| Ppp3cb | Ppp3cb | NW_006501274.1:3805001-3810000 | NW_006501274.1:3771055 | 0.449816326 | 1 | 0.303711258 | Y | Y |
| Ppp4r2 | Ppp4r2 | NW_006501059.1:10745001-10750000 | NW_006501059.1:1075246<br>1 | 0.215060706 | 1 | 0.393506161 | Y | Y |
| Prdm10 | Prdm10 | NW_006501112.1:67501-72500 | NW_006501112.1:41233 | 0.223160504 | 1 | 0.702748542 | Y | Y |
| Prelid3b | Slmo2 | NW_006501107.1:1347501-1352500 | NW_006501107.1:1347025 | 0.872322468 | 1 | 0.466386755 | Y | Y |
| Prkci | Prkci | NW_006501046.1:7220001-7225000 | NW_006501046.1:7233009 | 0.321046781 | 1 | 0.711316418 | Y | Y |
| Proser1 | Proser1 | NW_006501125.1:4952501-4957500 | NW_006501125.1:4933576 | 0.410476464 | 1 | 0.483062212 | Y | Y |

| | | | | | | | | |
|---|---|---|---|---|---|---|---|---|
| Prpf18 | Prpf18 | NW_006501315.1:122501-127500 | NW_006501315.1:143787 | 0.188050388 | 1 | 0.317324159 | Y | Y |
| Prph2 | Prph2 | NW_006502601.1:137501-142500 | NW_006502601.1:149423 | 0.124010848 | 0.999529439 | 0.549152477 | | |
| Ptbp1 | Ptbp1 | NW_006501136.1:2252501-2257500 | NW_006501136.1:2255139 | 0.23629325 | 1 | 0.3908413 | Y | Y |
| Ptk7 | Ptk7 | NW_006501519.1:277501-282500 | NW_006501519.1:249222 | 0.141564351 | 0.99982354 | 0.34765436 | Y | Y |
| Ptp4a1 | Ptp4a1 | NW_006501626.1:350001-355000 | NW_006501626.1:355355 | 0.127155964 | 0.99964708 | 0.47306722 | Y | Y |
| Ptprc | Ptprc | NW_006501213.1:1662501-1667500 | NW_006501213.1:1720723 | 0.151416492 | 0.99988236 | 0.439913657 | Y | Y |
| Ptprz1 | Ptprz1 | NW_006501405.1:145001-150000 | NW_006501405.1:188227 | 1.168603908 | 1 | 0.646040873 | | |
| Pura | Pura | NW_006501530.1:852501-857500 | NW_006501530.1:859847 | 0.530479889 | 1 | 0.745457171 | Y | Y |
| Pus3 | Pus3 | NW_006501164.1:3210001-3215000 | NW_006501164.1:3213374 | 0.416713357 | 1 | 0.645344534 | Y | Y |
| R3hdm1 | R3hdm1 | NW_006502075.1:135001-140000 | NW_006502075.1:75783 | 0.135193162 | 0.99982354 | 0.31025088 | Y | Y |
| R3hdm2 | R3hdm2 | NW_006501066.1:275001-280000 | NW_006501066.1:329995 | 0.340865041 | 1 | 0.640320297 | Y | Y |
| Rab22a | Rab22a | NW_006501107.1:587501-592500 | NW_006501107.1:591853 | 0.620628386 | 1 | 0.777960641 | Y | Y |
| Rab23 | Rab23 | NW_006501695.1:152501-157500 | NW_006501695.1:181849 | 0.18158965 | 1 | 0.467965179 | Y | Y |
| Rab33b | Rab33b | NW_006501125.1:6957501-6962500 | NW_006501125.1:6961448 | 0.421711178 | 1 | 0.539713027 | Y | Y |
| Rab39b | Rab39b | NW_006502407.1:27501-32500 | NW_006502407.1:30463 | 0.481984323 | 1 | 0.721642502 | | |
| Rab3d | Rab3d | NW_006501211.1:1740001-1745000 | NW_006501211.1:1736884 | 0.134784233 | 0.99982354 | 0.54539153 | Y | Y |
| Rab5b | Rab5b | NW_006501362.1:322501-327500 | NW_006501362.1:329564 | 0.340942623 | 1 | 0.807305776 | Y | Y |
| Rad17 | Rad17 | NW_006501218.1:2265001-2270000 | NW_006501218.1:2258489 | 0.272450943 | 1 | 0.339172179 | Y | Y |
| Rap1b | Rap1b | NW_006501066.1:9700001-9705000 | NW_006501066.1:9705585 | 0.181902731 | 1 | 0.842723633 | Y | Y |
| Raver1 | Raver1 | NW_006501353.1:565001-570000 | NW_006501353.1:582408 | 0.388791379 | 1 | 0.575043944 | | |
| Rb1cc1 | Rb1cc1 | NW_006501700.1:907501-912500 | NW_006501700.1:955295 | 0.115497217 | 0.999235339 | 0.313411504 | Y | Y |
| Rbm7 | Rbm7 | NW_006501163.1:722501-727500 | NW_006501163.1:733615 | 0.275432829 | 1 | 0.803662213 | Y | Y |
| Rc3h2 | Rc3h2 | NW_006501634.1:2000001-2005000 | NW_006501634.1:2010233 | 0.135193869 | 0.99982354 | 0.617137487 | Y | Y |
| Rcan2 | Rcan2 | NW_006501565.1:675001-680000 | NW_006501565.1:776601 | 0.306907132 | 1 | 0.54779585 | Y | Y |
| Rfesd | Rfesd | NW_006501335.1:3970001-3975000 | NW_006501335.1:3966421 | 0.173769711 | 1 | 0.71769767 | Y | Y |
| Rftn1 | Rftn1 | NW_006502151.1:25001-30000 | NW_006502151.1:27067 | 0.140394754 | 0.99982354 | 0.62052545 | Y | Y |
| Rfx3 | Rfx3 | NW_006501365.1:1085001-1090000 | NW_006501365.1:1139467 | 0.14638503 | 0.99982354 | 0.285188696 | Y | Y |
| Rhag | Rhag | NW_006501383.1:45001-50000 | NW_006501383.1:42804 | 0.253597607 | 1 | 0.480555014 | | |

| | | | | | | | | |
|---|---|---|---|---|---|---|---|---|
| Rhpn2 | Rhpn2 | NW_006501278.1:1392501-1397500 | NW_006501278.1:1415118 | 0.136183396 | 0.99982354 | 0.330723544 | Y | Y |
| Rims2 | Rims2 | NW_006501417.1:1140001-1145000 | NW_006501417.1:1177690 | 1.152457156 | 1 | 0.45611098 | Y | Y |
| Riok3 | Riok3 | NW_006501454.1:120001-125000 | NW_006501454.1:123552 | 0.336377727 | 1 | 0.535046956 | Y | Y |
| Rlim | Rlim | NW_006501568.1:1210001-1215000 | NW_006501568.1:1207402 | 0.113346043 | 0.999117699 | 0.391185061 | Y | Y |
| Rnf133 | Rnf133 | NW_006501405.1:760001-765000 | NW_006501405.1:765640 | 0.360300279 | 1 | 0.645982158 | | |
| Rnf148 | Rnf148 | NW_006501405.1:767501-772500 | NW_006501405.1:770013 | 0.305556555 | 1 | 0.507711419 | | |
| Rnf19a | Rnf19a | NW_006501220.1:190001-195000 | NW_006501220.1:195284 | 0.1126821 | 0.999058879 | 0.606268271 | Y | Y |
| Robo4 | Robo4 | NW_006501547.1:970001-975000 | NW_006501547.1:967380 | 0.15860588 | 0.99994118 | 0.405674842 | Y | Y |
| Rock1 | Rock1 | NW_006501756.1:560001-565000 | NW_006501756.1:617088 | 0.223627479 | 1 | 0.656704569 | Y | Y |
| Rp1 | Rp1 | NW_006501755.1:450001-455000 | NW_006501755.1:446830 | 0.219006362 | 1 | 0.554256863 | | |
| Rp9 | Rp9 | NW_006501574.1:3282501-3287500 | NW_006501574.1:3279937 | 0.241434756 | 1 | 0.405021657 | Y | Y |
| Rprd1a | Rprd1a | NW_006501173.1:4262501-4267500 | NW_006501173.1:4264113 | 0.322300014 | 1 | 0.613651912 | Y | Y |
| Rps25 | Rps25 | NW_006501652.1:627501-632500 | NW_006501652.1:635356 | 0.292948932 | 1 | 0.354949675 | Y | Y |
| Rps28 | Rps28 | NW_006501520.1:407501-412500 | NW_006501520.1:407384 | 0.150249294 | 0.99982354 | 0.617214631 | Y | Y |
| Rrm2b | Rrm2b | NW_006501220.1:1987501-1992500 | NW_006501220.1:1986425 | 0.379214901 | 1 | 0.373925762 | Y | Y |
| Rsad2 | Rsad2 | NW_006501185.1:1770001-1775000 | NW_006501185.1:1770231 | 0.315502644 | 1 | 0.72789209 | Y | Y |
| Rsbn1l | Rsbn1l | NW_006501532.1:842501-847500 | NW_006501532.1:850035 | 0.13018527 | 0.99976472 | 0.383196928 | Y | Y |
| Rtel1 | Rtel1 | NW_006501809.1:612501-617500 | NW_006501809.1:629460 | 0.372610922 | 1 | 0.39003345 | Y | Y |
| Rtl1 | Rtl1 | NW_006501050.1:10262501-10267500 | NW_006501050.1:1026285 4 | 0.49140436 | 1 | 0.72424018 | | |
| S1pr5 | S1pr5 | NW_006501353.1:390001-395000 | NW_006501353.1:394485 | 0.234870941 | 1 | 0.600849708 | | |
| Samd12 | Samd12 | NW_006501212.1:3475001-3480000 | NW_006501212.1:3674029 | 0.298256579 | 1 | 0.693423471 | | |
| Samd9 | NA | NW_006501550.1:1517501-1522500 | NW_006501550.1:1518990 | 0.180066242 | 1 | 0.636552559 | | |
| Satb1 | Satb1 | NW_006501412.1:115001-120000 | NW_006501412.1:46152 | 0.394246273 | 1 | 0.423457747 | Y | Y |
| Sc5d | Sc5d | NW_006501191.1:2307501-2312500 | NW_006501191.1:2312173 | 0.14599988 | 0.99982354 | 0.616014061 | Y | Y |
| Scamp1 | Scamp1 | NW_006501205.1:1367501-1372500 | NW_006501205.1:1370482 | 0.158713134 | 0.99994118 | 0.731817633 | Y | Y |
| Scgb2a2 | NA | NW_006501723.1:1507501-1512500 | NW_006501723.1:1529140 | 0.320343256 | 1 | 0.75710491 | | |
| Scn3b | Scn3b | NW_006501191.1:225001-230000 | NW_006501191.1:241653 | 0.202521559 | 1 | 0.76479798 | Y | |
| Sdr9c7 | Sdr9c7 | NW_006502313.1:75001-80000 | NW_006502313.1:70470 | 0.370545532 | 1 | 0.349423713 | | |

| | | | | | | | | |
|---|---|---|---|---|---|---|---|---|
| Sec24c | Sec24c | NW_006501274.1:3625001-3630000 | NW_006501274.1:3604352 | 0.211723909 | 1 | 0.77188478 | Y | Y |
| Sec62 | Sec62 | NW_006501046.1:6975001-6980000 | NW_006501046.1:6970253 | 0.171174231 | 1 | 0.658449364 | Y | Y |
| Serp1 | Serp1 | NW_006501814.1:2501-7500 | NW_006501814.1:6103 | 0.746847901 | 1 | 0.717956176 | Y | Y |
| Serpinb8 | Serpinb8 | NW_006501717.1:827501-832500 | NW_006501717.1:828709 | 0.225507377 | 1 | 0.701719473 | Y | Y |
| Sertm1 | NA | NW_006501125.1:3475001-3480000 | NW_006501125.1:3479015 | 0.234129447 | 1 | 0.607432988 | | |
| Sesn3 | Sesn3 | NW_006501653.1:1795001-1800000 | NW_006501653.1:1821509 | 0.41096463 | 1 | 0.83861651 | Y | Y |
| Setd7 | Setd7 | NW_006501125.1:6917501-6922500 | NW_006501125.1:6936230 | 0.72966523 | 1 | 0.702886934 | Y | Y |
| Sgol1 | Sgol1 | NW_006501911.1:162501-167500 | NW_006501911.1:165243 | 0.395301205 | 1 | 0.45725988 | | |
| Sh3glb1 | Sh3glb1 | NW_006501323.1:77501-82500 | NW_006501323.1:80790 | 0.164307317 | 1 | 0.494345897 | Y | Y |
| Shisa9 | Shisa9 | NW_006501248.1:2557501-2562500 | NW_006501248.1:2273605 | 0.116922787 | 0.999235339 | 0.610765561 | | |
| Shmt2 | Shmt2 | NW_006501066.1:242501-247500 | NW_006501066.1:243093 | 0.141332545 | 0.99982354 | 0.533727283 | Y | Y |
| Shoc2 | Shoc2 | NW_006501901.1:602501-607500 | NW_006501901.1:612142 | 0.120444152 | 0.999470619 | 0.607592403 | Y | Y |
| Siae | Siae | NW_006502520.1:80001-85000 | NW_006502520.1:80521 | 0.12196444 | 0.999470619 | 0.385596507 | Y | Y |
| Sik3 | Sik3 | NW_006502156.1:210001-215000 | NW_006502156.1:238947 | 0.478722965 | 1 | 0.589041365 | Y | Y |
| Sim2 | Sim2 | NW_006501195.1:500001-505000 | NW_006501195.1:527305 | 0.440526074 | 1 | 0.379642297 | | |
| Skil | Skil | NW_006501046.1:7290001-7295000 | NW_006501046.1:7303420 | 0.432567196 | 1 | 0.711456007 | Y | Y |
| Slc10a2 | Slc10a2 | NW_006501402.1:4060001-4065000 | NW_006501402.1:4051498 | 0.153322766 | 0.99988236 | 0.475347492 | | |
| Slc13a1 | Slc13a1 | NW_006501405.1:1217501-1222500 | NW_006501405.1:1168952 | 0.550816123 | 1 | 0.366241652 | | |
| Slc13a5 | Slc13a5 | NW_006501614.1:1160001-1165000 | NW_006501614.1:1152230 | 0.140159287 | 0.99982354 | 0.290410145 | | |
| Slc16a7 | Slc16a7 | NW_006501066.1:2427501-2432500 | NW_006501066.1:2482066 | 0.170931249 | 1 | 0.476238669 | Y | Y |
| Slc22a8 | Slc22a8 | NW_006501905.1:157501-162500 | NW_006501905.1:160412 | 0.484697429 | 1 | 0.736708499 | | |
| Slc25a27 | Slc25a27 | NW_006501603.1:357501-362500 | NW_006501603.1:413860 | 0.331515519 | 1 | 0.460869833 | Y | Y |
| Slc25a31 | Slc25a31 | NW_006501366.1:1605001-1610000 | NW_006501366.1:1601388 | 0.204701054 | 1 | 0.37103813 | | |
| Slc25a32 | Slc25a32 | NW_006501220.1:3102501-3107500 | NW_006501220.1:3107565 | 0.197892956 | 1 | 0.648200986 | Y | Y |
| Slc25a46 | Slc25a46 | NW_006501133.1:3937501-3942500 | NW_006501133.1:3941586 | 0.294557249 | 1 | 0.496117231 | Y | Y |
| Slc29a1 | Slc29a1 | NW_006501277.1:825001-830000 | NW_006501277.1:829518 | 0.114028427 | 0.999117699 | 0.29405343 | Y | Y |
| Slc2a2 | Slc2a2 | NW_006501046.1:7952501-7957500 | NW_006501046.1:7933046 | 0.74644472 | 1 | 0.622835596 | | |
| Slc2a5 | Slc2a5 | NW_006501092.1:1712501-1717500 | NW_006501092.1:1723115 | 0.151528409 | 0.99988236 | 0.466629507 | | Y |

| | | | | | | | | |
|---|---|---|---|---|---|---|---|---|
| Slc30a8 | Slc30a8 | NW_006501212.1:4715001-4720000 | NW_006501212.1:4713895 | 0.268477444 | 1 | 0.669561903 | | |
| Slc33a1 | Slc33a1 | NW_006501060.1:2717501-2722500 | NW_006501060.1:2701094 | 0.714147275 | 1 | 0.665023581 | Y | Y |
| Slc35a4 | Slc35a4 | NW_006502100.1:17501-22500 | NW_006502100.1:22904 | 0.134749789 | 0.99982354 | 0.733365815 | Y | Y |
| Slc35e3 | Slc35e3 | NW_006501066.1:9772501-9777500 | NW_006501066.1:9788588 | 0.218117771 | 1 | 0.649996405 | Y | Y |
| Slc39a5 | Slc39a5 | NW_006501362.1:610001-615000 | NW_006501362.1:619441 | 0.276231356 | 1 | 0.785898537 | Y | Y |
| Slc39a6 | Slc39a6 | NW_006501173.1:4340001-4345000 | NW_006501173.1:4333088 | 0.284816933 | 1 | 0.433462412 | Y | Y |
| Slc44a2 | Slc44a2 | NW_006501353.1:277501-282500 | NW_006501353.1:279785 | 0.135691484 | 0.99982354 | 0.335868781 | Y | Y |
| Slc4a9 | Slc4a9 | NW_006501530.1:1112501-1117500 | NW_006501530.1:1109932 | 0.425676261 | 1 | 0.682437477 | | |
| Slc7a1 | Slc7a1 | NW_006501160.1:1970001-1975000 | NW_006501160.1:1982012 | 0.125512692 | 0.999529439 | 0.438534762 | Y | Y |
| Slc7a11 | Slc7a11 | NW_006501125.1:7965001-7970000 | NW_006501125.1:8030196 | 0.15433534 | 0.99994118 | 0.370413267 | | Y |
| Slc7a14 | Slc7a14 | NW_006501046.1:7407501-7412500 | NW_006501046.1:7394777 | 0.222051261 | 1 | 0.630156305 | | |
| Slco1a2 | Slco1a4 | NW_006501111.1:427501-432500 | NW_006501111.1:462434 | 0.303516384 | 1 | 0.649910063 | | |
| Slco1b3 | Slco1b2 | NW_006501111.1:787501-792500 | NW_006501111.1:723715 | 0.306541626 | 1 | 0.521996784 | | |
| Slco1c1 | Slco1c1 | NW_006501111.1:877501-882500 | NW_006501111.1:915300 | 0.121686749 | 0.999470619 | 0.495493552 | | |
| Slco4c1 | Slco4c1 | NW_006501305.1:3752501-3757500 | NW_006501305.1:3773466 | 0.145977182 | 0.99982354 | 0.330392247 | | |
| Slf1 | Slf1 | NW_006501335.1:2942501-2947500 | NW_006501335.1:2952926 | 0.524983381 | 1 | 0.816821926 | Y | Y |
| Slmap | Slmap | NW_006501043.1:82501-87500 | NW_006501043.1:34073 | 0.111681227 | 0.999000059 | 0.277756502 | Y | Y |
| Sltm | Sltm | NW_006501153.1:867501-872500 | NW_006501153.1:840278 | 0.11546206 | 0.999235339 | 0.313827586 | Y | Y |
| Smarcc2 | Smarcc2 | NW_006501362.1:547501-552500 | NW_006501362.1:524165 | 0.524017353 | 1 | 0.310547764 | Y | Y |
| Smpd3 | Smpd3 | NW_006501344.1:1587501-1592500 | NW_006501344.1:1578174 | 0.12310052 | 0.999529439 | 0.466710749 | Y | Y |
| Snap91 | Snap91 | NW_006501243.1:1692501-1697500 | NW_006501243.1:1745241 | 0.422619577 | 1 | 0.737344147 | Y | |
| Sntb1 | Sntb1 | NW_006501212.1:1457501-1462500 | NW_006501212.1:1551811 | 0.174897755 | 1 | 0.549433139 | Y | Y |
| Snx14 | Snx14 | NW_006501243.1:205001-210000 | NW_006501243.1:201914 | 0.209443535 | 1 | 0.675180948 | Y | Y |
| Snx19 | Snx19 | NW_006501112.1:1100001-1105000 | NW_006501112.1:1142531 | 0.264771053 | 1 | 0.825557792 | Y | Y |
| Snx29 | Snx29 | NW_006501248.1:2815001-2820000 | NW_006501248.1:2818550 | 0.293155096 | 1 | 0.581355784 | Y | Y |
| Sohlh2 | NA | NW_006501125.1:3152501-3157500 | NW_006501125.1:3153426 | 0.243005816 | 1 | 0.493668541 | | |
| Sorcs3 | Sorcs3 | NW_006501231.1:2857501-2862500 | NW_006501231.1:2734875 | 0.377793514 | 1 | 0.509348431 | | |
| Sorl1 | Sorl1 | NW_006501191.1:2022501-2027500 | NW_006501191.1:2015644 | 0.32109782 | 1 | 0.776935219 | Y | Y |

| | | | | | | | | |
|---|---|---|---|---|---|---|---|---|
| Sox18 | Sox18 | NW_006501809.1:212501-217500 | NW_006501809.1:219531 | 0.308183303 | 1 | 0.337395733 | Y | Y |
| Sox2 | Sox2 | NW_006501046.1:2012501-2017500 | NW_006501046.1:2014523 | 0.245905307 | 1 | 0.682846206 | | |
| Spata24 | Spata24 | NW_006501530.1:192501-197500 | NW_006501530.1:187517 | 0.119672494 | 0.999352979 | 0.585559014 | Y | Y |
| Spata5 | Spata5 | NW_006501581.1:1130001-1135000 | NW_006501581.1:1141835 | 0.613235141 | 1 | 0.655018903 | Y | Y |
| Spdya | Spdya | NW_006501510.1:2822501-2827500 | NW_006501510.1:2826581 | 0.111898 | 0.999058879 | 0.378650788 | Y | Y |
| Spg20 | Spg20 | NW_006501125.1:3232501-3237500 | NW_006501125.1:3227026 | 0.346888368 | 1 | 0.575389749 | Y | Y |
| Spo11 | Spo11 | NW_006501290.1:200001-205000 | NW_006501290.1:196377 | 0.128859375 | 0.99976472 | 0.430311666 | | |
| Spryd4 | Spryd4 | NW_006501362.1:840001-845000 | NW_006501362.1:842167 | 0.149392327 | 0.99982354 | 0.56190089 | Y | Y |
| Srek1 | Srek1 | NW_006501218.1:5215001-5220000 | NW_006501218.1:5207074 | 0.202245022 | 1 | 0.777727733 | Y | Y |
| Srgap1 | Srgap1 | NW_006501066.1:5802501-5807500 | NW_006501066.1:5805306 | 0.143818419 | 0.99982354 | 0.429292102 | Y | Y |
| Srpra | Srpr | NW_006501164.1:2850001-2855000 | NW_006501164.1:2853432 | 0.200935143 | 1 | 0.651225343 | Y | Y |
| Ss18 | Ss18 | NW_006501225.1:2977501-2982500 | NW_006501225.1:3044834 | 0.321635488 | 1 | 0.561244553 | Y | Y |
| St14 | St14 | NW_006501112.1:377501-382500 | NW_006501112.1:331430 | 0.126365624 | 0.99958826 | 0.766631453 | Y | Y |
| Stard13 | Stard13 | NW_006501358.1:2035001-2040000 | NW_006501358.1:2023919 | 0.159799383 | 1 | 0.527672391 | Y | Y |
| Stard4 | Stard4 | NW_006501133.1:2315001-2320000 | NW_006501133.1:2318937 | 0.133651413 | 0.99982354 | 0.605784106 | Y | Y |
| Stat2 | Stat2 | NW_006501362.1:730001-735000 | NW_006501362.1:715394 | 0.368780642 | 1 | 0.631042678 | Y | Y |
| Stat6 | Stat6 | NW_006501066.1:97501-102500 | NW_006501066.1:86991 | 0.332628173 | 1 | 0.797535785 | Y | Y |
| Stk3 | Stk3 | NW_006501126.1:1162501-1167500 | NW_006501126.1:1167378 | 0.164859793 | 1 | 0.417880214 | Y | Y |
| Stk33 | Stk33 | NW_006501141.1:655001-660000 | NW_006501141.1:613931 | 0.123235082 | 0.999529439 | 0.298804347 | Y | |
| Strbp | Strbp | NW_006501634.1:1777501-1782500 | NW_006501634.1:1779831 | 0.58374213 | 1 | 0.344933188 | Y | Y |
| Stt3a | Stt3a | NW_006501547.1:235001-240000 | NW_006501547.1:260094 | 0.334597831 | 1 | 0.568497216 | Y | Y |
| Stt3b | Stt3b | NW_006501039.1:3560001-3565000 | NW_006501039.1:3576936 | 0.12624534 | 0.99958826 | 0.295833518 | Y | Y |
| Stx16 | Stx16 | NW_006501107.1:962501-967500 | NW_006501107.1:967816 | 1.980784716 | 1 | 0.794876727 | Y | Y |
| Sucnr1 | Sucnr1 | NW_006502570.1:45001-50000 | NW_006502570.1:50636 | 0.368382313 | 1 | 0.614720277 | | |
| Suox | Suox | NW_006501362.1:330001-335000 | NW_006501362.1:337245 | 0.294374871 | 1 | 0.809059757 | Y | Y |
| Supt20h | Supt20 | NW_006501125.1:3700001-3705000 | NW_006501125.1:3670022 | 1.035154477 | 1 | 0.530124362 | Y | Y |
| Sybu | Sybu | NW_006501143.1:1120001-1125000 | NW_006501143.1:1122867 | 0.128257566 | 0.9997059 | 0.450197427 | Y | Y |
| Synpo2l | Synpo2l | NW_006501274.1:3635001-3640000 | NW_006501274.1:3645905 | 0.300236158 | 1 | 0.696532682 | Y | |

| | | | | | | | | |
|---|---|---|---|---|---|---|---|---|
| Syt4 | Syt4 | NW_006501133.1:4092501-4097500 | NW_006501133.1:4092585 | 0.259276135 | 1 | 0.352559764 | | |
| Taf2 | Taf2 | NW_006501212.1:2162501-2167500 | NW_006501212.1:2223985 | 0.27981996 | 1 | 0.598118747 | Y | Y |
| Taf4b | Taf4b | NW_006501225.1:2807501-2812500 | NW_006501225.1:2749362 | 0.148877189 | 0.99982354 | 0.670119363 | Y | Y |
| Taf8 | Taf8 | NW_006501434.1:107501-112500 | NW_006501434.1:118896 | 0.243315982 | 1 | 0.406264085 | Y | Y |
| Tas2r40 | Tas2r144 | NW_006502289.1:392501-397500 | NW_006502289.1:396952 | 0.388900432 | 1 | 0.625296674 | | |
| Tatdn1 | Tatdn1 | NW_006501508.1:1107501-1112500 | NW_006501508.1:1094839 | 0.235596278 | 1 | 0.324532166 | Y | Y |
| Tbc1d30 | Tbc1d30 | NW_006501066.1:6335001-6340000 | NW_006501066.1:6377770 | 0.213586546 | 1 | 0.428093171 | Y | Y |
| Tbc1d31 | Tbc1d31 | NW_006501508.1:2162501-2167500 | NW_006501508.1:2185140 | 0.188933721 | 1 | 0.402320852 | Y | Y |
| Tbc1d5 | Tbc1d5 | NW_006501270.1:830001-835000 | NW_006501270.1:760780 | 0.118435343 | 0.999352979 | 0.6088377 | Y | Y |
| Tbcel | Tbcel | NW_006501191.1:2522501-2527500 | NW_006501191.1:2518389 | 0.156266989 | 0.99994118 | 0.593545609 | Y | Y |
| Tbk1 | Tbk1 | NW_006501066.1:6047501-6052500 | NW_006501066.1:6050074 | 0.229575581 | 1 | 0.457409322 | Y | Y |
| Tcfl5 | Tcfl5 | NW_006501107.1:4720001-4725000 | NW_006501107.1:4725010 | 0.111752444 | 0.999000059 | 0.60223682 | Y | |
| Tecta | Tecta | NW_006501191.1:2452501-2457500 | NW_006501191.1:2455605 | 0.401574262 | 1 | 0.589610035 | | |
| Tfap2b | Tfap2b | NW_006501316.1:582501-587500 | NW_006501316.1:610644 | 0.333232564 | 1 | 0.320191562 | | |
| Tfap2d | Tfap2d | NW_006501316.1:535001-540000 | NW_006501316.1:471136 | 0.136337003 | 0.99982354 | 0.417130709 | | |
| Tfpi2 | NA | NW_006501550.1:825001-830000 | NW_006501550.1:822940 | 0.371558464 | 1 | 0.625374721 | Y | Y |
| Thy1 | Thy1 | NW_006501652.1:255001-260000 | NW_006501652.1:259243 | 0.144126316 | 0.99982354 | 0.77190913 | Y | Y |
| Thyn1 | Thyn1 | NW_006501112.1:4632501-4637500 | NW_006501112.1:4637030 | 0.269255147 | 1 | 0.814382 | Y | Y |
| Tjap1 | Tjap1 | NW_006501277.1:42501-47500 | NW_006501277.1:48936 | 0.121765649 | 0.999470619 | 0.545128753 | Y | Y |
| Tm4sf4 | Tm4sf4 | NW_006501125.1:1112501-1117500 | NW_006501125.1:1103098 | 0.61802586 | 1 | 0.371991889 | | |
| Tmed1 | Tmed1 | NW_006501353.1:102501-107500 | NW_006501353.1:104690 | 0.232926208 | 1 | 0.734616888 | Y | Y |
| Tmem123 | NA | NW_006501245.1:3112501-3117500 | NW_006501245.1:3119438 | 0.174786873 | 1 | 0.80924225 | Y | Y |
| Tmem136 | Tmem136 | NW_006501191.1:3225001-3230000 | NW_006501191.1:3228634 | 0.629331725 | 1 | 0.8318634 | | |
| Tmem151b | Tmem151b | NW_006501277.1:882501-887500 | NW_006501277.1:885053 | 0.134965104 | 0.99982354 | 0.579539095 | | |
| Tmem161b | Tmem161b | NW_006501392.1:727501-732500 | NW_006501392.1:704143 | 0.150815723 | 0.99988236 | 0.605532665 | Y | Y |
| Tmem173 | Tmem173 | NW_006501530.1:287501-292500 | NW_006501530.1:286558 | 0.211087048 | 1 | 0.514626866 | | |
| Tmem212 | Tmem212 | NW_006501046.1:8850001-8855000 | NW_006501046.1:8852470 | 0.288964313 | 1 | 0.663217205 | | |
| Tmem218 | Tmem218 | NW_006501547.1:782501-787500 | NW_006501547.1:789311 | 0.230860084 | 1 | 0.680436482 | Y | Y |

| | | | | | | | | |
|---|---|---|---|---|---|---|---|---|
| Tmem241 | Tmem241 | NW_006501454.1:150001-155000 | NW_006501454.1:295072 | 0.135837522 | 0.99982354 | 0.433348619 | Y | Y |
| Tmem25 | Tmem25 | NW_006501694.1:425001-430000 | NW_006501694.1:421883 | 0.285050749 | 1 | 0.64020204 | | |
| Tmem30a | Tmem30a | NW_006501170.1:2050001-2055000 | NW_006501170.1:2063838 | 0.154116224 | 0.99994118 | 0.383144002 | Y | Y |
| Tmprss5 | Tmprss5 | NW_006501163.1:75001-80000 | NW_006501163.1:77050 | 0.452852035 | 1 | 0.555584541 | | |
| Tnfrsf11b | Tnfrsf11b | NW_006501212.1:2915001-2920000 | NW_006501212.1:2944364 | 0.822232781 | 1 | 0.601401845 | Y | Y |
| Tnfsf10 | Tnfsf10 | NW_006501046.1:9450001-9455000 | NW_006501046.1:9458612 | 0.685839483 | 1 | 0.715364129 | Y | Y |
| Tnik | Tnik | NW_006501046.1:8022501-8027500 | NW_006501046.1:8001418 | 0.752921638 | 1 | 0.72220975 | Y | Y |
| Tnks | Tnks | NW_006501073.1:5837501-5842500 | NW_006501073.1:5798585 | 0.133136963 | 0.99976472 | 0.341960404 | Y | Y |
| Tppp3 | Tppp3 | NW_006501344.1:700001-705000 | NW_006501344.1:705078 | 0.131074668 | 0.99976472 | 0.434558746 | Y | Y |
| Treh | Treh | NW_006501694.1:292501-297500 | NW_006501694.1:293933 | 0.39135596 | 1 | 0.77839712 | Y | Y |
| Trerf1 | NA | NW_006501434.1:260001-265000 | NW_006501434.1:256432 | 0.300657342 | 1 | 0.463249603 | Y | Y |
| Trhr | Trhr | NW_006501143.1:1565001-1570000 | NW_006501143.1:1568644 | 0.353210111 | 1 | 0.76558593 | | |
| Trib1 | Trib1 | NW_006501508.1:347501-352500 | NW_006501508.1:349251 | 0.234934622 | 1 | 0.399713904 | Y | Y |
| Trmt12 | Trmt12 | NW_006501508.1:1157501-1162500 | NW_006501508.1:1159281 | 0.124585029 | 0.999529439 | 0.482205234 | Y | Y |
| Trpc3 | Trpc3 | NW_006501581.1:1877501-1882500 | NW_006501581.1:1895457 | 0.264005249 | 1 | 0.437073598 | Y | Y |
| Trpc4 | Trpc4 | NW_006501125.1:4120001-4125000 | NW_006501125.1:4079806 | 0.523283629 | 1 | 0.551734657 | | |
| Trps1 | Trps1 | NW_006501212.1:6167501-6172500 | NW_006501212.1:6355972 | 0.700283913 | 1 | 0.720303487 | Y | Y |
| Tsc22d2 | Tsc22d2 | NW_006501125.1:47501-52500 | NW_006501125.1:47994 | 0.137010667 | 0.99982354 | 0.722113887 | Y | Y |
| Ttbk1 | Ttbk1 | NW_006501519.1:152501-157500 | NW_006501519.1:127997 | 0.32033128 | 1 | 0.504208911 | | |
| Ttc3 | Ttc3 | NW_006501195.1:800001-805000 | NW_006501195.1:876163 | 0.531798751 | 1 | 0.511368828 | Y | Y |
| Tubb1 | Tubb1 | NW_006501107.1:1335001-1340000 | NW_006501107.1:1339035 | 0.83279346 | 1 | 0.753392401 | Y | |
| Tyk2 | Tyk2 | NW_006501353.1:540001-545000 | NW_006501353.1:549708 | 0.2261345 | 1 | 0.545401342 | Y | Y |
| Tyr | Tyr | NW_006501303.1:1025001-1030000 | NW_006501303.1:1027779 | 0.489096359 | 1 | 0.454359703 | | |
| Ubash3b | Ubash3b | NW_006501191.1:980001-985000 | NW_006501191.1:984448 | 0.257643691 | 1 | 0.820210623 | Y | Y |
| Ube2d2 | Ube2d2a | NW_006501530.1:347501-352500 | NW_006501530.1:358108 | 0.281209061 | 1 | 0.740100561 | Y | Y |
| Ubr2 | Ubr2 | NW_006502601.1:110001-115000 | NW_006502601.1:131217 | 0.674058259 | 1 | 0.40962415 | Y | Y |
| Ubr5 | Ubr5 | NW_006501220.1:2045001-2050000 | NW_006501220.1:2021701 | 0.431769332 | 1 | 0.614117085 | Y | Y |
| Ubxn2b | Ubxn2b | NW_006501440.1:205001-210000 | NW_006501440.1:218508 | 0.248797675 | 1 | 0.558003201 | Y | Y |

| | | | | | | | | |
|---|---|---|---|---|---|---|---|---|
| Ufm1 | Ufm1 | NW_006501125.1:4560001-4565000 | NW_006501125.1:4562937 | 0.41024091 | 1 | 0.640067049 | Y | Y |
| Usp12 | Usp12 | NW_006501160.1:3645001-3650000 | NW_006501160.1:3653686 | 0.201827619 | 1 | 0.329162176 | Y | Y |
| Usp13 | Usp13 | NW_006501046.1:3812501-3817500 | NW_006501046.1:3749415 | 0.448101081 | 1 | 0.728700535 | Y | Y |
| Usp15 | Usp15 | NW_006501066.1:4535001-4540000 | NW_006501066.1:4554044 | 0.375356578 | 1 | 0.81213664 | Y | Y |
| Usp2 | Usp2 | NW_006501652.1:297501-302500 | NW_006501652.1:308333 | 0.276378595 | 1 | 0.338843134 | Y | Y |
| Usp28 | Usp28 | NW_006501163.1:165001-170000 | NW_006501163.1:208277 | 1.319689397 | 1 | 0.822995994 | Y | Y |
| Usp40 | Usp40 | NW_006501258.1:3632501-3637500 | NW_006501258.1:3591648 | 0.135632341 | 0.99982354 | 0.611773426 | Y | Y |
| Usp42 | Usp42 | NW_006501217.1:610001-615000 | NW_006501217.1:613060 | 0.415194993 | 1 | 0.399382082 | Y | Y |
| Usp54 | Usp54 | NW_006501274.1:3757501-3762500 | NW_006501274.1:3698854 | 0.212839534 | 1 | 0.753743329 | Y | Y |
| Uspl1 | Uspl1 | NW_006501160.1:1045001-1050000 | NW_006501160.1:1043815 | 0.173878104 | 1 | 0.490160409 | Y | Y |
| Vapb | Vapb | NW_006501107.1:672501-677500 | NW_006501107.1:672711 | 0.629474316 | 1 | 0.745117889 | Y | Y |
| Vcl | Vcl | NW_006501274.1:3285001-3290000 | NW_006501274.1:3258644 | 0.270427689 | 1 | 0.64466678 | Y | Y |
| Vps13b | Vps13b | NW_006501126.1:147501-152500 | NW_006501126.1:348523 | 0.555391701 | 1 | 0.508989468 | Y | Y |
| Vps26b | Vps26b | NW_006501112.1:4630001-4635000 | NW_006501112.1:4634966 | 0.437021466 | 1 | 0.814382 | Y | Y |
| Vps50 | Vps50 | NW_006501550.1:1270001-1275000 | NW_006501550.1:1190395 | 0.272541284 | 1 | 0.335133958 | Y | Y |
| Wdr33 | Wdr33 | NW_006501133.1:3610001-3615000 | NW_006501133.1:3611405 | 0.156701428 | 0.99994118 | 0.520468344 | Y | Y |
| Wdyhv1 | Wdyhv1 | NW_006501508.1:1930001-1935000 | NW_006501508.1:1940770 | 0.201321982 | 1 | 0.396925051 | Y | Y |
| Wwtr1 | Wwtr1 | NW_006501125.1:1032501-1037500 | NW_006501125.1:1085020 | 0.517480701 | 1 | 0.424484949 | Y | Y |
| Xpot | Xpot | NW_006501066.1:5977501-5982500 | NW_006501066.1:6007627 | 0.35158545 | 1 | 0.660754714 | Y | Y |
| Ythdc1 | Ythdc1 | NW_006501246.1:672501-677500 | NW_006501246.1:677194 | 0.163742889 | 1 | 0.439008547 | Y | Y |
| Zbtb16 | Zbtb16 | NW_006501163.1:552501-557500 | NW_006501163.1:560833 | 0.810268977 | 1 | 0.834575583 | Y | Y |
| Zbtb39 | Zbtb39 | NW_006503545.1:22501-27500 | NW_006503545.1:21754 | 0.181607913 | 1 | 0.79453339 | Y | Y |
| Zbtb44 | Zbtb44 | NW_006501112.1:400001-405000 | NW_006501112.1:401926 | 0.155758194 | 0.99994118 | 0.795021512 | Y | Y |
| Zbtb45 | Zbtb45 | NW_006502295.1:132501-137500 | NW_006502295.1:135967 | 0.119439326 | 0.999352979 | 0.625563439 | Y | Y |
| Zbtb6 | Zbtb6 | NW_006501634.1:1932501-1937500 | NW_006501634.1:1939509 | 0.158985582 | 0.99994118 | 0.668882822 | Y | Y |
| Zc3h7a | Zc3h7a | NW_006501248.1:3472501-3477500 | NW_006501248.1:3473153 | 0.11380346 | 0.999117699 | 0.406969941 | Y | Y |
| Zdbf2 | Zdbf2 | NW_006501089.1:3870001-3875000 | NW_006501089.1:3867963 | 0.547789979 | 1 | 0.753437655 | Y | Y |
| Zdhhc13 | Zdhhc13 | NW_006502473.1:77501-82500 | NW_006502473.1:103218 | 0.16024751 | 1 | 0.454665052 | Y | Y |

| | | | | | | | | |
|---|---|---|---|---|---|---|---|---|
| Zfpm2 | Zfpm2 | NW_006501143.1:4705001-4710000 | NW_006501143.1:4475014 | 0.890188802 | 1 | 0.724507939 | Y | |
| Zhx2 | Zhx2 | NW_006501508.1:2290001-2295000 | NW_006501508.1:2310667 | 0.302098058 | 1 | 0.689244731 | Y | Y |
| Zmat3 | Zmat3 | NW_006501046.1:4425001-4430000 | NW_006501046.1:4431199 | 0.498246172 | 1 | 0.765959424 | Y | Y |
| Znf202 | Zfp202 | NW_006501191.1:157501-162500 | NW_006501191.1:158918 | 0.224790018 | 1 | 0.797955297 | Y | Y |
| Znf24 | Zfp24 | NW_006501173.1:3707501-3712500 | NW_006501173.1:3712047 | 0.130003347 | 0.99976472 | 0.551992919 | Y | Y |
| Znf317 | Zfp317 | NW_006501353.1:1682501-1687500 | NW_006501353.1:1681344 | 0.173104072 | 1 | 0.780760564 | Y | Y |
| Znf318 | Zfp318 | NW_006501519.1:67501-72500 | NW_006501519.1:86801 | 0.168715008 | 1 | 0.711441689 | Y | Y |
| Znf436 | Zfp46 | NW_006501184.1:352501-357500 | NW_006501184.1:365773 | 0.119701627 | 0.999352979 | 0.335576005 | Y | Y |
| Znf451 | Zfp451 | NW_006501695.1:107501-112500 | NW_006501695.1:110330 | 0.592989237 | 1 | 0.440343449 | Y | Y |
| Znf512 | Zfp512 | NW_006501275.1:992501-997500 | NW_006501275.1:972394 | 0.277071823 | 1 | 0.411930462 | Y | Y |
| Znf518b | Zfp518b | NW_006502417.1:560001-565000 | NW_006502417.1:568026 | 0.166811309 | 1 | 0.428362074 | Y | Y |
| Znf521 | Zfp521 | NW_006501225.1:3900001-3905000 | NW_006501225.1:3874481 | 0.111977135 | 0.999058879 | 0.610890201 | Y | Y |
| Znf638 | Zfp638 | NW_006501567.1:367501-372500 | NW_006501567.1:345926 | 0.774193454 | 1 | 0.70096813 | Y | Y |
| Znf706 | Zfp706 | NW_006501220.1:1085001-1090000 | NW_006501220.1:1083386 | 0.181100313 | 1 | 0.719282913 | Y | Y |
| Znf831 | Zfp831 | NW_006501107.1:1585001-1590000 | NW_006501107.1:1527397 | 0.312387998 | 1 | 0.782492114 | | |
| Znfx1 | Znfx1 | NW_006501708.1:275001-280000 | NW_006501708.1:284143 | 1.274860735 | 1 | 0.625369087 | Y | Y |
| Zscan30 | Zscan30 | NW_006501173.1:3657501-3662500 | NW_006501173.1:3653759 | 0.120502274 | 0.999470619 | 0.594913734 | | |
| Zw10 | Zw10 | NW_006501163.1:127501-132500 | NW_006501163.1:123819 | 0.809071372 | 1 | 0.307782725 | Y | Y |

**Table S14** Inversions limits and haplotype frequencies across populations of interest. Data from (12).

| Chr. | inversion | left_breakpoint | right_breakpoint | pAnc_highland MountEvansCO | pAnc_lowland1 LincolnNE | pAnc_lowland2 MercedCA |
|---|---|---|---|---|---|---|
| 6 | inv6.0 | 0 | 43800000 | 0.086956522 | 0.84285714 | 1 |
| 7 | inv7.0 | 0 | 43400000 | 0.086956522 | 0.98571429 | 0.96666667 |
| 18 | inv18.0 | 34243699 | 45379251 | 0.989130435 | 0 | 0.1 |
| 22 | inv22.0 | 48636744 | 54689292 | 0.065217391 | 1 | 0.9 |
| 19 | inv19.0 | 1.00E+06 | 26700000 | 0.369565217 | 1 | 0.96666667 |
| 21 | inv21.0 | 39500000 | 70200000 | 0.391304348 | 0.91428571 | 0.93333333 |

**Table S15** Hypergeometric tests for selection target enrichment within inversions.

| Inversion | Selection_Targets (gene count) | All genes | P.value |
|---|---|---|---|
| inv18.0 | 51 | 112 | 9.21E-46 |
| inv19.0 | 40 | 123 | 2.41E-29 |
| inv21.0 | 35 | 17 | 3.44E-19 |
| inv22.0 | 3 | 116 | 0.0061997 |
| inv6.0 | 76 | 274 | 5.14E-78 |
| inv7.3 | 9 | 41 | 1.05E-05 |
| Genome-wide | 627 | 14345 | N/A |

**Table S16** GO enrichment results for top 1 percent of genes in weighted ranking in the LZ

| p_value | Term size | Query size | Intersection size | precision | recall | term_id | source | term_name |
|---|---|---|---|---|---|---|---|---|
| 0.01591697 | 93 | 135 | 8 | 0.05925926 | 0.08602151 | GO:0032963 | GO:BP | collagen metabolic process |
| 0.00406983 | 1199 | 135 | 29 | 0.21481481 | 0.02445194 | GO:0005576 | GO:CC | extracellular region |
| 0.00053226 | 280 | 135 | 14 | 0.1037037 | 0.04912281 | GO:0004175 | GO:MF | endopeptidase activity |
| 0.0036994 | 426 | 135 | 16 | 0.11851852 | 0.03712297 | GO:0008233 | GO:MF | peptidase activity |
| 0.00894361 | 81 | 135 | 7 | 0.05185185 | 0.08641975 | GO:0004222 | GO:MF | metalloendopeptidase activity |
| 0.02922175 | 67 | 135 | 6 | 0.04444444 | 0.08955224 | GO:0005518 | GO:MF | collagen binding |
| 0.03702717 | 136 | 135 | 8 | 0.05925926 | 0.05839416 | GO:0008237 | GO:MF | metallopeptidase activity |
| 0.0475289 | 2 | 135 | 2 | 0.01481481 | 1 | GO:0004687 | GO:MF | myosin light chain kinase activity |
| 0.0022846 | 71 | 135 | 7 | 0.05185185 | 0.09859155 | REAC:R-MMU-1474228 | REAC | Degradation of the extracellular matrix |
| 0.0047494 | 180 | 135 | 10 | 0.07407407 | 0.05555556 | REAC:R-MMU-1474244 | REAC | Extracellular matrix organization |
| 0.03268557 | 47 | 135 | 5 | 0.03703704 | 0.10638298 | REAC:R-MMU-375276 | REAC | Peptide ligand-binding receptors |
| 0.00081494 | 2714 | 135 | 54 | 0.4 | 0.01989683 | TF:M04632 | TF | Factor: SREBP-1; motif: NNNGTGGGGTGAN |
| 0.00453347 | 3760 | 135 | 65 | 0.48148148 | 0.01728264 | TF:M01168 | TF | Factor: SREBP; motif: NNNNYCACNCCANNN |
| 0.00559967 | 2564 | 135 | 50 | 0.37037037 | 0.01950078 | TF:M01173 | TF | Factor: SREBP-1; motif: NSNNTCACNCCANNN |
| 0.00135015 | 18 | 135 | 4 | 0.02962963 | 0.22222222 | WP:WP441 | WP | Matrix metalloproteinases |

**Table S17** Total read counts and reads mapped to features across all samples used in this study.

| Tissue | SampleID | Reads_Mapped_To_Features (millions) | Total_Reads (millions) | Strain | Trt | maternalID |
|---|---|---|---|---|---|---|
| JZ/DEC. | JZ002 | 37.89 | 65.77 | ME | 1N | ME15 |
| JZ/DEC. | JZ003 | 28.87 | 49.27 | ME | 1N | ME15 |
| JZ/DEC. | JZ006 | 31.50 | 52.88 | ME | 2H | ME22 |
| JZ/DEC. | JZ007 | 34.22 | 56.90 | ME | 2H | ME22 |
| JZ/DEC. | JZ009 | 27.03 | 45.54 | ME | 2H | ME24 |
| JZ/DEC. | JZ010 | 21.38 | 35.74 | ME | 2H | ME24 |
| JZ/DEC. | JZ013 | 35.31 | 59.93 | ME | 1N | ME5 |
| JZ/DEC. | JZ015 | 28.38 | 48.10 | ME | 1N | ME5 |
| JZ/DEC. | JZ021 | 24.22 | 40.59 | ME | 2H | ME10 |
| JZ/DEC. | JZ023 | 21.95 | 36.17 | ME | 2H | ME10 |
| JZ/DEC. | JZ025 | 34.78 | 58.17 | ME | 1N | ME16 |
| JZ/DEC. | JZ026 | 36.28 | 61.45 | ME | 1N | ME16 |
| JZ/DEC. | JZ030 | 20.69 | 34.63 | ME | 1N | ME23 |
| JZ/DEC. | JZ032 | 30.60 | 51.67 | ME | 1N | ME23 |
| JZ/DEC. | JZ039 | 32.00 | 58.36 | BW | 1N | BW2 |
| JZ/DEC. | JZ040 | 33.62 | 62.16 | BW | 1N | BW2 |
| JZ/DEC. | JZ041 | 26.80 | 48.30 | BW | 1N | BW2 |
| JZ/DEC. | JZ047 | 25.77 | 43.52 | BW | 2H | BW16 |
| JZ/DEC. | JZ048 | 21.67 | 37.48 | BW | 2H | BW16 |
| JZ/DEC. | JZ049 | 22.73 | 39.40 | BW | 2H | BW16 |
| JZ/DEC. | JZ051 | 26.82 | 47.16 | BW | 2H | BW3 |
| JZ/DEC. | JZ052 | 32.45 | 56.98 | BW | 2H | BW3 |
| JZ/DEC. | JZ060 | 24.11 | 39.76 | ME | 1N | ME30 |
| JZ/DEC. | JZ061 | 23.20 | 37.93 | ME | 1N | ME30 |
| JZ/DEC. | JZ063 | 31.00 | 53.34 | BW | 2H | BW15 |
| JZ/DEC. | JZ064 | 32.63 | 55.24 | BW | 2H | BW15 |
| JZ/DEC. | JZ066 | 33.70 | 56.99 | ME | 2H | ME11 |
| JZ/DEC. | JZ068 | 30.97 | 53.06 | ME | 2H | ME11 |
| JZ/DEC. | JZ070 | 34.42 | 57.60 | BW | 2H | BW12 |
| JZ/DEC. | JZ071 | 16.26 | 27.09 | BW | 2H | BW12 |
| JZ/DEC. | JZ072 | 31.72 | 52.50 | ME | 1N | ME39 |
| JZ/DEC. | JZ074 | 30.23 | 50.51 | ME | 1N | ME39 |
| JZ/DEC. | JZ075 | 39.04 | 62.06 | ME | 2H | ME43 |
| JZ/DEC. | JZ078 | 32.34 | 50.97 | ME | 2H | ME43 |
| JZ/DEC. | JZ081 | 23.96 | 41.47 | ME | 1N | ME40 |
| JZ/DEC. | JZ082 | 27.61 | 46.65 | ME | 1N | ME40 |

| | | | | | | |
|---|---|---|---|---|---|---|
| JZ/DEC. | JZ087 | 21.26 | 36.22 | BW | 2H | BW21 |
| JZ/DEC. | JZ088 | 23.80 | 40.30 | BW | 2H | BW21 |
| JZ/DEC. | JZ092 | 31.37 | 51.29 | BW | 2H | BW32 |
| JZ/DEC. | JZ093 | 24.63 | 40.94 | BW | 2H | BW32 |
| JZ/DEC. | JZ094 | 34.40 | 59.37 | BW | 2H | BW28 |
| JZ/DEC. | JZ095 | 30.16 | 51.09 | BW | 2H | BW28 |
| JZ/DEC. | JZ096 | 30.98 | 52.70 | BW | 2H | BW28 |
| JZ/DEC. | JZ097 | 35.31 | 59.12 | BW | 2H | BW28 |
| JZ/DEC. | JZ098 | 24.39 | 43.83 | BW | 1N | BW26 |
| JZ/DEC. | JZ099 | 34.28 | 62.51 | BW | 1N | BW26 |
| JZ/DEC. | JZ101 | 33.26 | 57.34 | BW | 2H | BW34 |
| JZ/DEC. | JZ103 | 31.26 | 52.44 | BW | 2H | BW36 |
| JZ/DEC. | JZ104 | 28.18 | 50.98 | BW | 1N | BW39 |
| JZ/DEC. | JZ106 | 31.73 | 57.50 | BW | 1N | BW25 |
| JZ/DEC. | JZ108 | 27.06 | 49.37 | BW | 1N | BW45 |
| JZ/DEC. | JZ109 | 31.12 | 55.39 | BW | 1N | BW45 |
| JZ/DEC. | JZ110 | 29.68 | 53.50 | BW | 1N | BW45 |
| JZ/DEC. | JZ111 | 31.36 | 56.07 | BW | 1N | BW43 |
| JZ/DEC. | JZ112 | 27.30 | 49.25 | BW | 1N | BW43 |
| JZ/DEC. | JZ114 | 30.12 | 52.91 | BW | 1N | BW38 |
| JZ/DEC. | JZ116 | 26.98 | 48.57 | BW | 1N | BW38 |
| JZ/DEC. | JZ117 | 27.43 | 49.84 | BW | 1N | BW25 |
| JZ/DEC. | JZ121 | 30.91 | 55.66 | BW | 1N | BW38 |
| JZ/DEC. | JZ122 | 21.57 | 38.39 | BW | 1N | BW41 |
| JZ/DEC. | JZ123 | 34.54 | 59.89 | BW | 1N | BW41 |
| JZ/DEC. | JZ124 | 28.40 | 48.70 | BW | 1N | BW41 |
| JZ/DEC. | JZ129 | 35.62 | 60.88 | ME | 2H | ME203 |
| JZ/DEC. | JZ130 | 27.09 | 47.64 | ME | 2H | ME203 |
| JZ/DEC. | JZ131 | 20.18 | 35.06 | ME | 2H | ME203 |
| JZ/DEC. | JZ132 | 31.05 | 53.36 | ME | 2H | ME203 |
| JZ/DEC. | JZ134 | 31.02 | 55.19 | ME | 1N | ME202 |
| JZ/DEC. | JZ135 | 21.06 | 36.14 | ME | 1N | ME202 |
| JZ/DEC. | JZ138 | 30.48 | 52.51 | ME | 1N | ME208 |
| JZ/DEC. | JZ142 | 27.92 | 47.93 | ME | 1N | ME208 |
| JZ/DEC. | JZ146 | 28.81 | 51.06 | ME | 1N | ME232 |
| JZ/DEC. | JZ147 | 33.82 | 60.08 | ME | 1N | ME232 |
| JZ/DEC. | JZ166 | 39.31 | 67.10 | ME | 2H | ME243 |
| JZ/DEC. | JZ167 | 31.38 | 53.84 | ME | 2H | ME243 |
| JZ/DEC. | JZ168 | 37.21 | 64.36 | ME | 2H | ME236 |

| | | | | | | |
|---|---|---|---|---|---|---|
| JZ/DEC. | JZ169 | 32.25 | 54.89 | ME | 2H | ME236 |
| JZ/DEC. | JZ211 | 31.55 | 53.05 | ME | 2H | ME282 |
| JZ/DEC. | JZ214 | 33.06 | 56.55 | ME | 2H | ME282 |
| LZ | LZ002 | 39.30 | 62.94 | ME | 1N | ME15 |
| LZ | LZ003 | 32.29 | 54.06 | ME | 1N | ME15 |
| LZ | LZ006 | 27.28 | 47.87 | ME | 2H | ME22 |
| LZ | LZ007 | 27.96 | 48.79 | ME | 2H | ME22 |
| LZ | LZ009 | 29.98 | 53.27 | ME | 2H | ME24 |
| LZ | LZ010 | 34.35 | 60.72 | ME | 2H | ME24 |
| LZ | LZ013 | 33.14 | 55.36 | ME | 1N | ME5 |
| LZ | LZ015 | 27.73 | 47.13 | ME | 1N | ME5 |
| LZ | LZ021 | 30.14 | 53.80 | ME | 2H | ME10 |
| LZ | LZ023 | 29.61 | 53.23 | ME | 2H | ME10 |
| LZ | LZ025 | 36.75 | 61.95 | ME | 1N | ME16 |
| LZ | LZ026 | 36.71 | 63.00 | ME | 1N | ME16 |
| LZ | LZ030 | 30.37 | 52.29 | ME | 1N | ME23 |
| LZ | LZ032 | 39.23 | 66.89 | ME | 1N | ME23 |
| LZ | LZ039 | 36.20 | 62.17 | BW | 1N | BW2 |
| LZ | LZ040 | 26.78 | 46.28 | BW | 1N | BW2 |
| LZ | LZ041 | 29.40 | 50.75 | BW | 1N | BW2 |
| LZ | LZ047 | 31.79 | 52.02 | BW | 2H | BW16 |
| LZ | LZ048 | 47.45 | 78.78 | BW | 2H | BW16 |
| LZ | LZ049 | 29.19 | 47.96 | BW | 2H | BW16 |
| LZ | LZ051 | 36.74 | 60.38 | BW | 2H | BW3 |
| LZ | LZ052 | 32.21 | 56.72 | BW | 2H | BW3 |
| LZ | LZ060 | 31.49 | 55.74 | ME | 1N | ME30 |
| LZ | LZ061 | 28.19 | 49.29 | ME | 1N | ME30 |
| LZ | LZ063 | 35.75 | 60.14 | BW | 2H | BW15 |
| LZ | LZ064 | 27.49 | 46.26 | BW | 2H | BW15 |
| LZ | LZ066 | 24.55 | 44.69 | ME | 2H | ME11 |
| LZ | LZ068 | 30.95 | 56.99 | ME | 2H | ME11 |
| LZ | LZ070 | 29.21 | 50.36 | BW | 2H | BW12 |
| LZ | LZ071 | 31.44 | 52.65 | BW | 2H | BW12 |
| LZ | LZ072 | 28.29 | 49.29 | ME | 1N | ME39 |
| LZ | LZ074 | 32.29 | 57.50 | ME | 1N | ME39 |
| LZ | LZ075 | 26.27 | 46.24 | ME | 2H | ME43 |
| LZ | LZ078 | 27.05 | 49.93 | ME | 2H | ME43 |
| LZ | LZ081 | 24.45 | 43.11 | ME | 1N | ME40 |
| LZ | LZ082 | 29.30 | 52.49 | ME | 1N | ME40 |

| | | | | | | |
|---|---|---|---|---|---|---|
| LZ | LZ087 | 28.48 | 47.73 | BW | 2H | BW21 |
| LZ | LZ088 | 30.57 | 51.20 | BW | 2H | BW21 |
| LZ | LZ092 | 34.78 | 58.51 | BW | 2H | BW32 |
| LZ | LZ093 | 35.10 | 59.28 | BW | 2H | BW32 |
| LZ | LZ094 | 32.68 | 56.08 | BW | 2H | BW28 |
| LZ | LZ095 | 30.91 | 52.57 | BW | 2H | BW28 |
| LZ | LZ096 | 28.83 | 47.69 | BW | 2H | BW28 |
| LZ | LZ097 | 30.96 | 50.41 | BW | 2H | BW28 |
| LZ | LZ098 | 27.38 | 48.03 | BW | 1N | BW26 |
| LZ | LZ099 | 34.33 | 57.87 | BW | 1N | BW26 |
| LZ | LZ101 | 29.89 | 49.49 | BW | 2H | BW34 |
| LZ | LZ103 | 31.98 | 53.89 | BW | 2H | BW36 |
| LZ | LZ104 | 27.78 | 47.54 | BW | 1N | BW39 |
| LZ | LZ106 | 35.30 | 61.10 | BW | 1N | BW25 |
| LZ | LZ108 | 33.23 | 57.77 | BW | 1N | BW45 |
| LZ | LZ109 | 34.02 | 56.93 | BW | 1N | BW45 |
| LZ | LZ110 | 33.55 | 56.52 | BW | 1N | BW45 |
| LZ | LZ111 | 25.80 | 41.60 | BW | 1N | BW43 |
| LZ | LZ112 | 29.58 | 49.07 | BW | 1N | BW43 |
| LZ | LZ114 | 31.86 | 53.04 | BW | 1N | BW38 |
| LZ | LZ116 | 36.48 | 58.49 | BW | 1N | BW38 |
| LZ | LZ117 | 29.23 | 49.17 | BW | 1N | BW25 |
| LZ | LZ121 | 37.24 | 61.03 | BW | 1N | BW38 |
| LZ | LZ122 | 36.33 | 61.67 | BW | 1N | BW41 |
| LZ | LZ123 | 37.96 | 62.80 | BW | 1N | BW41 |
| LZ | LZ124 | 30.62 | 53.81 | BW | 1N | BW41 |
| LZ | LZ129 | 27.38 | 49.05 | ME | 2H | ME203 |
| LZ | LZ130 | 36.59 | 64.19 | ME | 2H | ME203 |
| LZ | LZ131 | 25.28 | 45.42 | ME | 2H | ME203 |
| LZ | LZ132 | 25.20 | 45.38 | ME | 2H | ME203 |
| LZ | LZ134 | 28.34 | 49.37 | ME | 1N | ME202 |
| LZ | LZ135 | 17.21 | 31.12 | ME | 1N | ME202 |
| LZ | LZ138 | 29.98 | 54.23 | ME | 1N | ME208 |
| LZ | LZ142 | 37.22 | 66.52 | ME | 1N | ME208 |
| LZ | LZ146 | 25.30 | 45.82 | ME | 1N | ME232 |
| LZ | LZ147 | 35.43 | 60.23 | ME | 1N | ME232 |
| LZ | LZ166 | 32.17 | 57.87 | ME | 2H | ME243 |
| LZ | LZ167 | 28.28 | 51.28 | ME | 2H | ME243 |
| LZ | LZ168 | 35.60 | 62.85 | ME | 2H | ME236 |

| | | | | | | |
|---|---|---|---|---|---|---|
| LZ | LZ169 | 29.23 | 51.52 | ME | 2H | ME236 |
| LZ | LZ211 | 31.62 | 55.43 | ME | 2H | ME282 |
| LZ | LZ214 | 32.71 | 56.88 | ME | 2H | ME282 |

**Legend for Dataset S1**
Individual phenotype data from pups, dams, and placental histology. Includes pup birthweights, in utero mass data, maternal physiological data, and placental histology data.

**Legend for Dataset S2**
Sequence data summary data. Includes annotation for placenta-specific genes in *Mus* added to the *Peromyscus maniculatus* annotation file and output tables for JZ/DEC. and LZ RNAseq data. RNAseq output data include columns with yes/no information on gene set membership for all gene sets discussed in the manuscript.
